# Marionette: *E. coli* containing 12 highly-optimized small molecule sensors

**DOI:** 10.1101/285866

**Authors:** Adam J. Meyer, Thomas H. Segall-Shapiro, Christopher A. Voigt

## Abstract

Cellular processes are carried out by many interacting genes and their study and optimization requires multiple levers by which they can be independently controlled. The most common method is via a genetically-encoded sensor that responds to a small molecule (an “inducible system”). However, these sensors are often suboptimal, exhibiting high background expression and low dynamic range. Further, using multiple sensors in one cell is limited by cross-talk and the taxing of cellular resources. Here, we have developed a directed evolution strategy to simultaneously select for less background, high dynamic range, increased sensitivity, and low crosstalk. Libraries of the regulatory protein and output promoter are built based on random and rationally-guided mutations. This is applied to generate a set of 12 high-performance sensors, which exhibit >100-fold induction with low background and cross-reactivity. These are combined to build a single “sensor array” and inserted into the genomes of *E. coli* MG1655 (wild-type), DH10B (cloning), and BL21 (protein expression). These “Marionette” strains allow for the independent control of gene expression using 2,4-diacetylphophloroglucinol (DAPG), cuminic acid (Cuma), 3-oxohexanoyl-homoserine lactone (OC6), vanillic acid (Van), isopropyl β-D-1-thiogalactopyranoside (IPTG), anhydrotetracycline (aTc), L-arabinose (Ara), choline chloride (Cho), naringenin (Nar), 3,4-dihydroxybenzoic acid (DHBA), sodium salicylate (Sal), and 3-hydroxytetradecanoyl-homoserine lactone (OHC14).

---

Advances in biology are often tied to new methods that use external stimuli to control the levels of gene expression^1–3^. Pioneered in the early 1980s, so-called inducible systems were developed that allow genes to be turned on by adding a small molecule inducer to the growth media^4^. These consist of a protein transcription factor (*e.g*., LacI) whose binding to a DNA operator in a promoter is controlled by the inducer (*e.g*., IPTG). Initially co-opted from natural regulatory networks, over the years many versions were designed to improve performance. In the 1990s, additional systems were developed that responded to other inducers, notably arabinose and aTc, which became common tools in the field. In 1997, Lutz and Bujard published a seminal paper that combined three (IPTG, arabinose, aTc) that could be easily interchanged on a two-plasmid system^5^. Its organizational simplicity, compatibility, and quantified response functions were revolutionary. Beyond providing a new tool to biologists to control multiple genes with independent “strings,” it facilitated researchers with quantitative backgrounds to enter biology^6–7^. Armed with the new ability to control two genes with precision, physicists and engineers built the first synthetic genetic circuits, performed single molecule experiments inside cells, deconstructed the origins of noise in gene expression, determined how enzyme balancing impacts metabolic flux, elucidated rules underlying the assembly of molecular machines, and built synthetic symbiotic microbial communities, just to highlight a few^8–19^.

Sensor performance is quantified by its response function; in other words, how the concentration of inducer changes the activity of the output promoter (Figure 1a). Often, this promoter retains a residual activity in the absence of inducer (“leakiness”). This hampers the ability to explore low expression levels or keep a gene in the off state, particularly needed for toxic proteins^20–21^. Another important parameter is the dynamic range, defined as the ratio of promoter activity in the on and off states. When this is large, both high and low expression can be explored as well as many intermediate states. The sensitivity is the concentration of inducer that turns a sensor on (defined as 50% activation). A lower sensitivity reduces the amount of a chemical that must be added to the media. Further, when multiple sensors are combined into one cell, they can interfere with each other’s response functions (Figure 1b). Some small molecules bind non-cognate regulators and this can lead to off-target activation (cross reactivity) or competitive inhibition with the cognate small molecule (antagonism)^22–27^. Finally, each sensor requires cellular resources (*e.g*., ribosomes) to function and the activation of one sensor can influence another indirectly due to resource competition^28^. Each sensor also requires ~1-2 kb of DNA and this becomes increasingly difficult to carry on plasmids. These challenges limit the number of sensors that can be put in a single cell and the maximum reported to date is four^29^.

**Figure 1:**
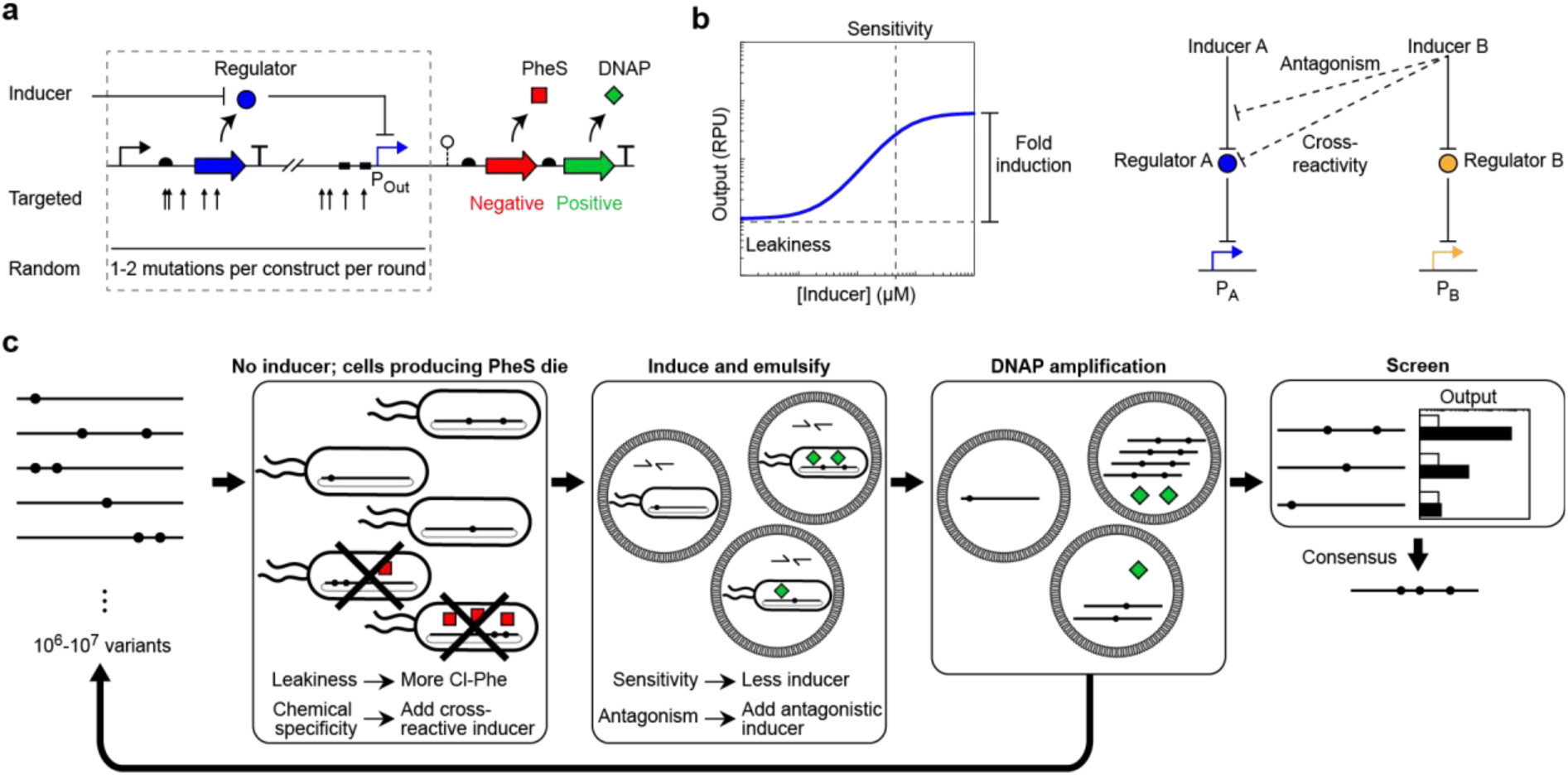
A dual selection for sensor optimization. **a)** A regulator (blue circle) is expressed from a weak constitutive promoter and controls expression from the output promoter (P_Out_) while itself being controlled by an externally applied inducer molecule. Transcription from P_Out_ determines the expression level of an aminoacyl-tRNA synthetase (PheS; red square) and a DNA polymerase (KOD or PK6; green diamond). Initial libraries may contain targeted degeneracy in the regulator RBS, regulator CDS, or P_Out_ (arrows), and random mutations are added throughout the entire library during each round of selection. **b)** A response function captures the activity of P_Out_ at various levels of inducer. Regulator A affects promoter A, and is itself affected by inducer A. Inducer B may affect Regulator A (chemical cross-reactivty) or interfere with inducer A’s action (antagonism). **c)** The dual selection schema is shown (see text for details). Dots denote mutations; half arrows represent PCR primers.

Sensors can be improved using biophysical models, rational engineering and directed evolution^5, 23, 26, 30–36^. Improving performance requires that screens or selections be performed in the presence and absence of inducer^37^. Such dual selections have been accomplished by sorting cells based low and high fluorescence, utilizing proteins that can be both toxic and selective (*e.g.,* TetA or HSV-TK), or by deploying separate positive and negative selections^26, 35, 37–44^. This has been applied to improving the response functions and eliminating cross-reactivity between pairs of regulators^23, 26, 35, 45^.

Here, we have developed a selection methodology that allows us to intervene at multiple steps in order to simultaneously select for improved response functions and decreased crosstalk (Figure 1c). First, we built 12 sensors that respond to different small molecules and made directed changes in order to improve function. These sensors were then subjected to multiple rounds of negative-positive selection. Negative selection is facilitated by a promiscuous A294G mutant of the phenylalanine aminoacyl tRNA-synthetase (PheS)^46^. In the absence of inducer, leaky transcription of PheS leads to the charging of phenylalanyl-tRNA with the non-canonical amino acid 4-Chloro-DL-phenylalanine (Cl-Phe). Adding more Cl-Phe increases the stringency of the selection by making PheS transcription more toxic^43, 47^. Positive selection is facilitated by a thermostable DNA polymerase (DNAP), either KOD DNAP from the archaea *Thermococcus kodakarensis*^48^ (for stringent replication) or the engineered PK6 DNAP^49^ (to introduce mutations). In the presence of inducer, the level of DNAP can be quantified by PCR amplification of the sensor library after the cells are emulsified with primers^50–51^. Multiple properties of the response function can be simultaneously improved during one cycle of negative-positive selection and stringency altered by changing the concentrations of Cl-Phe and inducer. Further, cross reactions can be selected against by adding the inducer or regulator from a problematic system. The dual selection is applied over multiple rounds to create a highly-optimized set of sensors. From these, we identify 12 that can be used together in a single strain, and these are combined and integrated into the genomes of *E. coli* MG1655, DH10B, and BL21 in order to create the Marionette family of strains. Genome integration increases stability and reduces the cellular resources required to maintain regulator expression. It also simplifies the use of the inducible systems, where only the output promoters need to be incorporated into a design (*e.g*., for the expression of multiple proteins from a plasmid).

An initial set of 17 putative sensors were designed. Each sensor consists of a weak constitutive promoter (P_LacI_, Supplementary Table 1) driving the expression of the regulatory gene and an output promoter that is acted on by the regulator. The regulatory genes were either codon optimized and synthesized or cloned (Methods and Supplementary Table 4). The output promoters were either obtained from the literature or, in the case of P_Van_, rationally designed by inserting cognate operator sequences into unregulated promoters (Supplementary Table 1). The activity of the output promoter was measured through the expression of yellow fluorescent protein (YFP) using flow cytometry and reported in relative promoter units (RPUs) (Supplementary Figure 1 and Methods). Each complete sensor was cloned into the same sensor plasmid architecture (Supplementary Figure 2). Some sensors require additional genes, which are encoded as an operon with the regulatory protein. For the Ara-inducible system, the transporter *araE* was included in order to produce a graded response^52^. For the Ery-inducible system, the ribosome methylase *ery*^*R*^ was included to confer resistance to Ery^53^.

Rational mutations were made to improve some sensors prior to performing the direction evolution experiments. Multiple versions of each sensor, each with different a promoter and RBS used to drive the expression of the regulator, were tested (Supplementary Figure 3). The version with the largest dynamic range was chosen for further optimization. Then, a number of potential improvements gleaned from the literature or rationally designed were evaluated, the results of which are shown in Supplementary Figure 4. After this step, we reduced the set of sensors to 14, removing copper, glucaric acid and paraquat inducible systems because their responses were too small for subsequent optimization. The response functions of the initial sensors are shown in Figure 2 (grey curves). Each function was obtained by fitting the experimental data to the equation

**Figure 2:**
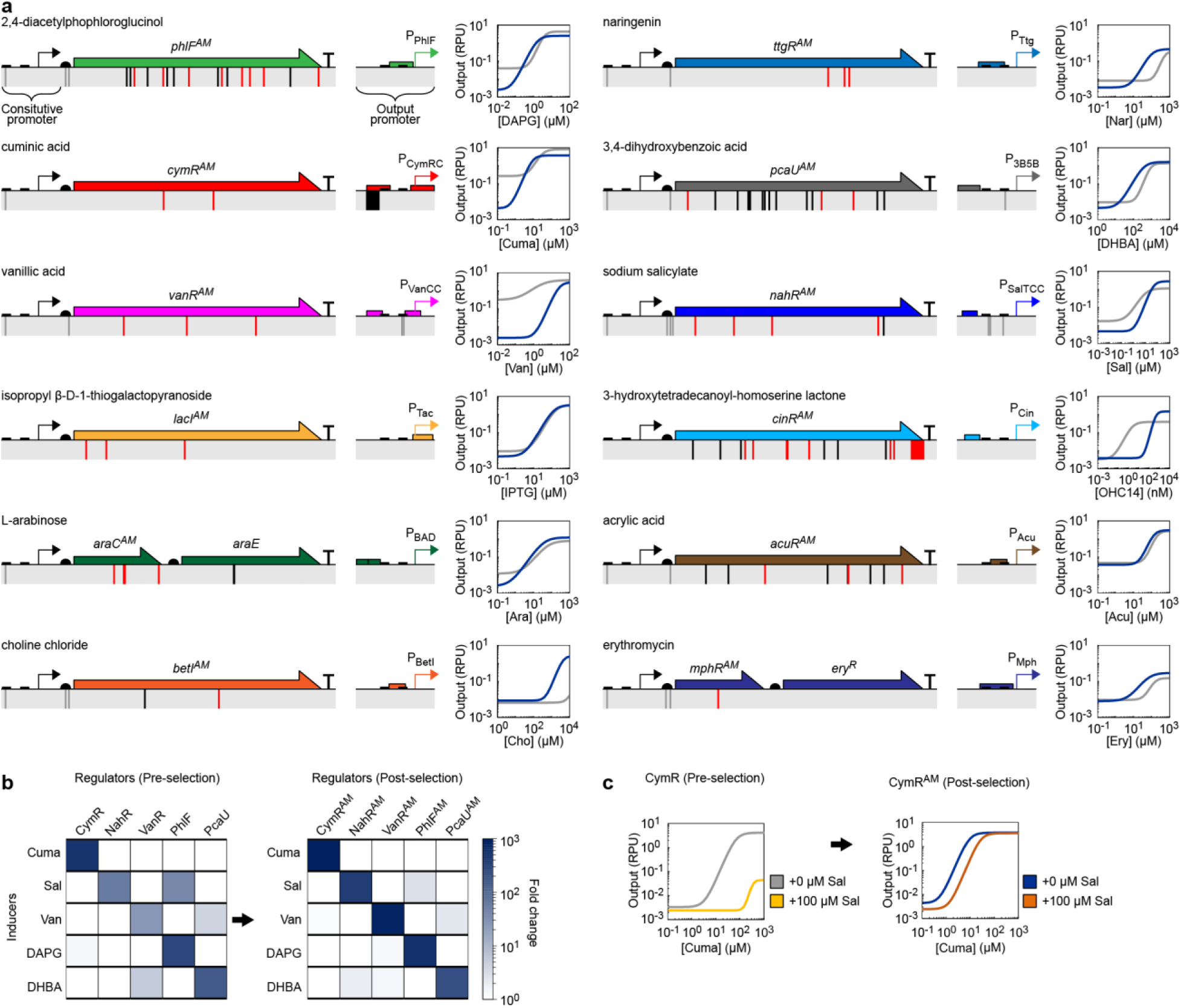
Improved sensor performance. **a)** Each genetically encoded sensor is shown, with coding (red) and non-coding (grey/black) mutations noted. Mutations in grey were also applied to the parental sensor. The corresponding response functions comparing the evolved (blue) and parental (grey) sensors. The fit of Equation 1 to the mean of three replicates performed on different days is shown (see Supplementary Figure 5 for data). The response function parameters for evolved sensors are provided in Table 1. **b)** Chemical cross-reactivity heat-map of reference (left) and evolved (right) sensors. Inducer concentrations were: 100 μM Cuma, 100 μM Sal, 100 μM Van, 10 μM DAPG, and 1 mM DHBA. The mean of three replicates performed on different days is shown (see Supplementary Figure 8 for data). **c)** Response function with Sal (light/dark orange) and without Sal (grey/blue) for parental CymR (left) and evolved CymR^AM^ (right). The fit of Equation 1 to the mean of three replicates from different days is shown (see Supplementary Figure 7 for data). Sequences of promoters and regulators are provided in Supplementary Table 1 and 4.

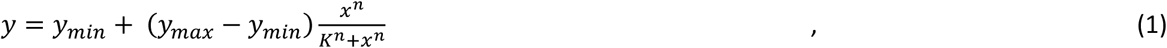

where *y* is the promoter activity in RPU, *x* is the concentration of the small molecule, *y*_*min*_ is the leakiness, *y*_*max*_/*y*_*min*_ is the dynamic range, *K* is the sensitivity, and *n* is the cooperativity. The raw data points, including error bars, used for this fit are shown in Supplementary Figure 5. While they all show some response, the high leakiness, low dynamic range and low sensitivity are apparent for many. Of the 14, the aTc- and OC6-sensors produced a good enough response to not require additional optimization (Table 1).

**Table 1.**
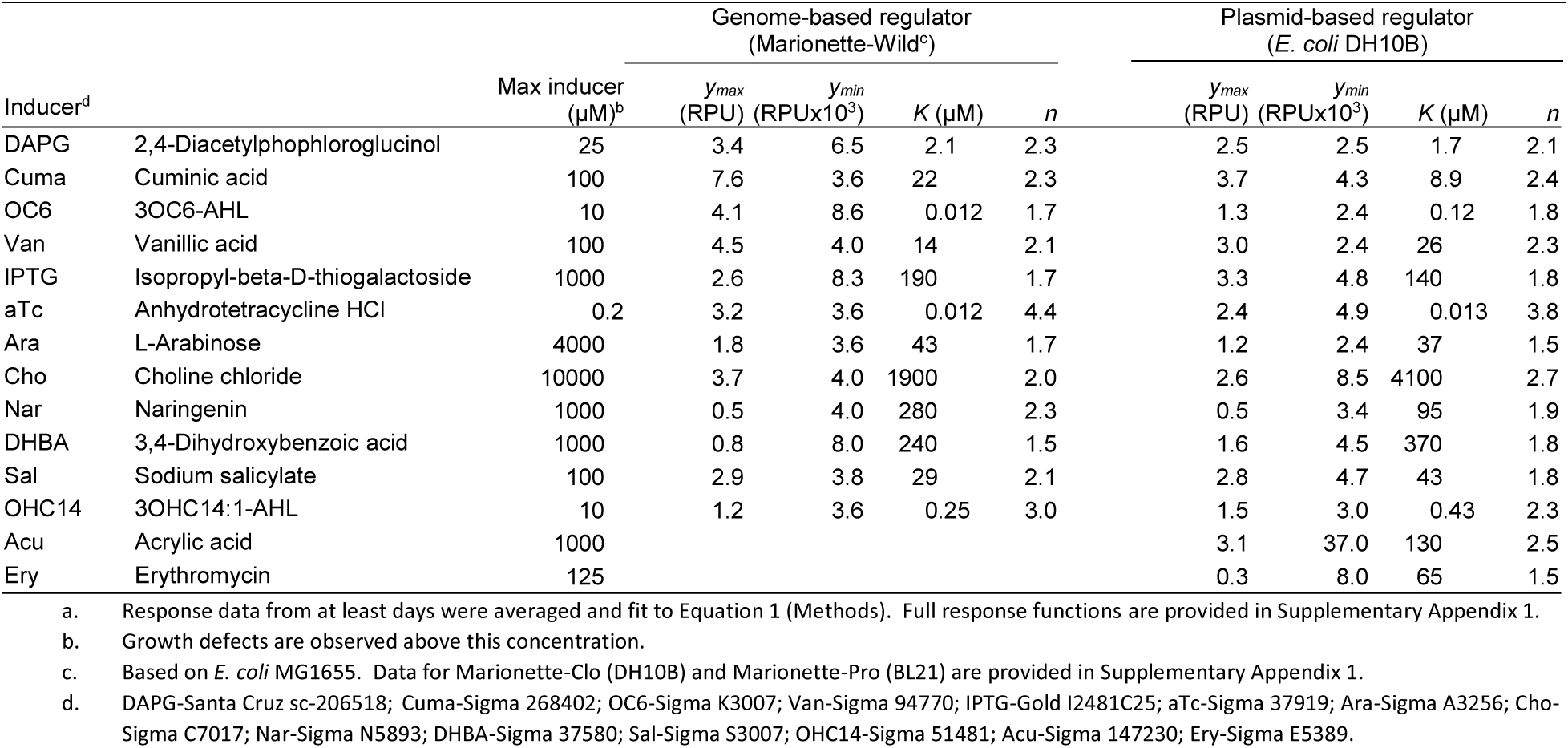
Sensor response function parameters^a^.

The remaining 12 were then subjected to directed evolution using the dual selection (Figure 1c, Supplementary Figure 6). For each sensor, a library was constructed and cloned upstream of the operon containing PheS and DNAP (Figure 1a) on a dual selection plasmid. The initial library contained a mixture of random and rational mutations. For a subset of sensors, the output promoter was mutagenized: selected bases in the −10 box and −35 box were randomized. To control regulator expression, critical bases in the ribosome binding site (RBS) were randomized. For LacI and AraC, we partially mutagenized key amino acid residues based on prior work^23, 33–34^. Specifically, the genes were PCR amplified with primers of limited degeneracy (*e.g.,* VNA, WKK, or NDC) thus allowing LacI Q18, F161, W220, Q291, and L296 and AraC L133, E165, E169, and C280 to sample a subset of possible amino acids (Methods). The specific mutations made to the initial library for each sensor are shown in Supplementary Appendix 2.

Multiple rounds of the dual selection were performed with the initial library. Between 4 and 23 rounds were performed, depending on how many issues needed to be corrected for each sensor. Different interventions were performed during each round to bias solutions to address problems identified for each sensor. The conditions for each round are presented in detail in Supplementary Appendix 2. Typically, the stringency of the negative selection was increased at a particular round by increasing the concentration of Cl-Phe from 2 mM to 4 mM. This biases against leakiness in the absence of inducer. During positive selection, inducer was added to the surviving cells, leading to the expression of DNAP. During early rounds, the maximum amount of inducer was added. For some sensors, we sought to increase the sensitivity of the response by reducing the amount of inducer after each round; for example, BetI was induced with 1 mM Cho in the final round of selection, down from 5 mM Cho in the first round. After induction, the cells were encapsulated, lysed, and PCR amplified using the DNAP expressed by the sensor (Methods). In early rounds, additional random mutations throughout the sensor were introduced by using the PK6 DNAP during the positive selection (yielding an average of 1-2 mutations per kilobase). In later rounds, the stringent KOD DNAP was used to reduce the diversity in the population. After the amplification step of the positive selection, the constructs were recloned into the selection plasmid. Recloning allows the selection plasmid to be reset each round, thus preventing the accumulation of cheaters, and offers the opportunity to change the DNAP as needed. In some cases, in an effort to combine multiple beneficial mutations into a single variant, gene shuffling^54^ was used between rounds (Methods).

During the selection rounds, additional interventions were included in order to reduce crosstalk between systems. For example, crosstalk between the IPTG- and Ara-sensors is well known, where IPTG reduces the output of the AraC/P_BAD_ system^23^ (Supplementary Figure 7). To identify mutants that reduce this, 1000 μM IPTG was included during all rounds of selection for the Ara sensor. IPTG was also included in the negative selection to prevent the evolution of an IPTG-induced AraC mutant. Crosstalk between Van and DHBA was also observed and eliminated through the negative selection (Figure 2b, Supplementary Figure 8). Salycilate was also found to antagonize the Cuma sensor (Figure 2c, Supplementary Figure 7) and cross react with the DAPG sensor (Figure 2b, Supplementary Figure 8) and was thus added to both the positive and negative selections.

After all of the rounds of selection are complete, the library was assembled into a YFP screening plasmid (Supplementary Figure 6). Several clones were picked and assayed for output expression in the presence and absence of inducer (Supplementary Appendix 2). In addition, when selecting against crosstalk, the clones were screened for induction by these molecules. The mutants showing highest improvement (leakiness, sensitivity, dynamic range, and orthogonality) were identified and sequenced (Supplementary Appendix 2). Based on the mutations observed, one or more consensus sequences were constructed and re-screened to identify the best variant. The final sequences of the evolved sensors, including the mutations identified are provided in Supplementary Table 4. Mutations were identified throughout the sensors, including the promoter/RBS controlling regulator expression, synonymous and non-synonymous mutations throughout the regulator genes, and mutations/substitutions in the output promoters (Figure 2a). On average, about eight cumulative mutations were made to each sensor as a result of the rounds of dual selection.

The improvements in the response functions are shown in Figure 2a (blue curves). The fit parameters to Equation 1 are provide in Table 1 (raw data are provided in Supplementary Figure 5). There is marked improvement in many of the response functions, sometimes showing orders of magnitude changes in the leakiness, dynamic range, and sensitivity. The Ery- and Acu-sensors showed slight improvements in their response functions, but were not chosen to be part of the final set because of their low dynamic range.

CymR, NahR, VanR, PhlF, and PcaU each responds to a substituted benzene. Therefore, we examined the activity of this set of sensors against all five inducers. The optimized sensors showed significant reduction in the crosstalk while maintaining high activity with their cognate inducer (Figure 2b, Supplementary Figure 8). Improvements in the antagonism between CymR and Sal were also tested (Figure 2c, Supplementary Figure 7). In the presence of 100 μM Sal, the ability for Cuma to induce its sensor drops by 1200-fold. The sensor obtained by rounds of positive selection in the presence of Cuma reduces this by two orders of magnitude. There is also slight antagonism of AraC by IPTG, which also improved as a result of the selection (Supplementary Figure 7). Collectively, these improvements allow for all of these sensors to be used simultaneously in a single cell.

The best 12 sensors were then combined to form a “sensor array” that was inserted into the genome of *E. coli* MG1655 to create “Marionette-Wild” (Figure 3 and Supplementary Table 6). Genomic insertion has several benefits: it stabilizes the cluster and simplifies the use of multiple systems without building large plasmids containing the regulators. The array consists of the 12 regulatory genes and *araE* transporter organized into several operons (Figure 3a and Supplementary Figure 9). The genes were organized into three operons controlled by three medium-strength constitutive promoters. Each gene is encoded by its own ribosome binding site (RBS), which was rationally designed using the RBS Calculator^55^ in order to achieve equivalent expression as when encoded on the plasmid (Methods and Supplementary Figure 10). Strong terminators were included before and after the sensor array in order to insulate the array from context effects. Phage transduction was used to move the sensor array to create two additional cell lines: the recA-deficient *E. coli* DH10B strain for cloning “Marionette-Clo” and the protease-deficient *E. coli* BL21 for protein expression “Marionette-Pro” (Methods).

**Figure 3:**
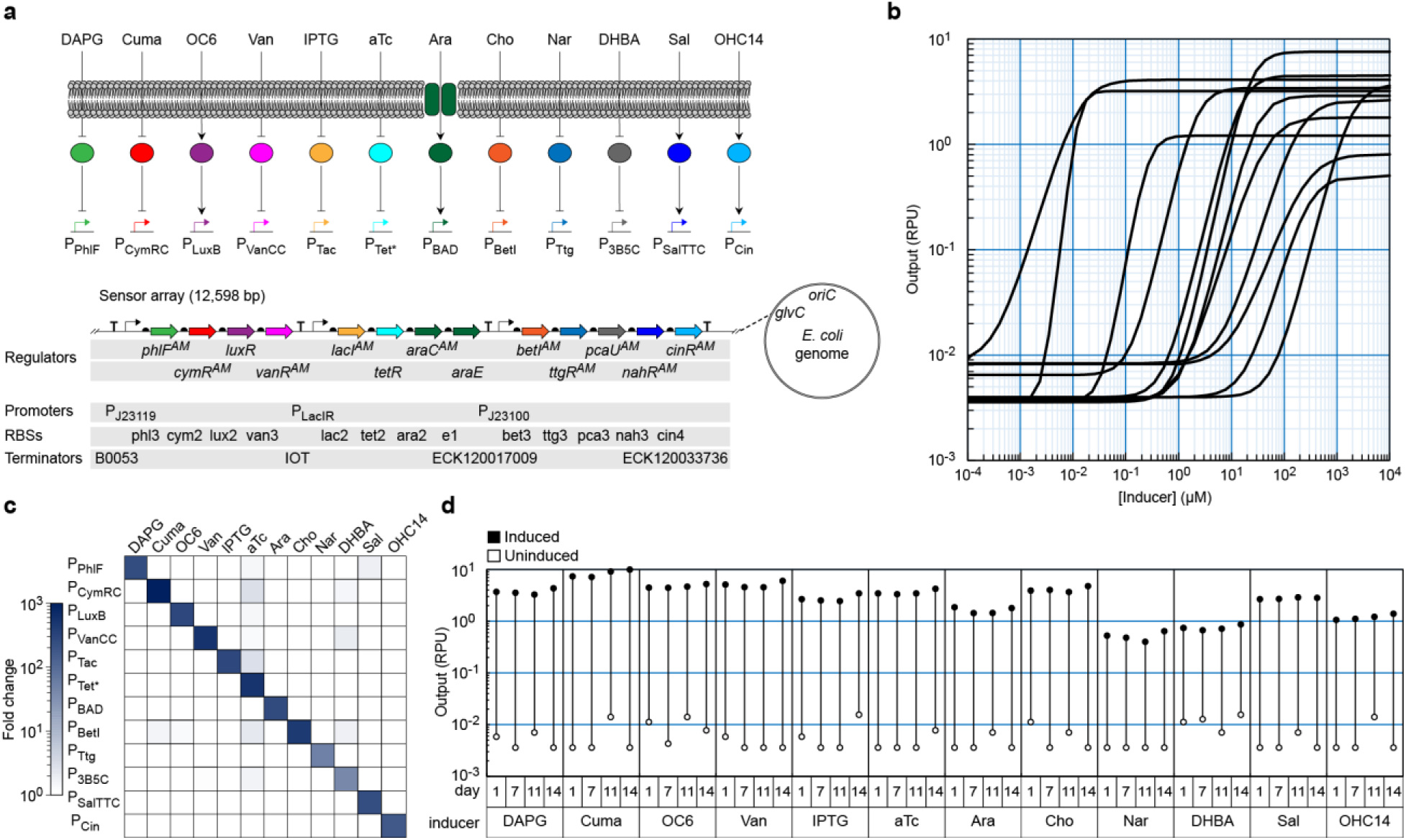
Marionette-Wild performance. **a)** The molecular and genetic schematic of the Marionette cluster and its location in the *E. coli* genome. The cluster was inserted in the direction of leading strand replication, between 3,860,010 and 3,861,627 in *E. coli* MG1655 (NCBI accession number NC_000913), between 3,957,594 and 3,959,211 in *E. coli* DH10B (NC_010473), and between 3,720,027 and 3,721,644 in *E. coli* BL21 (CP010816). **b)** Overlaid response functions for each of the 12 output promoters with their cognate inducers. The fit of Equation 1 to the mean of three replicates performed on different days is shown (see Supplementary Figure 11 for data). The response function parameters are provided in Table 1. **c)** Chemical cross-reactivity heat-map of the 12 output promoters with each inducer. Inducer concentrations were: 25 μM DAPG, 500 μM Cuma, 10 μM OC6, 100 μM Van, 1 mM IPTG, 200 nM aTc, 4 mM Ara, 10 mM Cho, 1 mM Nar, 2.5 mM DHBA, 250 μM Sal, and 10 μM OHC14. The mean of three replicates performed on different days is shown (see Supplementary Figure 13 for data). **d)** Output of uninduced (open circles) and induced (closed circles) cultures on day 1, 7, 11, and 14 days of passaging. Inducer concentrations were: 25 μM DAPG, 500 μM Cuma, 10 μM OC6, 100 μM Van, 1 mM IPTG, 200 nM aTc, 4 mM Ara, 10 mM Cho, 1 mM Nar, 2.5 mM DHBA, 250 μM Sal, and 10 μM OHC14. A single evolutionary trajectory is shown. Passaging details and data from other days are provided in Supplementary Figure 17.

The responses of the 12 genomically-encoded sensors in Marionette-Wild are shown in Figure 3b, the parameters derived from the fits to Equation 1 are shown in Table 1, and the raw data points are provided in Supplementary Figure 11 and Supplementary Appendix 1. Each response was measured by transforming the strain with a p15a (Supplementary Figure 2) plasmid containing the output promoter fused to YFP. Each response function shows at least 100-fold induction with similar levels of on- and off-states. The technical information for the use of each sensor is organized as a series of datasheets in the Supplementary Information. The response of each sensor was also measured in Marionette-Clo and Marionette-Pro (Supplementary Appendix 1). The performances of the sensors closely match that of Marionette-Wild with several exceptions; notably, the responses to IPTG (P_Tac_) and choline (P_BetI_) are leakier.

All of the sensors follow similar induction dynamics, with induction after 15 minutes and full induction by 2 hours. Interestingly, those sensors based on activators were slower to turn on, as compared to those based on repressors (Supplementary Figure 12). There is little cross reactivity from the non-cognate inducers (Figure 3c and Supplementary Figure 13). The response functions for three different promoters were also measured in the presence of the maximum levels of all 11 other inducers, and there was little change in dynamic range (Supplementary Figure 14). The sensor responses were measured during exponential growth. To evaluate performance in stationary phase, cells were grown overnight (~20 hours) and response functions were measured (Methods). The responses closely match those measured during exponential growth, with several exceptions (Supplementary Figure 12). Induction on plates where the inducers are added to LB-agar resulted in similar responses as those observed in early stationary phase (Supplementary Figure 15).

We tested whether carrying the sensor array impacts the growth rate of the three Marionette strains (Supplementary Figure 16). The Marionette-Wild strain grows with a doubling time of 29.0 ± 1.9 min as compared 27.1 ± 1.2 for wild-type *E. coli* MG1655 (Methods). The other two Marionette strains grow slightly faster, albeit within error (Supplementary Figure 16). Growth of Marionette-Wild was also evaluated in the presence of all 12 inducers at maximum levels and only a modest effect was observed (Supplementary Figure 16).

Carrying the sensor array requires the continuous expression of 12 regulatory proteins and a transporter. This could lead to a draw on cellular resources that confers a selective advantage to eliminating the array. While genomic insertion improves evolutionary stability^21, 56–57^, it could still be disrupted over time, particularly for the *recA*-positive^58^ Marionette-Wild. To address this, we performed three independent experiments to assess the evolutionary stability of Marionette-Wild. First, we determined whether Marionette could reliably control a plasmid based promoter, even after extended passage. The 12 reporter strains were passaged for 14 days in liquid culture without inducer, diluting the cells 10^6^-fold each day (10^84^-fold dilution total). On each day, a subset of cells from each line were grown and assayed with and without inducer (Figure 3d and Supplementary Figure 17). Second, the Marionette-Wild strain was passaged for nine days, streaking cultures on agar plates and inoculating single colonies into liquid culture each day. On the tenth day, the culture was transformed with each of the 12 reporter plasmids and assayed with and without the cognate inducer for each reporter. Third, we determined if serial transfer would lead to the emergences of subpopulations^59^. We passaged the Marionette-Wild strain for nine days in liquid culture, diluting the cells 10^6^-fold each day (10^54^-fold dilution total). On the tenth day, the culture was transformed with each of the 12 reporter plasmids and assayed with and without the cognate inducer for each reporter. For all three evolution experiments, the sensors perform indistinguishably after growth and passaging (Supplementary Figure 17). There is no decline in the fold-induction over time and there was no emergence of “broken” subpopulations by flow cytometry (Supplementary Figure 18).

This work represents a dramatic expansion in our ability to study and control genes in cells. The Marionette strains enable the modular control of up to 12 genes, simply by placing each one under the control of a small (50 to 289 base pair) inducible promoter. This means that a single construct can be built and then the expression levels perturbed in many ways through the combination of different small molecules. This could be to determine the role of proteins in a natural system, picking apart the stoichiometric requirements for a molecular machine^60–61^. This can also be part of rapid optimization of metabolic pathways, where the ideal stoichiometry can be identified without the need to build megabase-scale libraries^62–63^. Through the use of CRISPRi^64^, sRNA^65^, or other tools, endogenous genes can be inducibly down-regulated as well as up-regulated, thus enabling exquisite control of natural processes and metabolic flux. Further, dynamic systems can be probed by controlling the timing of the induction of each component in order to determine the role of ordered gene expression^15, 66^. The ability to control gene expression has been a major limitation in genetic engineering; now, pulling the strings on Marionette enables unprecedented genetic control.

## Acknowledgements

Supported by U.S. Office of Naval Research Multidisciplinary University Research Initiative grant # N00014-16-1-2388 (AJM, THSS, and CAV). We would like to thank Aditya M. Kunjapur and Kristala L. J. Prather for providing DNA templates for the amplification of P_Van_, P_3B5_, *vanR*, and *pcaU*.

## Accession

The following plasmids and strains can be acquired from Addgene: pAJM.711 (P_PhlF_-YFP) 108512; pAJM.712 (P_CymRC_-YFP) 108513; pAJM.713 (P_LuxB_-YFP) 108514; pAJM.714 (P_VanCC_-YFP) 108515; pAJM.715 (P_Tac_-YFP) 108516; pAJM.717 (P_Tet*_-YFP) 108517; pAJM.716 (P_BAD_-YFP) 108518; pAJM.718 (P_BetI_-YFP) 108519; pAJM.719 (P_Ttg_-YFP) 108520; pAJM.1459 (P_3B5C_-YFP) 108521; pAJM.721 (P_SalTTC_-YFP) 108522; pAJM.944 (P_Cin_-YFP) 108523; pAJM.847 (PhlF^AM^ + P_PhlF_-YFP) 108524; pAJM.657 (CymR^AM^ + P_CymRC_-YFP) 108525; pAJM.474 (LuxR + P_LuxB_-YFP) 108526; pAJM.773 (VanR^AM^ + P_VanCC_-YFP) 108527; pAJM.336 (LacI^AM^ + P_Tac_-YFP) 108528; pAJM.011 (TetR + P_Tet*_-YFP) 108529; pAJM.677 (AraC^AM^ + AraE + P_BAD_-YFP) 108530; pAJM.683 (BetI^AM^ + P_BetI_-YFP) 108531; pAJM.661 (TtgR^AM^ + P_Ttg_-YFP) 108532; pAJM.690 (PcaU_AM_ + P_3B5B_-YFP) 108533; pAJM.771 (NahR^AM^ + P_SalTTC_-YFP) 108534; pAJM.1642 (CinR^AM^ + P_Cin_-YFP) 108535; pAJM.884 (AcuR^AM^ + P_Acu_-YFP) 108536; pAJM.969 (MphR^AM^ + EryR + P_Mph_-YFP) 108537; sAJM.1504 (Marionette-Clo) 108251; sAJM.1505 (Marionette-Pro) 108253; sAJM.1506 (Marionette-Wild) 108254.

## Author contributions

A.J.M and C.A.V. conceived the study and designed the experiments; A.J.M performed the experiments; A.J.M and T.H.S.S. analyzed the data; A.J.M and C.A.V. wrote the manuscript with input from all the authors.

## Methods

### Strains, plasmids, and media

*Escherichia coli* DH10B (New England Biolabs, Ipswich, MA – USA) was used for all routine cloning and directed evolution. Plasmid-based regulator systems were characterized in *E. coli* DH10B. Marionette-Wild, -Clo, and -Pro were derived from *E. coli* MG1655^67^, *E. coli* DH10B, and *E. coli* BL21 (New England Biolabs, Ipswich, MA – USA) cells respectively. *E. coli* JTK164H was used to clone RK6 suicide vectors^68^. All other plasmids contain p15A origins of replication and kanamycin resistance (Supplementary Figures 2 and 6). LB-Miller media (BD, Franklin Lakes, NJ - USA) was used for directed evolution, stability assays, and cytometry assays unless specifically noted. 2xYT liquid media (BD, Franklin Lakes, NJ - USA) and LB + 1.5% agar (BD, Franklin Lakes, NJ – USA) plates were used for routine cloning and strain maintenance. M9 media (1x M9 Salts [Millipore Sigma, St. Louis, MO - USA], 2 mM MgSO_4_, 100 μM CaCl_2_, and 0.2% Casamino acids) supplemented with either 0.4% glucose or 0.4% glycerol was used for cytometry assays where noted.

### Chemical transformation

For routine transformations, strains were made competent for chemical transformation. Overnight cultures (250 μl for *E. coli* DH10B derived cells, 100 μl for *E. coli* MG1655 and BL21 derived cells) were subcultured into 100 ml SOB media (BD, Franklin Lakes, NJ – USA) and grown at 37 °C, 250 rpm for 3 hours. Cultures were centrifuged (4500*g*, 4 °C, 10 minutes) and pellets were resuspended in 15 ml TFBI buffer^51^ (30 mM KOAc, 50 mM MnCl_2_, 100 mM RbCl, 10 mM CaCl_2_, and 15% v/v glycerol, pH 5.0). After 1 hour on ice, cells were centrifuged (4500*g*, 4 °C, 10 minutes) and pellets were resuspended in 2 ml TFBII buffer (10 mM NaMOPS pH 7.0, 75 mM CaCl_2_, 10 mM RbCl, and 15% v/v glycerol). Competent cells were stored at −80 °C until use.

### Response function measurements (mid-log phase)

All measurements shown were taken by cytometry of cells in mid-log growth except when noted. Glycerol stocks of strains containing the plasmids of interest were streaked on LB + 1.5% Agar plates and grown overnight at 37 °C. Single colonies were inoculated into 1 ml LB + antibiotics in 2-ml 96-deepwell plates (USA Scientific, Orlando, FL - USA) sealed with an AeraSeal film (Excel Scientific, Victorville, CA - USA) and grown at 37 °C, 900 rpm overnight in a Multitron Pro shaker incubator (INFORS HT, Bottmingen, Switzerland). The overnight growths were diluted 1:200 into 1 ml LB + antibiotics in 2-ml 96-deepwell plates + AeraSeal film and grown at 37 °C, 900 rpm. After 2 hours the growths were diluted (*E. coli* DH10B/Marionette-Clo 1:500; *E. coli* BL21/Marionette-Pro 1:2,000; *E. coli* MG1655/Marionette-Wild 1:5,000) into prewarmed LB + antibiotics + inducer where necessary in 2-ml 96-deepwell plates + AeraSeal film and grown at 37 °C, 900 rpm for 5 hours. After growth, 20 μl of culture sample was diluted into 180 μl PBS + 200 μg/ml kanamycin to inhibit translation.

### Response function measurements (stationary phase)

Measurements were taken from cells in stationary phase to generate data shown in Supplementary Appendix 1 and Supplementary Figures 12 and 13. Glycerol stocks of strains containing the plasmids of interest were streaked on LB + 1.5% Agar plates and grown overnight at 37 °C. Single colonies were inoculated into 1 ml LB + antibiotics in 2-ml 96-deepwell plates (USA Scientific, Orlando, FL - USA) sealed with an AeraSeal film (Excel Scientific, Victorville, CA - USA) and grown at 37 °C, 900 rpm overnight in a Multitron Pro shaker incubator (INFORS HT, Bottmingen, Switzerland). The overnight growths were diluted 1:200 into 1 ml LB + antibiotics in 2-ml 96-deepwell plates + AeraSeal film and grown at 37 °C, 900 rpm. After 2 hours the growths were diluted (*E. coli* DH10B/Marionette-Clo 1:500; *E. coli* BL21/Marionette-Pro 1:2,000; *E. coli* MG1655/Marionette-Wild 1:5,000) into prewarmed LB + antibiotics + inducer where necessary in 2-ml 96-deepwell plates + AeraSeal film and grown at 37 °C, 900 rpm for 20 hours. After growth, 2 μl of culture sample was diluted into 198 μl PBS + 200 μg/ml kanamycin to inhibit translation.

### Time course (mid-log phase)

For Supplemental Figure 12: mid-log induction time course, glycerol stocks of strains containing the plasmids of interest were streaked on LB + 1.5% Agar plates and grown overnight at 37 °C. Single colonies were inoculated into 1 ml LB + antibiotics in 2-ml 96-deepwell plates (USA Scientific, Orlando, FL - USA) sealed with an AeraSeal film (Excel Scientific, Victorville, CA - USA) and grown at 37 °C, 900 rpm overnight in a Multitron Pro shaker incubator (INFORS HT, Bottmingen, Switzerland). The overnight growths were diluted 1:200 into 1 ml LB + antibiotics in 2-ml 96-deepwell plates + AeraSeal film and grown at 37 °C, 900 rpm. After 2 hours the growths were diluted 1:500 into prewarmed LB + antibiotics. After 0, 1, 2, 3, 3.5, 4, 4.25, 4.5, or 4.75 hours, cultures were further diluted 1:10 into prewarmed LB + antibiotics + inducer where necessary in 2-ml 96-deepwell plates + AeraSeal film and grown at 37 °C, 900 rpm for 5, 4, 3, 2, 1.5, 1, 0.75, 0.5, or 0.25 hours (5 hours total after the initial growth). After growth, 20 μl of culture sample was diluted into 180 μl PBS + 200 μg/ml kanamycin to inhibit translation.

### Time course (mid-log phase to stationary phase)

For Supplemental Figure 12: mid-log to stationary induction time course, glycerol stocks of strains containing the plasmids of interest were streaked on LB + 1.5% Agar plates and grown overnight at 37 °C. Single colonies were inoculated into 1 ml LB + antibiotics in 2-ml 96-deepwell plates (USA Scientific, Orlando, FL - USA) sealed with an AeraSeal film (Excel Scientific, Victorville, CA - USA) and grown at 37 °C, 900 rpm overnight in a Multitron Pro shaker incubator (INFORS HT, Bottmingen, Switzerland). The overnight growths were diluted 1:200 into 1 ml LB + antibiotics in 2-ml 96-deepwell plates + AeraSeal film and grown at 37 °C, 900 rpm. After 2 hours the growths were diluted 1:5000 into prewarmed LB + antibiotics + inducer where necessary in 2-ml 96-deepwell plates + AeraSeal film and grown at 37 °C, 900 rpm for 5, 6, 7, 8, 9, 10, or 20 hours. After growth, 2 to 20 μl of culture sample was diluted into 180-198 μl PBS + 200 μg/ml kanamycin to inhibit translation.

### Cytometry analysis

Fluorescence characterization with cytometry was performed on a BD LSR Fortessa flow cytometer with HTS attachment (BD, Franklin Lakes, NJ - USA). Cells diluted in PBS + kanamycin were run at a rate of 0.5 μl/s. The events were gated by forward scatter height (mid-log: 1,000-10,000; stationary: 500-5000) and side scatter area (mid-log: 1,000-10,000; stationary: 500-5,000) to reduce false events. After gating, thousands of events were used for analysis. For each sample, the median YFP fluorescence was calculated. All output values are reported in terms of relative promoter units (RPU). For a given promoter measurement, the strain (*E. coli* DH10B, Marionette-Wild, *etc*) is transformed with the plasmid. The strain is then assayed alongside a strain containing the RPU standard plasmid (Supplementary Figure 1, Supplementary Table 8) as well as an autofluorescence control. The median autofluorescence value is subtracted from the all other median fluorescence values, including that of the RPU standard. The experimental sample value is then divided by the RPU standard value.

### Library generation

Portions of the initial libraries were created by PCR using degenerate oligonucleotides (Integrated DNA Technologies Coralville, IA - USA). These fragments were joined into a degenerate, full-length sensor module by overlap PCR. Sensor modules were assembled into selection vectors by Golden Gate assembly^70^. Linear insert and plasmid selection vector were mixed at 1:1 molar ratio totaling ~ 1 μg DNA along with 5 μl 10x T4 ligase Buffer, 1 μl T4 DNA ligase (2,000,000 U/ml), 2 μl BbsI (10,000 U/ml) (all from New England Biolabs, Ipswich, MA - USA) in 50 μl total. Reactions were cycled 45 times between 2 minutes at 37 °C and 5 minutes at 16 °C, and then incubated for 30 minutes at 50 °C, 30 minutes at 37 °C, and 10 minutes at 80 °C in a DNA Engine cycler (Bio-Rad, Hercules, CA – USA). An additional 1 μl BbsI was then added, and the assembly was incubated for 1 hour at 37 °C. Assemblies were then purified using Zymo Spin I columns (Zymo Research, Irvine, CA - USA). Host cells were transformed with library plasmid by electroporation. Supplementary Appendix 2 contains a depiction of all degeneracy found in the “Initial library” for each selection.

### Negative selection

LB media + 8 mM Cl-Phe (4-Chloro-DL-phenylalanine [Millipore Sigma, St. Louis, MO - USA]) was mixed, autoclaved, and stored at room temperature. Cl-Phe has a tendency to adhere to glassware, and care was taken to avoid disturbing the water + LB powder + Cl-Phe powder mixture prior to autoclaving. LB media + 8 mM Cl-Phe was mixed with plain LB media to achieve the desired concentration of Cl-Phe (See Interventions:[Cl-Phe] for each selection in Supplementary Appendix 2). Following transformation and outgrowth, cultures were diluted into 7 ml LB + Cl-Phe supplemented with antibiotics as well as any cross-reactive inducers (See Interventions:[Negative ligand] for each selection in Supplementary Appendix 2). Cultures were grown at 37 °C for 12-16 hours.

### Positive selection

Following negative selection, cultures were diluted 1:100 into 2 ml LB + antibiotics in culture tubes and grown at 37 °C, 250 rpm for 2 hours. The cultures were then diluted 1:100 into 2 ml prewarmed LB + antibiotics + inducer (See Interventions:[Positive ligand] for each selection in Supplementary Appendix 2) and grown at 37 °C, 250 rpm for 4 hours. Following induction, 1 ml of culture was centrifuged at (5,000*g*, 25 °C, 10 minutes). Supernatant was removed and the cell pellet was resuspended in 50 μl 1x CPR buffer (50 mM Tris-HCl pH 8.8, 10 mM KCl, 2 mM MgSO_4_, 10 mM (NH_4_)_2_SO_4_). 5 μl resuspension was added to 95 μl of 1 x CPR buffer plus 0.4 μM CPR primers and 200 μM dNTPs. This aqueous phase added to a 2 ml centrifuge tube containing 438 μl Tegosoft DEC (Evonik, Essen, Germany), 42 μl AbilWE09 (Evonik, Essen, Germany) and 120 μl mineral oil (Millipore Sigma, St. Louis, MO - USA) and the rubber stopped from a 1 ml syringe. The mixture was vortexed at maximum setting for 2 minutes. The emulsion was split evenly five 0.2 ml PCR tubes, and thermal cycled (95 °C for 5 minutes; 20 cycles of [95 °C for 30 seconds, 55 °C for 30 seconds, 72 °C for 2 minutes/kb]; and 72 °C for 5 min). Emulsions were then centrifuged (10,000*g*, 25 °C, 10 minutes) and the upper (oil) phase was removed. Then, 100 μl H^2^O and 500 μl chloroform were added and the mixture was pipetted to disrupt the pellet. The resuspension was then transfered to a 1.5 ml heavy-gel phase-lock tube (5 Prime, San Francisco, CA - USA) and centrifuged (16,000*g*, 25 °C, 2 minutes). The upper (aqueous) phase was collected, and DNA was purified using Zymo Spin I columns. The library was amplified in a recovery PCR using Accuprime *Pfx* and nested recovery primers, gel purified, and assembled as described above. A depiction of the CPR primer and recovery primer amplification are provided in Supplementary Figure 6.

### Library shuffling

Between some rounds of selection (See Interventions:Notes for each selection in Supplementary Appendix 2), libraries were shuffled upon themselves such that fragments of each mutant mutually serve as both primer and template for a primer-less PCR amplification^54^. 1 μg linear, library DNA (from the recovery PCR) was added to a mild DNAse reaction (500 mM Tris pH 7.4, 100 mM MnCl2, 0.5 U DNAse [New England Biolabs, Ipswich, MA - USA]) and lightly digested for 3 minutes at 15 °C. Fragmented DNA was purified using Zymo Spin I columns and reassembled in a primer-less PCR in 1 x KAPA HiFi Master Mix (KAPA Biosystems, Wilmington, MA – USA) by thermal cycling (95 °C for 2 minutes; 35 cycles of [95 °C for 30 seconds, 65 °C for 90 seconds, 62 °C for 90 seconds, 59 °C for 90 seconds, 56 °C for 90 seconds, 53 °C for 90 seconds, 50 °C for 90 seconds, 47 °C for 90 seconds, 44 °C for 90 seconds, 41 °C for 90 seconds, 68 °C for 90 seconds]; and 72 °C for 4 minutes). The reassembly was purified using Zymo Spin I columns, reamplified using Accuprime *Pfx* and CPR primers, gel purified, and assembled as described above.

### On/off screen

At the end of each selection, libraries were assembled into the YFP screening plasmid (Supplementary Figure 6). Host cells were transformed and plated on LB-agar. Between 35 and 92 individual clones were picked and assayed by cytometry as described. Cells were grown with no inducer, in the presence of cognate inducer, and when necessary in the presence of relevant non-cognate inducers. Measurements were made of cells in mid-log phase by cytometry as described. The most promising clones, as judged by dynamic range and orthogonality, were mini-prepped and the sensor region was sequenced.

### Genomic integration

In preparation of recombineering, cells were transformed with a plasmid containing Ara-inducible λ Red recombination machinery with a temperature sensitive origin of replication^71^. 50 μl of overnight culture was subcultured in 50 ml LB medium and grown at 30 °C, 250 rpm for 2 hours. 2 mM Ara was added, and the culture continued to grow at 30 °C, 250 rpm for 3 hours. The culture was then centrifuged (4500*g*, 4 °C, 10 minutes) and washed with ice cold 10% glycerol four times, with the fourth resuspension in 200 μl 10% glycerol. Recombineering-ready cells were stored at −80 °C until use. For the first insertion, six genes (*phlF*^*AM*^, *cymR*^*AM*^, *luxR*, *vanR*^*AM*^, *lacI*^*AM*^, and *tetR*) were Golden Gate assembled using BsaI into an RK6 suicide vector (which needs Pir protein in order to propagate)^68^. Pir-expressing *E. coli* JTK164H cells were transformed, and plasmids were purified, verified, and linearized with BpiI leaving homology to the *glvC* pseudogene. Recombineering-ready *E. coli* MG1655 cells were electroporated and transformed with gel-purified, linearized inserts. After an outgrowth of one hour at 37 °C, transformations were plated on LB-agar plates + antibiotic (5 μg/ml chloramphenicol). Colonies were picked and grown at 37°C in LB + antibiotic, and the presence of the insert was verified by colony PCR. For the second insertion, this strain was made recombineering-ready and the process was repeated with the next set of genes (*araC*^*AM*^, *araE*, *betI*^*AM*^, and *ttgR*^*AM*^) with the insert containing homology to *tetR* and *glvC* and conferring resistance to 20 μg/ml spectinomycin. For the third insertion, this strain was made recombineering-ready and the process was repeated with the final set of genes (*pcaU*^*AM*^, *nahR*^*AM*^, and *cinR*^*AM*^) with the insert containing homology to *ttgR*^*AM*^ and *glvC* and conferring resistance to 5 μg/ml chloramphenicol. See Supplementary Figure 9 for schematic details.

### Phage transduction to transfer the Marionette cluster

50 μl of overnight culture of Marionette-Wild was subcultured in 5 ml LB medium + 0.2% glucose and 5 mM CaCl_2_ and grown at 37 °C, 250 rpm for 30 minutes. 100 μl p1 phage lysate (10^9^ pfu/ml) was added and the culture was grown at 37 °C, 250 rpm until lysis (3 hours)^72^. 50 μl chloroform was added and the culture continued at 37 °C, 250 rpm for 5 minutes. Culture was centrifuged (9200*g*, 25 °C, 10 minutes). Supernatant was filtered (0.45 μM) and stored at 4 °C until use. 1.5 ml of overnight culture of *E. coli* MG1655, BL21, or DH10B (the DH10B cells contained a temperature-sensitive plasmid transiently expressing RecA) was centrifuged (10,000*g*, 25 °C, 5 minutes). The cell pellet was resuspended in 750 μl P1 buffer (5 mM MgSO_4_ and 10 mM CalCl_2_) and up to 100 μl Marionette-P1 phage lysate was added. After 30 minutes at 25 °C, 1 ml LB + 200 μl 1 M sodium citrate was added and the culture was grown at 37 °C, 250 rpm for 30 minutes. The culture was centrifuged (10,000*g*, 25 °C, 2 minutes) and the pellets were resuspended in 100 μl LB and plated on LB-agar plates + 5 μg/ml chloramphenicol and 5 mM sodium citrate. Colonies were picked and grown at 37 °C in LB + 5 μg/ml chloramphenicol and 5 mM sodium citrate, and the presence of the insert was verified by colony PCR. The entire Marionette cluster was verified by sequencing PCR amplicons of the cluster from the genome of Marionette-Wild.

### Sensor induction on plates

Cells were transformed with reporter plasmids and, following outgrowth at 37°C for one hour, plated on LB-agar (LB-Miller powder + 1.5 % agar [BD, Franklin Lakes, NJ – USA]) with and without the appropriate inducer. After an overnight incubation at 37°C for 16 hours, plates were incubated at 4°C for one hour and imaged using a ChemiDocMP Imaging system (Bio-Rad, Hercules, CA - USA) and Image Lab 4.0 software (Bio-Rad, Hercules, CA - USA) employing blue epi illumination, a 530/28 filter and a 0.1 second exposure. Raw images were rotated and cropped using XnView (XnSoft, Reims, France). Individual colonies were scraped from the plate, resuspended in 200 μl PBS + 200 μg/ml kanamycin, and assayed by cytometry as described in Cytometry analysis.

### Plate reader assays to measure growth rates

Glycerol stocks of strains of interest were streaked on LB + 1.5% Agar plates and grown overnight at 37 °C. Single colonies were inoculated into 1 ml LB in 2-ml 96-deepwell plates (USA Scientific, Orlando, FL - USA) sealed with an AeraSeal film (Excel Scientific, Victorville, CA - USA) and grown at 37 °C, 900 rpm overnight in a Multitron Pro shaker incubator (INFORS HT, Bottmingen, Switzerland). The overnight growths were diluted 1:200 into 1 ml LB in 2-ml 96-deepwell plates + AeraSeal film and grown at 37 °C, 900 rpm. After 2 hours the growths were diluted (*E. coli* DH10B/Marionette-Clo 1:500; *E. coli* BL21/Marionette-Pro 1:2,000; *E. coli* MG1655/Marionette-Wild 1:5,000) into prewarmed LB + inducer where necessary in 2-ml 96-deepwell plates. 100 µl of this culture was immediately transferred to a 300-μl 96-well black walled optical bottom plates (Thermo Scientific Nunc, Waltham, MA – USA) sealed with a BreathEasy film (Sigma-Aldrich, St. Louis, MO – USA), and grown in a Synergy H1 plate reader (BioTek, Winooski, VT – USA) at 37 °C, 1000 rpm. OD_600_ was measured every 20 minutes over 12 hours of growth. OD_600_ readings were also taken from wells containing media with no cells, and for each time point, readings from such wells were subtracted from the appropriate sample measurements to remove background. OD_600_ values were converted to equivalent 1 cm path length measurements using a standard curve.

### Calculation of growth rates

To calculate doubling times, the last measurement with OD_600_ < 0.1 and the first measurement with OD_600_ > 0.4 were identified, and the doubling time was calculated assuming exponential growth between those two points. Doubling time is calculated as elapsed time (in minutes) divided by the number of doublings that occurred in that time (log_2_[final OD_600_/initial OD_600_]).

### Evolutionary stability (passaging with reporters)

Marionette-Wild was transformed with each of the 12 reporter plasmids and plated on LB-agar. Single colonies were inoculated into 1 ml LB + antibiotics in 2- ml 96-deepwell plates (USA Scientific, Orlando, FL - USA) sealed with an AeraSeal film (Excel Scientific, Victorville, CA - USA) and grown at 37 °C, 900 rpm overnight in a Multitron Pro shaker incubator (INFORS HT, Bottmingen, Switzerland). The overnight growths were diluted 1:200 into 1 ml LB + antibiotics in 2-ml 96-deepwell plates + AeraSeal film and grown at 37 °C, 900 rpm. After 2 hours, the growths were diluted 1:5000 into prewarmed LB + antibiotics + inducer where necessary in 2-ml 96-deepwell plates + AeraSeal film and grown at 37 °C, 900 rpm for 5 hours. After growth, 20 μl of culture sample was diluted into 180 μl PBS + 200 μg/ml kanamycin to inhibit translation and assayed by cytometry as described in Cytometry analysis. Uninduced cultures were allowed to grow at 37 °C overnight (17 hours) and the process (beginning with the 1:200 dilution of the overnight culture) was repeated each day for 14 days.

### Evolutionary stability (passaging on plates)

A single colony of Marionette-Wild was inoculated into 1 ml LB in 2-ml 96-deepwell plates (USA Scientific, Orlando, FL - USA) sealed with an AeraSeal film (Excel Scientific, Victorville, CA - USA) and grown at 37 °C, 900 rpm for 12 hours in a Multitron Pro shaker incubator (INFORS HT, Bottmingen, Switzerland). The overnight growth was streaked onto LB-agar plates and grown at 37 °C for 12 hours. This process was repeated 9 times. The final culture was made chemically competent as described in Chemical transformation, transformed with relevant reporter plasmids, and assayed with cytometry as described in Cytometry analysis.

### Evolutionary stability (passaging in liquid culture)

A single colony of Marionette-Wild was inoculated into 1 ml LB in 2-ml 96-deepwell plates (USA Scientific, Orlando, FL - USA) sealed with an AeraSeal film (Excel Scientific, Victorville, CA - USA) and grown at 37 °C, 900 rpm for 12 hours in a Multitron Pro shaker incubator (INFORS HT, Bottmingen, Switzerland). The overnight growths were diluted 1:200 into 1 ml LB in 2-ml 96-deepwell plates + AeraSeal film and grown at 37 °C, 900 rpm. After 2 hours, the growths were diluted 1:5000 into prewarmed LB in 2-ml 96-deepwell plates + AeraSeal film and grown at 37 °C, 900 rpm for 22 hours. This process was repeated 9 times. The final culture was made chemically competent as described in Chemical transformation, transformed with relevant reporter plasmids, and assayed with cytometry as described in Cytometry analysis.

### Computational methods (fitting to the response function)

To parameterize the response function, error minimization was performed using the Solver function in Excel software (Microsoft, Redmond, WA - USA). ^Equation 1 was entered with *y*_*min*_, *y*_*max*_, *K*, and *n* as tunable parameters, *x* (inducer concentration) as the^ independent variable and *y* as the output. For each *x*, the error between the measured RPU value and the output of the function was determined. The total error was determined by summing the normalized square of the error ([*y*-measured RPU value]^2^ /*y*) for each *x*. The Solver function minimized the total error by tuning *y*_*min*_, *y*_*max*_, *K*, and *n*.

### Computational methods (partial randomization of amino acids)

Initial libraries for the selection of LacI and AraC utilized partial randomization of amino acids. Previous literature informed decisions regarding the desirability of including each possible residue in the library. The CASTER 2.0 tool^73^ was used to design degenerate oligodeoxynucleotides that can sample the desired amino acids while limiting undesired amino acids and stop codons.

### Computational methods (RBS design)

The initial RBS for each regulator in the Marionette cluster was designed using the RBS Calculator v1.1^55^ using the *E. coli* DH10B 16S rRNA setting. The “Pre-Sequence” included the last 100 bp of the upstream CDS (where appropriate) as well as any scars used in Golden Gate assembly. The “Target Translation Initiation Rates” were chosen based on the RBS strength of the plasmid-based sensor (found using the “Reverse Engineer RBSs” tool) and compensating for the reduction in copy number and increase in promoter strength associated with the move to the genomic system. After initial assembly and testing, a decision was made based on the appearance of the response function as to whether the RBS was too low, too high, or good. Low RBSs were rationally mutated to more closely resemble the consensus Shine-Dalgarno sequence (TAAGGAGGT) while high RBSs were rationally mutated to less closely resemble the consensus Shine-Dalgarno sequence. All rationally designed RBS variants were checked using the “Reverse Engineer RBSs” to guard against making dramatic, unanticipated changes to the RBS strength. Rational mutations are noted in Supplementary Table 2. Translation Initiation Rate (in arbitrary units) for each RBS variant is provided in Supplementary Figure 10.

