## Supplementary Materials for "Marionette: *E. coli* containing 12 highly-optimized small molecule sensors"

### Supplementary Appendices

**Supplementary Appendix 1: Sensor specification sheets**

Each page details the performance specifications of 14 sensors.

| **Regulator:$P_{Out}$:Inducer** | **Notes** |
|---|---|
| Information about the inducer including recommendations for preparation and storage of stocks. Recommended maximum induction concentrations are based on induction saturation, solubility, cellular toxicity, cross-reactivity, and antagonism. | Notes that may prove useful to the user. Known instances of cross reactivity are reported; values are based on fold-induction. Known instances of antagonism are reported; values are based on the change in fold-induction at the midpoint of the response function. |

#### Plasmid-based regulator

Information about the plasmid-based regulator system in *E. coli* DH10B. Response of the output promoter as a function of inducer concentration in *E. coli* DH10B in mid-log phase is plotted. The mean of three replicates performed on different days is shown, and error bars represent the standard deviation. Dashed lines indicate the extension of the response function beyond inducer concentrations for which data have been collected. The grey boxed region denotes inducer concentrations above the recommended maximum induction concentration. Recommended maximum induction concentrations are based on induction saturation, solubility, cellular toxicity, cross-reactivity, and antagonism.

Each response function was obtained by fitting the experimental data to Equation 1 (Methods) and also appears in Figure 2a. The response function parameters are provided in Table 1.

The plasmid map(s) used to generate these data are shown, and their sequences are provided in Supplementary Table 8.

Cytometry histograms for no induction (white), moderate induction (grey), and maximum induction (black) are shown. Raw values are white cell corrected and RPU normalized.

#### Genome-based regulator

Information about the genome-based regulator system in Marionette strains. Response of the output promoter as a function of inducer concentration in Marionette-Wild in mid-log phase is plotted. The mean of three replicates performed on different days is shown, and error bars represent the standard deviation. Dashed lines indicate the extension of the response function beyond inducer concentrations for which data have been collected. The grey boxed region denotes inducer concentrations above the recommended maximum induction concentration. Recommended maximum induction concentrations are based on induction saturation, solubility, cellular toxicity, cross-reactivity, and antagonism.

Each response function was obtained by fitting the experimental data to Equation 1 (Methods) and also appears in Figure 3b. The response function parameters are provided in Table 1.

The plasmid map(s) used to generate these data are shown, and their sequences are provided in Supplementary Table 8.

Cytometry histograms for no induction (white), moderate induction (grey), and maximum induction (black) are shown. Raw values are white cell corrected and RPU normalized.

Outputs in the presence (black bars) and absence (white bars) of the appropriate inducer in Marionette-Wild, -Clo, and -Pro in mid-log and stationary phase are shown. The mean of three replicates performed on different days is shown, and error bars represent the standard deviation.

### $PhlF^{AM}$:$P_{PhlF}$:DAPG

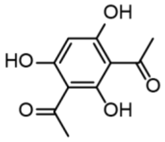

| | |
|---|---|
| Inducer | 2,4-Diacetylphloroglucinol |
| Source | Santa Cruz sc-206518 |
| Stock | 25 mM |
| Solvent | DMF |
| Max induction | 25 µM |
| Storage | -20 °C |

#### Notes

-PhlF is a TetR family repressor from *Pseudomonas fluorescens*

-$PhlF^{AM}$:$P_{PhlF}$ is 2.6-fold induced by 100 µM Sal

#### Plasmid-based regulator

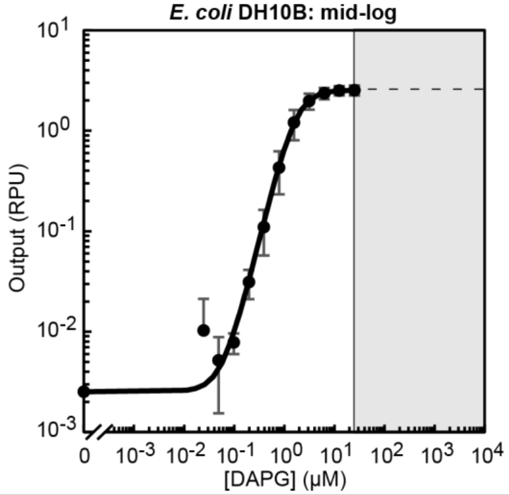

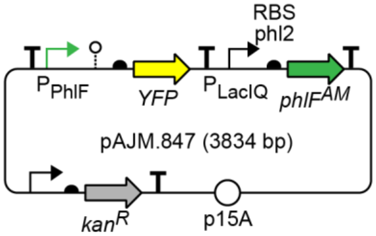

| | |
|---|---|
| Max (RPU) | 2.5 |
| Min (RPU) | $2.5\times10^{-3}$ |
| K (µM) | $1.7\times10^{0}$ |
| n | 2.1 |

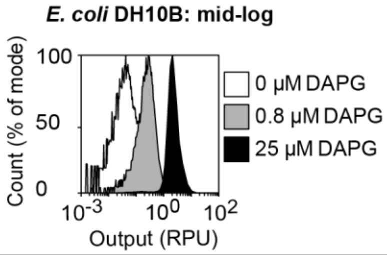

#### Genome-based regulator

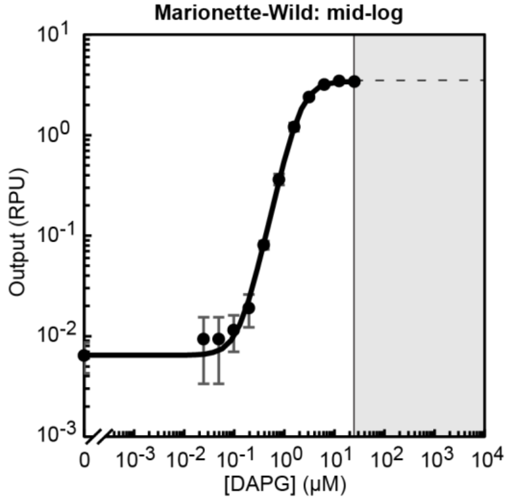

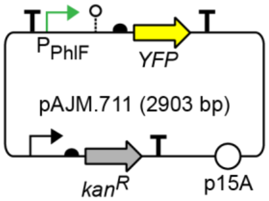

| | |
|---|---|
| Max (RPU) | 3.4 |
| Min (RPU) | $6.5\times10^{-3}$ |
| K (µM) | $2.1\times10^{0}$ |
| n | 2.3 |

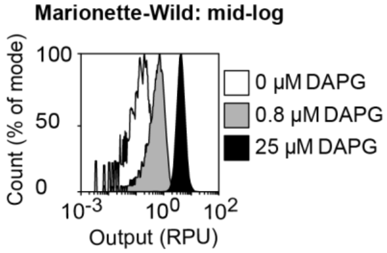

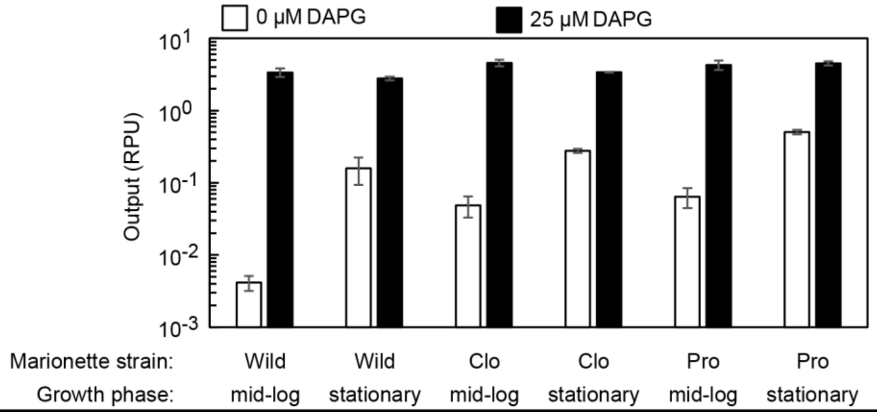

### CymR$^{AM}$:P$_{CymRC}$:Cuma

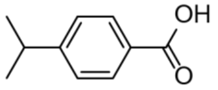

| | |
|---|---|
| Inducer | Cuminic acid |
| Source | Sigma 268402 |
| Stock | 100 mM |
| Solvent | EtOH |
| Max induction | 100 μM |
| Storage | -20 °C |

**Notes**

-CymR is a TetR family repressor from *Pseudomonas putida*

-CymR$^{AM}$:P$_{CymRC}$:Cuma induction is 13-fold antagonized by 100 μM Sal

#### Plasmid-based regulator

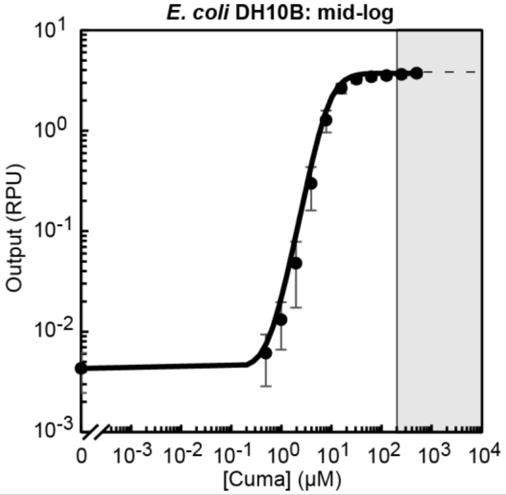

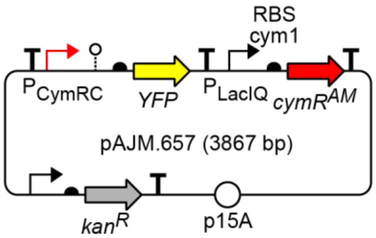

| | |
|---|---|
| Max (RPU) | 3.7 |
| Min (RPU) | $4.3 \times 10^{-3}$ |
| K (μM) | $8.9 \times 10^{0}$ |
| n | 2.4 |

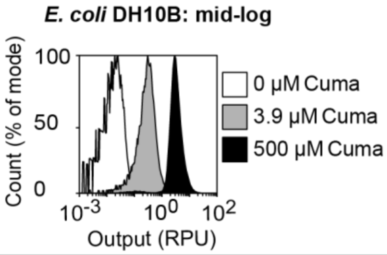

#### Genome-based regulator

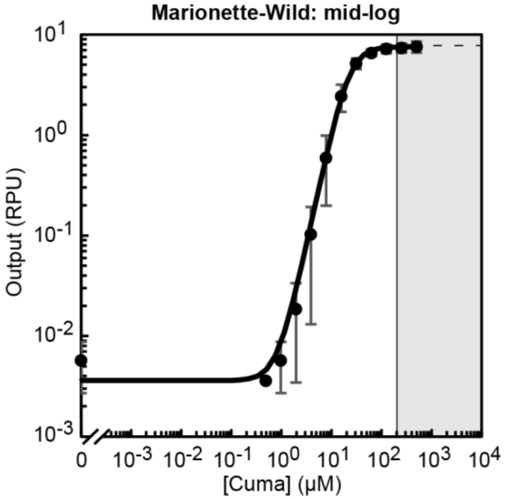

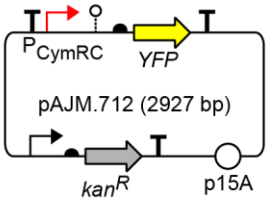

| | |
|---|---|
| Max (RPU) | 7.6 |
| Min (RPU) | $3.6 \times 10^{-3}$ |
| K (μM) | $2.2 \times 10^{1}$ |
| n | 2.3 |

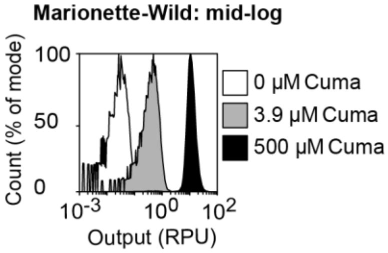

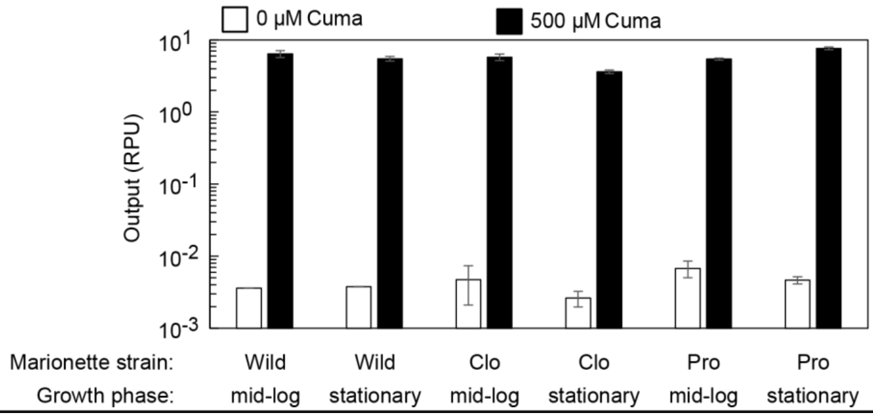

### LuxR:$P_{LuxB}$:OC6

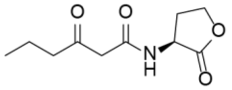

| | |
|---|---|
| Inducer | 3OC6-HSL |
| Source | Sigma K3007 |
| Stock | 10 mM |
| Solvent | DMF |
| Max induction | 10 µM |
| Storage | -20 °C |

#### Notes

-LuxR is a LuxR family activator from *Vibrio fischeri*

#### Plasmid-based regulator

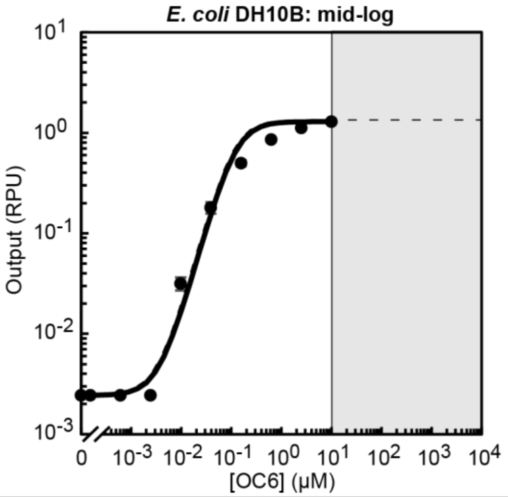

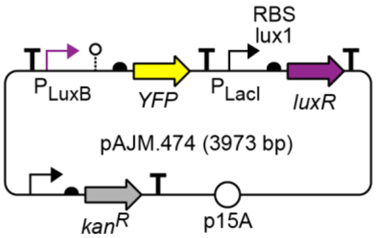

| | |
|---|---|
| Max (RPU) | 1.3 |
| Min (RPU) | $2.4 \times 10^{-3}$ |
| K (nM) | $1.2 \times 10^{-1}$ |
| n | 1.8 |

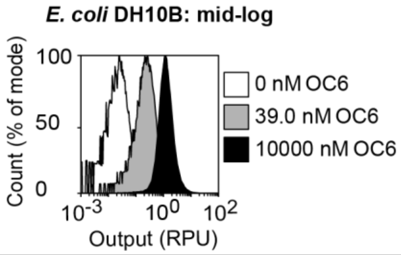

#### Genome-based regulator

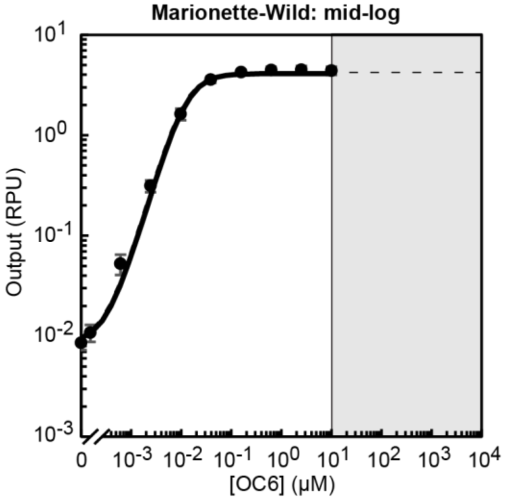

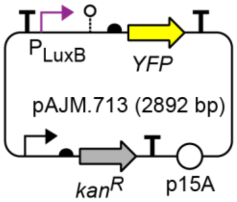

| | |
|---|---|
| Max (RPU) | 4.1 |
| Min (RPU) | $8.6 \times 10^{-3}$ |
| K (nM) | $1.2 \times 10^{-2}$ |
| n | 1.7 |

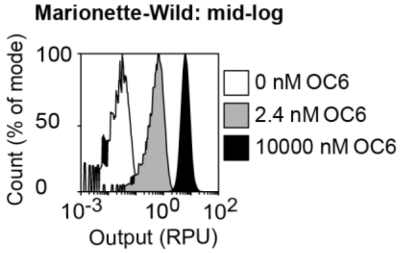

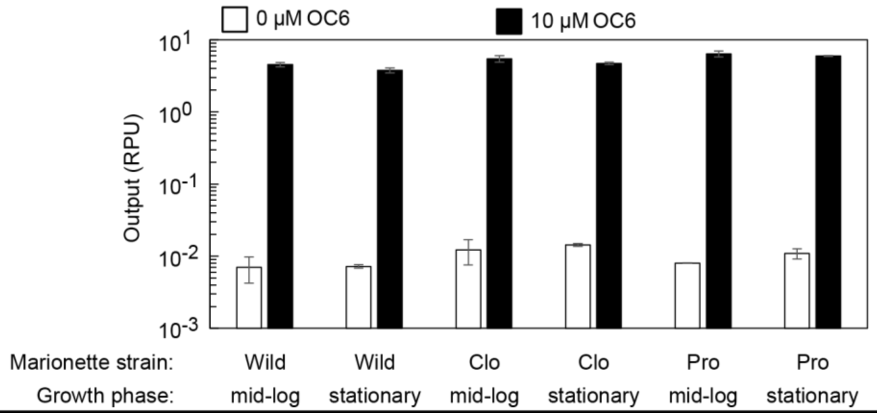

### VanR$^{AM}$:P$_{VanCC}$:Van

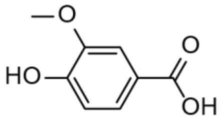

| | |
|---|---|
| Inducer | Vanillic acid |
| Source | Sigma 94770 |
| Stock | 100 mM |
| Solvent | EtOH |
| Max induction | 100 µM |
| Storage | -20 °C |

**Notes**

-VanR is a GntR family repressor from *Caulobacter crescentus*

-PcaU$^{AM}$:P$_{3B5}$ is 2.2-fold induced by 100 µM Van

-VanR$^{AM}$:P$_{VanCC}$ is 1.3-fold induced by 1 mM DHBA

#### Plasmid-based regulator

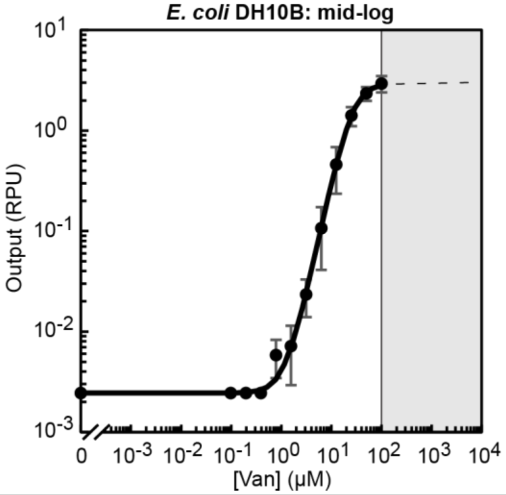

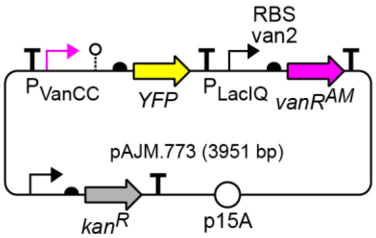

| | |
|---|---|
| Max (RPU) | 3.0 |
| Min (RPU) | 2.4x10$^{-3}$ |
| K (µM) | 2.6x10$^{1}$ |
| n | 2.3 |

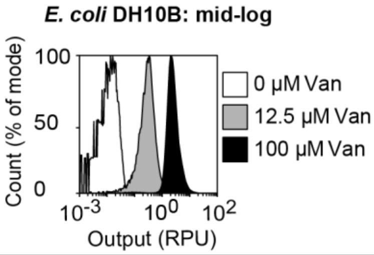

#### Genome-based regulator

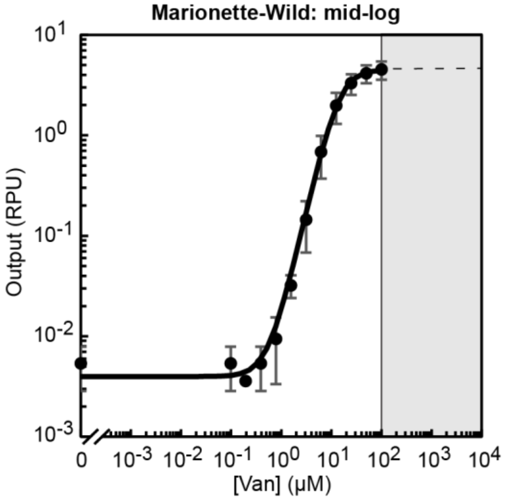

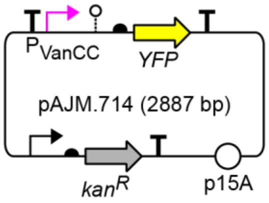

| | |
|---|---|
| Max (RPU) | 4.5 |
| Min (RPU) | 4.0x10$^{-3}$ |
| K (µM) | 1.4x10$^{1}$ |
| n | 2.1 |

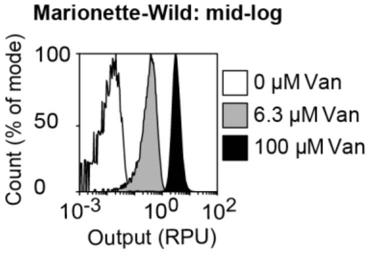

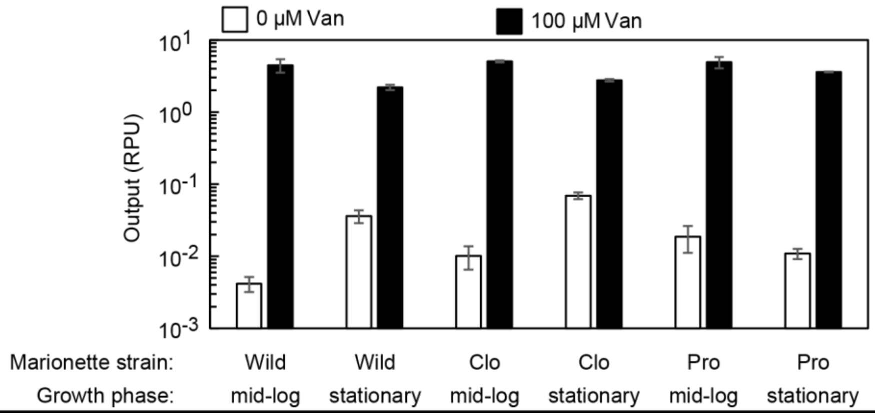

### LacI$^{AM}$:P$_{Tac}$:IPTG

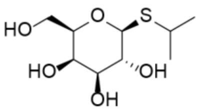

| | |
|---|---|
| Inducer | Isopropyl-β-D-thiogalactoside |
| Source | Gold - I2481C25 |
| Stock | 1 M |
| Solvent | Water |
| Max induction | 1 mM |
| Storage | -20 °C |

**Notes**

-LacI is a LacI family repressor from *Escherichia coli;* native gene is present in MG1655, DH10B, and BL21, including Marionette strains

-AraC$^{AM}$:P$_{BAD}$:Ara induction is 1.5-fold inhibited by 1 mM IPTG

-The P$_{Tac}$ used contains a symmetrical lac operator[2]

#### Plasmid-based regulator

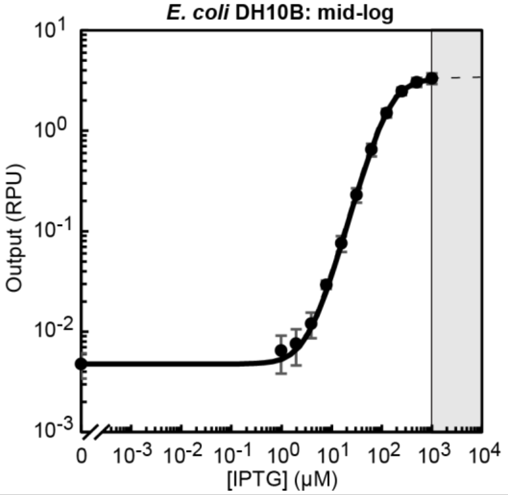

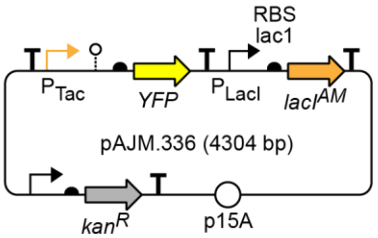

| | |
|---|---|
| Max (RPU) | 3.3 |
| Min (RPU) | 4.8x10$^{-3}$ |
| K (µM) | 1.4x10$^{2}$ |
| n | 1.8 |

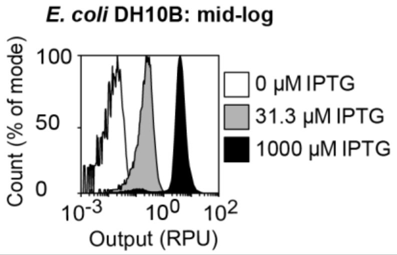

#### Genome-based regulator

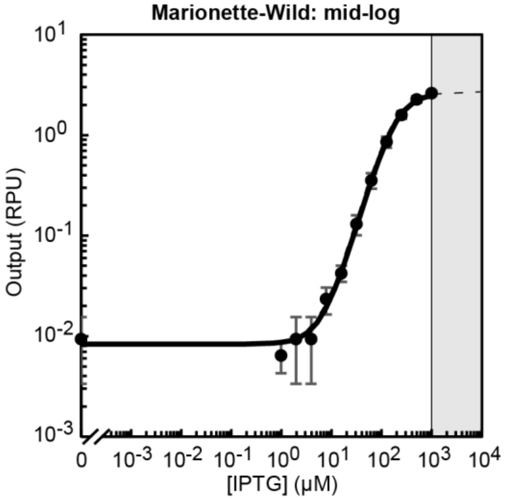

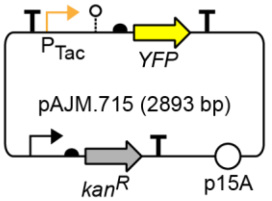

| | |
|---|---|
| Max (RPU) | 2.6 |
| Min (RPU) | 8.3x10$^{-3}$ |
| K (µM) | 1.9x10$^{2}$ |
| n | 1.7 |

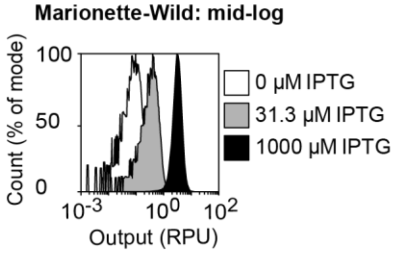

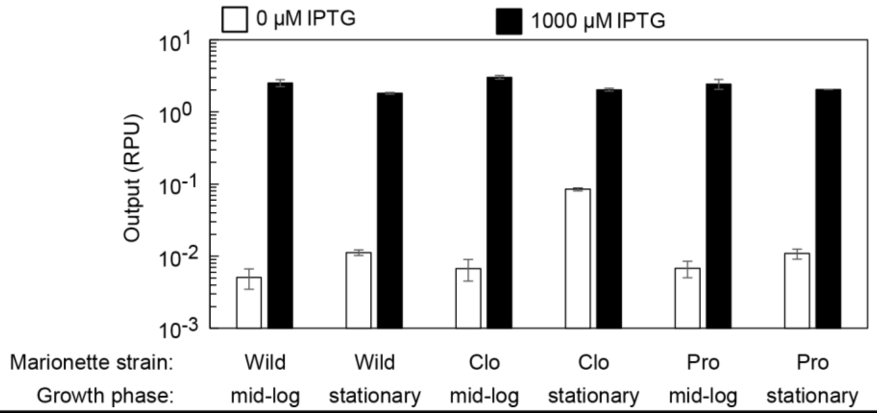

### TetR:$P_{Tet*}$:aTc

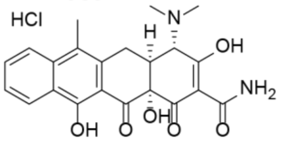

| | |
|---|---|
| Inducer | Anhydrotetracycline HCl |
| Source | Sigma 37919 |
| Stock | 200 µM |
| Solvent | 1:1 Water:EtOH |
| Max induction | 200 nM |
| Storage | -20 °C |

#### Notes

-TetR is a TetR family repressor from an *Escherichia coli* mobile element; not presenct in MG1655, DH10B, or BL21

#### Plasmid-based regulator

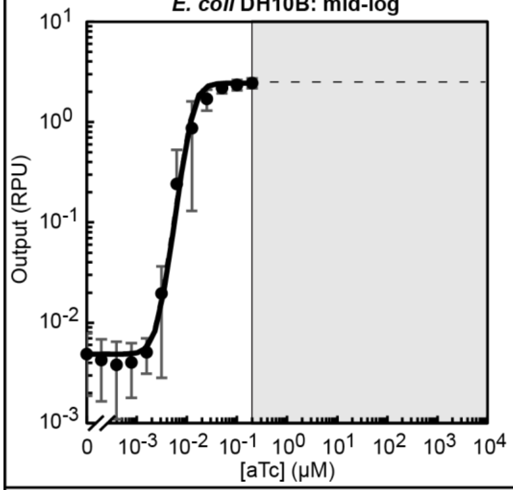

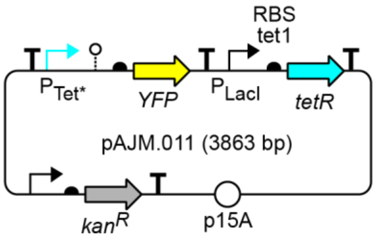

| | |
|---|---|
| Max (RPU) | 2.4 |
| Min (RPU) | $4.9 \times 10^{-3}$ |
| K (µM) | $1.3 \times 10^{-2}$ |
| n | 3.8 |

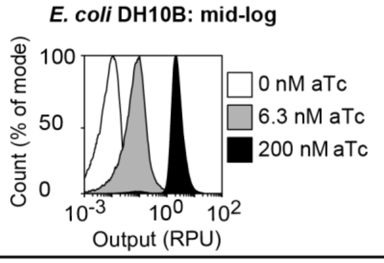

#### Genome-based regulator

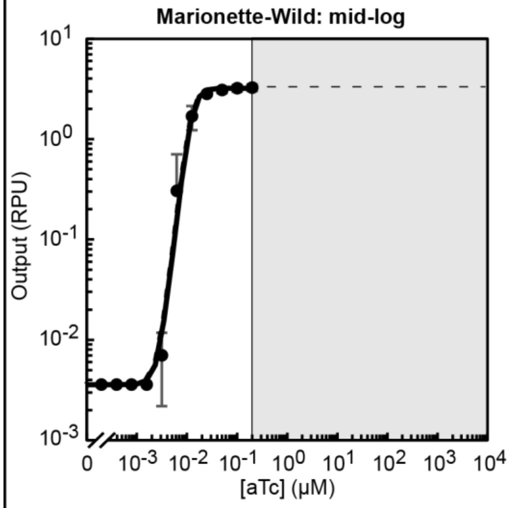

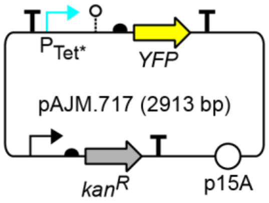

| | |
|---|---|
| Max (RPU) | 3.2 |
| Min (RPU) | $3.6 \times 10^{-3}$ |
| K (µM) | $1.2 \times 10^{-2}$ |
| n | 4.4 |

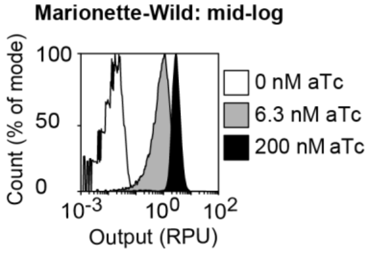

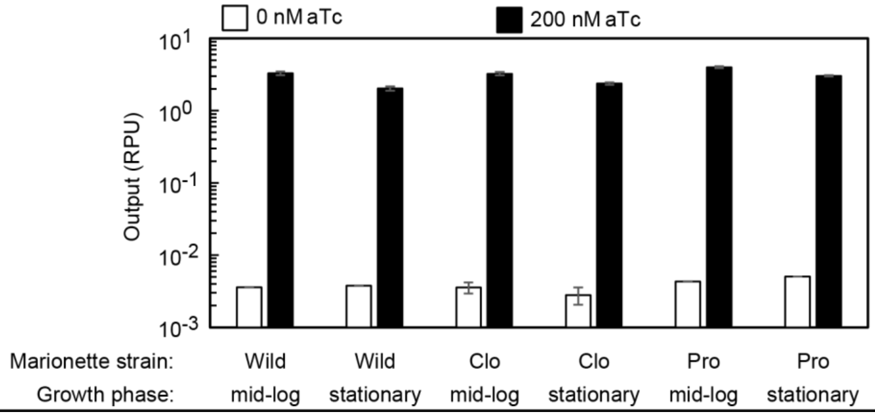

### AraC$^{AM}$:P$_{BAD}$:Ara

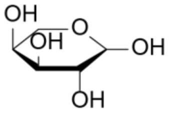

| | |
|---|---|
| Inducer | L-Arabinose |
| Source | Sigma A3256 |
| Stock | 1 M |
| Solvent | Water |
| Max induction | 4 mM |
| Storage | -20 °C |

**Notes**

-AraC is a AraC family activator/repressor from *Escherichia coli;* native gene is present in MG1655 and BL21, including Marionette strains

-AraC$^{AM}$:P$_{BAD}$:Ara induction is 1.5-fold inhibited by 1 mM IPTG

-*E. coli* MG1655 and BL21 can catabolize Ara

#### Plasmid-based regulator

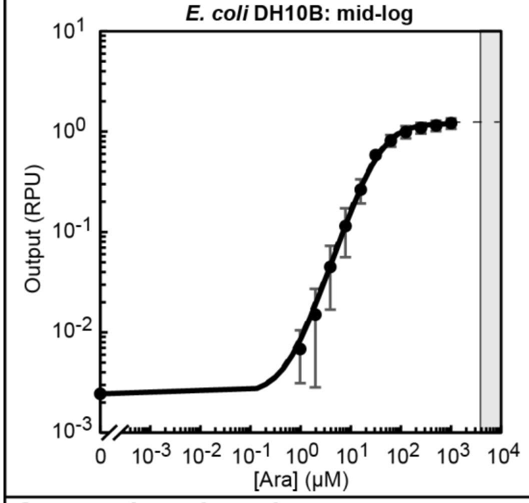

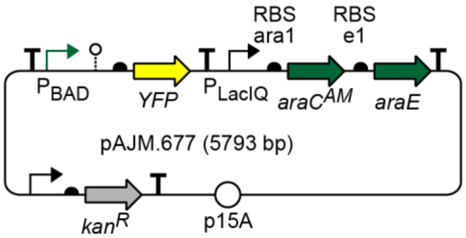

| | |
|---|---|
| Max (RPU) | 1.2 |
| Min (RPU) | 2.4x10$^{-3}$ |
| K (μM) | 3.7x10$^1$ |
| n | 1.5 |

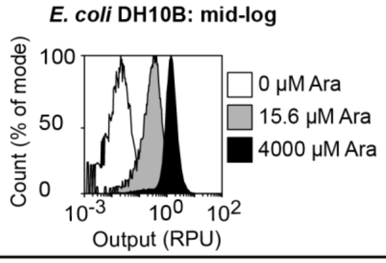

#### Genome-based regulator

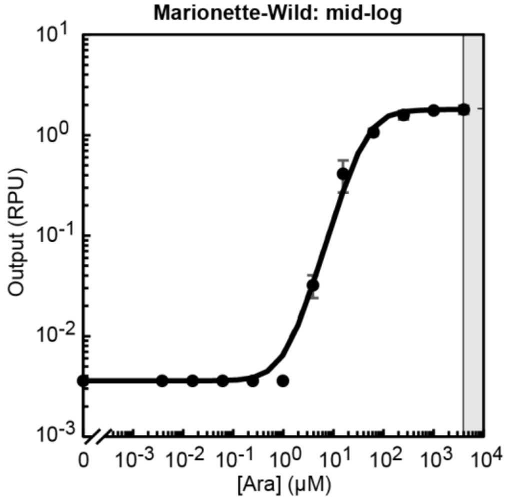

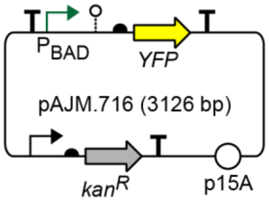

| | |
|---|---|
| Max (RPU) | 1.8 |
| Min (RPU) | 3.6x10$^{-3}$ |
| K (μM) | 4.3x10$^1$ |
| n | 1.7 |

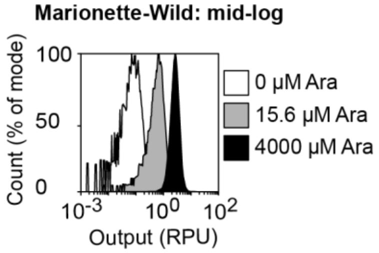

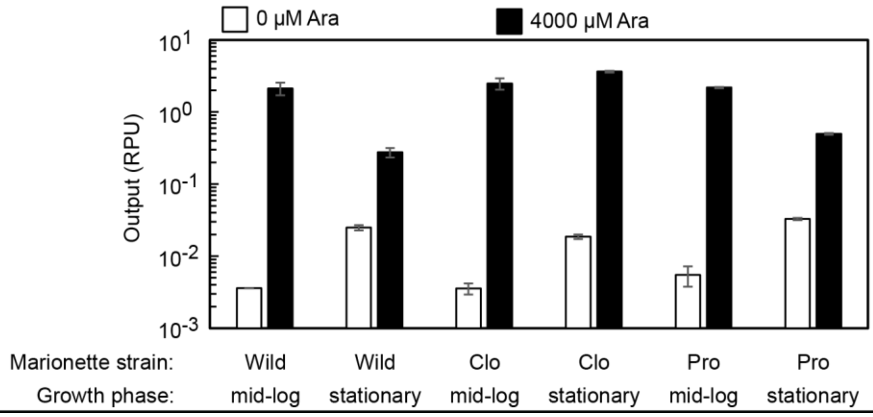

### $BetI^{AM}$:$P_{BetI}$:Cho

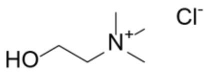

| | |
|---|---|
| Inducer | Choline chloride |
| Source | Sigma C7017 |
| Stock | 1 M |
| Solvent | Water |
| Max induction | 10 mM |
| Storage | -20 °C |

#### Notes

-BetI is a TetR family repressor from *Escherichia coli;* native gene is present in MG1655, DH10B, and BL21, including Marionette strains

-*Escherichia coli* produces Cho, resulting in induction of $P_{BetI}$ in stationary phase

#### Plasmid-based regulator

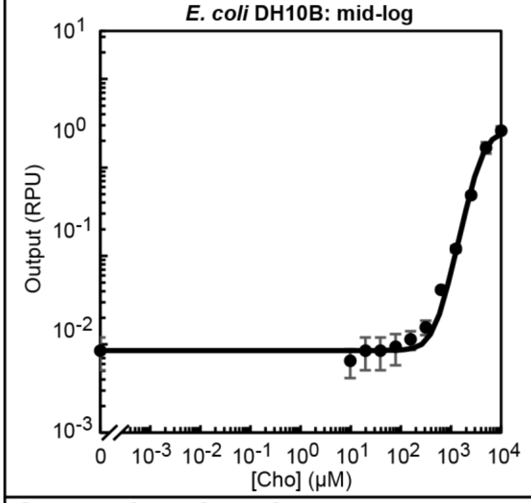

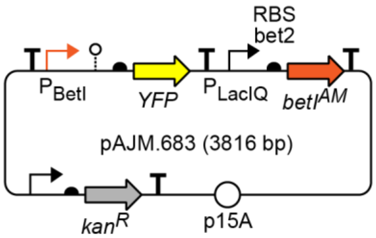

| | |
|---|---|
| Max (RPU) | 2.6 |
| Min (RPU) | $8.5\times10^{-3}$ |
| K (μM) | $4.1\times10^{3}$ |
| n | 2.7 |

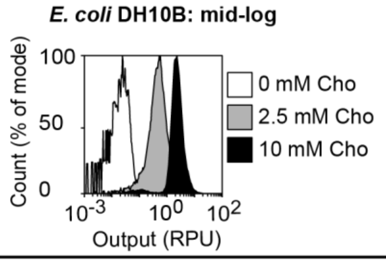

#### Genome-based regulator

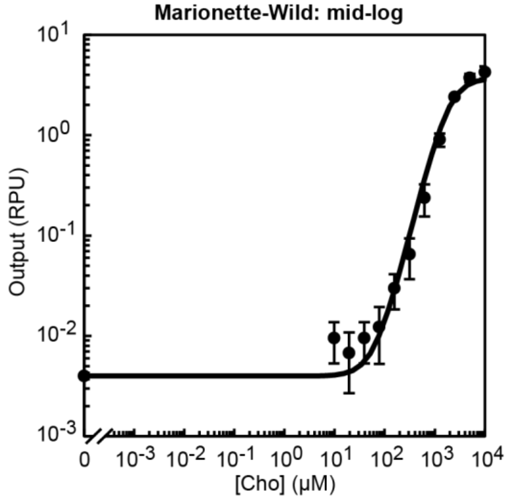

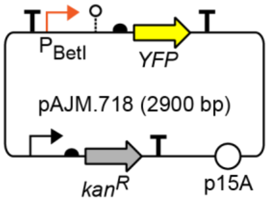

| | |
|---|---|
| Max (RPU) | 3.7 |
| Min (RPU) | $4.0\times10^{-3}$ |
| K (μM) | $1.9\times10^{3}$ |
| n | 2.0 |

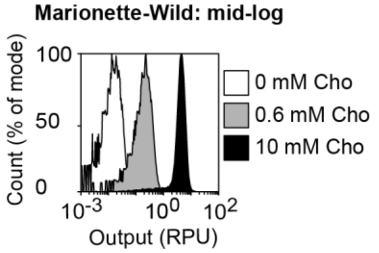

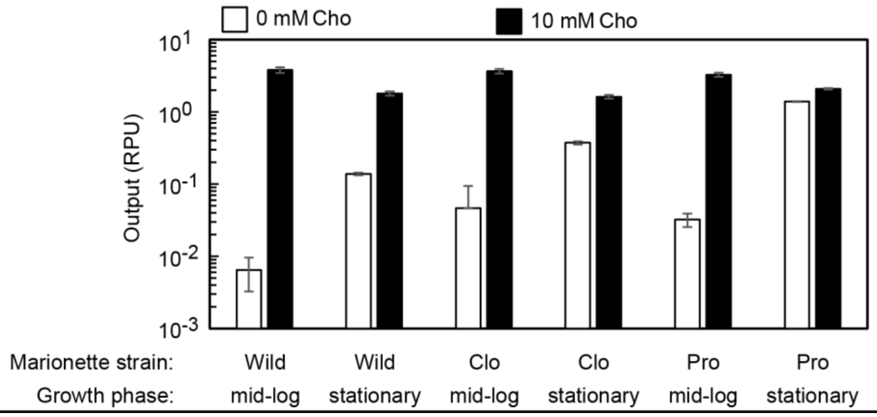

### $TtgR^{AM}$:$P_{Ttg}$:Nar

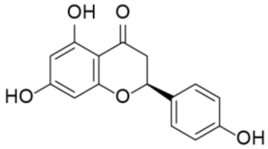

| | |
|---|---|
| Inducer | Naringenin |
| Source | Sigma N5893 |
| Stock | 1 M |
| Solvent | DMSO |
| Max induction | 1 mM |
| Storage | -20 °C |

**Notes**

-TtgR is a TetR family repressor from *Pseudomonas putida*

-Nar forms a precipitate in media; this does not affect induction

#### Plasmid-based regulator

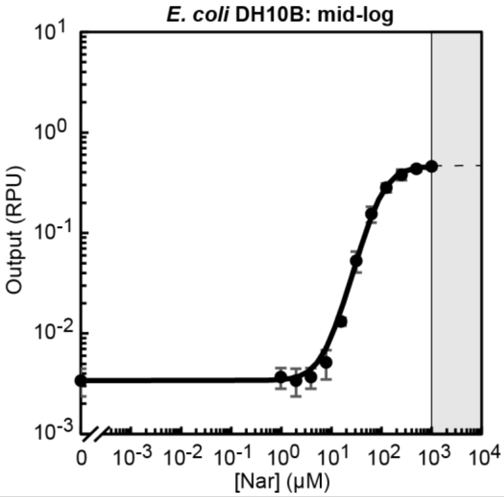

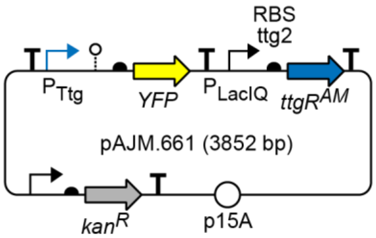

| | |
|---|---|
| Max (RPU) | 0.5 |
| Min (RPU) | $3.4 \times 10^{-3}$ |
| K (μM) | $9.5 \times 10^{1}$ |
| n | 1.9 |

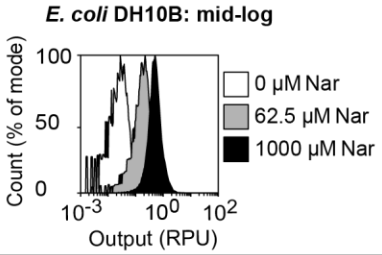

#### Genome-based regulator

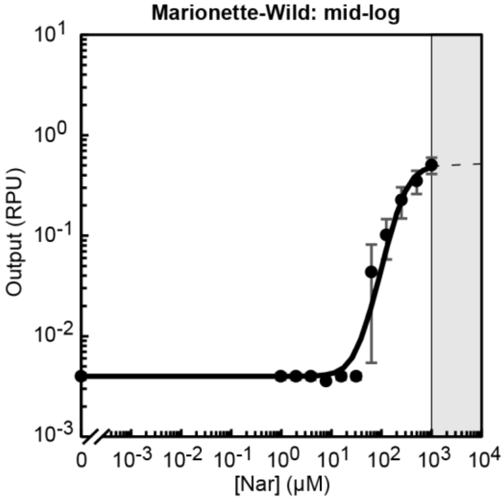

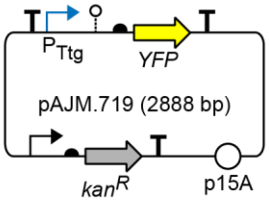

| | |
|---|---|
| Max (RPU) | 0.5 |
| Min (RPU) | $4.0 \times 10^{-3}$ |
| K (μM) | $2.8 \times 10^{2}$ |
| n | 2.3 |

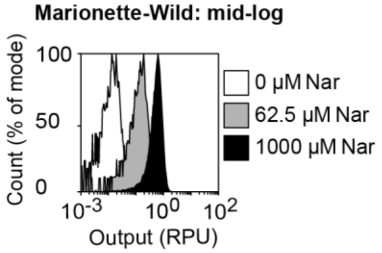

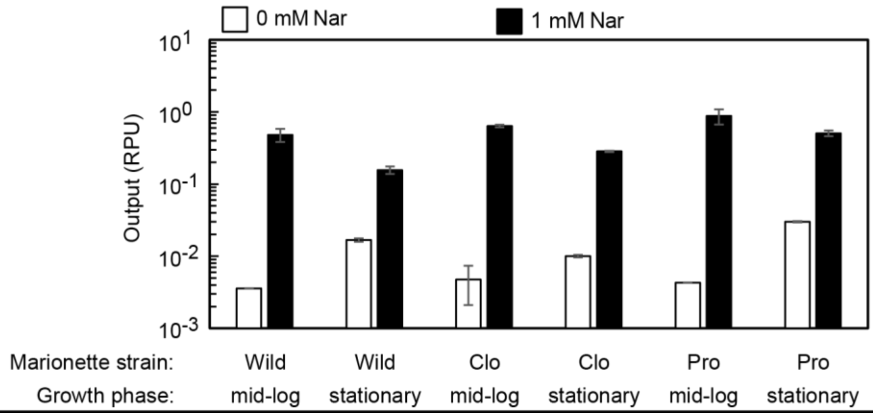

### PcaU$^{AM}$:P$_{3B5}$:DHBA

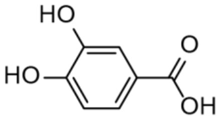

| | |
|---|---|
| Inducer | 3,4-Dihydroxybenzoic acid |
| Source | Sigma 37580 |
| Stock | 1 M |
| Solvent | EtOH |
| Max induction | 1 mM |
| Storage | -20 °C |

**Notes**

-PcaU is a IclR family repressor from *Acinetobacter sp. ADP1*

-PcaU$^{AM}$:P$_{3B5}$ is 2.2-fold induced by 100 μM Van

-VanR$^{AM}$:P$_{VanCC}$ is 1.3-fold induced by 1 mM DHBA

#### Plasmid-based regulator

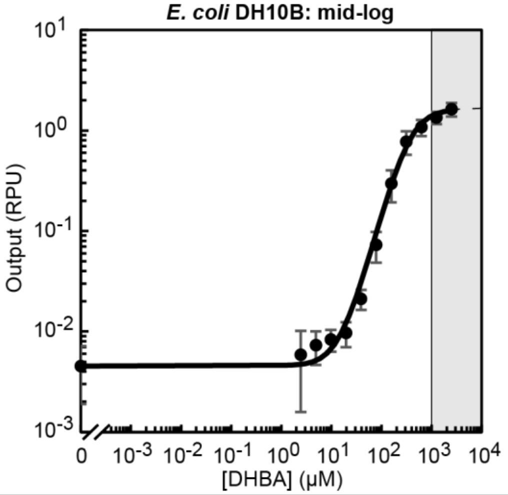

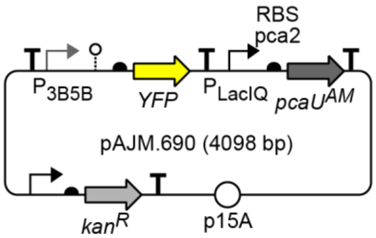

| | |
|---|---|
| Max (RPU) | 1.63 |
| Min (RPU) | 0.0045 |
| K (μM) | 369.22 |
| n | 1.82 |

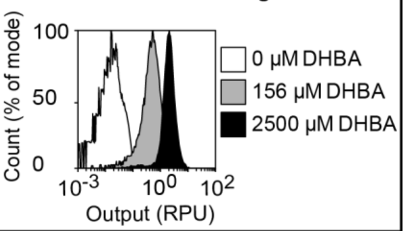

#### Genome-based regulator

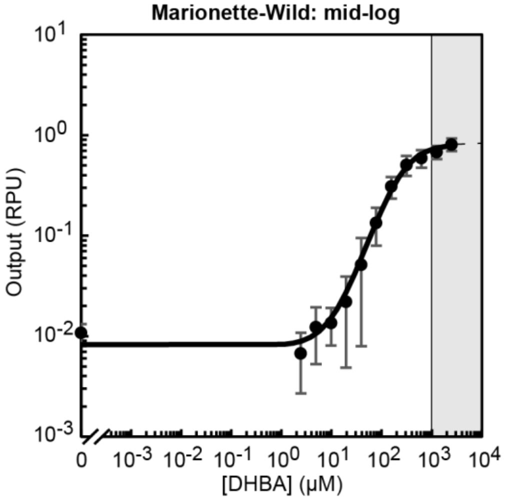

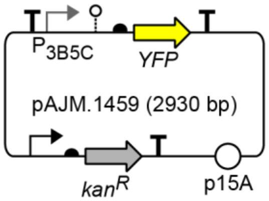

| | |
|---|---|
| Max (RPU) | 0.81 |
| Min (RPU) | 0.0080 |
| K (μM) | 241.69 |
| n | 1.47 |

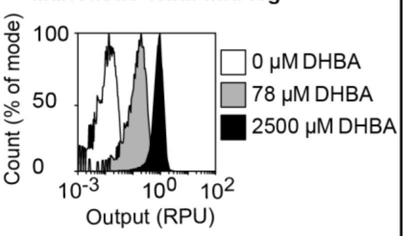

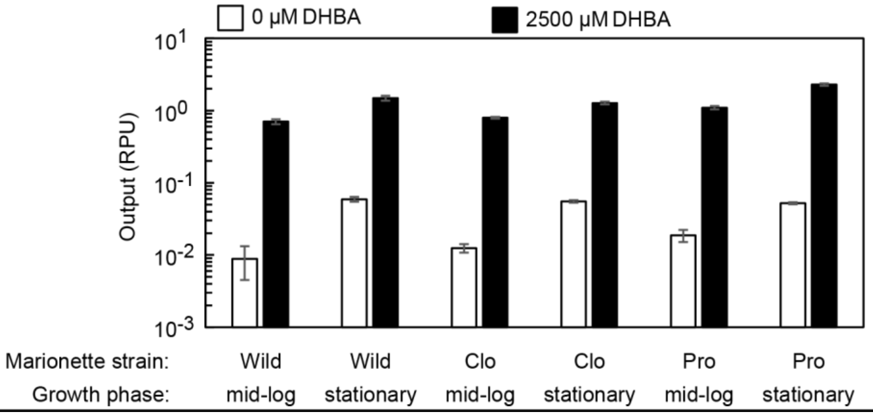

### $NahR^{AM}$:$P_{SalTTC}$:Sal

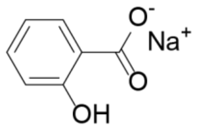

| | |
|---|---|
| Inducer | Sodium salicylate |
| Source | Sigma S3007 |
| Stock | 1 M |
| Solvent | Water |
| Max induction | 100 µM |
| Storage | -20 °C |

#### Notes

-NahR is a LysR family activator from *Pseudomonas putida*
-$PhlF^{AM}$:$P_{PhlF}$ is 2.6-fold induced by 100 µM Sal
-$CymR^{AM}$:$P_{CymRC}$:Cuma induction is 13-fold antagonized by 100 µM Sal

#### Plasmid-based regulator

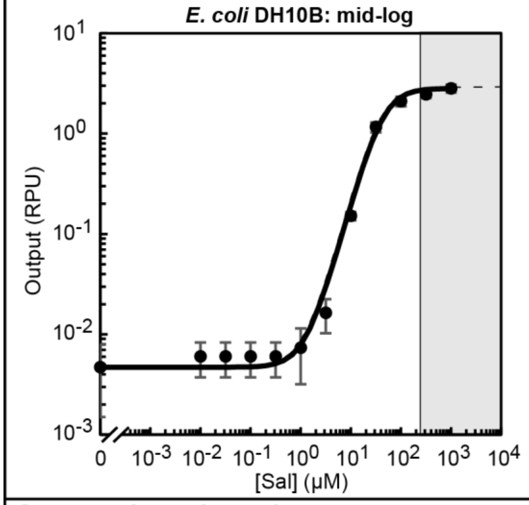

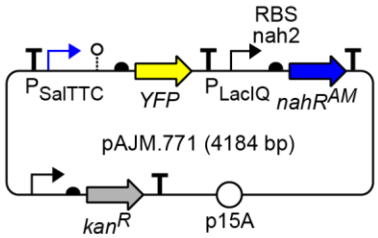

| | |
|---|---|
| Max (RPU) | 2.8 |
| Min (RPU) | $4.7 \times 10^{-3}$ |
| K (µM) | $4.3 \times 10^{1}$ |
| n | 1.8 |

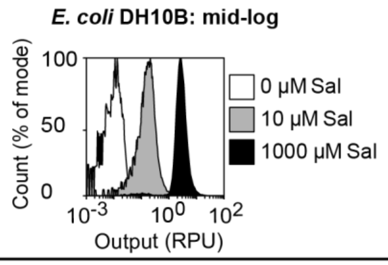

#### Genome-based regulator

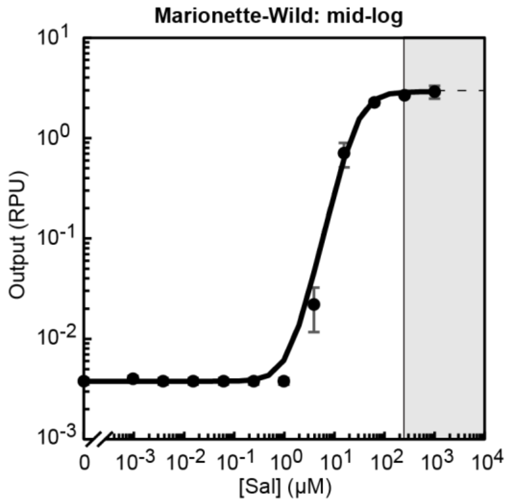

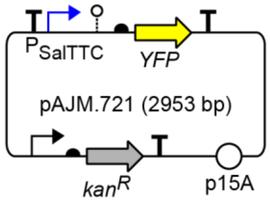

| | |
|---|---|
| Max (RPU) | 2.9 |
| Min (RPU) | $3.8 \times 10^{-3}$ |
| K (µM) | $2.9 \times 10^{1}$ |
| n | 2.1 |

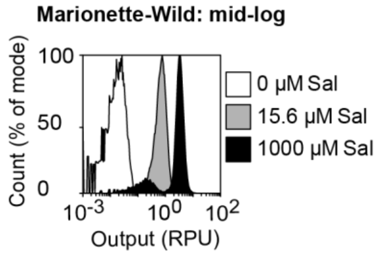

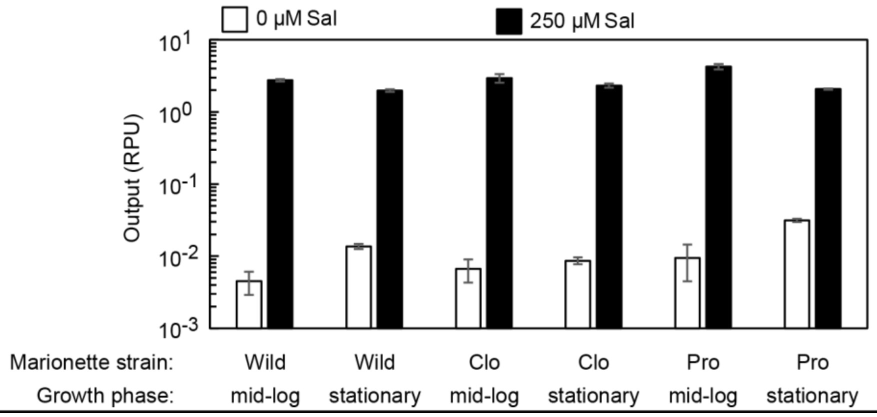

### CinR$^{AM}$:P$_{Cin}$:OHC14

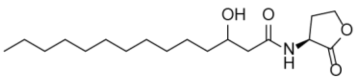

| | |
|---|---|
| Inducer | 3OHC14-HSL |
| Source | Sigma 51481 |
| Stock | 10 mM |
| Solvent | DMF |
| Max induction | 10 µM |
| Storage | -20 °C |

#### Notes

-CinR is a LuxR family activator from *Rhizobium leguminosarum*

#### Plasmid-based regulator

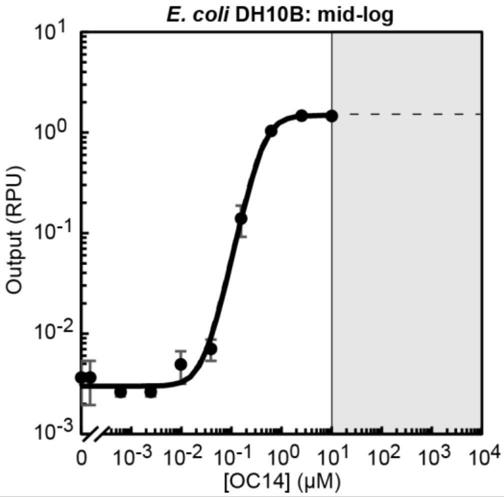

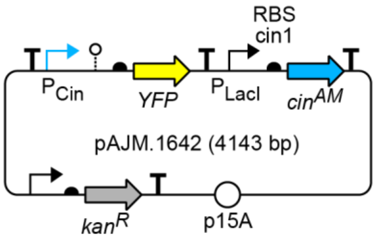

| | |
|---|---|
| Max (RPU) | 1.5 |
| Min (RPU) | $3.0\times10^{-3}$ |
| K (µM) | $4.3\times10^{-1}$ |
| n | 2.3 |

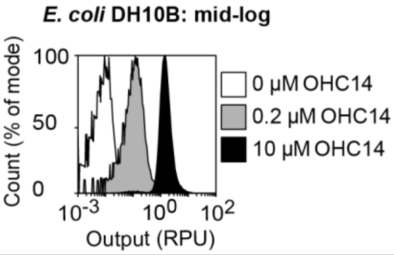

#### Genome-based regulator

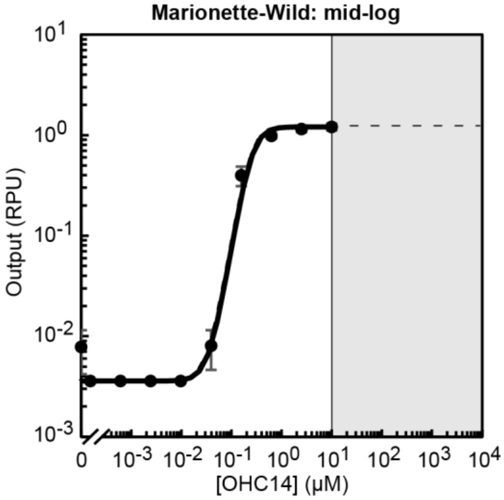

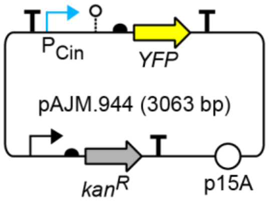

| | |
|---|---|
| Max (RPU) | 1.2 |
| Min (RPU) | $3.6\times10^{-3}$ |
| K (µM) | $2.5\times10^{-1}$ |
| n | 3.0 |

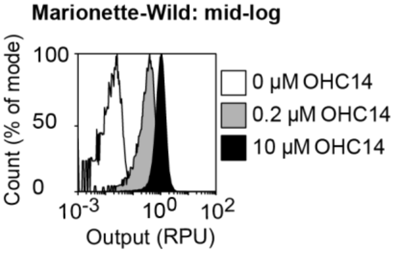

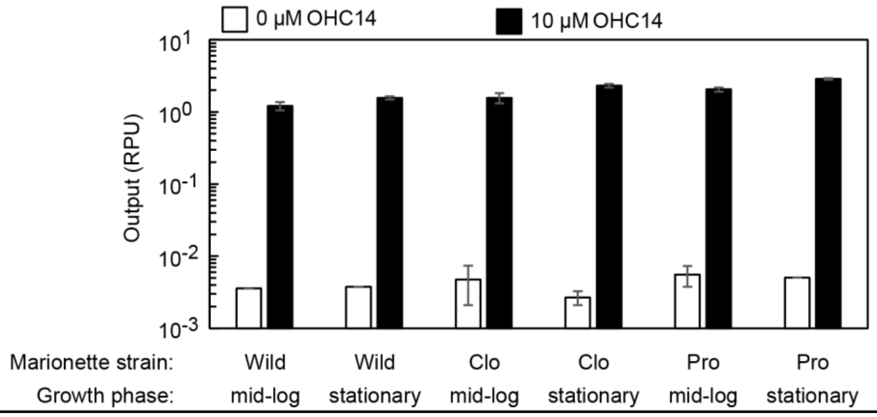

### AcuR$^{AM}$:P$_{Acu}$:Acr

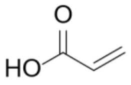

| | |
|---|---|
| Inducer | Acrylic acid |
| Source | Sigma 147230 |
| Stock | 1 M |
| Solvent | Water |
| Max induction | 1 mM |
| Storage | 4 °C |

#### Notes

-AcuR is a TetR family repressor from *Rhodobacter sphaeroides*

#### Plasmid-based regulator

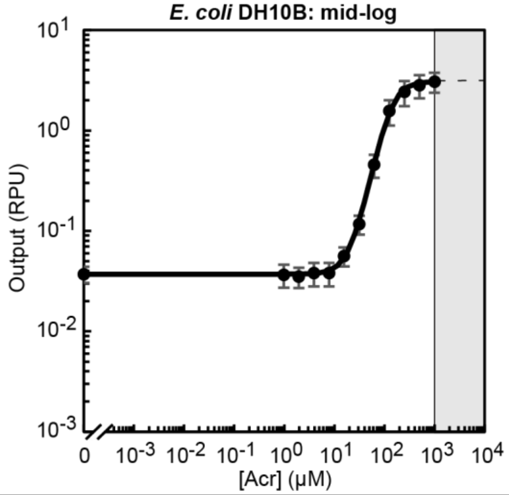

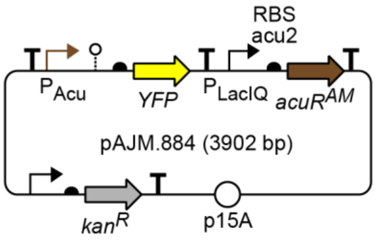

| | |
|---|---|
| Max (RPU) | 3.1 |
| Min (RPU) | $3.7 \times 10^{-2}$ |
| K (μM) | $1.3 \times 10^{2}$ |
| n | 2.5 |

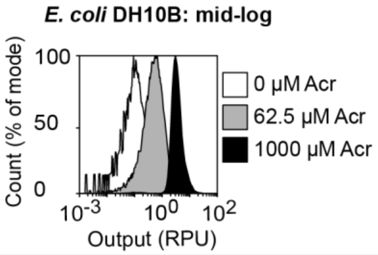

### $MphR^{AM}:P_{Mph}:Ery$

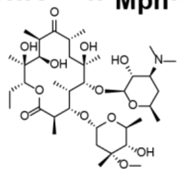

| | |
|---|---|
| Inducer | Erythromycin |
| Source | Sigma E5389 |
| Stock | 100 mM |
| Solvent | EtOH |
| Max induction | 125 µM |
| Storage | 4 °C |

**Notes**

-MphR is a TetR family repressor from *Escherichia coli* clinical isolates; not present in MG1655, DH10B, or BL21

-Ery inhibits translation in *Escherichia coli*

-$Ery^R$ methylates the ribosome, confering Ery resistance up to 125 µM

**Plasmid-based regulator**

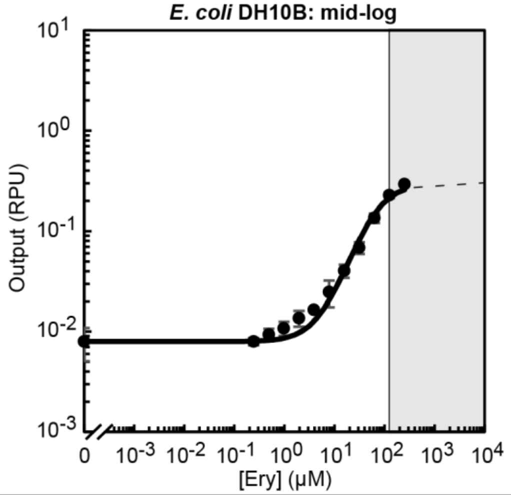

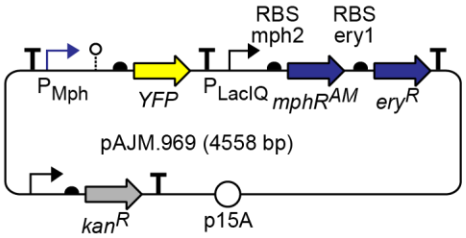

| | |
|---|---|
| Max (RPU) | 0.3 |
| Min (RPU) | $8.0 \times 10^{-3}$ |
| K (µM) | $6.5 \times 10^1$ |
| n | 1.5 |

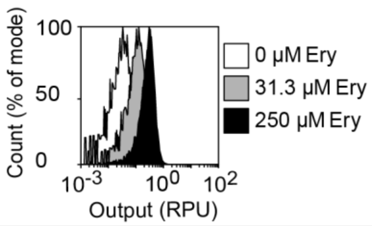

**Supplementary Appendix 2: Sensor evolution**

Each page details the process of directed evolution to optimize 12 sensors.

#### Regulator:$P_{Out}$:Inducer

Left: Coding (red) and non-coding (grey/black) mutations in the evolved sensor are noted. Mutations in grey are also found the parental sensor. Right: response functions of the evolved (blue) and parental (grey) sensors. All data are from *E. coli* DH10B cells in mid-log growth in LB media. The mean of three replicates from different days is shown, and error bars represent the standard deviation.

#### Initial library

Degeneracy (orange) was added to the regulator CDS, regulator RBS, and the output promoter. In the case of LacI, positions were partially randomized using multiple degenerate oligonucleotides mixed at the percentage shown. Black diamonds denote BbsI restriction sites. N=A25:C25:T25:G25; W=A50:T50; S=C50:G50; M=A50:C50; K=G50:T50; D=33A:33G:33T; V=33A:33C:33G.

#### Directed evolution interventions

The selection parameters for each round of selection are noted. R1=Round 1 of selection. DNAP=DNA polymerase (error-prone PK6 vs high-fidelity KOD). "Shuffle" indicates that libraries were subject to gene shuffling prior to the next round of selection.

The first two rounds of the LacI selection utilized the positive selection plasmid and the negative selection plasmid, respectively. All other rounds utilize dual selection plasmids (Supplementary Figure 6).

#### On/off screen

RPU values for each clone with inducer(s) added as noted. All data are from *E. coli* DH10B cells in mid-log growth in LB media. A single replicate is shown.

#### Sequencing

Non-synonymous mutations are listed as amino acid changes while synonymous mutations are not listed (*i.e.,* Q18M denotes that the 18$^{th}$ amino acid mutated from glutamine to methionine). Blue denotes that the exact mutation appears in the final mutant (Regulator$^{AM}$). * = stop codon.

#### Consensus mutant testing

Response functions for parental (WT) sensors, evolved sensors, and notable sensors from the literature are shown. Where relevant, response functions with cross-reactive inducers or antagonistic inducers are shown. Nonsynonymous mutations are shown. Full sequences of promoters and proteins (parental and evolved) are provided in Supplementary Tables 1 and 4. All data are from *E. coli* DH10B cells in mid-log growth in LB media. The mean of three replicates from different days is shown, and error bars represent the standard deviation.

### PhlF:$P_{PhlF}$:DAPG

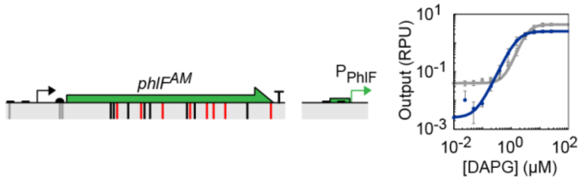

### Directed evolution interventions

| PhlF | [Cl-Phe] | [Negative ligand] | DNAP | [Positive ligand] | Notes |
|---|---|---|---|---|---|
| R1 | 2 mM | | PK6 | 2.5 μM DAPG | |
| R2 | 2 mM | | PK6 | 2.5 μM DAPG | |
| R3 | 2 mM | | PK6 | 2.5 μM DAPG | |
| R4 | 2 mM | | PK6 | 2.5 μM DAPG | |
| R5 | 4 mM | | PK6 | 2.5 μM DAPG | |
| R6 | 4 mM | | PK6 | 2.5 μM DAPG | |
| R7 | 4 mM | | KOD | 2.5 μM DAPG | |
| R8 | 4 mM | | KOD | 2.5 μM DAPG | |
| R9 | 4 mM | 10 μM Sal | PK6 | 10 μM Sal + 2.5 μM DAPG | |
| R10 | 4 mM | 10 μM Sal | PK6 | 10 μM Sal + 2.5 μM DAPG | |
| R11 | 4 mM | 10 μM Sal | PK6 | 10 μM Sal + 2.5 μM DAPG | |
| R12 | 4 mM | 20 μM Sal | PK6 | 20 μM Sal + 2.5 μM DAPG | |
| R13 | 4 mM | 30 μM Sal | PK6 | 30 μM Sal + 2.5 μM DAPG | Shuffle |
| R14 | 4 mM | 40 μM Sal | PK6 | 40 μM Sal + 2.5 μM DAPG | |
| R15 | 4 mM | 40 μM Sal | PK6 | 40 μM Sal + 2.5 μM DAPG | |
| R16 | 4 mM | 50 μM Sal | PK6 | 50 μM Sal + 2.5 μM DAPG | |
| R17 | 4 mM | 60 μM Sal | PK6 | 60 μM Sal + 2.5 μM DAPG | |
| R18 | 4 mM | 75 μM Sal | PK6 | 2.5 μM DAPG | |
| R19 | 4 mM | 100 μM Sal | KOD | 2.5 μM DAPG | |
| R20 | 4 mM | 100 μM Sal | KOD | 100 μM Sal + 2.5 μM DAPG | |
| R21 | 4 mM | 100 μM Sal | KOD | 100 μM Sal + 2.5 μM DAPG | Shuffle |
| R22 | 4 mM | 100 μM Sal | KOD | 100 μM Sal + 100 μM Van + 2.5 μM DAPG | |
| R23 | 4 mM | 100 μM Sal | KOD | 100 μM Sal + 100 μM Van + 2.5 μM DAPG | |

### Initial library (889 bp)

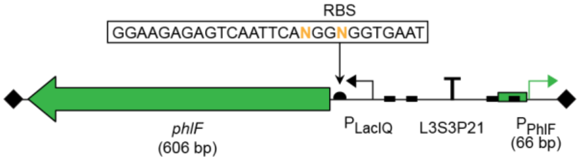

### On/off screen

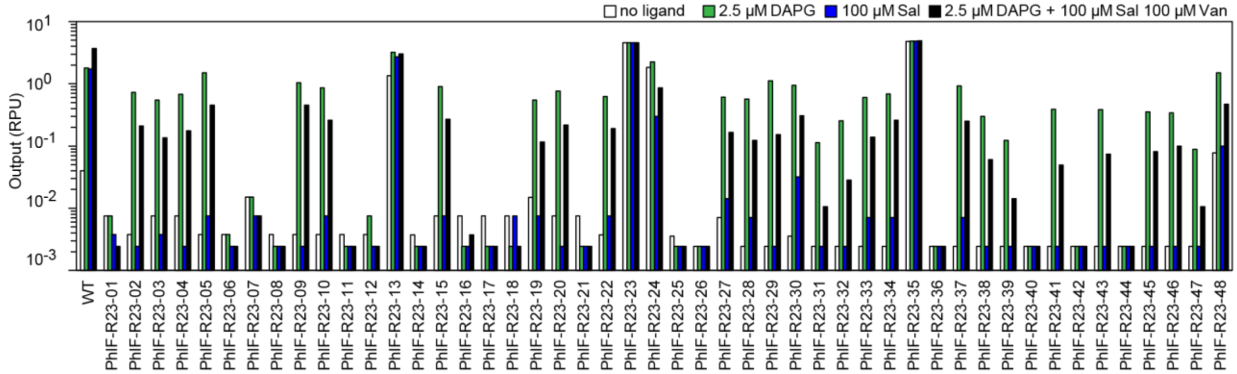

### Sequencing

| Mutant | Mutations |
|---|---|
| PhlF-R23-02 | S50G, S73I, R94H, A121V, M137I, G197D |
| PhlF-R23-05 | S50G, S73I, C111S, A121V, Q125L, M137I, G197C |
| PhlF-R23-09 | T4I, S50G, S73I, R94H, A121V, M137I, N144D, R200C |
| PhlF-R23-10 | S50G, S73I, A121V, M137I |
| PhlF-R23-15 | S50G, S73I, A121V, M137I |
| PhlF-R23-20 | S50G, S73I, R94C, C111S, A121V, Q125L, M137I, V179I, G197R |
| PhlF-R23-22 | S50G, S73I, M137I, K153E, T155I |
| PhlF-R23-28 | L38M, S50G, S73I, A121V, M137I, G197D, R200C |
| PhlF-R23-33 | L38M, S50G, S73I, M137I, T155I |
| PhlF-R23-34 | T4I, S50G, S73I, A121V, T124A, M137I, N144I, S148N |
| PhlF-R23-37 | S50G, S73I, R94H, L123M, M137I, T155I, P196Q |
| PhlF-R23-46 | S50G, S73I, M137I, V179A |

### Consensus mutant testing

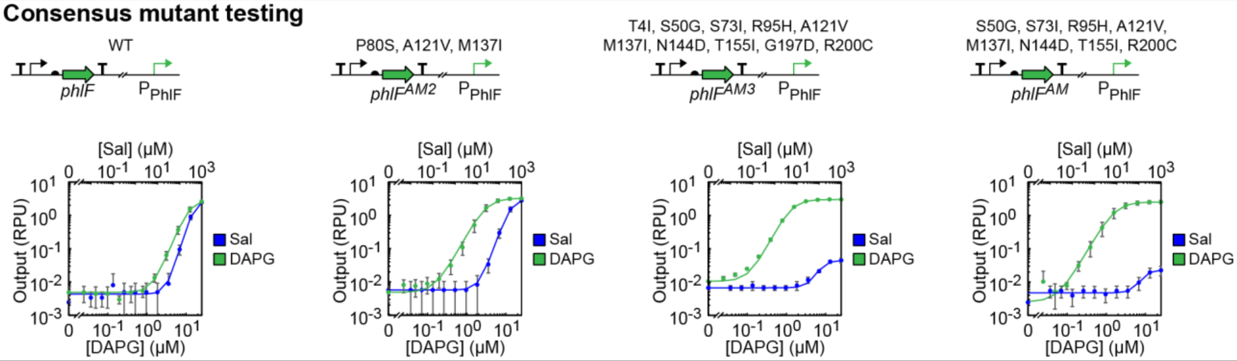

### CymR:$P_{CymR}$:Cuma

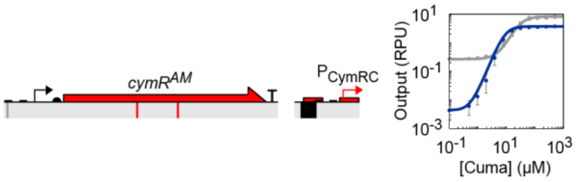

### Directed evolution interventions

| CymR | [Cl-Phe] | [Negative ligand] | DNAP | [Positive ligand] | Notes |
|---|---|---|---|---|---|
| R1 | 2 mM | | PK6 | 50 μM Cuma | |
| R2 | 2 mM | | PK6 | 50 μM Cuma | |
| R3 | 4 mM | | PK6 | 50 μM Cuma | |
| R4 | 4 mM | 10 μM Sal | PK6 | 10 μM Sal + 50 μM Cuma | |
| R5 | 4 mM | 10 μM Sal | PK6 | 10 μM Sal + 50 μM Cuma | |
| R6 | 4 mM | 20 μM Sal | PK6 | 20 μM Sal + 50 μM Cuma | |
| R7 | 4 mM | 30 μM Sal | PK6 | 30 μM Sal + 50 μM Cuma | |
| R8 | 4 mM | 40 μM Sal | PK6 | 40 μM Sal + 50 μM Cuma | |
| R9 | 4 mM | 50 μM Sal | PK6 | 50 μM Sal + 50 μM Cuma | |
| R10 | 4 mM | 75 μM Sal | PK6 | 75 μM Sal + 50 μM Cuma | |
| R11 | 4 mM | | KOD | 100 μM Sal + 50 μM Cuma | |
| R12 | 4 mM | 100 μM Sal | KOD | 100 μM Sal + 50 μM Cuma | |
| R13 | 4 mM | | KOD | 100 μM Sal + 50 μM Cuma | Shuffle |
| R14 | 4 mM | | KOD | 100 μM Sal + 50 μM Cuma | |
| R15 | 4 mM | | KOD | 100 μM Sal + 50 μM Cuma | |

### Initial library (913 bp)

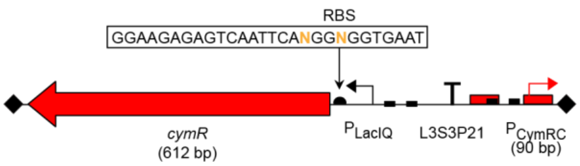

### On/off screen

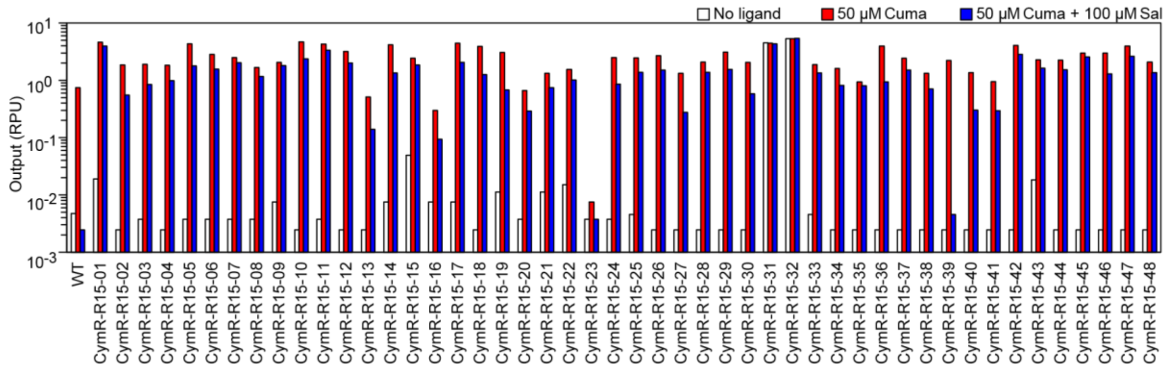

### Sequencing

| Mutant | Mutations |
|---|---|
| CymR-R15-07 | Q16R, A21T, A38V, L69R, E100D, S110G, Q128L, G144N, A171V |
| CymR-R15-09 | A38V, L69R, E100D, S110G, P121Q, Q128L, V138I, G144N, A171V |
| CymR-R15-11 | S110G, P121L, Q128L, A171T, R195H |
| CymR-R15-17 | A79V, S110G, A122V, A171T |
| CymR-R15-28 | V26I, L69R, E75D, E100D, S110G, R129H, R153H, A171V |
| CymR-R15-35 | G25A, L69R, A79V, E100D, S110G, Q128L, A171V |
| CymR-R15-42 | S110G, M141I, A171T, S189N |
| CymR-R15-44 | A43V, L69R, E100D, S110G, A171V, V172I |
| CymR-R15-45 | A43V, L69R, E100D, S110G, A171V, V172I, A199V |
| CymR-R15-47 | A23S, S110G, E125K, A171T |
| CymR-R15-48 | A38V, L69R, E100D, S110G, R133H, A171V, K200R |

### Consensus mutant testing

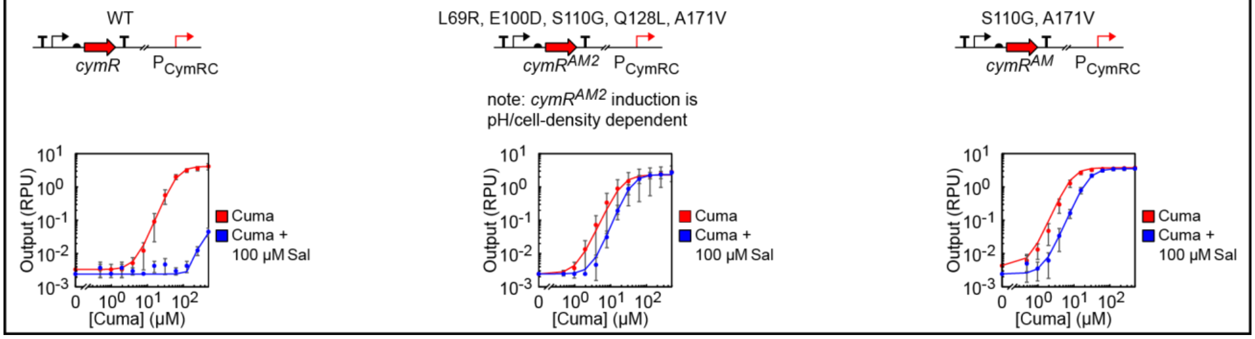

### VanR:$P_{Van}$:Van

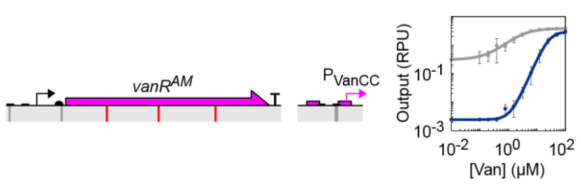

### Directed evolution interventions

| VanR | [Cl-Phe] | [Negative ligand] | DNAP | [Positive ligand] | Notes |
|---|---|---|---|---|---|
| R1 | 2 mM | | PK6 | 10 μM Van | |
| R2 | 2 mM | | PK6 | 10 μM Van | |
| R3 | 2 mM | | PK6 | 10 μM Van | |
| R4 | 2 mM | | PK6 | 10 μM Van | |
| R5 | 4 mM | | PK6 | 10 μM Van | |
| R6 | 4 mM | | PK6 | 10 μM Van | |
| R7 | 4 mM | | KOD | 10 μM Van | |
| R8 | 4 mM | | KOD | 10 μM Van | |
| R9 | 4 mM | 625 μM DHBA | KOD | 10 μM Van | Shuffle |
| R10 | 4 mM | 1250 μM DHBA | KOD | 10 μM Van | |
| R11 | 4 mM | 2500 μM DHBA | KOD | 10 μM Van | |

### Initial library (1006 bp)

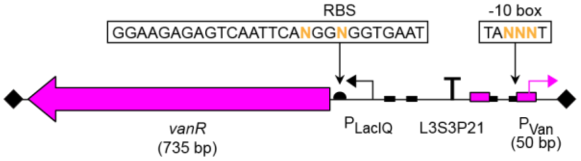

### On/off screen

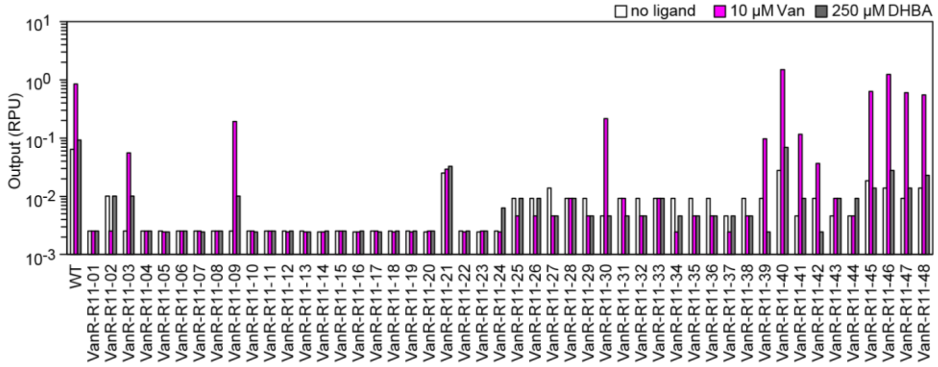

### Sequencing

| Mutant | Mutations |
|---|---|
| VanR-R11-03 | M3I, T49I, A93T, P179T E205K |
| VanR-R11-09 | T49I, R108C, S155G |
| VanR-R11-30 | T49I, A97T |
| VanR-R11-39 | T49I, H106N, A107T, A111V, P179S, E205K |
| VanR-R11-40 | R5H, T49I, A121T L144M, A187V |
| VanR-R11-41 | T49I, A111V, P179S |
| VanR-R11-45 | P4S, T49I, A103T, A118V |
| VanR-R11-46 | R5H, T49I |
| VanR-R11-47 | T49I, A107T, A111V, N126D, E226, A229T |
| VanR-R11-48 | A15T, T49I, A111V, A187V, V195I, A238T |

### Consensus mutant testing

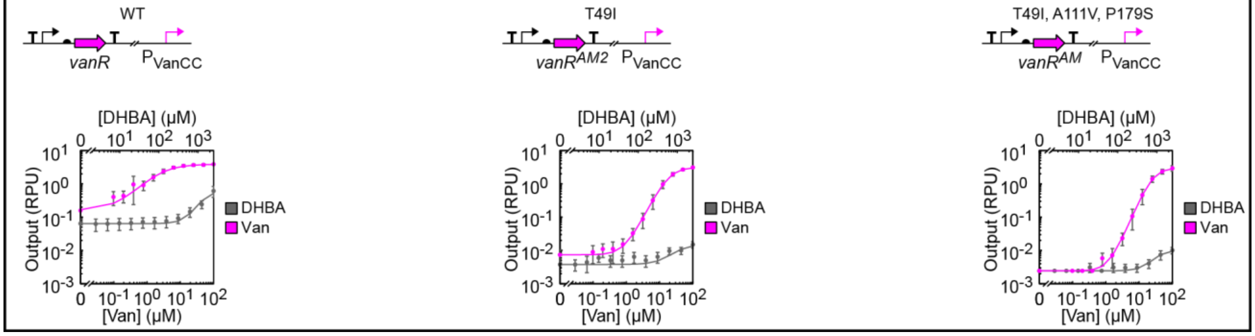

### LacI:$P_{Tac}$:IPTG

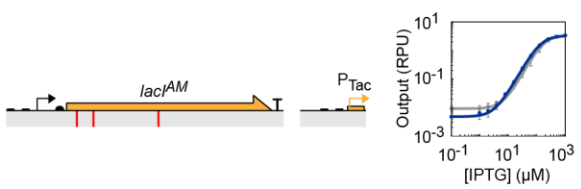

### Directed evolution interventions

| LacI | [Cl-Phe] | [Negative ligand] | DNAP | [Positive ligand] | Notes |
|---|---|---|---|---|---|
| R1 | - | | PK6 | 50 μM IPTG | Postive selection |
| R2 | - | 0 μM IPTG | PK6 | - | Negative selection |
| R3 | 4 mM | | PK6 | 50 μM IPTG | Dual selection |
| R4 | 4 mM | | PK6 | 50 μM IPTG | Dual selection |
| R5 | 4 mM | | PK6 | 50 μM IPTG | Dual selection |
| R6 | 4 mM | | KOD | 50 μM IPTG | Dual selection |
| R7 | 4 mM | | KOD | 50 μM IPTG | Dual selection |

### Initial library (1359 bp)

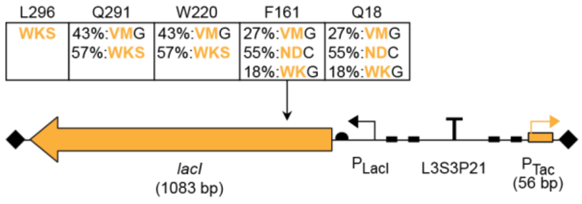

### On/off screen

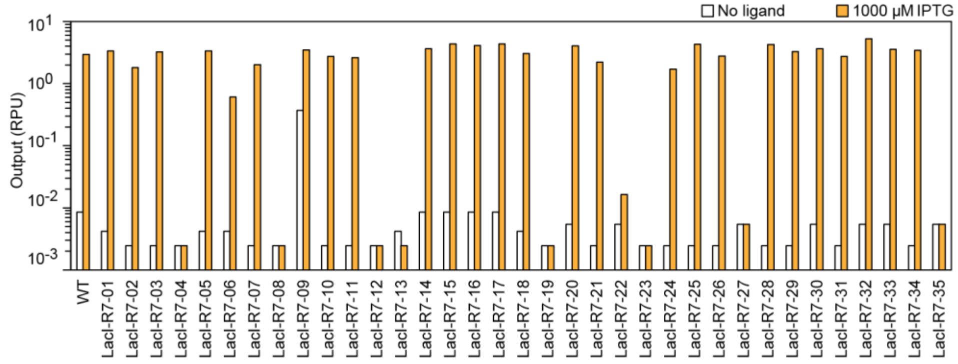

### Sequencing

| Mutant | Q18 | A43 | F161 | W220 | Q291 | L296 |
|---|---|---|---|---|---|---|
| LacI-R7-01 | Q | A | W | W | Q | M |
| LacI-R7-03 | M | V | Y | W | Q | L |
| LacI-R7-05 | M | A | Y | W | Q | M |
| LacI-R7-07 | Q | A | Y | W | Q | L |
| LacI-R7-10 | M | V | Y | W | Q | L |
| LacI-R7-11 | Q | A | Y | W | Q | L |
| LacI-R7-14 | Q | A | Y | W | Q | L |
| LacI-R7-16 | Q | A | Y | W | Q | L |
| LacI-R7-17 | Q | A | Y | W | Q | L |
| LacI-R7-18 | Q | A | Y | W | Q | L |
| LacI-R7-20 | Q | A | Y | W | Q | L |
| LacI-R7-21 | Q | A | Y | W | Q | L |
| LacI-R7-25 | M | V | Y | W | Q | L |
| LacI-R7-28 | M | V | Y | W | Q | L |
| LacI-R7-29 | Q | A | Y | W | Q | L |
| LacI-R7-30 | Q | A | Y | W | Q | L |
| LacI-R7-31 | H | A | Y | W | Q | I |
| LacI-R7-32 | Q | V | Y | W | Q | L |
| LacI-R7-33 | Q | V | W | W | Q | I |

### Consensus mutant testing

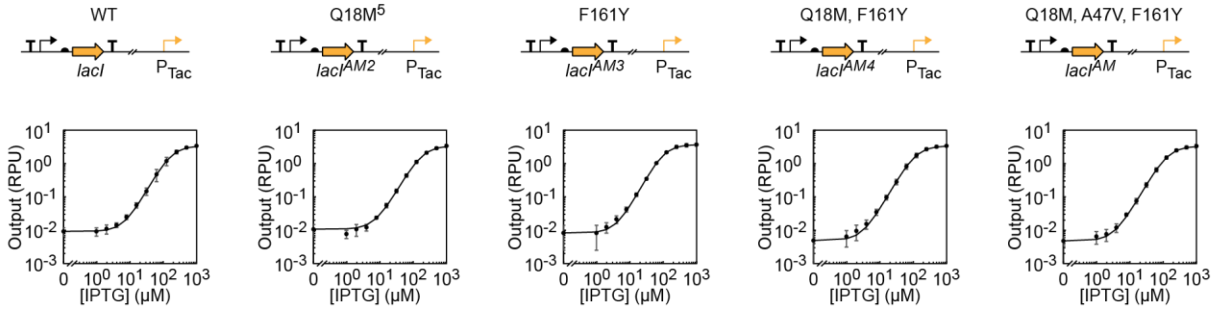

### AraC:P$_{BAD}$:Ara

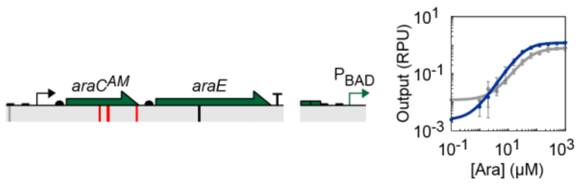

### Directed evolution interventions

| AraC | [Cl-Phe] | [Negative ligand] | DNAP | [Positive ligand] | Notes |
|---|---|---|---|---|---|
| R1 | 2 mM | 1000 μM IPTG | PK6 | 1000 μM IPTG + 50 μM Ara | |
| R2 | 2 mM | 1000 μM IPTG | PK6 | 1000 μM IPTG + 50 μM Ara | |
| R3 | 4 mM | 1000 μM IPTG | PK6 | 1000 μM IPTG + 50 μM Ara | |
| R4 | 4 mM | 1000 μM IPTG | PK6 | 1000 μM IPTG + 50 μM Ara | |

### Initial library (2848 bp)

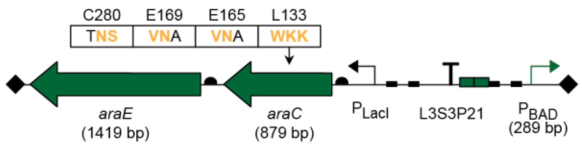

### On/off screen

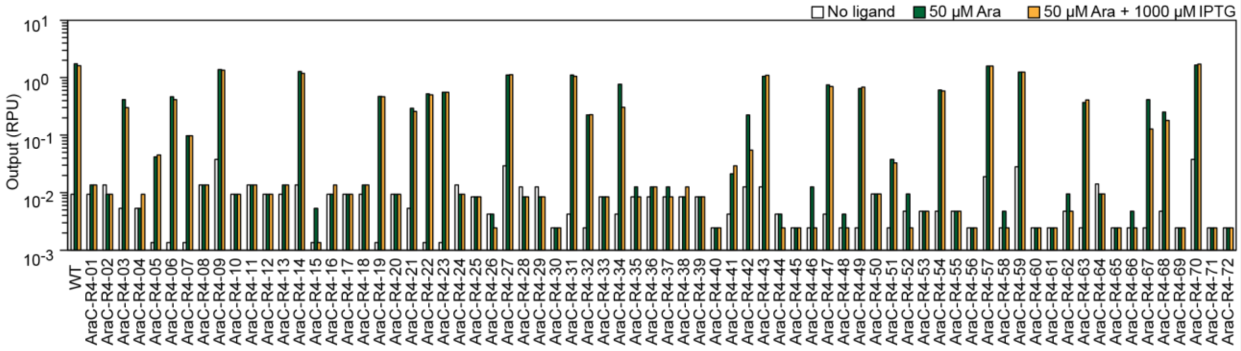

### Sequencing

| Mutant | L133 | E165 | E169 | C280 |
|---|---|---|---|---|
| AraC-R4-06 | R | L | K | W |
| AraC-R4-14 | L | R | G | C |
| AraC-R4-19 | I | P | Q | C |
| AraC-R4-21 | F | A | A | S |
| AraC-R4-22 | S | R | G | L |
| AraC-R4-23 | W | G | I | L |
| AraC-R4-31 | F | L | G | L |
| AraC-R4-32 | R | I | L | W |
| AraC-R4-43 | M | E | G | W |
| AraC-R4-47 | M | R | A | F |
| AraC-R4-49 | F | L | L | Y |
| AraC-R4-54 | S | V | V | L |
| AraC-R4-57 | I | V | E | F |
| AraC-R4-63 | W | G | E | L |

### Consensus mutant testing

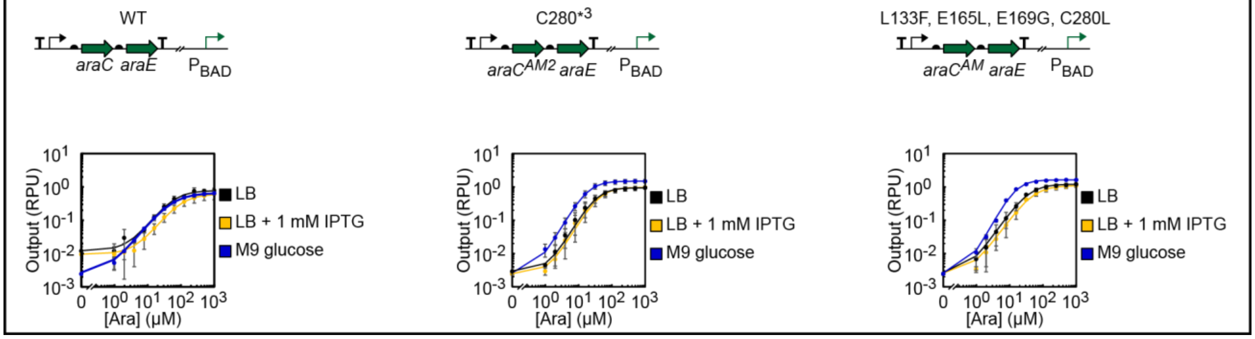

### BetI:$P_{BetI}$:Cho

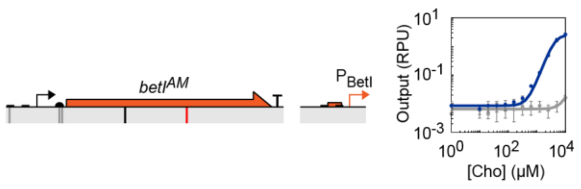

### Directed evolution interventions

| BetI | [Cl-Phe] [Negative ligand] | DNAP | [Positive ligand] | Notes |
|---|---|---|---|---|
| R1 | 2 mM | PK6 | 5 mM Cho | |
| R2 | 2 mM | PK6 | 5 mM Cho | |
| R3 | 4 mM | PK6 | 5 mM Cho | |
| R4 | 4 mM | PK6 | 4 mM Cho | |
| R5 | 4 mM | PK6 | 3 mM Cho | |
| R6 | 4 mM | PK6 | 2 mM Cho | |
| R7 | 4 mM | KOD | 1 mM Cho | |
| R8 | 4 mM | KOD | 1 mM Cho | |

### Initial library (871 bp)

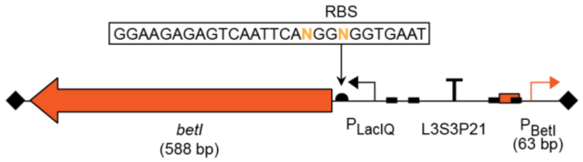

### On/off screen

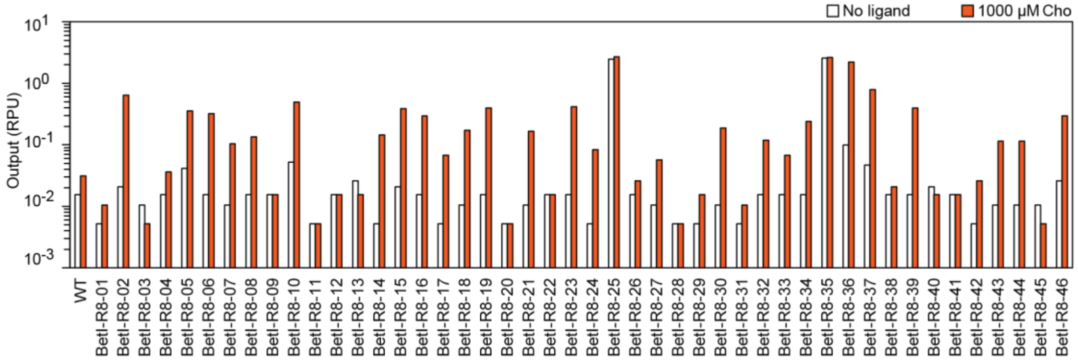

### Sequencing

| Mutant | Mutations |
|---|---|
| BetI-R8-02 | M121I, Q127H, A180V |
| BetI-R8-06 | R37C, S115I |
| BetI-R8-14 | S115I, D195N, *196RYGSA* |
| BetI-R8-15 | S115I, K177E, H186Q |
| BetI-R8-16 | S115I, K173E |
| BetI-R8-19 | D70N, S115N, A180V |
| BetI-R8-23 | H77Y, S115I |
| BetI-R8-30 | G93D, Q149H, A180V, R185H |
| BetI-R8-36 | I32V, S115I, S182N, T189I |
| BetI-R8-39 | R51H, H77L, S102N, A180V, D195N, *196G* |

### Consensus mutant testing

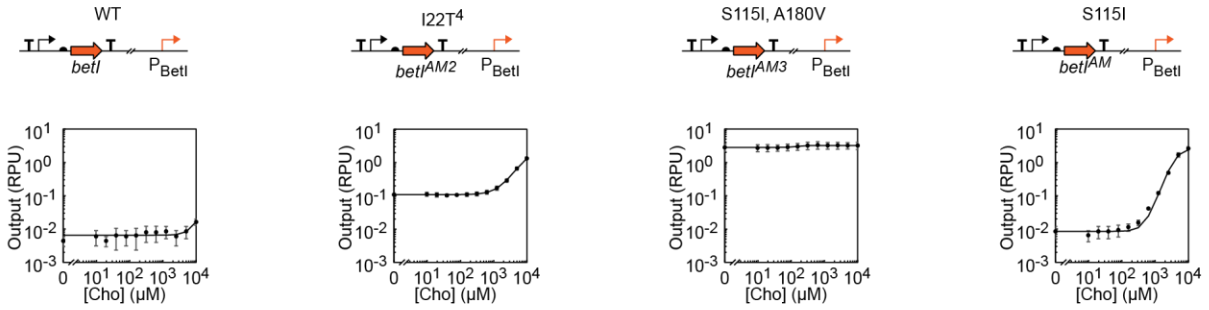

### TtgR:$P_{Ttg}$:Nar

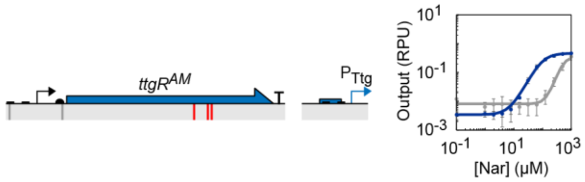

### Directed evolution interventions

| TtgR | [Cl-Phe] | [Negative ligand] | DNAP | [Positive ligand] | Notes |
|---|---|---|---|---|---|
| R1 | 2 mM | | PK6 | 250 µM Nar | |
| R2 | 2 mM | | PK6 | 250 µM Nar | |
| R3 | 4 mM | | PK6 | 250 µM Nar | |
| R4 | 4 mM | | PK6 | 250 µM Nar | |
| R5 | 4 mM | | PK6 | 250 µM Nar | |
| R6 | 4 mM | | PK6 | 250 µM Nar | |
| R7 | 4 mM | | KOD | 250 µM Nar | |
| R8 | 4 mM | | KOD | 250 µM Nar | |

### Initial library (904 bp)

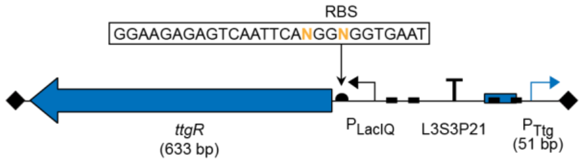

### On/off screen

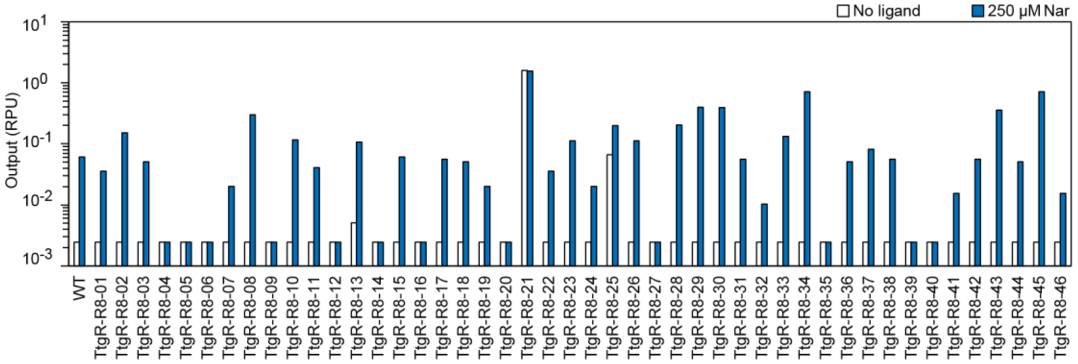

### Sequencing

| Mutant | Mutations |
|---|---|
| TtgR-R8-08 | R90H, R130H, A144T |
| TtgR-R8-29 | A61T, H67L, G140A, E161G |
| TtgR-R8-30 | A61T, R130C, A144T, R151H, K193N, A207T |
| TtgR-R8-34 | I17V, H67R, D71N, A144T, A146V, A148V |
| TtgR-R8-43 | R130H, A148V |
| TtgR-R8-45 | S79N, R130H, A144T, A148V, *211Q* |

### Consensus mutant testing

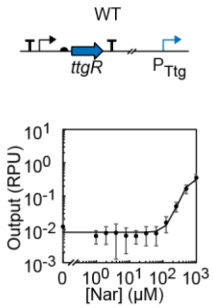

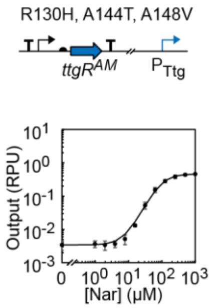

### PcaU:$P_{3B5}$:DHBA

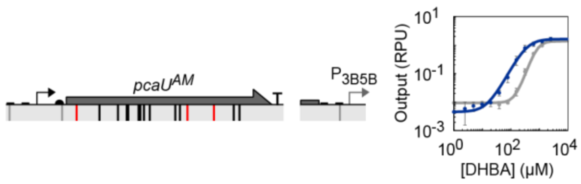

### Initial library (1153 bp)

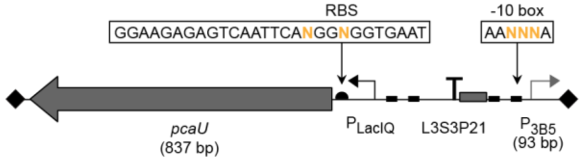

### Directed evolution interventions

| PcaU | [Cl-Phe] | [Negative ligand] | DNAP | [Positive ligand] | Notes |
|---|---|---|---|---|---|
| R1 | 2 mM | | PK6 | 500 μM DHBA | |
| R2 | 2 mM | | PK6 | 500 μM DHBA | |
| R3 | 4 mM | | PK6 | 500 μM DHBA | |
| R4 | 4 mM | 10 μM Van | PK6 | 10 μM Van + 250 μM DHBA | |
| R5 | 4 mM | 20 μM Van | PK6 | 20 μM Van + 250 μM DHBA | |
| R6 | 4 mM | 20 μM Van | PK6 | 20 μM Van + 250 μM DHBA | |
| R7 | 4 mM | 30 μM Van | PK6 | 30 μM Van + 250 μM DHBA | |
| R8 | 4 mM | 40 μM Van | PK6 | 40 μM Van + 250 μM DHBA | |
| R9 | 4 mM | 50 μM Van | PK6 | 50 μM Van + 250 μM DHBA | |
| R10 | 4 mM | 75 μM Van | PK6 | 250 μM DHBA | |
| R11 | 4 mM | 100 μM Van | KOD | 250 μM DHBA | |
| R12 | 4 mM | 100 μM Van | KOD | 100 μM Van + 250 μM DHBA | |
| R13 | 4 mM | 100 μM Van | KOD | 250 μM DHBA | Shuffle |
| R14 | 4 mM | 100 μM Van | KOD | 200 μM DHBA | |
| R15 | 4 mM | 100 μM Van | KOD | 100 μM DHBA | |

### On/off screen

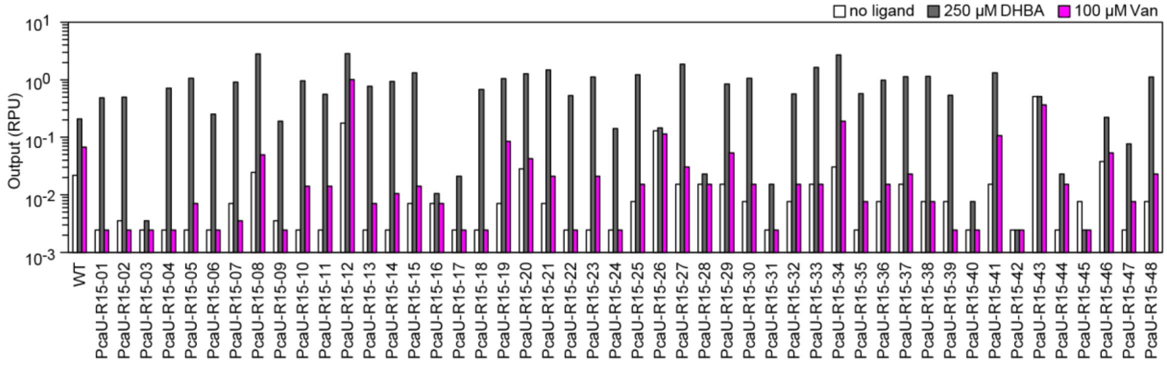

### Sequencing

| Mutant | Mutations |
|---|---|
| PcaU-R11-04 | K14N, A165T, N201H |
| PcaU-R15-01 | G102S, A165T, K212N, K256E |
| PcaU-R15-02 | G98A, G102S, P107S, P162S, V241I |
| PcaU-R15-04 | A165T, K212N |
| PcaU-R15-05 | G102S, A165T, S209N |
| PcaU-R15-07 | A165T, K212N |
| PcaU-R15-13 | A165T, Q181K, K212N, Q237R |
| PcaU-R15-18 | A165T, K212N |
| PcaU-R15-22 | A165T, N201H, R206C, Q261H |
| PcaU-R15-25 | A165T, Q192R, I260V |
| PcaU-R15-27 | A165T, K212M |
| PcaU-R15-30 | A165T, L193V, K212E |
| PcaU-R15-33 | A165T, V241I, N276I |
| PcaU-R15-36 | A165T |
| PcaU-R15-37 | T20N, A165T, S122R,R206C, Q237R, V241I |
| PcaU-R15-38 | K9R, A165T, A174T, Q261R,H262Y |
| PcaU-R15-48 | M1I, A165T, Q192R |

### Consensus mutant testing

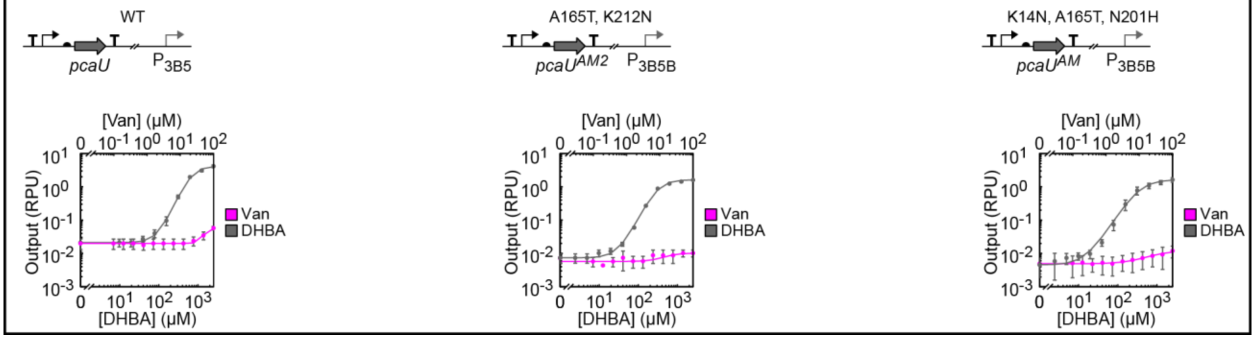

### NahR:$P_{Sal}$:Sal

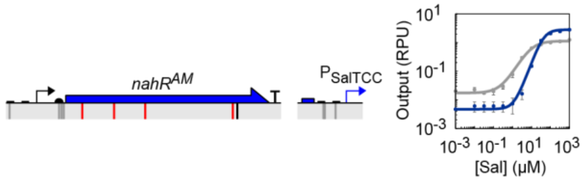

### Directed evolution interventions

| NahR | [Cl-Phe] [Negative ligand] | DNAP | [Positive ligand] | Notes |
|---|---|---|---|---|
| R1 | 2 mM | PK6 | 50 μM Sal | |
| R2 | 2 mM | PK6 | 50 μM Sal | |
| R3 | 2 mM | PK6 | 50 μM Sal | |
| R4 | 2 mM | PK6 | 50 μM Sal | |
| R5 | 4 mM | PK6 | 50 μM Sal | |
| R6 | 4 mM | PK6 | 50 μM Sal | |
| R7 | 4 mM | KOD | 50 μM Sal | |
| R8 | 4 mM | KOD | 50 μM Sal | |
| R9 | 4 mM | KOD | 30 μM Sal | Shuffle |
| R10 | 4 mM | KOD | 20 μM Sal | |
| R11 | 4 mM | KOD | 10 μM Sal | |

### Initial library (1240 bp)

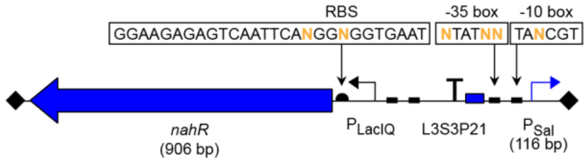

### On/off screen

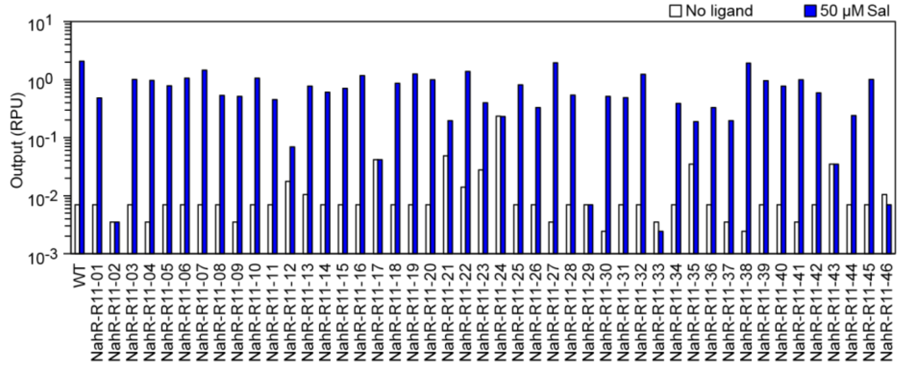

### Sequencing

| Mutant | Mutations |
|---|---|
| NahR-R11-04 | P72H, N144D, A230T, R248C, S260C |
| NahR-R11-07 | P72H, R248C |
| NahR-R11-16 | V118A, Q168K, R248L |
| NahR-R11-19 | P72H, R248C, D251N |
| NahR-R11-27 | L6M, I25V, P72H, V118A, L154H, R183C, R248C |
| NahR-R11-38 | L6M, I25V, P72H, V118A, L154H, I200V, R248C |
| NahR-R11-41 | I25V, Q49R, V118A, V197A, R248C |

### Consensus mutant testing

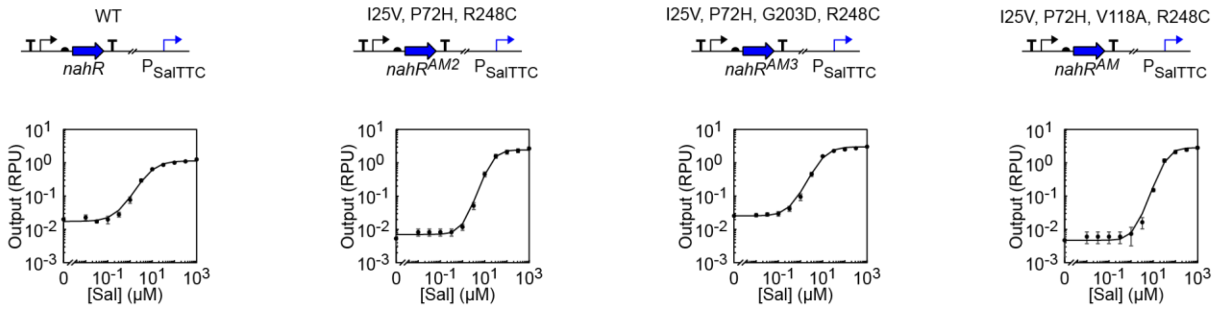

### CinR:$P_{Cin}$:OHC14

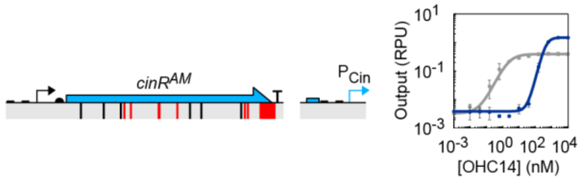

### Directed evolution interventions

| CinR | [Cl-Phe] | [Negative ligand] | DNAP | [Positive ligand] | Notes |
|---|---|---|---|---|---|
| R1 | 2 mM | | PK6 | 5 µM OHC14 | |
| R2 | 2 mM | | PK6 | 5 µM OHC14 | |
| R3 | 2 mM | | PK6 | 5 µM OHC14 | |
| R4 | 2 mM | | PK6 | 1.25 µM OHC14 | |
| R5 | 2 mM | | PK6 | 1.25 µM OHC14 | |
| R6 | 4 mM | | PK6 | 1.25 µM OHC14 | |
| R7 | 4 mM | 10 µM OC12 | PK6 | 1.25 µM OHC14 | Shuffle |
| R8 | 4 mM | 10 µM OC12 | PK6 | 1.25 µM OHC14 | |
| R9 | 4 mM | 10 µM OC12 | KOD | 1.25 µM OHC14 | |

### Initial library (1172 bp)

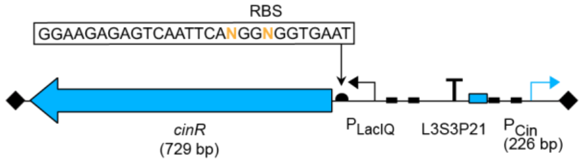

### On/off screen

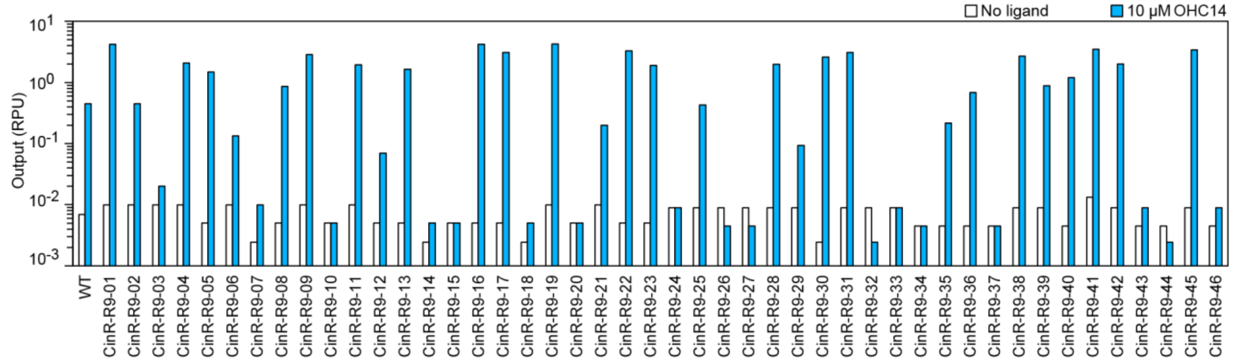

### Sequencing

| Mutant | Mutations |
|---|---|
| CinR-R9-01 | R86H, R151H, P177S, K220R |
| CinR-R9-05 | A103V, A136V, R149C, H170Y, L187M, K220R, *242YRIRMTRRNW* |
| CinR-R9-08 | A19V, R137P, P139S, E142K, A178V, G204D, K220R, *242YRIRMTRRNW* |
| CinR-R9-11 | L70I, Q87L, G115D, T144I, L146F, R149C, Y196N, K220R, *242YRIRMTRRNW* |
| CinR-R9-13 | R28H, K50M, Q87P, V106L, H111Y, P121T, Q127R, D141N, K220R, *242YRIRMTRRNW* |
| CinR-R9-16 | N71S, T98A, A136V, K197N, A230V, *242YRIRMTRRNW* |
| CinR-R9-17 | S72C, V80I, G114D, G115D, R137H, K220R, A224T, *242YRIRMTRRNW* |
| CinR-R9-28 | M1I, A63T, R67C, P97L, I138M, E142K, R149H, G204D, N240D, *242YKDPHDAT* |
| CinR-R9-30 | S72C, V80I, G114D, G115D, R137H, K220R, A224T, *242YRIRMTRRNW* |
| CinR-R9-31 | Y73H, I138M, V166I, *242YRIRMTRRNW* |
| CinR-R9-36 | A32T, Y73H, A103T, G115S, L146F, K220R, *242YRIRMTRRNW* |
| CinR-R9-39 | T98M, E100G, P175L, K220R, *242YRIRMTRRNW* |
| CinR-R9-40 | T5I, G204D, K220R, *242YRIRMTRRNW* |
| CinR-R9-42 | Q16R, A25V, A48T, A63D, K220R, Q235R, R237H |
| CinR-R9-45 | K18M, A19T, K81R, D141E, N152D, K220R, A230T, *242YRIRMTRRNW* |

### Consensus mutant testing

WT

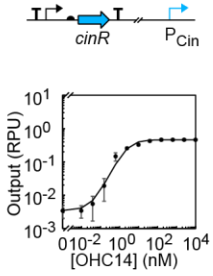

S72C, V80I, G114D, G115D, R137H, C150Y, K220R, A224T, R237C, *242YRIRMTRRNW*

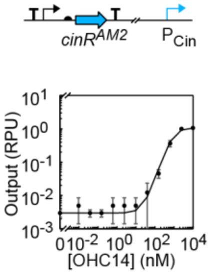

S72C, V80I, G114D, G115D, R137H, K220R, A224T, *242YRIRMTRRNW*

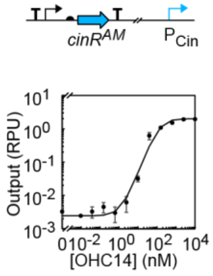

### AcuR:$P_{Acu}$:Acr

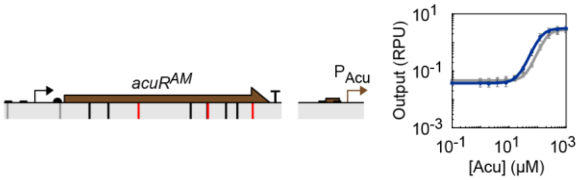

### Directed evolution interventions

| AcuR | [Cl-Phe] [Negative ligand] | DNAP | [Positive ligand] | Notes |
|---|---|---|---|---|
| R1 | 2 mM | PK6 | 100 µM Acr | |
| R2 | 2 mM | PK6 | 100 µM Acr | |
| R3 | 4 mM | PK6 | 100 µM Acr | |
| R4 | 4 mM | PK6 | 100 µM Acr | |
| R5 | 4 mM | PK6 | 100 µM Acr | |
| R6 | 4 mM | PK6 | 100 µM Acr | |
| R7 | 4 mM | KOD | 100 µM Acr | |

### Initial library (957 bp)

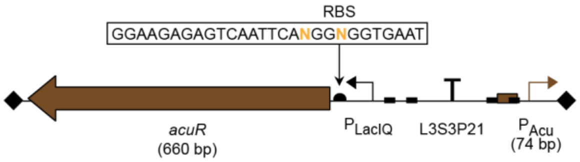

### On/off screen

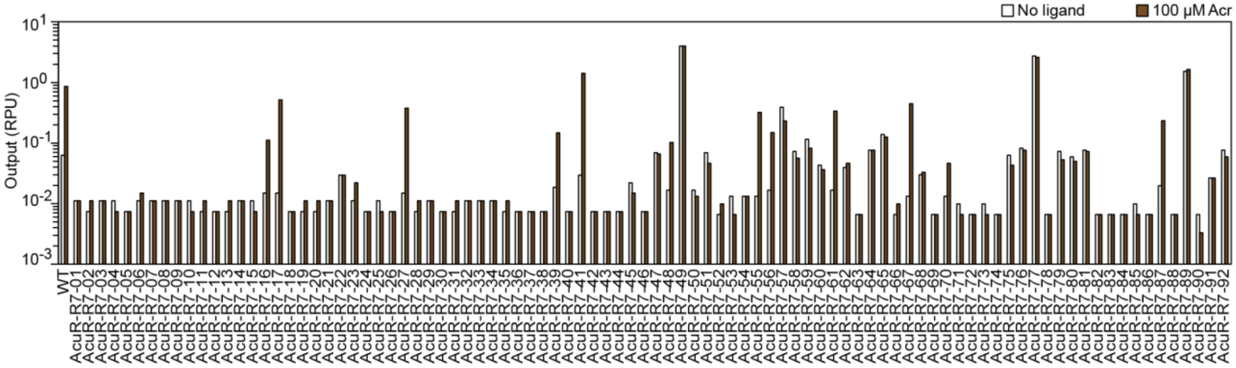

### Sequencing

| Mutant | Mutations |
|---|---|
| AcuR-R7-16 | G147C, R161H, Q218L |
| AcuR-R7-17 | V10D, P138Q, E150K, A171V |
| AcuR-R7-21 | T6I, A117V |
| AcuR-R7-39 | A28V, D73N, R211H |
| AcuR-R7-41 | P138S |
| AcuR-R7-48 | Q30R, A87V |
| AcuR-R7-55 | P138Q, Q218R, E219N, *220NDKDPHDAT* |
| AcuR-R7-56 | R34H, G49S, G167S |
| AcuR-R7-61 | T4I, D24E, A117V |
| AcuR-R7-67 | E80K, R154H, L202M |
| AcuR-R7-87 | D24N, R122S |

### Consensus mutant testing

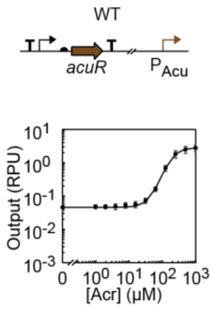

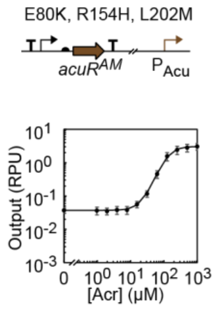

### MphR:$P_{Mph}$:Ery

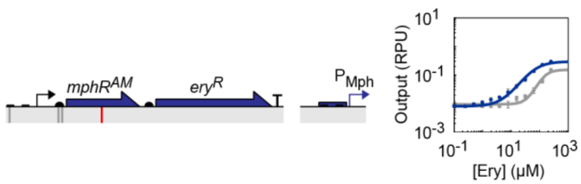

### Directed evolution interventions

| MphR | [Cl-Phe] | [Negative ligand] | DNAP | [Positive ligand] | Notes |
|---|---|---|---|---|---|
| R1 | 2 mM | | PK6 | 100 μM Ery | |
| R2 | 2 mM | | PK6 | 100 μM Ery | |
| R3 | 4 mM | | PK6 | 100 μM Ery | |
| R4 | 4 mM | | PK6 | 100 μM Ery | |
| R5 | 4 mM | | PK6 | 100 μM Ery | |
| R6 | 4 mM | | PK6 | 100 μM Ery | |
| R7 | 4 mM | | KOD | 100 μM Ery | |

### Initial library (1613 bp)

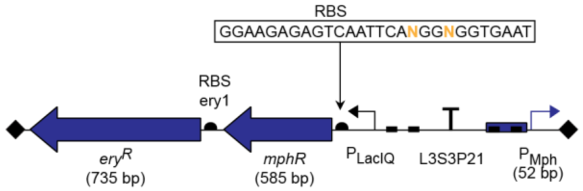

### On/off screen

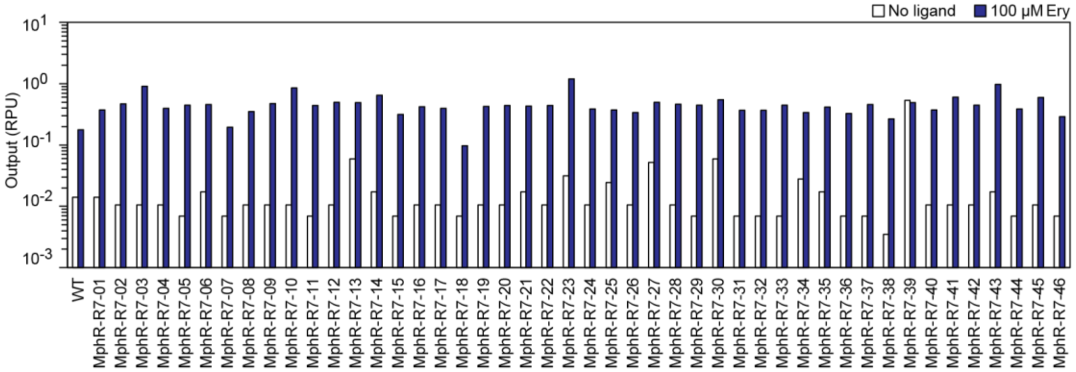

### Sequencing

| Mutant | Mutations |
|---|---|
| MphR-R7-03 | N94D |
| MphR-R7-05 | E11K, N94D, N97D |
| MphR-R7-10 | L30M, N94D, N97D, P182Q, D186N |
| MphR-R7-11 | N94D, R96S, Q188R |
| MphR-R7-29 | P4L, G76S, N94D |
| MphR-R7-31 | N94D, A174V |
| MphR-R7-32 | T53I, N94D, H193R |
| MphR-R7-33 | N94D, A142V |
| MphR-R7-37 | K7I, N94D, D160N |
| MphR-R7-38 | E11D, N94D, Q188R |
| MphR-R7-41 | N94D, A192V |
| MphR-R7-43 | N50D, V88I, A171T, E183G, Q191R, H193N |
| MphR-R7-44 | N50D, P136L |
| MphR-R7-45 | N94D |

### Consensus mutant testing

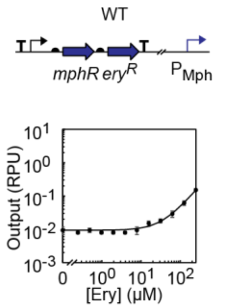

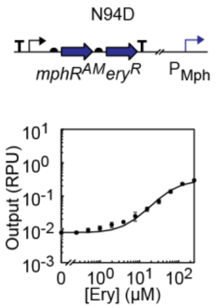

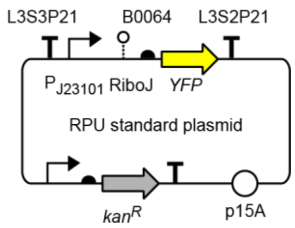

**Supplementary Figure 1: RPU Standard plasmid**

The RPU standard plasmid is used to normalize fluorescence output[6] (Methods). Promoter ($P_{J23101}$) sequence is provided in Supplementary Table 1. Insulator (RiboJ) sequence is provided in Supplementary Table 2. RBS (B0064) sequence is provided in Supplementary Table 3. Protein (YFP) sequence is provided in Supplementary Table 4. Terminator (L3S3P21, L3S2P21) sequences are provided in Supplementary Table 5. Full plasmid sequence is provided in Supplementary Table 8.

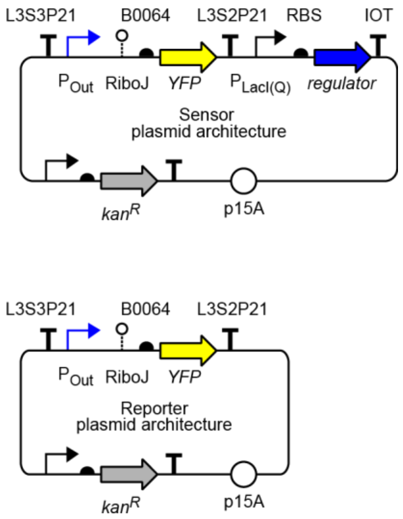

**Supplementary Figure 2: General plasmid architectures**

All sensors are assembled into the same sensor plasmid architecture for characterization. All output promoters are assembled into the same reporter plasmid architecture for use in Marionette strains. Promoter ($P_{LacI}$, $P_{LacIQ}$) sequences are provided in Supplementary Table 1. Insulator (RiboJ) sequence is provided in Supplementary Table 2. RBS (B0064) sequence is provided in Supplementary Table 3. Protein (YFP) sequence is provided in Supplementary Table 4. Terminator (L3S3P21, L3S2P21, IOT) sequences are provided in Supplementary Table 5.

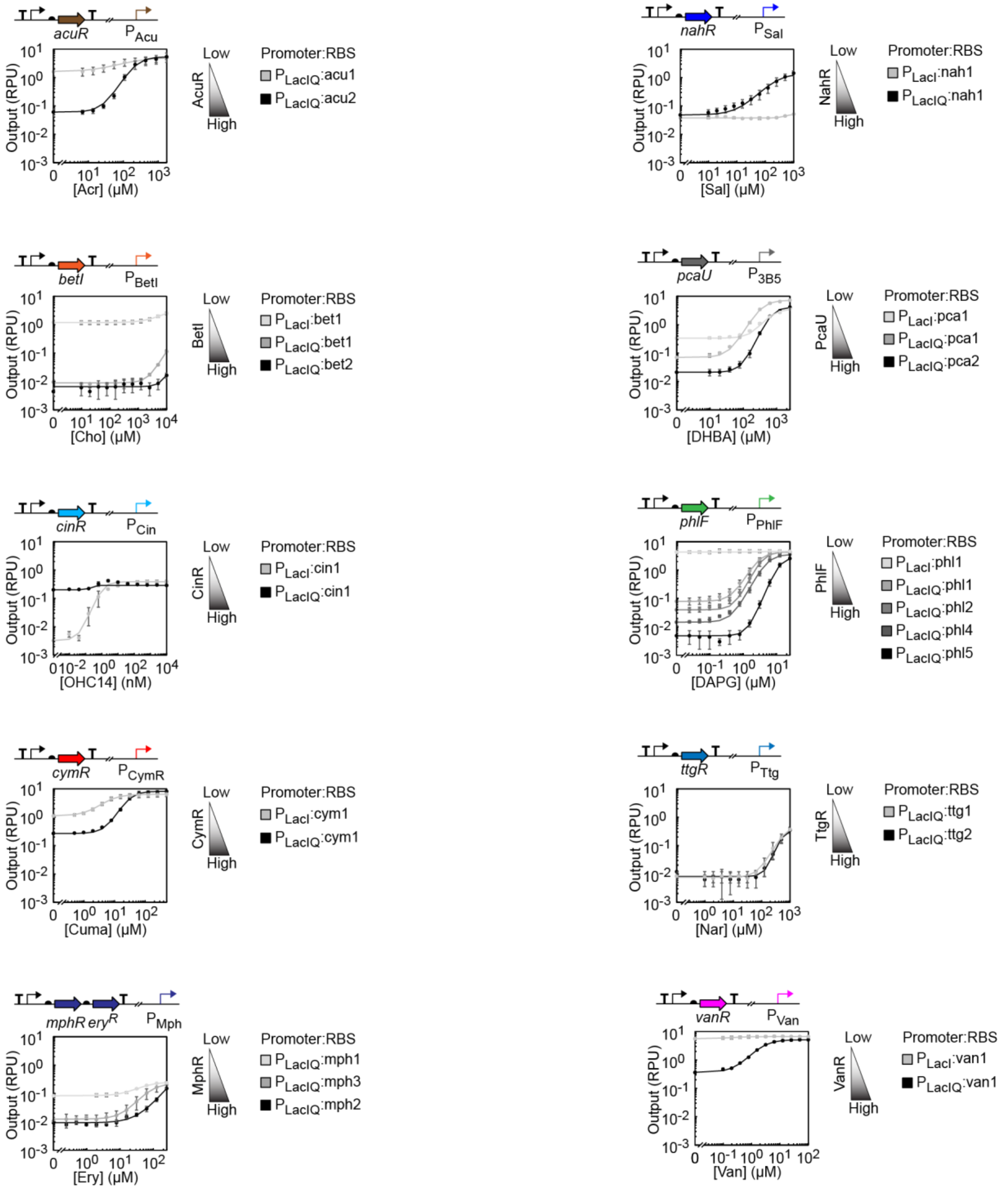

**Supplementary Figure 2: Initial sensor tuning**

The promoter and/or RBS driving regulator expression was tuned, and the response function for each construct is shown. Darker color denotes higher expression of the regulator, as determined by the relative promoter and RBS strengths. All data are from *E. coli* DH10B cells in mid-log growth in LB media. The mean of three replicates performed on different days is shown, and error bars represent the standard deviation. Promoter and RBSs are noted, and their sequences are provided in Supplementary Tables 1 and 3.

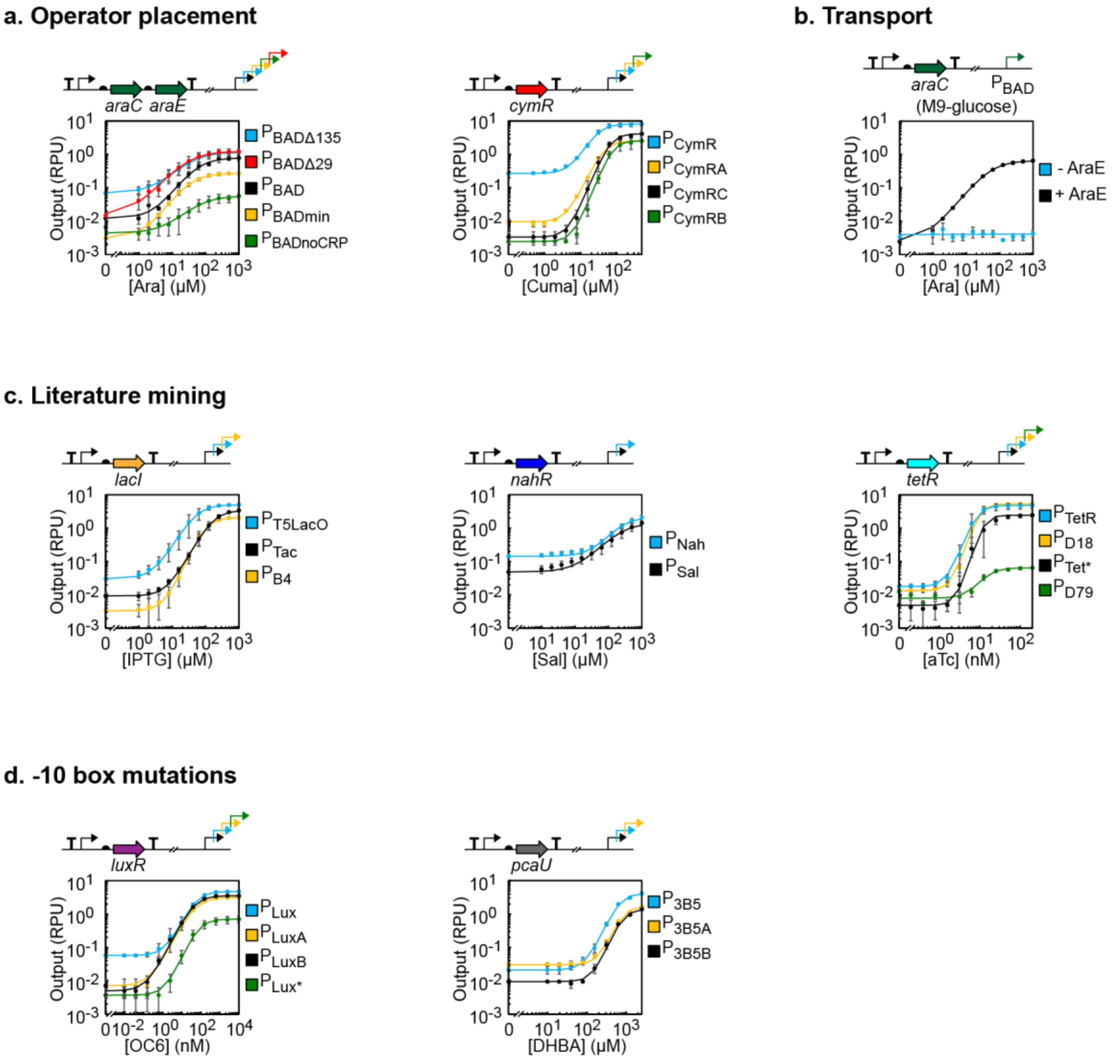

**Supplementary Figure 4: Initial sensor engineering**

Output promoter mutations were made based on either rational design or inspired by the literature. Multiple promoters in the diagram denote that multiple promoter variants were tested. All data are from *E. coli* DH10B cells in mid-log growth in LB media (except M9-glucose was used in section b. The mean of three replicates performed on different days is shown, and error bars represent the standard deviation. Promoter variants are noted, and their sequences are provided in Supplementary Table 1.

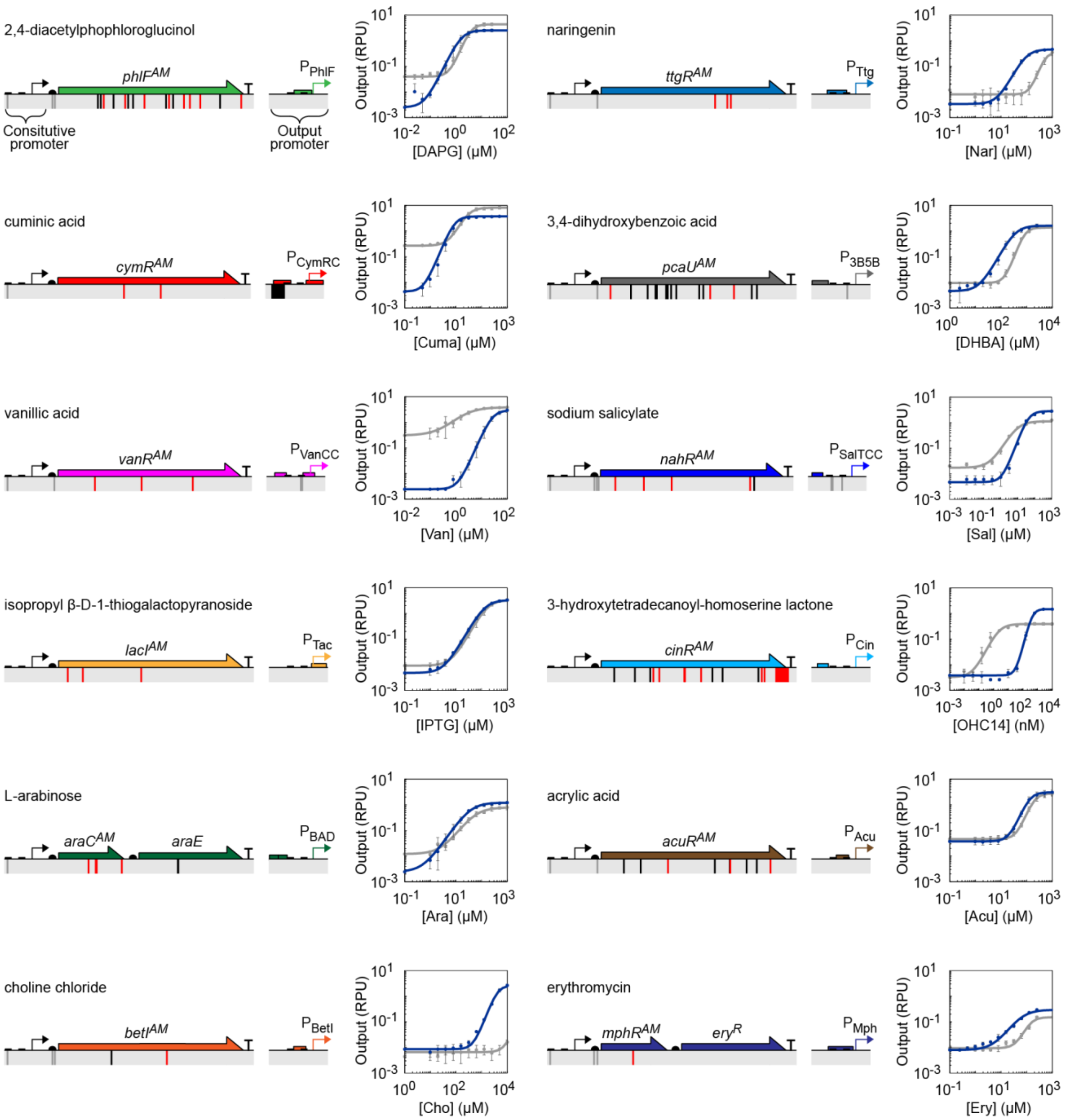

**Supplementary Figure 5: Effect of sensor evolution on response functions**

Each genetically encoded sensor is shown, with coding (red) and non-coding (grey/black) mutations noted. Mutations in grey were also applied to the parental sensor. The corresponding response functions comparing the evolved (blue) and parental (grey) sensors. All data are from *E. coli* DH10B cells in mid-log growth in LB media. The mean of three replicates performed on different days is shown, and error bars represent the standard deviation. Response functions were obtained by fitting the experimental data to Equation 1 and are shown in Figure 2a. Response function parameters for evolved sensors are provided in Table 1. Sequences of promoters and regulators are provided in Supplementary Table 1 and 4.

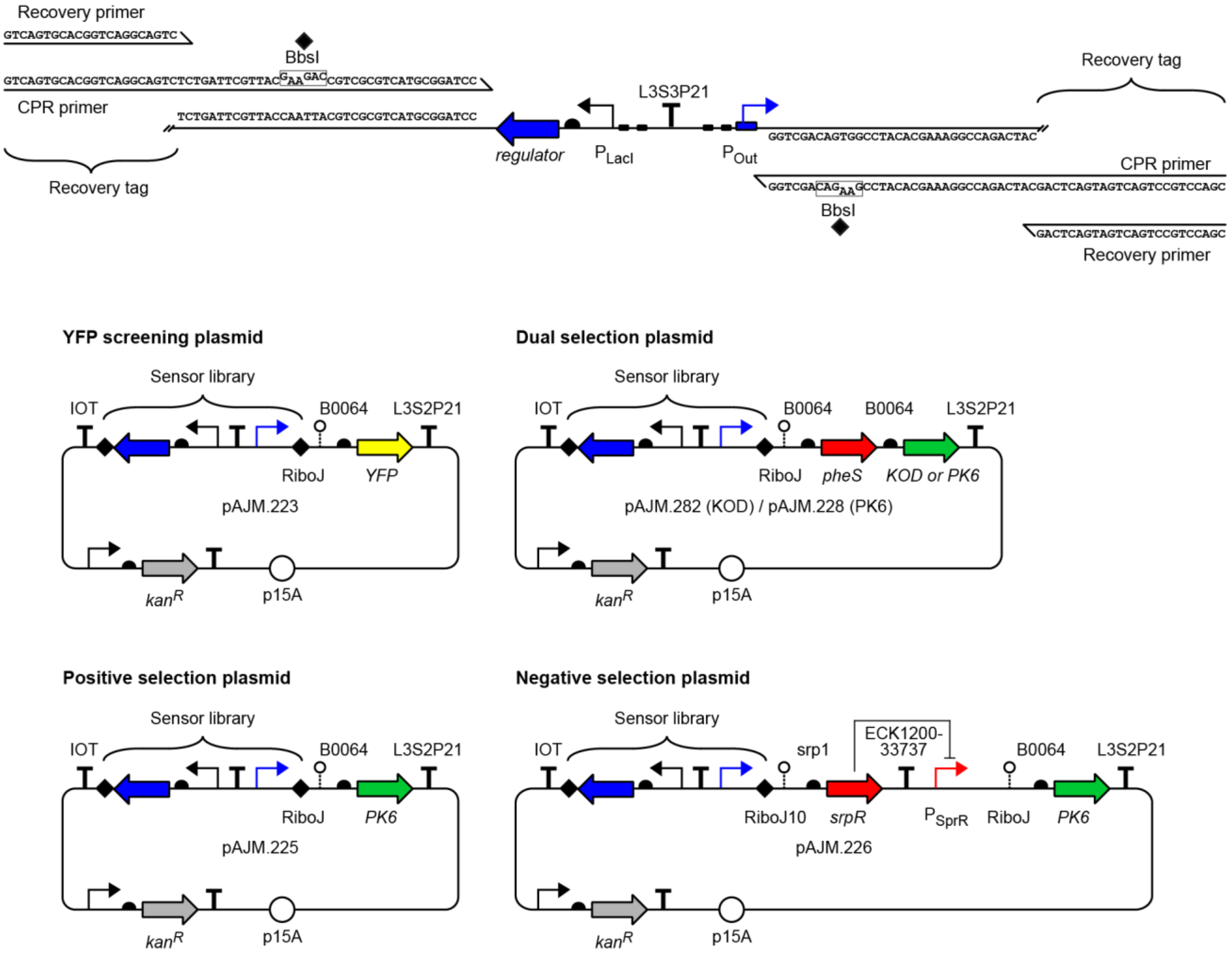

**Supplementary Figure 6: Directed evolution selection plasmids**

Sensor libraries were constructed as divergent operons flanked by BbsI sites (black diamonds). Most libraries were Golden Gate assembled into a dual selection plasmids during selection and into the YFP screening plasmid for on/off screening. The first two rounds of the LacI selection utilized the positive and negative selection plasmids, respectively. CPR primers are used in the emulsion PCR; they re-introduce BbsI sites and append recovery tags to the library. Recovery primers are used in the recovery PCR; they anneal to the recovery tags. Promoter ($P_{SrpR}$) sequence is provided in Supplementary Table 1. Insulator (RiboJ, RiboJ10) sequences are provided in Supplementary Table 2. RBS (B0064, srp1) sequences are provided in Supplementary Table 3. Protein (YFP, PheS, PK6, KOD, SrpR) sequences are provided in Supplementary Table 4. Terminator (L3S3P21, L3S2P21, IOT, ECK120033737) sequences are provided in Supplementary Table 5.

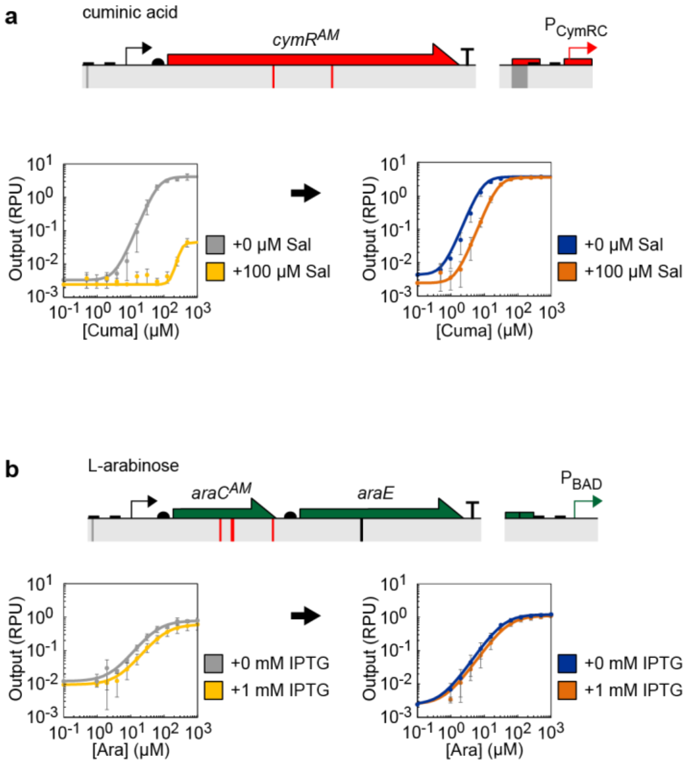

**Supplementary Figure 7: Chemical antagonism**

**a)** Response function with Sal (light/dark orange) and without Sal (grey/blue) for reference CymR (left) and evolved CymR$^{AM}$ (right). **b)** Response function with IPTG (light/dark orange) and without (grey/blue) IPTG for reference AraC (left) and evolved AraC$^{AM}$ (right). All data are from *E. coli* DH10B cells in mid-log growth in LB media. The mean of three replicates performed on different days is shown, and error bars represent the standard deviation. Response functions were obtained by fitting the experimental data to Equation 1 and are shown in Figure 2c. Response function parameters for evolved sensors are provided in Table 1. Sequences of promoters and regulators are provided in Supplementary Table 1 and 4.

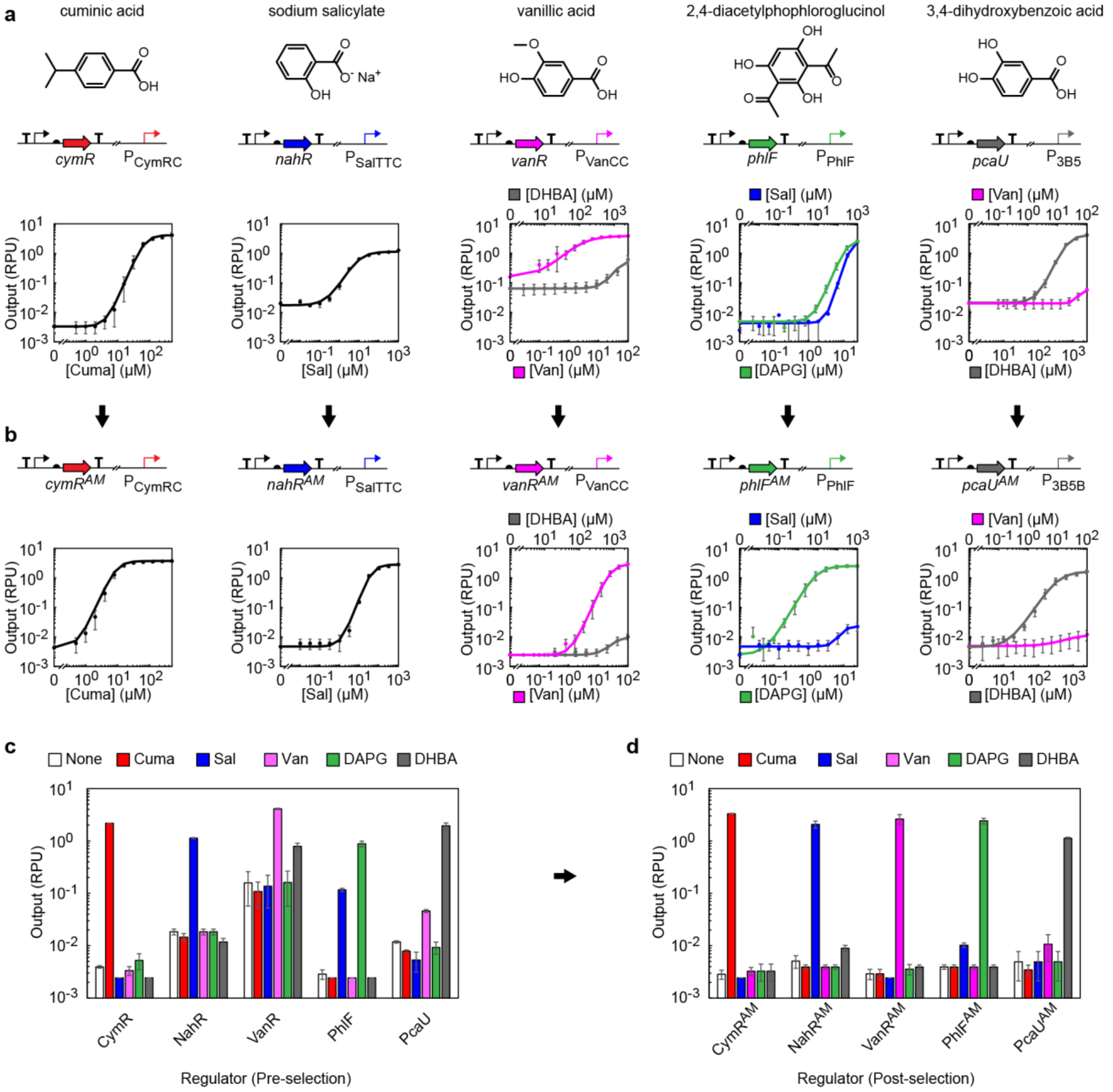

**Supplementary Figure 8: Chemical cross-reactivity**

CymR, NahR, VanR, PhlF, and PcaU respond to similar chemical inducers. **a)** Response functions for the wild-type sensors with cognate inducers and cross-reactive inducers (where appropriate). **b)** Response functions for the evolved sensors with cognate inducers and cross-reactive inducers (where appropriate). Response functions were obtained by fitting the experimental data to Equation 1. Response function parameters for evolved sensors are provided in Table 1. **c)** Cross-reactivity assay of wild-type sensors. **d)** Cross-reactivity assay of evolved sensors. Cross-reactivity heat maps are provided in Figure 2b. Inducer concentrations are 100 µM Cuma, 100 µM Sal, 100 µM Van 10 µM DAPG, 1 mM DHBA. All data are from *E. coli* DH10B cells in mid-log growth in LB media. The mean of three replicates performed on different days is shown, and error bars represent the standard deviation. Sequences of promoters and regulators are provided in Supplementary Table 1 and 4.

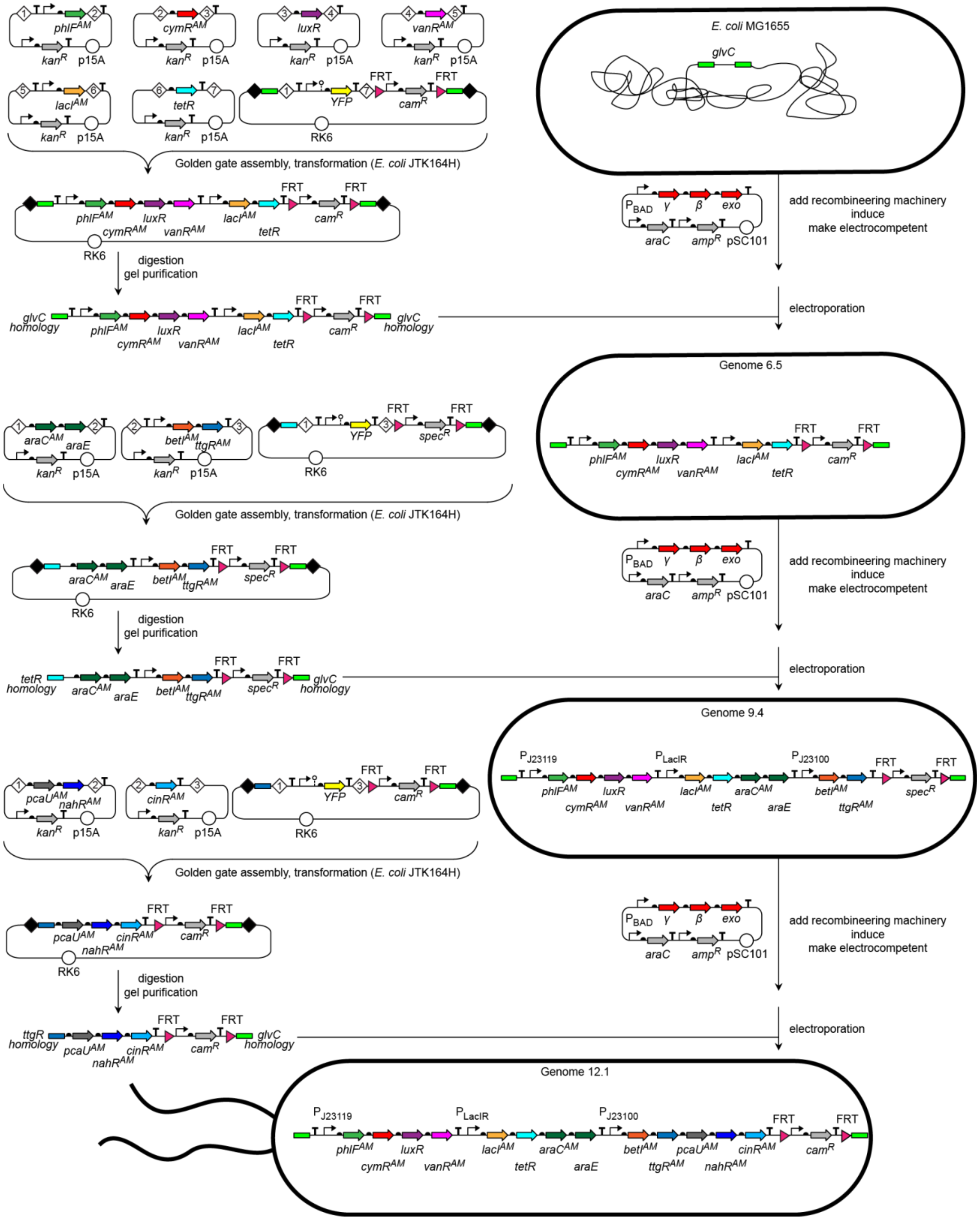

**Supplementary Figure 9: Genomic integration of the Marionette cluster**

Regulators were assembled into RK6 suicide vectors. Regulators were serially integrated into *glvC* locus of the *E. coli* MG1655 genome using λ Red recombineering (Methods). All part sequences are provided in Supplementary Tables 1-5. The sequence of the full cluster is provided in Supplementary Table 6.

| Construct | RBS | Score | Note | RBS | Score | Note | RBS | Score | Note | RBS | Score | Note | RBS | Score | Note | RBS | Score | Note |
|---|---|---|---|---|---|---|---|---|---|---|---|---|---|---|---|---|---|---|
| Plasmid | phl2 | 6066 | | cym1 | 5300 | | lux1 | 23403 | | van2 | 10916 | | lac1 | 3379 | | tet1 | 25607 | |
| Genome 6.1 | phl6 | 620 | low | cym3 | 391 | low | lux2 | 4056 | good | van4 | 13069 | low | lac3 | 328 | low | tet3 | 62294 | high |
| Genome 6.2 | phl3 | 1999 | good | cym2 | 4631 | good | lux2 | 4056 | good | van5 | 26850 | low | lac4 | 973 | low | tet4 | 52502 | high |
| Genome 6.3 | phl3 | 1999 | good | cym2 | 4631 | good | lux2 | 4056 | good | van3 | 42112 | good | lac5 | 3868 | low | tet2 | 17908 | good |
| Genome 6.4 | phl3 | 1999 | good | cym2 | 4631 | good | lux2 | 4056 | good | van3 | 42112 | good | lac5 | 3868 | low | tet2 | 17908 | good |
| Genome 6.5 | phl3 | 1999 | good | cym2 | 4631 | good | lux2 | 4056 | good | van3 | 42112 | good | lac2 | 10889 | good | tet2 | 17908 | good |

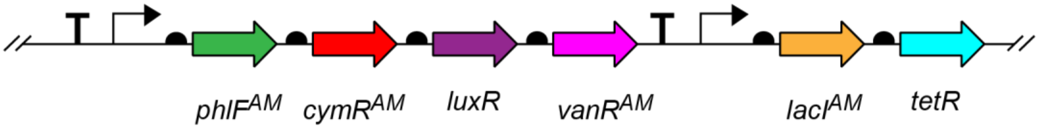

| Construct | RBS | Score | Note | RBS | Score | Note | RBS | Score | Note | RBS | Score | Note |
|---|---|---|---|---|---|---|---|---|---|---|---|---|
| Plasmid | ara1 | 2953 | | e1 | 930 | | bet2 | 1147 | | ttg2 | 17865 | |
| Genome 9.1 | ara2 | 6374 | good | e1 | 930 | good | bet2 | 1147 | low | ttg4 | 19594 | high |
| Genome 9.2 | ara2 | 6374 | good | e1 | 930 | good | bet2 | 1147 | low | ttg3 | 17908 | good |
| Genome 9.3 | ara2 | 6374 | good | e1 | 930 | good | bet4 | 1720 | low | ttg3 | 17908 | good |
| Genome 9.4 | ara2 | 6374 | good | e1 | 930 | good | bet3 | 8155 | good | ttg3 | 17908 | good |

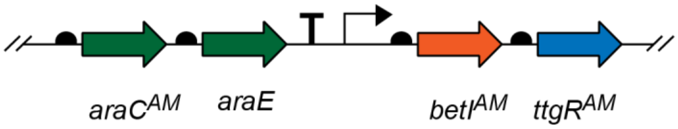

| Construct | RBS | Score | Note | RBS | Score | Note | RBS | Score | Note |
|---|---|---|---|---|---|---|---|---|---|
| Plasmid | pca2 | 17865 | | nah2 | 2450 | | cin1 | 22373 | |
| Genome 11.1 | pca4 | 43031 | low | nah3 | 3135 | good | | | |
| Genome 11.2 | pca3 | 105849 | good | nah3 | 3135 | good | | | |
| Genome 12.1 | pca3 | 105849 | good | nah3 | 3135 | good | cin2 | 2076 | good |

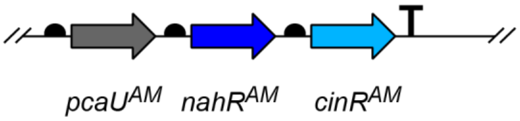

**Supplementary Figure 10: Tuning the RBSs in the Marionette cluster**

The Marionette cluster was integrated in three sections (Supplementary Figure 9). The RBS strength of each regulator was tuned by iteratively designing, building and testing multiple versions of each section. The RBS name, score (Translation Initiation Rate from the RBS Calculator; in arbitrary units), and a qualitative assessment of the RBS strength are provided for each regulator in each construct. See Methods for full description; all part sequences are provided in Supplementary Tables 1-5.

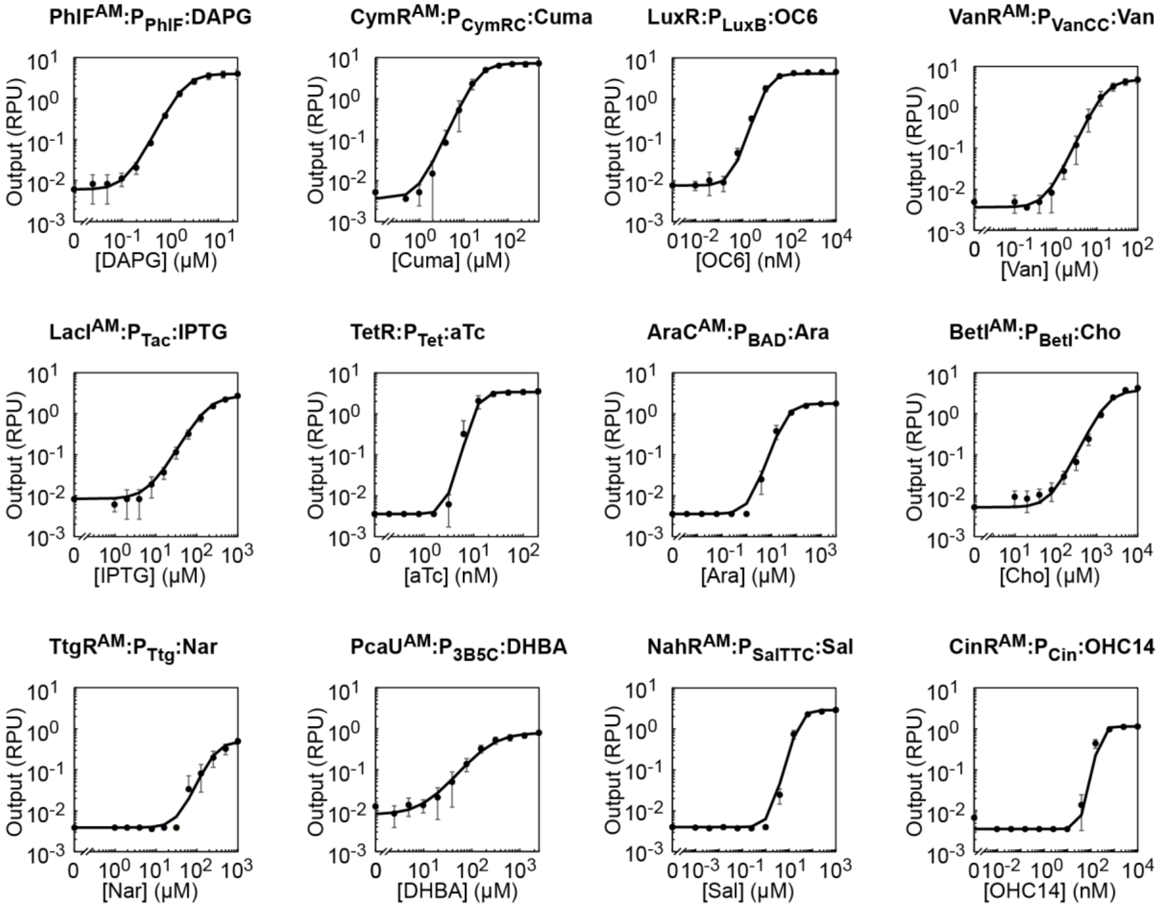

**Supplementary Figure 11: Marionette-Wild response functions**

Response functions for each of the 12 output promoters with their cognate inducers are shown. All data are from cells in mid-log growth in LB media. The mean of three replicates performed on different days is shown, and error bars represent the standard deviation. Response functions were obtained by fitting the experimental data to Equation 1 and are shown in Figure 3b. Response function parameters for genomic sensors are provided in Table 1. Sequences of promoters are provided in Supplementary Table 1. Sequences of reporter plasmids are provided in Supplementary Table 8.

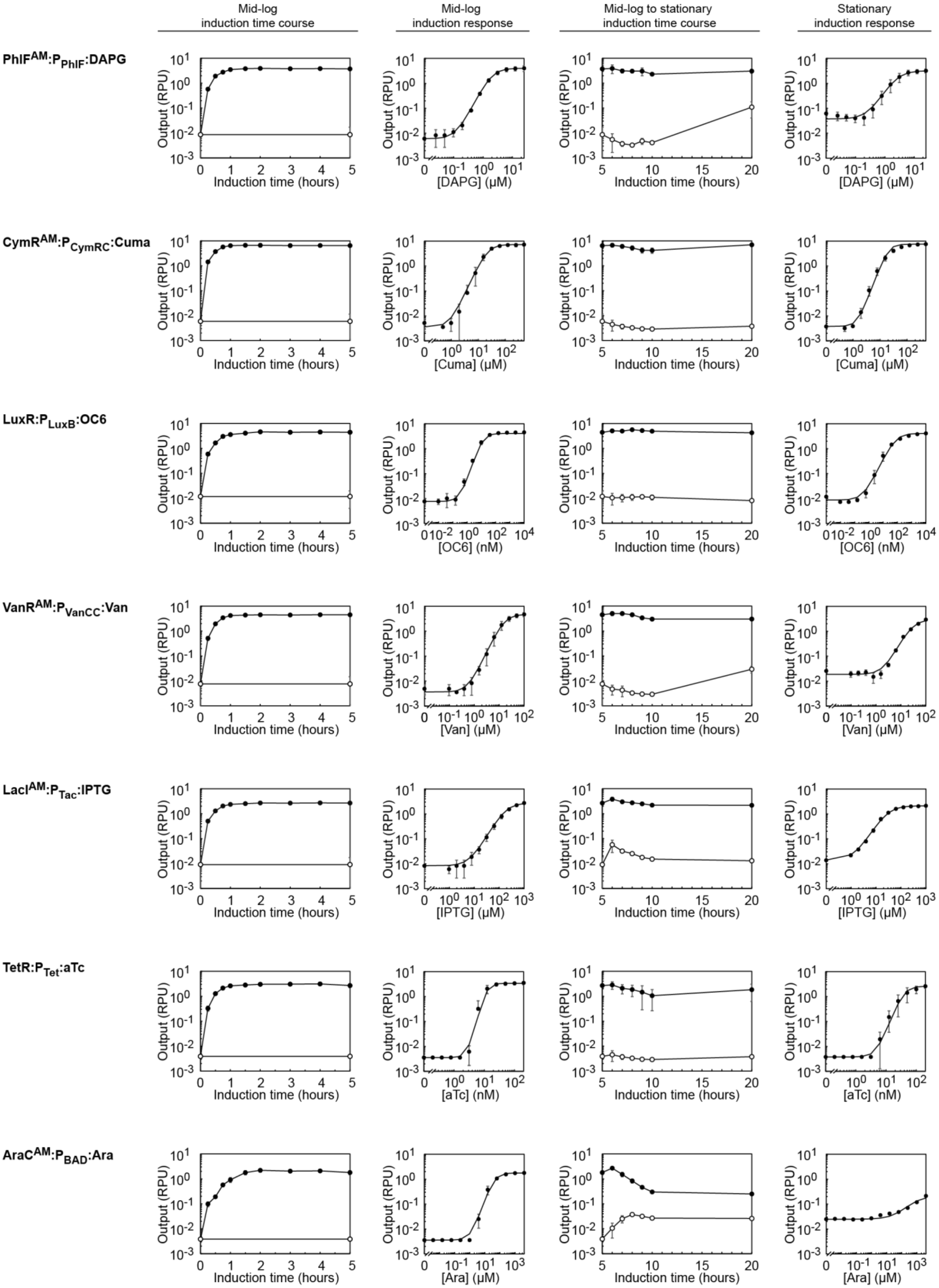
Mid-log induction time course
Mid-log induction response
Mid-log to stationary induction time course
Stationary induction response
$PhlF^{AM}$:$P_{PhlF}$:DAPG
$CymR^{AM}$:$P_{CymRC}$:Cuma
LuxR:$P_{LuxB}$:OC6
$VanR^{AM}$:$P_{VanCC}$:Van
$LacI^{AM}$:$P_{Tac}$:IPTG
TetR:$P_{Tet}$:aTc
$AraC^{AM}$:$P_{BAD}$:Ara
Output (RPU)
Induction time (hours)
[DAPG] (μM)
[Cuma] (μM)
[OC6] (nM)
[Van] (μM)
[IPTG] (μM)
[aTc] (nM)
[Ara] (μM)

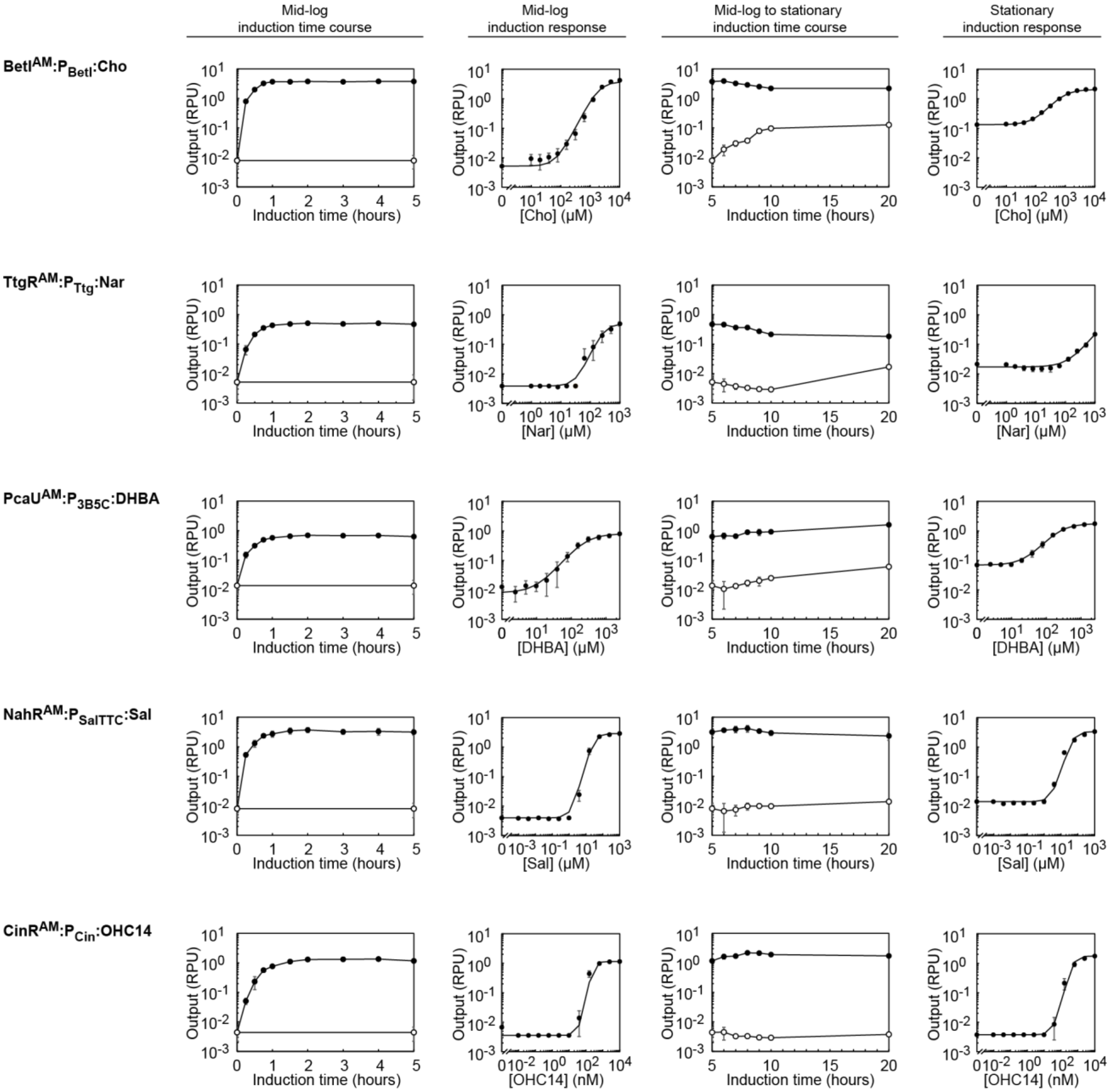

**Supplementary Figure 12: Marionette-Wild induction time course**

**First column)** Time course data from mid-log phase are shown. Inducer was added at different times (black) or not at all (white). **Second column)** Inducer response function after five hours of induction. **Third column)** Time course data from mid-log to stationary phase are shown. Inducer was added at different times (black) or not at all (white). **Forth column)** Inducer response function after 20 hours of induction. See Methods for growth and induction details. The mean of three replicates performed on different days is shown, and error bars represent the standard deviation. Inducer concentrations were: 25 µM DAPG, 100 µM Cuma, 10 µM OC6, 100 µM Van, 1 mM IPTG, 200 nM aTc, 4 mM Ara, 10 mM Cho, 1 mM Nar, 1 mM DHBA, 100 µM Sal, and 10 µM OHC14. LuxR, AraC, NahR, and CinR are activators, while the other regulators are repressors.

Marionette-Wild mid-log

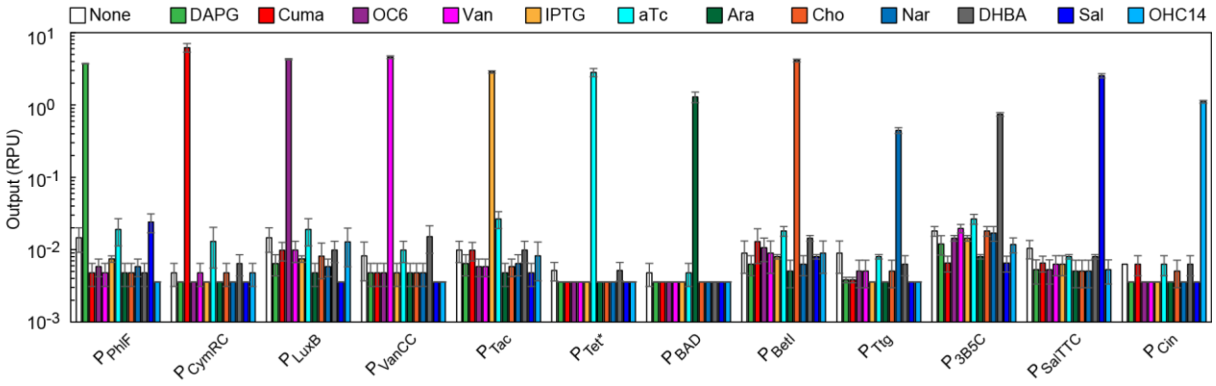

Marionette-Wild stationary

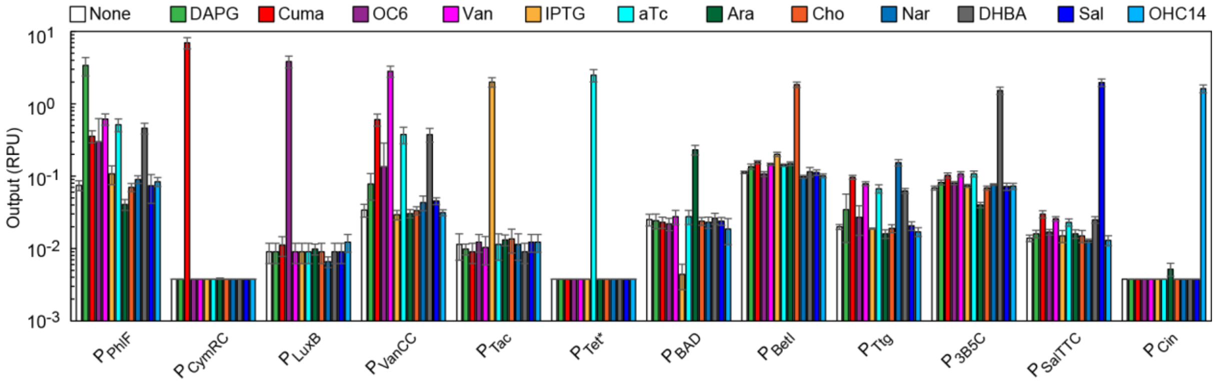

**Supplementary Figure 13: Inducer cross reactivity in Marionette-Wild in mid-log and stationary phase**
Cross-reactivity assay in mid-log and stationary phase (Methods). A cross-reactivity heat map of the mid-log data is provided in Figure 3c. The mean of three replicates performed on different days is shown, and error bars represent the standard deviation. Inducer concentrations were: 25 µM DAPG, 500 µM Cuma, 10 µM OC6, 100 µM Van, 1 mM IPTG, 200 nM aTc, 4 mM Ara, 10 mM Cho, 1 mM Nar, 2.5 mM DHBA, 250 µM Sal, and 10 µM OHC14.

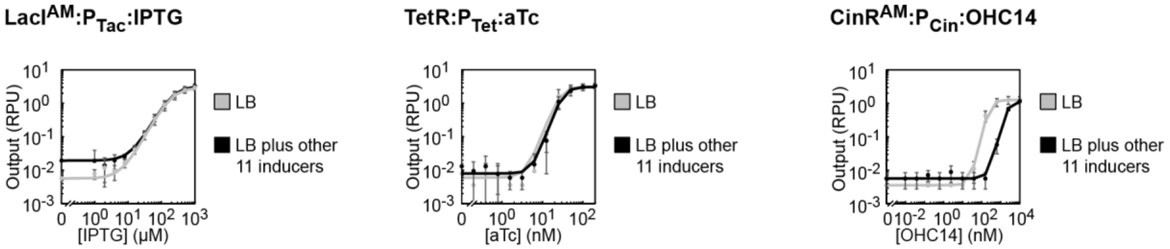

**Supplementary Figure 14: Impact of simultaneous presence of all inducers on response functions**

Marionette-Wild was transformed with each of three reporter plasmids ($P_{Tac}$-YFP, $P_{Tet*}$-YFP, $P_{Cin}$-YFP). Response functions for each output reporter with the appropriate inducer were determined in LB (grey) or in LB containing the other 11 inducers (black). Measurements were taken of cells from mid-log phase. The mean of three replicates performed on different days is shown, and error bars represent the standard deviation. Inducer concentrations were: 6.25 µM DAPG, 100 µM Cuma, 1 µM OC6, 100 µM Van, 1 mM IPTG, 50 nM aTc, 1 mM Ara, 5 mM Cho, 1 mM Nar, 625 µM DHBA, 100 µM Sal, and 1 µM OHC14.

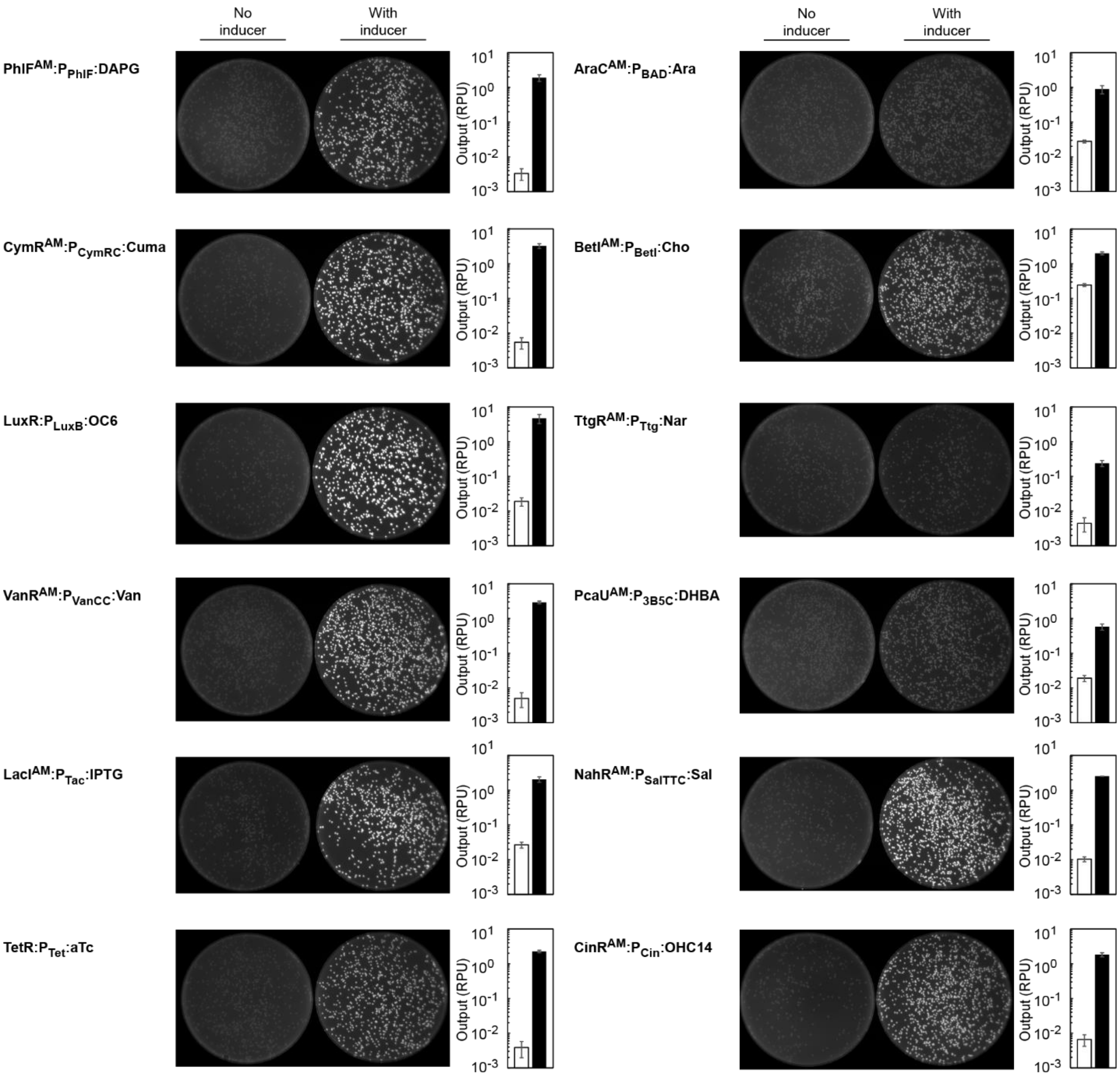

**Supplementary Figure 15: Induction of sensors on plates**

Marionette-Wild was transformed with each of the 12 reporter plasmids and plated on LB-agar with and without the appropriate inducer. After overnight growth at 37 ˚C, plates were incubated at 4 ˚C for one hour and imaged (Methods). Individual colonies from plates with (black) and without (white) inducer were picked, resuspended in PBS, and assayed by cytometry. The mean of three replicates performed on different days is shown, and error bars represent the standard deviation. Inducer concentrations were: 25 µM DAPG, 100 µM Cuma, 10 µM OC6, 100 µM Van, 1 mM IPTG, 200 nM aTc, 4 mM Ara, 10 mM Cho, 1 mM Nar, 1 mM DHBA, 100 µM Sal, and 10 µM OHC14.

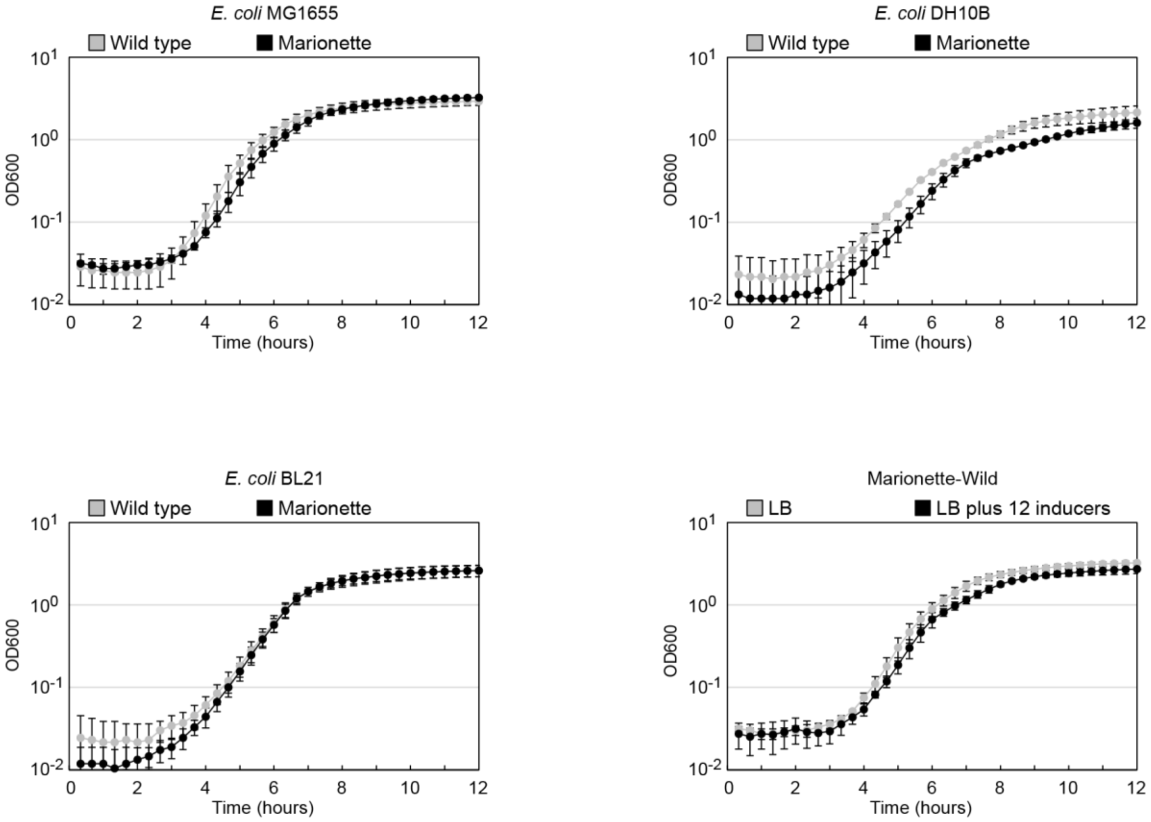

**Supplementary Figure 16: Growth impact of maintaining genomically encoded Marionette cluster**

Marionette-Wild, -Clo, and -Pro (black) as well as their parental strains (*E. coli* MG1655, DH10B, and BL21 respectively; grey) were grown in LB. Marionette-Wild was also grown in LB plus all 12 inducers. OD600 measurements were taken every 20 minutes (Methods). The Marionette-Wild strain grows with a doubling time of 29.0 $\pm$ 1.9 min as compared 27.1 $\pm$ 1.2 for wild-type *E. coli* MG1655. The Marionette-Clo strain grows with a doubling time of 42.3 $\pm$ 3.8 min as compared 45.1 $\pm$ 3.8 for wild-type *E. coli* DH10B. The Marionette-Pro strain grows with a doubling time of 31.1 $\pm$ 1.1 min as compared 34.5 $\pm$ 1.6 for wild-type *E. coli* Bl21. The Marionette-Wild strain grows with a doubling time of 29.0 $\pm$ 1.9 min in LB as compared 31.4 $\pm$ 2.3 for LB plus all 12 inducers. The mean of three replicates performed on different days is shown, and error bars represent the standard deviation. Inducer concentrations were: 25 µM DAPG, 100 µM Cuma, 10 µM OC6, 100 µM Van, 1 mM IPTG, 200 nM aTc, 4 mM Ara, 10 mM Cho, 1 mM Nar, 1 mM DHBA, 100 µM Sal, and 10 µM OHC14.

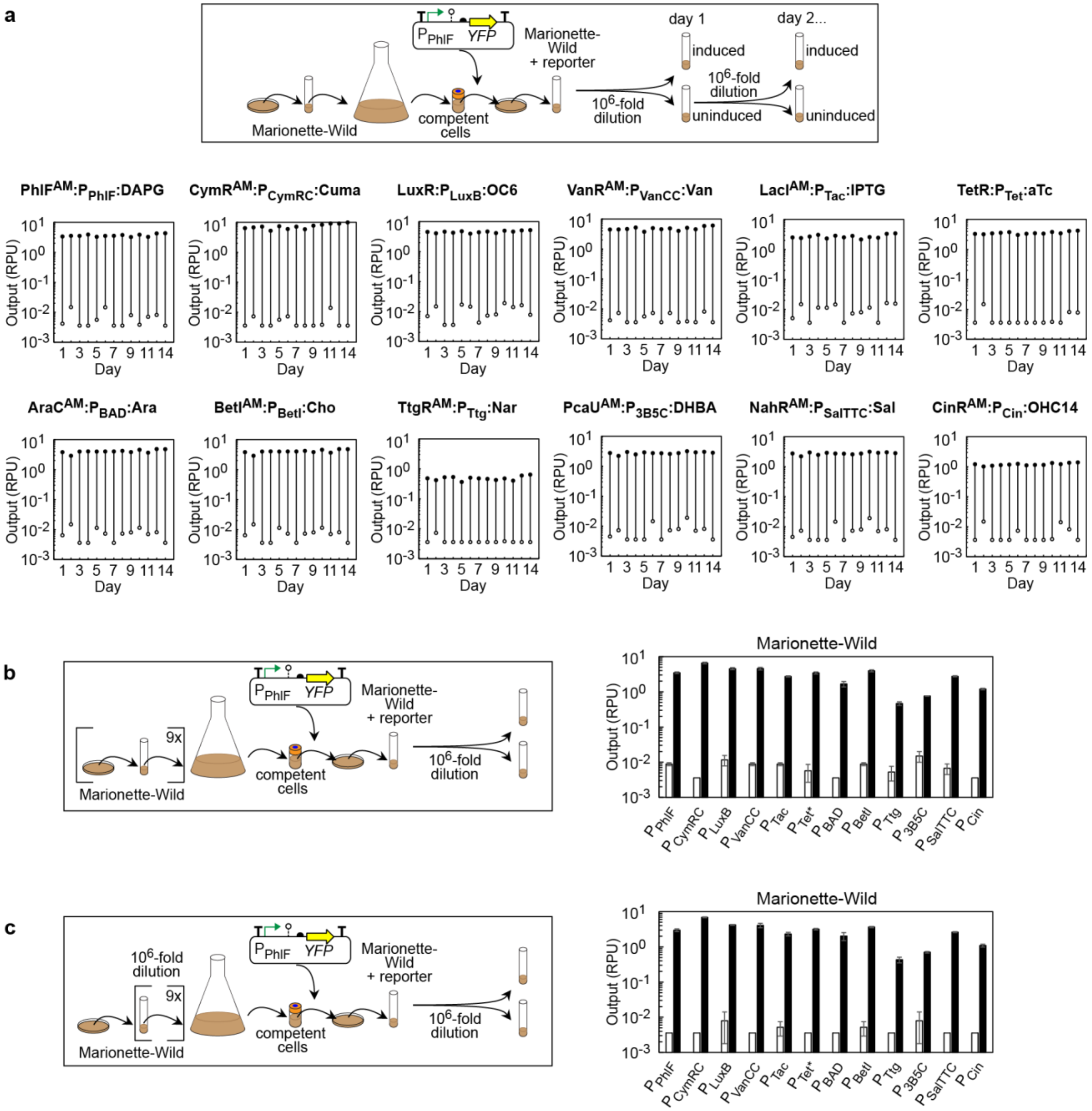

**Supplementary Figure 17: Evolutionary stability of Marionette-Wild**

**a)** Marionette-Wild was transformed with each of the 12 reporter plasmids and grown for 14 days. Each day, cells were assayed with (black) and without (white) the corresponding inducer (Methods). These data are the result of a single experiment. Data from days 1, 7, 11, and 14 are provided in Figure 3d. **b)** Marionette-Wild was passaged between plates and liquid culture nine times and was then transformed with each of the 12 reporter plasmids and assayed with (black) and without (white) the corresponding inducer (Methods). **c)** Marionette-Wild was diluted $10^6$-fold in liquid culture nine times and was then transformed with each of the 12 reporter plasmids and assayed with (black) and without (white) the corresponding inducer (Methods). The mean of three replicates performed on different days is shown, and error bars represent the standard deviation. All data are from cells in mid-log growth in LB media. Inducer concentrations were: 25 µM DAPG, 500 µM Cuma, 10 µM OC6, 100 µM Van, 1 mM IPTG, 200 nM aTc, 4 mM Ara, 10 mM Cho, 1 mM Nar, 2.5 mM DHBA, 250 µM Sal, and 10 µM OHC14.

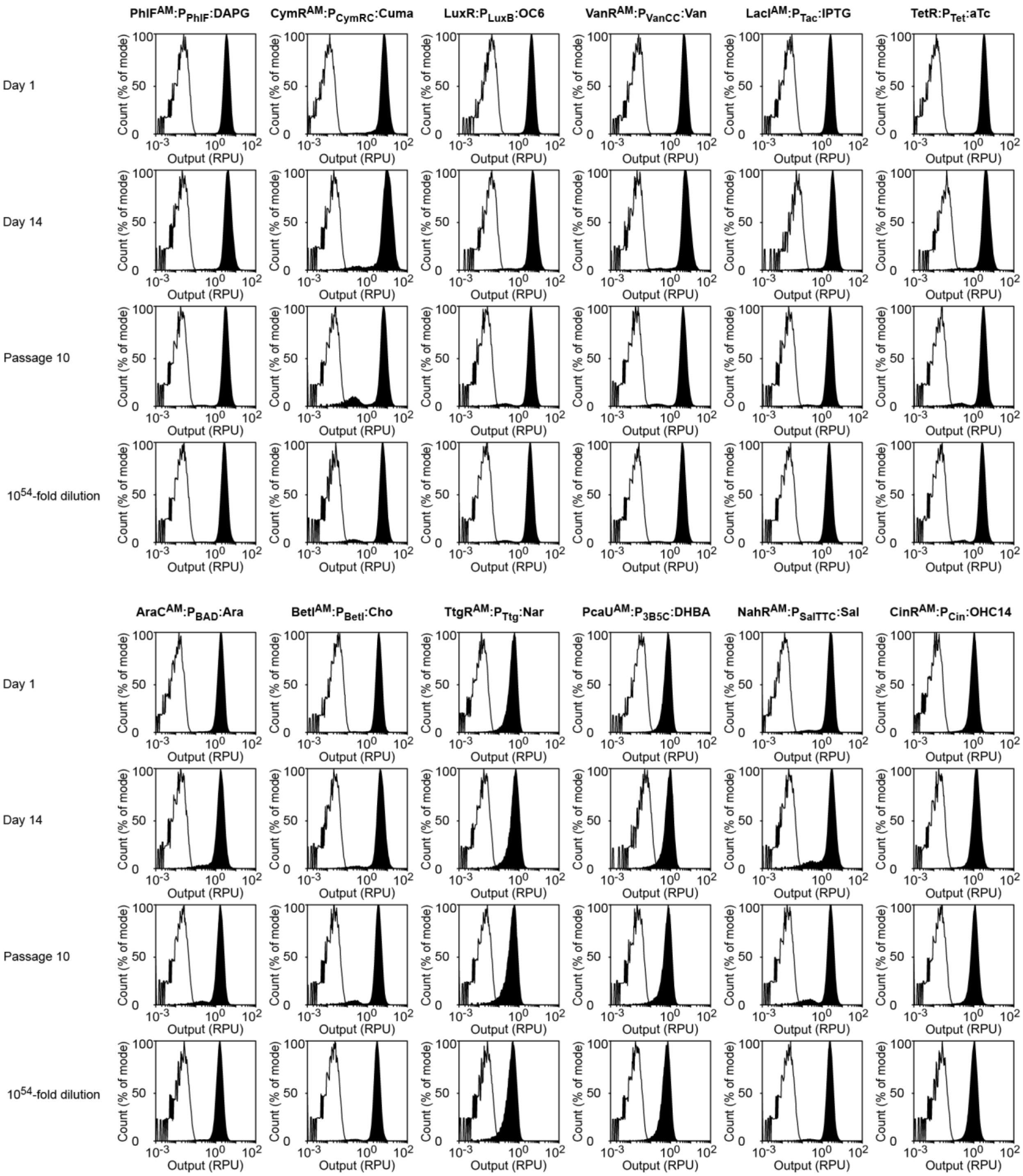

**Supplementary Figure 18: Fluorescence distributions of Marionette-Wild after passaging**

Cytometry histograms with (black) and without (white) the corresponding inducer are shown. Raw values are white cell corrected and RPU normalized. Data are from Day 1, Day 14, Passage 10, and $10^{54}$-fold dilution populations of the evolutionary stability assays (Methods, Supplementary Figure 17). Inducer concentrations were: 25 µM DAPG, 500 µM Cuma, 10 µM OC6, 100 µM Van, 1 mM IPTG, 200 nM aTc, 4 mM Ara, 10 mM Cho, 1 mM Nar, 2.5 mM DHBA, 250 µM Sal, and 10 µM OHC14.

### Supplementary Table 1: Promoters

| Regulator | Promoter | DNA sequence[a] | Source[b] |
|---|---|---|---|
| AcuR | $P_{Acu}$ | CGCTAGCAAGTAAGGCCGACGCTTCACAACCGCACTTGATT**TAATAGACCATACCGTCTATTA**TTTCTGGCCAT | 7-8 |
| AraCE | $P_{BAD}$ | A**GAAACCAATTGTCCATA**TTGCATCAGACATTGCCGTCACTGCGTCTTTTACTGGCTCTTCTCGCTAACCAAACCGGTAACCCCGCTTATTAAAAGCATTCTGTAACAAAGCGGGACCAAAGCCATGACAAAAACGCG**TAACAAAAGTGTCTATAA**TC**ACGGCAGAAAAGTCCACA**TTGATTATTTGCACGGCGTCACACTTTGCTATGCCA**TAGCATTTTTATCCAT**AAGAT**TAGCGGATCCTA**CCTGACGCTTTTTATCGCAACTCTCTACTGTTTCTCCATACCCG | 9 |
| AraCE | $P_{BAD\Delta30}$ | ATTGCCGTCACTGCGTCTTTTACTGGCTCTTCTCGCTAACCAAACCGGTAACCCCGCTTATTAAAAGCATTCTGTAACAAAGCGGGACCAAAGCCATGACAAAAACGCG**TAACAAAAGTGTCTATAA**TC**ACGGCAGAAAAGTCCACA**TTGATTATTTGCACGGCGTCACACTTTGCTATGCCA**TAGCATTTTTATCCAT**AAGAT**TAGCGGATCCTA**CCTGACGCTTTTTATCGCAACTCTCTACTGTTTCTCCATACCCG | Rational |
| AraCE | $P_{BAD\Delta135}$ | ACG**TAACAAAAGTGTCTATAA**TC**ACGGCAGAAAAGTCCACA**TTGATTATTTGCACGGCGTCACACTTTGCTATGCCA**TAGCATTTTTATCCAT**AAGAT**TAGCGGATCCTA**CCTGACGCTTTTTATCGCAACTCTCTACTGTTTCTCCATACCCG | Rational |
| AraCE | $P_{BADnoCRP}$ | A**GAAACCAATTGTCCATA**TTGCATCAGACATTGCCGTCACTGCGTCTTTTACTGGCTCTTCTCGCTAACCAAACCGGTAACCCCGCTTATTAAAAGCATTCTGTAACAAAGCGGGACCAAAGCCATGACAAAAACGCG**TAACAAAAGTGTCTATAA**TC**ACGGCAGAAAAGTCCACA**ATGTTGCGTTTCCTCGATCTTCTAACCGCACCCCCA**TAGCATTTTTATCCAT**AAGAT**TAGCGGATCCTA**CCTGACGCTTTTTATCGCAACTCTCTACTGTTTCTCCATACCCG | Rational |
| AraCE | $P_{BADmin}$ | **TAGCATTTTTATCCAT**AAGAT**TAGCGGATCCTA**CCTGACGCTTTTTATCGCAACTCTCTACTGTTTCTCCATACCCG | Rational |
| BetI | $P_{BetI}$ | AGCGCGGGTGAGAGGGATTCGTTACCAATAGACA**ATTGATTGGACGTTCAATATAA**TGCTAGC | 6, 10 |
| CinR | $P_{Cin}$ | CCCTTTGTGCGTCCAAACGGACGCACGGCGCTCTAAAGCGGGTCGCGATCTTTCAGATTCGCTCCTCGCGCTTTCAGTCTTTGTTTTGGCGCATGTCGTTATCGCAAAACCGCTGCACACTTTTGCGCGACATGCTCTGATCCCCCTCATCT**GGGGGGGCCTATCTGAGGGAA**TTTCCGATCCGGCTCGCCTGAACCATTCTGCTTTCCACGAACTTGAAAACGCT | iGEM R0078 |
| CymR | $P_{CymR}$ | GAAATCATAAAAAATTTATTTGCTTTCAGGAAAATTTTTCTGTATAATAGATTC**AACAAACAGACAATCTGGTCTGTTTGTATTAT** | Rational[11] |
| | $P_{CymRA}$ | GAAATCATCAGAACTTTATTTTCTTTCAGGAAAATTTTTCTGTATAATAGATTC**AACAAACAGACAATCTGGTCTGTTTGTATTAT** | Rational |
| | $P_{CymRB}$ | **AACAAACAGACAATCTGGTCTGTTTGTAT**TATTTTCTTTCAGGAAAATTTTTCTGTATAATAGATTC**AACAAACAGACAATCTGGTCTGTTTGTATTAT** | Rational |
| | $P_{CymRC}$ | **AACAAACAGACAATCTGGTCTGTTTGTAT**TATGGAAAATTTTTCTGTATAATAGATTC**AACAAACAGACAATCTGGTCTGTTTGTATTAT** | Rational |
| LacI | $P_{Tac}$ | TGTTGACAATTAATCATCGGCTCGTATAATGTGTGG**AATTGTGAGCGCTCACAATT** | 6 |
| | $P_{T5LacO}$ | AATCATAAAAAATTTATTTGCT**TTGTGAGCGGATAACAATT**ATAATAGATTCAA**TTGTGAGCGGATAACAATT** | pQE (c) |
| | $P_{B4}$ | **AATTGTGAGCGGATAACAATT**GAC**TTGTGAGCGGATAACAA**TGATACTTCGTGC | 12 |
| LuxR | $P_{Lux}$ | **ACCTGTAGGATCGTACAGGT**TTACGCAAGAAAATGGTTTGTTATAGTCGAATAAA | 13 |
| | $P_{LuxA}$ | **ACCTGTAGGATCGTACAGGT**TTACGCAAGAAAATGGTTTGTTATTGTCGAATAAA | Rational |
| | $P_{LuxB}$ | **ACCTGTAGGATCGTACAGGT**TTACGCAAGAAAATGGTTTGTTACAGTCGAATAAA | Rational |
| | $P_{Lux*}$ | **ACCTGTAGGATCGTACAGGT**TTACGCAAGAAAATGGTTTGTTACTTTCGAATAAA | 13 |
| MphR | $P_{Mph}$ | G**GATTGAATATAACCGACGTGACTGTTACATTTAGG**TGGCTAAACCCGTCAA | 7, 14 |
| NahR | $P_{Sal}$ | GGGGCCTCGCTTGGGTTATTGCTGGTGCCCGGCCGGG**CGCAATATTCATGTTGATGATTTATTATATATC**GAGTGGTGTATTTATCAATATTGTTTGCTCCGTTATCGTTATTAAC | 15 |
| | $P_{SalTTC}$ | GGGGCCTCGCTTGGGTTATTGCTGGTGCCCGGCCGGG**CGCAATATTCATGTTGATGATTTATTATATATC**GAGTGGTGTATTTATTTATATTGTTTGCTCCGTTACCGTTATTAAC | Evolved |
| | $P_{Nah}$ | ACATGACTACTTCAGACGATT**CGCAGTATTCACGCTGGTGATAAACAAATTCAACT**ATGCTTTATTGACAAATAAAAGCACGCTCACCATCATCGCGAATAC | 15-16 |
| PcaU | $P_{3B5}$ | **TTTTGTTCGATTATCGAACAAATTATTGAAATATCGAACAAAACCT**CTAAACTACTGTGGCACTGAATCAAAAAATTATAAACAATGATCAGA | 17 |
| | $P_{3B5A}$ | **TTTTGTTCGATTATCGAACAAATTATTGAAATATCGAACAAAACCT**CTAAACTACTGTGGCACTGAATCAAAAAATTATAAACTATGATCAGA | Rational |
| | $P_{3B5B}$ | **TTTTGTTCGATTATCGAACAAATTATTGAAATATCGAACAAAACCT**CTAAACTACTGTGGCACTGAATCAAAAAATTATAAACCATGATCAGA | Rational |
| | $P_{3B5C}$ | **TTTTGTTCGATTATCGAACAAATTATTGAAATATCGAACAAAACCT**CTAAACTACTGTGGCACTGAATCAAAAAATTATAAACCCTGATCAGA | Rational |
| PhlF | $P_{PhlF}$ | CGACGTACGGTGGAATCTGATTCGTTACCAATTGAC**ATGATACGAAACGTACCGTATCGTTAAGGT** | 10 |
| TetR | $P_{Tet*}$ | TTTTCAGCAGGACGCACTGACC**TCCCTATCAGTGATAGAGA**TTGACA**TCCCTATCAGTGATAGAGA**TACTGAGCAC | 13 |
| | $P_{D18}$ | TACAACGTCGTGTTAGCTGCCTTTCGTCTTCAATAATTCTTGACA**TCCCTATCAGTGATAGAGA**TACTTGAATA | 12 |
| | $P_{D79}$ | TACAACGTCGTGTTAGCTAGCCTTTCGTCTTCAATAATTCTTAGACA**TCCCTATCAGTGATAGAGA**TACTTGAATA | 12 |

| | | | |
|---|---|---|---|
| TetR | $P_{TetR}$ | **TCCCTATCAGTGATAGAGA**TTGACA**TCCCTATCAGTGATAGA**TATAATGAGCAC | 10 |
| TtgR | $P_{Ttg}$ | CACCCAGCAG**TATTTACAAACAACCATGAATGTAAGTA**TATTCCTTAGCAA | 7, 18 |
| VanR | $P_{Van}$ | **ATTGGATCCAA**TTGACAGCTAGCTCAGTCCTAGGTATA**ATTGGATCCAAT** | Rational |
| | $P_{VanCC}$ | **ATTGGATCCAA**TTGACAGCTAGCTCAGTCCTAGGTACC**ATTGGATCCAAT** | Evolved |
| SrpR | $P_{SrpR}$ | GATTCGTTACCAATTGACAGCTAGCTCAGTCCTAGGT**ATATACATACATGCTTGTTTGTTTGTAAAC** | 10 |
| σ70 | $P_{LacI}$ | GCGGCGCGCCATCGAATGGCGCAAAACCTTTCGCGGTATGGCATGATAGCGCCC | *E. coli* genome |
| | $P_{LacIQ}$ | GCGGCGCGCCATCGAATGGTGCAAAACCTTTCGCGGTATGGCATGATAGCGCCC | Rational[19] |
| | $P_{LacIR}$ | GCGGCGCGCCATCGAATGGTGAAAAACCTTTCGCGGTATGGCATGATAGCGCCC | Rational |
| | $P_{J23101}$ | TTTACAGCTAGCTCAGTCCTAGGTATTATGCTAGC | iGEM J23101 |
| | $P_{J23100}$ | TTGACGGCTAGCTCAGTCCTAGGTACAGTGCTAGC | iGEM J23100 |
| | $P_{J23119}$ | TTGACAGCTAGCTCAGTCCTAGGTATAATGCTAGC | iGEM J23119 |

a) -35 box, -10 box, and +1 underlined.  Operators in bold.  Evolved or rational point mutations in blue.
b) Rational mutations were made by altering the -10 box, adding or eliminating operators, or (where noted) inspired by previous work.  Evolved mutations arose during selection.
c) Qiagen (Hilden, Germany)

### Supplementary Table 2: Insulators

| Insulator | DNA sequence | Source |
|---|---|---|
| RiboJ | AGCTGTCACCGGATGTGCTTTCCGGTCTGATGAGTCCGTGAGGACGAAACAGCCTCTACAAATAATTTTGTTTAA | 20 |
| RiboJ10 | AGCGCTCAACGGGTGTGCTTCCCGTTCTGATGAGTCCGTGAGGACGAAAGCGCCTCTACAAATAATTTTGTTTAA | 6 |

### Supplementary Table 3: RBSs

| Protein | RBS | DNA sequence[a] | Source[b] |
|---|---|---|---|
| AcuR | acu1 | GGAAGAGAGTCAATT**CAGGGTGGT**GAAT | *E. coli* genome |
| | acu2 | GGAAGAGAGTCAATT**CAGGGAGGT**GAAT | Evolved |
| AraC | ara1 | GGAAGAGAGTCAATT**CAGGGTGGT**GAAT | *E. coli* genome |
| | ara2 | GGATCCCAATGATGTGGGGTCCCGCTTT**CAAGGAGGT**AAAT | RBS calculator[16, 21] |
| AraE | e1 | TAAGATCCTATTCCAGCGGGATTAA**AGAGGAGCGA**TTAAGC | Evolved |
| BetI | bet1 | GGAAGAGAGTCAATT**CAGGGTGGT**GAAT | *E. coli* genome |
| | bet2 | GGAAGAGAGTCAATT**CACGGGGGT**GAAT | Evolved |
| | bet3 | GGAAGAGAGTCATT**CATGGAGGT**GAAT | rational |
| | bet4 | GGAAGAGAGTCAATT**CACGGAGGT**GAAT | rational |
| CinR | cin1 | GGAAGAGAGTCAATT**CAGGGTGGT**GAAT | *E. coli* genome |
| | cin2 | AGGTCCGAGACGCCCGTC**AACGGAGAA**CGGCGA | RBS calculator[16, 21] |
| CymR | cym1 | GGAAGAGAGTCAATT**CAGGGTGGT**GAAT | *E. coli* genome |
| CymR | cym2 | AATGGAATATTCATAGTAAATACCA**AACGGAGAT**TCTT | rational |
| | cym3 | AATGGAATATTCATAGTAAATACCA**AACGGATAT**TCTT | RBS calculator[16, 21] |
| LacI | lac1 | GGAAGAGAGTCAATT**CAGGGTGGT**GAAT | *E. coli* genome |
| | lac2 | GGAAGAGAGTCAATT**CATGGAGGT**GAAT | Rational |
| | lac3 | GGAAGAGAGTCAATT**CAGGGTAGA**GAAC | Rational |
| | lac4 | GGAAGAGAGTCAATT**CAGGGTAGA**GAAT | Rational |
| | lac5 | GGAAGAGAGTCAATT**CAGGGCGGT**GAAT | Rational |

| | | | |
|---|---|---|---|
| LuxR | lux1 | GGAAGAGAGTCAATTCAGGGTGGTGAAT | *E. coli* genome |
| | lux2 | AGGTACTACGCGAGGACTTAAGCCT**TTCGGAGGG**GCAGA | RBS calculator[16, 21] |
| MphR | mpr1 | GGAAGAGAGTCAATT**CAGGGTGGT**GAAT | *E. coli* genome |
| | mph2 | GGAAGAGAGTCAATT**TAGGGAGGT**GAAT | evolved |
| | mph3 | GGAAGAGAGTCAATT**CAGGGAGGT**GAAT | Rational |
| NahR | nah1 | GGAAGAGAGTCAATT**CAGGGTGGT**GAAT | *E. coli* genome |
| | nah2 | GGAAGAGAGTCAATT**CATGGGGGT**TGAAT | Evolved |
| | nah3 | ACCCCCTATAAGAA**AAAGACTTA**ACTATCC | RBS calculator[16, 21] |
| PcaU | pca1 | GGAAGAGAGTCAATT**CAGGGTGGT**GAAT | *E. coli* genome |
| | pca2 | GGAAGAGAGTCAATT**CAGGGAGGT**GAAT | Evolved |
| | pca3 | CGCTTACAATAGACGAACAAT**AAAGGAGGA**ATTAACCG | Rational |
| | pca4 | CGCTTACAATAGACGAACAAT**TAAGGAGAA**ATTAACCG | RBS calculator[16, 21] |
| PhlF | phl1 | GGAAGAGAGTCAATT**CAGGGTGGT**GAAT | *E. coli* genome |
| | phl2 | GGAAGAGAGTCAATT**CATGGGGGT**GAAT | Evolved |
| | phl3 | CTATGGACTATGTTTGA**AAGGGAGAA**ATACTAG | [6] |
| | phl4 | GGAAGAGAGTCAATT**CAAGGGGGT**GAAT | Rational |
| | phl5 | GGAAGAGAGTCAATT**CAAGGTGGT**GAAT | Rational |
| | phl6 | CTATGGACTATGTTTGA**AAGGGATAC**ATACTAG | Rational |
| TetR | tet1 | GGAAGAGAGTCAATT**CAGGGTGGT**GAAT | *E. coli* genome |
| | tet2 | GTAATAAT**CAAGGTGGC**AAAAA | Rational |
| | tet3 | GTAATAAT**CCAGGAGGA**AAAAA | RBS calculator[16, 21] |
| | tet4 | GTAATAAT**CCGGGAGGC**AAAAA | Rational |
| TtgR | ttg1 | GGAAGAGAGTCAATT**CAGGGTGGT**GAAT | *E. coli* genome |
| | ttg2 | GGAAGAGAGTCAATT**CAGGGAGGT**GAAT | Rational |
| | ttg3 | TTACGAATTAGA**TAAGTAGGT**AAAAC | Rational |
| | ttg4 | TTACGAATTAGA**TAAGAAGGT**AAAAC | RBS calculator[16, 21] |
| VanR | van1 | GGAAGAGAGTCAATT**CAGGGTGGT**GAAT | *E. coli* genome |
| | van2 | GGAAGAGAGTCAATT**CAGGGGGGT**GAATA | Evolved |
| | van3 | GCTTAAACTAACGAACGTAAA**TAAGGAGGA**TAGAC | RBS calculator[16, 21] |
| | van4 | GCTT**CAGGGGGGT**GAATA | Rational |
| | van5 | GCTT**TAAGGAGGT**GAATA | Rational |
| $Ery^R$ | ery1 | TAATATTGAAAAAGGAAGAGT | [7] |
| YFP | B0064 | TACTAGAGAAAGAGGGGAAATACTAG | iGEM B0064 |
| PheS | B0064 | TACTAGAGAAAGAGGGGAAATACTAG | iGEM B0064 |
| KOD | B0064 | TACTAGAGAAAGAGGGGAAATACTAG | iGEM B0064 |
| PK6 | B0064 | TACTAGAGAAAGAGGGGAAATACTAG | iGEM B0064 |
| SrpR | srp1 | GAGTCTATGGACTATGTTTTCACAGAGGAGGTACCAGG | [6] |

a) Evolved or rational point mutations in blue.
b) Rational mutations were made by altering the Shine-Dalgarno sequence (bold). Evolved mutations arose during selection. The RBS Calculator was used to design RBSs *de novo*.

### Supplementary Table 4: Proteins

| Protein | Amino acid sequence[a] | Source[b] |
|---|---|---|
| AcuR | MPLTDTPPSVPQKPRRGRPRGAPDASLAHQSLIRAGLEHLTEKGYSSVGVDEILKAARVPKGSFYHYFRNKADFGLALIEAYDTYFARLLDQAFLDGSLAPLARLRLFTRMAEEGMARHGFRRGCLVGNLGQEMGALPDDFRAALIGVLETWQRRTAQLFREAQACGELSADHDPDALAEAFWIGWEGAILRAKLELRPDPLHSFTRTFGRHFVTRTQE | Addgene 62569[7] |
| AcuR$^{AM}$ | MPLTDTPPSVPQKPRRGRPRGAPDASLAHQSLIRAGLEHLTEKGYSSVGVDEILKAARVPKGSFYHYFRNKADFGLALI**K**AYDTYFARLLDQAFLDGSLAPLARLRLFTRMAEEGMARHGFRRGCLVGNLGQEMGALPDDFRAALIGVLETWQ**H**RTAQLFREAQACGELSADHDPDALAEAFWIGWEGAILRAKLELRPDP**M**HSFTRTFGRHFVTRTQE | Evolved |
| AraC | MAEAQNDPLLPGYSFNAHLVAGLTPIEANGYLDFFIDRPLGMKGYILNLTIRGQGVVKNQGREFVCRPGDILLFPPGEIHHYGRHPEAREWYHQWVYFRPRAYWHEWLNWPSIFANTGFFRPDEAHQPHFSDLFGQIINAGQGEGRYSELLAINLLEQLLLRRMEAINESLHPPMDNRVREACQYISDHLADSNFDIASVAQHVCLSPSRLSHLFRQQLGISVLSWREDQRISQAKLLLSTTRMPIATVGRNVGFDDQLYFSRVFKKCTGASPSEFRAGCEEKVNDVAVKLS | 9 |
| AraC$^{AM}$ | MAEAQNDPLLPGYSFNAHLVAGLTPIEANGYLDFFIDRPLGMKGYILNLTIRGQGVVKNQGREFVCRPGDILLFPPGEIHHYGRHPEAREWYHQWVYFRPRAYWHEWLNWPSIFANTGFFRPDEAHQPHFSD**F**FGQIINAGQGEGRYSELLAINLLEQLLLRRM**L**AIN**G**SLHPPMDNRVREACQYISDHLADSNFDIASVAQHVCLSPSRLSHLFRQQLGISVLSWREDQRISQAKLLLSTTRMPIATVGRNVGFDDQLYFSRVFKKCTGASPSEFRAG**L**EEKVNDVAVKLS | Evolved |
| AraE | MVTINTESALTPRSLRDTRRMNMFVSVAAAVAGLLFGLDIGVIAGALPFITDHFVLTSRLQEWVVSSMMLGAAIGALFNGWLSFRLGRKYSLMAGAILFVLGSIGSAFATSVEMLIAARVVLGIAVGIASYTAPLYLSEMASENVRGKMISMYQLMVTLGIVLAFLSDTAFSYSGNWRAMLGVLALPAVLLIILVVFLPNSPRWLAEKGRHIEAEEVLRMLRDTSEKAREELNEIRESLKLKQGGWALFKINRNVRRAVFLGMLLQAMQQFTGMNIIMYYAPRIFKMAGFTTTEQQMIATLVVGLTFMFATFIAVFTVDKAGRKPALKIGFSVMALGTLVLGYCLMQFDNGTASSGLSWLSVGMTMMCIAGYAMSAAPVVWILCSEIQPLKCRDFGITCSTTTNWVSNMIIGATFLTLLDSIGAAGTFWLYTALNIAFVGITFWLIPETKNVTLEHIERKLMAGEKLRNIGV | Addgene 18987[9] |
| BetI | MPKLGMQSIRRRQLIDATLEAINEVGMHDATIAQIARRAGVSTGIISHYFRDKNGLLEATMRDITSQLRDAVLNRLHALPQGSAEQRLQAIVGGNFDETQVSSAAMKAWLAFWASSMHQPMLYRLQQVSSRRLLSNLVSEFRRELPREQAQEAGYGLAALIDGLWLRAALSGKPLDKTRANSLTRHFITQHLPTD | 10 |
| BetI$^{AM}$ | MPKLGMQSIRRRQLIDATLEAINEVGMHDATIAQIARRAGVSTGIISHYFRDKNGLLEATMRDITSQLRDAVLNRLHALPQGSAEQRLQAIVGGNFDETQVSSAAMKAWLAFWA**I**SMHQPMLYRLQQVSSRRLLSNLVSEFRRELPREQAQEAGYGLAALIDGLWLRAALSGKPLDKTRANSLTRHFITQHLPTD | Evolved |
| CinR | MIENTYSEKFESAFEQIKAAANVDAAIRILQAEYNLDFVTYHLAQTIASKIDSPFVRTTYPDAWVSRYLLNSYVKVDPIVKQGFERQLPFDWSEVEPTPEAYAMLVDAQKHGIGGNGYSIPVADKAQRRALLSLNARIPADEWTELVRRCRNEWIEIAHLIHRKAVYELHGENDPVPALSPREIECLHWTALGKDYKDISVILGISEHTTRDYLKTARFKLGCATISAAASRAVQLRIINP | iGEM C0077[22-23] |
| CinR$^{AM}$ | MIENTYSEKFESAFEQIKAAANVDAAIRILQAEYNLDFVTYHLAQTIASKIDSPFVRTTYPDAWVSRYLLN**C**YVKVDPI**I**KQGFERQLPFDWSEVEPTPEAYAMLVDAQKHGI**DD**NGYSIPVADKAQRRALLSLNA**H**IPADEWTELVRRCRNEWIEIAHLIHRKAVYELHGENDPVPALSPREIECLHWTALGKDYKDISVILGISEHTTRDYLKTARF**R**LGC**T**TISAAASRAVQLRIINP**YRIRMTRRNW** | Evolved |
| CymR | MSPKRRTQAERAMETQGKLIAAALGVLREKGYAGFRIADVPGAAGVSRGAQSHHFPTKLELLLATFEWLYEQITERSRARLAKLKPEDDVIQQMLDDAAEFFLDDDFSISLDLIVAADRDPALREGIQRTVERNRFVVEDMWLGVLVSRGLSRDDAEDILWLIFNSVRGLAVRSLWQKDKERFERVRNSTLEIARERYAKFKR | 10-11 |
| CymR$^{AM}$ | MSPKRRTQAERAMETQGKLIAAALGVLREKGYAGFRIADVPGAAGVSRGAQSHHFPTKLELLLATFEWLYEQITERSRARLAKLKPEDDVIQQMLDDAAEFFLDDDFSI**G**LDLIVAADRDPALREGIQRTVERNRFVVEDMWLGVLVSRGLSRDDAEDILWLIFNSVRGL**V**VRSLWQKDKERFERVRNSTLEIARERYAKFKR | Evolved |
| LacI | MKPVTLYDVAEYAGVSYQTVSRVVNQASHVSAKTREKVEAAMAELNYIPNRVAQQLAGKQSLLIGVATSSLALHAPSQIVAAIKSRADQLGASVVVSMVERSGVEACKAAVHNLLAQRVSGLIINYPLDDQDAIAVEAACTNVPALFLDVSDQTPINSIIFSHEDGTRLGVEHLVALGHQQIALLAGPLSSVSARLRLAGWHKYLTRNQIQPIAEREGDWSAMSGFQQTMQMLNEGIVPTAMLVANDQMALGAMRAITESGLRVGADISVVGYDDTEDSSCYIPPLTTIKQDFRLLGQTSVDRLLQLSQGQAVKGNQLLPVSLVKRKTTLAPNTQTASPRALADSLMQLARQVSRLESGQ | 13 |
| LacI$^{AM}$ | MKPVTLYDVAEYAGVSY**M**TVSRVVNQASHVSAKTREKVEAAM**V**ELNYIPNRVAQQLAGKQSLLIGVATSSLALHAPSQIVAAIKSRADQLGASVVVSMVERSGVEACKAAVHNLLAQRVSGLIINYPLDDQDAIAVEAACTNVPALFLDVSDQTPINSII**Y**SHEDGTRLGVEHLVALGHQQIALLAGPLSSVSARLRLAGWHKYLTRNQIQPIAEREGDWSAMSGFQQTMQMLNEGIVPTAMLVANDQMALGAMRAITESGLRVGADISVVGYDDTEDSSCYIPPLTTIKQDFRLLGQTSVDRLLQLSQGQAVKGNQLLPVSLVKRKTTLAPNTQTASPRALADSLMQLARQVSRLESGQ | Evolved |
| LuxR | MKNINADDTYRIINKIKACRSNNDINQCLSDMTKMVHCEYYLLAIIYPHSMVKSDISILDNYPKKWRQYYDDANLIKYDPIVDYSNSNHSPINWNIFENNAVNKKSPNVIKEAKTSGLITGFSFPIHTANNGFGMLSFAHSEKDNYIDSLFLHACMNIPLIVPSLVDNYRKINIANNKSNNDLTKREKECLAWACEGKSSWDISKILGCSERTVTFHLTNAQMKLNTTNRCQSISKAILTGAIDCPYFKN | 13 |
| MphR | MPRPKLKSDDEVLEAATVVLKRCGPIEFTLSGVAKEVGLSRAALIQRFTNRDTLLVRMMERGVEQVRHYLNAIPIGAGPQGLWEFLQVLVRSMNTRNDFSVNYLISWYELQVPELRTLAIQRNRAVVEGIRKRLPPGAPAAAELLLHSVIAGATMQWAVDPDGELADHVLAQIAAILCLMFPEHDDFQLLQAHA | Addgene 62570[7] |
| MphR$^{AM}$ | MPRPKLKSDDEVLEAATVVLKRCGPIEFTLSGVAKEVGLSRAALIQRFTNRDTLLVRMMERGVEQVRHYLNAIPIGAGPQGLWEFLQVLVRSM**D**TRNDFSVNYLISWYELQVPELRTLAIQRNRAVVEGIRKRLPPGAPAAAELLLHSVIAGATMQWAVDPDGELADHVLAQIAAILCLMFPEHDDFQLLQAHA | Evolved |
| NahR | MELRDLDLNLLVVFNQLLVDRRVSITAENLGLTQPAVSNALKRLRTSLQDPLFVRTHQGMEPTPYAAHLAEPVTSAMHALRNALQHHESFDPLTSERTFTLAMTDIGEIYFMPRLMDVLAHQAPNCVISTVRDSSMSLMQALQNGTVDLAVGLLPNLQTGFFQRRLLQNHYVCLCRKDHPVTREPLTLERFCSYGHVRVIAAGTGHGEVDTYMTRVGIRRDIRLEVPHFAAVGHILQRTDLLATVPIRLADCCVEPFGLSALPHPVVLPEIAINMFWHAKYHKDLANIWLRQLMFDLFTD | 16 |
| NahR$^{AM}$ | MELRDLDLNLLVVFNQLLVDRRVS**V**TAENLGLTQPAVSNALKRLRTSLQDPLFVRTHQGMEPTPYAAHLAE**H**VTSAMHALRNALQHHESFDPLTSERTFTLAMTDIGEIYFMPRLMD**A**LAHQAPNCVISTVRDSSMSLMQALQNGTVDLAVGLLPNLQTGFFQRRLLQNHYVCLCRKDHPVTREPLTLERFCSYGHVRVIAAGTGHGEVDTYMTRVGIRRDIRLEVPHFAAVGHILQRTDLLATVPI**C**LADCCVEPFGLSALPHPVVLPEIAINMFWHAKYHKDLANIWLRQLMFDLFTD | Evolved |
| PcaU | MWSNMDDKKVKEEKILHNSTNKKIIRHEDFVAGISKGMAILDSFGTDRHRLNITMAAEKTGMTRAAARRHLLTLEYLGYLESDGHYFYLTPKILKFSGSYLGGAQLPKISQPLLNLLTTQTSLIYSVMVLDGYEAITIARSAAHQQTDRVNPYGLHLGNRLPAHATSAGKILLAYLDDHAQQEWLNQYPLQRLTKYTYTNNIDFLRLLSEIKEQGWCYSSEEHELGVHALAVPIYGQQSRVVAALNIVSPTMRTTKEYLIQHILPLLQETARELRNIL | 17, 24 |
| PcaU$^{AM}$ | MWSNMDDKKVKEE**N**ILHNSTNKKIIRHEDFVAGISKGMAILDSFGTDRHRLNITMAAEKTGMTRAAARRHLLTLEYLGYLESDGHYFYLTPKILKFSGSYLGGAQLPKISQPLLNLLTTQTSLIYSVMVLDGYEAITIARSAAHQQTDRVNPYGLHLGNRLPAH**T**TSAGKILLAYLDDHAQQEWLNQYPLQRLTKYTYTN**H**IDFLRLLSEIKEQGWCYSSEEHELGVHALAVPIYGQQSRVVAALNIVSPTMRTTKEYLIQHILPLLQETARELRNIL | Evolved |
| PhlF | MARTPSRSSIGSLRSPHTHKAILTSTIEILKECGYSGLSIESVARRAGASKPTIYRWWTNKAALIAEVYENESEQVRKFPDLGSFKADLDFLLRNLWKVWRETICGEAFRCVIAEAQLDPATLTQLKDQFMERRREMPKKLVENAISNGELPKDTNRELLLDMIFGFCWYRLLTEQLTVEQDIEEFTFLLINGVCPGTQR | 10 |

| | | |
|---|---|---|
| $\text{PhlF}^{\text{AM}}$ | MARTPSRSSIGSLRSPHTHKAILTSTIEILKECGYSGLSIESVARRAGA**G**KPTIYRWWTNKAALIAEVYENE**I**EQVRKFPDLGSFKADLDFLL<br>**H**NLWKVWRETICGEAFRCVIAEAQLDP**V**TLTQLKDQFMERRRE**I**PKKLVE**D**AISNGELPKD**I**NRELLLDMIFGFCWYRLLTEQLTVEQDIEEF<br>TFLLINGVCPGTQ**C** | Evolved |
| TetR | MSRLDKSKVINSALELLNEVGIEGLTTRKLAQKLGVEQPTLYWHVKNKRALLDALAIEMLDRHHTHFCPLEGESWQDFLRNNAKSFRCALLSH<br>RDGAKVHLGTRPTEKQYETLENQLAFLCQQGFSLENALYALSAVGHFTLGCVLEDQEHQVAKEERETPTTDSMPPLLRQAIELFDHQGAEPAF<br>LFGLELIICGLEKQLKCESGS | 13 |
| TtgR | MVRRTKEEAQETRAQIIEAAERAFYKRGVARTTLADIAELAGVTRGAIYWHFNNKAELVQALLDSLHETHDHLARASESEDEVDPLGCMRKLL<br>LQVFNELVLDARTRRINEILHHKCEFTDDMCEIRQQRQSAVLDCHKGITLALANAVRRGQLPGELDAERAAVAMFAYVDGLIRRWLLLPDSVD<br>LLGDVEKWVDTGLDMLRLSPALRK | 7, 10 |
| $\text{TtgR}^{\text{AM}}$ | MVRRTKEEAQETRAQIIEAAERAFYKRGVARTTLADIAELAGVTRGAIYWHFNNKAELVQALLDSLHETHDHLARASESEDEVDPLGCMRKLL<br>LQVFNELVLDARTRRINEILHHKCEFTDDMCEIRQQ**H**QSAVLDCHKGITL**T**LAN**V**VRRGQLPGELDAERAAVAMFAYVDGLIRRWLLLPDSVD<br>LLGDVEKWVDTGLDMLRLSPALRK | Evolved |
| VanR | MDMPRIKPGQRVMMALRKMIASGEIKSGERIAEIPTAAALGVSRMPVRTALRSLEQEGLVVRLGARGYAARGVSSDQIRDAIEVRGVLEGFAA<br>RRLAERGMTAETHARFVALIAEGEALFAAGRLNGEDLDRYAAYNQAFHDTLVSAAGNGAVESALARNGFEPFAAAGALALDLMDLPAEYEHLL<br>AAHRQHQAVLDAVSCGDAEGAERIMRDHALAAIRNAKVFEAAASAGAPLGAAWSIRAD | 25 |
| $\text{VanR}^{\text{AM}}$ | MDMPRIKPGQRVMMALRKMIASGEIKSGERIAEIPTAAALGVSRMPVR**I**ALRSLEQEGLVVRLGARGYAARGVSSDQIRDAIEVRGVLEGFAA<br>RRLAERGMTAETHARFV**V**LIAEGEALFAAGRLNGEDLDRYAAYNQAFHDTLVSAAGNGAVESALARNGFEPFAAAGALALDLMDL**S**AEYEHLL<br>AAHRQHQAVLDAVSCGDAEGAERIMRDHALAAIRNAKVFEAAASAGAPLGAAWSIRAD | Evolved |
| $\text{Ery}^{\text{R}}$ | MNEKNIKHSQNFITSKHNIDKIMTNIRLNEHDNIFEIGSGKGHFTLELVKRCNFVTAIEIDHKLCKTTENKLVDHDNFQVLNKDILQFKFPKN<br>QSYKIYGNIPYNISTDIIRKIVFDSIANEIYLIVEYGFAKRLLNTKRSLALLLMAEVDISILSMVPREYFHPKPKVNSSLIRLSRKKSRISHK<br>DKQKYNYFVMKWVNKEYKKIFTKNQFNNSLKHAGIDDLNNISFEQFLSLFNSYKLFNK | Addgene 62570[7] |
| YFP | MVSKGEELFTGVVPILVELDGDVNGHKFSVSGEGEGDATYGKLTLKFICTTGKLPVPWPTLVTTFGYGLQCFARYPDHMKLHDFFKSAMPEGY<br>VQERTIFFKDDGNYKTRAEVKFEGDTLVNRIELKGIDFKEDGNILGHKLEYNYNSHNVYIMADKQKNGIKVNFKIRHNIEDGSVQLADHYQQN<br>TPIGDGPVLLPDNHYLSYQSALSKDPNEKRDHMVLLEFVTAAGITLGMDELYK | 6 |
| PheS | MSHLAELVASAKAAISQASDVAALDNVRVEYLGKKGHLTLQMTTLRELPPEERPAAGAVINEAKEQVQQALNARKAELESAALNARLAAETID<br>VSLPGRRIENGGLHPVTRTIDRIESFFGELGFTVATGPEIEDDYHNFDALNIPGHHPARADHDTFWFDTTRLLRTQTSGVQIRTMKAQQPPIR<br>IIAPGRVYRNDYDQTHTPMFHQMEGLIVDTNISFTNLKGTLHDFLRNFFEEDLQIRFRPSYFPFTEPSAEVDVMGKNGKWLEVLGCGMVHPNV<br>LRNVGIDPEVYSGFGFGMGMERLTMLRYGVTDLRSFFENDLRFLKQFK | 26 |
| KOD | MILDTDYITEDGKPVIRIFKKENGEFKIEYDRTFEPYFYALLKDDSAIEEVKKITAERHGTVVTVKRVEKVQKKFLGRPVEVWKLYFTHPQDV<br>PAIRDKIREHPAVIDIYEYDIPFAKRYLIDKGLVPMEGDEELKMLAFDIETLYHEGEEFAEGPILMISYADEEGARVITWKNVDLPYVDVVST<br>EREMIKRFLRVVKEKDPDVLITYNGDNFDFAYLKKRCEKLGINFALGRDGSEPKIQRMGDRFAVEVKGRIHFDLYPVIRRTINLPTYTLEAVY<br>EAVFGQPKEKVYAEEITTAWETGENLERVARYSMEDAKVTYELGKEFLPMEAQLSRLIGQSLWDVSRSSTGNLVEWFLLRKAYERNELAPNKP<br>DEKELARRRQSYEGGYVKEPERGLWENIVYLDFRSLYPSIIITHNVSPDTLNREGCKEYDVAPQVGHRFCKDFPGFIPSLLGDLLEERQKIKK<br>KMKATIDPIERKLLDYRQRAIKILANSYYGYYGYARARWYCKECAESVTAWGREYITMTIKEIEEKYGFKVIYSDTDGFFATIPGADAETVKK<br>KAMEFLKYINAKLPGALELEYEGFYKRGFFVTKKKYAVIDEEGKITTRGLEIVRRDWSEIAKETQARVLEALLKDGDVEKAVRIVKEVTEKLS<br>KYEVPPEKLVIHEQITRDLKDYKATGPHVAVAKRLAARGVKIRPGTVISYIVLKGSGRIGDRAIPFDEFDPTKHKYDAEYYIENQVLPAVERI<br>LRAFGYRKEDLRYQKTRQVGLSAWLKPKGTLE | 27 |
| PK6 | MILDTDYITEDGKPVIRIFKKENGEFKIEYDRTFEPYFYALLKDDSAIEEVKKITAERHGTVVTVKRVEKVQKKFLGRPVEVWKLYFTHPQDV<br>PAIRDKIREHPAVIDIYEYDIPFAKRYLIDKGLIPMEGEEELKILAFDIETLYHEGGEFGKGPIIMISYADENEAKVITWKNIDLPYVEVVSS<br>EREMIKRFLRIIREKDPDIIVTYNGDSFDFPYLAKRAEKLGIKLTIGRDGSEPKMQRIGDMTAVEVKGRIHFDLYPVIRRTINLPTYTLEAVY<br>EAVFGQPKEKVYAEEITTAWETGENLERVARYSMEDAKVTYELGKEFLPMEAQLSRLIGQSLWDVSRSSTGNLVEWFLLRKAYERNELAPNKP<br>DEKELARRRQSYEGGYVKEPERGLWENIVYLDFRSLYPSIIITHNVSPDTLNREGCKEYDVAPQVGHRFCKDFPGFIPSLLGHLLEERQKIKT<br>KMKETQDPIEKILLDYRQKAIKLLANSFYGYYGYARARWYCKECAESVTAWGREYITMTIKEIEEKYGFKVIYSDTDGFFATIPGADAETVKK<br>KAMEFLKYINAKLPGALELEYEGFYKRGFFVTKKKYAVIDEEGKITTRGLEIVRRDWSEIAKETQARVLEALLKDGDVEKAVRIVKEVTEKLS<br>KYEVPPEKLVIHEQITRDLKDYKATGPHVAVAKRLAARGVKIRPGTVISYIVLKGSGRIGDRAIPFDEFDPTKHKYDAEYYIENQVLPAVLRI<br>LEGFGYRKEDLRYQKTRQVGLSAWLKPKGTLE | 28 |
| SrpR | MARKTAAEAEETRQRIIDAALEVFVAQGVSDATLDQIARKAGVTRGAVYWHFNGKLEVLQAVLASRQHPLELDFTPDLGIERSWEAVVVAMLD<br>AVHSPQSKQFSEILIYQGLDESGLIHNRMVQASDRFLQYIHQVLRHAVTQGELPINLDLQTSIGVFKGLITGLLYEGLRSKDQQAQIIKVALG<br>SFWALLREPPRFLLCEEAQIKQVKSFE | 10 |

a) Evolved mutations in blue, bold, underlined.
b) Evolved mutations arose during selection.

**Supplementary Table 5: Terminators**

| Terminator | Sequence | Source |
|---|---|---|
| B0053 | TCCGGCAAAAAAACGGGCAAGGTGTCACCACCCTGCCCTTTTTCTTTAAAACCGAAAAGATTACTTCGCGTT | iGEM B0053 |
| ECK120017009 | GATCTAACTAAAAAGGCCGCTCTGCGGCCTTTTTTCTTTTCACT | 29 |
| ECK120033736 | AACGCATGAGAAAGCCCCCGGAAGATCACCTTCCGGGGGCTTTTTTATTGCGC | 29 |
| ECK120033737 | GGAAACACAGAAAAAAGCCCGCACCTGACAGTGCGGGCTTTTTTTTTCGACCAAAGG | 29 |
| IOT | TAATTGGTAACGAATCAGACAATTGACGGCTCGAGGGAGTAGCATAGGGTTTGCAGAATCCCTGCTTCGTCCATTTGAC<br>AGGCACATTATGCATCGATGATAAGCTGTCAAACATGAGCAGATCCTCTACGCCGGACGCATCGTGGCCGGCATCACCG<br>GCGCCACAGGTGCGGTTGCTGGCGCCTATATCGCCGACATCACCGATGGGGAAGATCGGGCTCGCCACTTCGGGCTCAT<br>GAGCAAATATTTTATCTG | 10 |
| L3S2P21 | CTCGGTACCAAATTCCAGAAAAGAGGCCTCCCGAAAGGGGGGCCTTTTTTCGTTTTGGTCC | 29 |
| L3S3P21 | CCAATTATTGAAGGCCTCCCTAACGGGGGGCCTTTTTTTGTTTCTGGTCTCCC | 29 |

### Supplementary Table 6: Marionette cluster sequence

| Cluster | Sequence[a,b] |
|---|---|
| Marionette | TAACCAATCAGGCTTCCTACTTACAGAATTGAGAAAAGAGGATGTGGAACGCTTACTAGCTCCGGCAAAAAAACGGGCAAGGTGTCACCACCCTGCCCTTTTTCTTTAAAACCGA<br>AAAGATTACTTCGCGTTGAGATTTTGAGACACAAGGTCGAATCGCACCAAGACAGGTTTGTCCATTGACAGCTAGCTCAGTCCTAGGTATAATGCTAGCCTATGGACTATGTTTG<br>AAAGGGAGAAATACTAGATGGCACGTACCCCGAGCCGTAGCAGCATTGGTAGCCTGCGTAGTCCGCATACCCATAAAGCAATTCTGACCAGCACCATTGAAATCCTGAAAGAATG<br>TGGTTATAGCGGTCTGAGCATTGAAAGCGTGGCACGTCGCGCCGGTGCAGGCAAACCGACCATTTATCGTTGGTGGACCAACAAAGCAGCACTGATTGCCGAAGTGTATGAAAAT<br>GAAATCGAACAGGTACGTAAATTTCCGGATTTGGGTAGCTTTAAAGCCGATCTGGATTTTCTGCTGCATAATCTGTGGAAAGTTTGGCGTGAAACCATTTGTGGTGAAGCATTTC<br>GTTGTGTTATTGCAGAAGCACAGTTGGACCCTGTAACCCTGACCCAACTGAAAGATCAGTTTATGGAACGTCGTCGTGAGATACCGAAAAAACTGGTTGAAGATGCCATTAGCAA<br>TGGTGAACTGCCGAAAGATATCAATCGTGAACTGCTGCTGGATATGATTTTTGGTTTTTGTTGGTATCGCCTGCTGACCGAACAGTTGACCGTTGAACAGGATATTGAAGAATTT<br>ACCTTCCTGCTGATTAATGGTGTTTGTCCGGGTACACAGTGTTGATAAAATGGAATATTCATAGTAAATACCAAACGGAGATTCTTATGAGCCCGAAACGTCGTACCCAGGCAGA<br>ACGTGCAATGGAAACCCAGGGTAAACTGATTGCAGCAGCACTGGGTGTTCTGCGTGAAAAAGGTTATGCAGGTTTTCGTATTGCAGATGTTCCGGGTGCAGCCGGTGTTAGCCGT<br>GGTGCACAGAGCCATCATTTTCCGACCAAACTGGAACTGCTGCTGGCAACCTTTGAATGGCTGTATGAGCAGATTACCGAACGTAGCCGTGCACGTCTGGCAAAACTGAAACCGG<br>AAGATGATGTTATTCAGCAGATGCTGGATGATGCAGCAGAATTTTTTCTGGATGATGATTTTAGCATCGGCCTGGATCTGATTGTTGCAGCAGATCGTGATCCGGCACTGCGTGA<br>AGGTATTCAGCGTACCGTTGAACGTAATCGTTTTGTTGTTGAAGATATGTGGCTGGGTGTGCTGGTGAGCCGTGGTCTGAGCCGTGATGATGCCGAAGATATTCTGTGGCTGATT<br>TTTAACAGCGTTCGTGGTCTGGTAGTTCGTAGCCTGTGGCAGAAAGATAAAGAACGTTTTGAACGTGTGCGTAATAGCACCCTGGAAATTGCACGTGAACGTTATGCAAAATTCA<br>AACGTTGAAGGTACTACGCGAGGACTTAAGCCTTTCGGAGGGGCAGAATGAAAAACATAAATGCCGACGACACATACAGAATAATTAATAAAATTAAAGCTTGTAGAAGCAATAA<br>TGATATTAATCAATGCTTATCTGATATGACTAAAATGGTACATTGTGAATATTATTTACTCGCGATCATTTATCCTCATTCTATGGTTAAATCTGATATTTCAATCCTAGATAAT<br>TACCCTAAAAAATGGAGGCAATATTATGATGACGCTAATTTGATAAAATATGATCCTATAGTAGATTATTCTAACTCCAATCATTCACCAATTAATTGGAATATATTTGAAAACA<br>ATGCTGTAAATAAAAAATCTCCAAATGTAATTAAAGAAGCGAAAACATCAGGTCTTATCACTGGGTTTAGTTTCCCTATTCATACGGCTAACAATGGCTTCGGAATGCTTAGTTT<br>TGCACATTCAGAAAAAGACAACTATATAGATAGTTTATTTTTACATGCGTGTATGAACATACCATTAATTGTTCCTTCTCTAGTTGATAATTATCGAAAAATAAATATAGCAAAT<br>AATAAATCAAACAACGATTTAACCAAAAGAGAAAAAGAATGTTTAGCGTGGGCATGCGAAGGAAAAAGCTCTTGGGATATTTCAAAAATATTAGGTTGCAGTGAGCGTACTGTCA<br>CTTTCCATTTAACCAATGCGCAAATGAAACTCAATACAACAAACCGCTGCCAAAGTATTTCTAAAGCAATTTTAACAGGAGCAATTGATTGCCCATACTTTAAAAATTGAGCTTA<br>AACTAACGAACGTAAATAAGGAGGATAGACATGGACATGCCTCGTATTAAACCGGGTCAGCGTGTTATGATGGCACTGCGTAAAATGATTGCAAGCGGTGAAATCAAAAGTGGTG<br>AACGTATTGCAGAAATTCCGACCGCAGCAGCACTGGGTGTTAGCCGTATGCCGGTTCGTATCGCACTGCGTTCACTGGAACAAGAAGGTCTGGTTGTTCGTCTGGGTGCACGTGG<br>TTATGCAGCCCGTGGTGTTAGCAGCGATCAGATTCGTGATGCAATTGAAGTTCGTGGTGTTCTGGAAGGTTTTGCAGCACGTCGTCTGGCAGAACGTGGTATGACCGCAGAAACC<br>CATGCACGTTTTGTTGTACTGATTGCAGAAGGTGAAGCACTGTTTGCAGCCGGTCGCCTGAATGGTGAAGATCTGGATCGTTATGCCGCATATAATCAGGCATTTCATGATACCC<br>TGGTTAGCGCAGCAGGTAATGGTGCAGTTGAAAGCGCACTGGCACGTAATGGTTTTGAACCGTTTGCAGCAGCCGGTGCACTGGCCCTGGATCTGATGGACCTGTCTGCCGAATA<br>TGAACATCTGCTGGCAGCACATCGTCAGCATCAGGCAGTTCTGGATGCAGTTAGCTGTGGTGATGCCGAAGGTGCAGAACGTATTATGCGTGATCATGCACTGGCAGCAATTCGT<br>AATGCAAAAGTTTTTGAAGCAGCAGCAAGCGCAGGCGCACCGCTGGGTGCAGCATGGTCAATTCGTGCAGATTGAATAGTAATTGGTAACGAATCAGACAATTGACGGCTCGAGG<br>GAGTAGCATAGGGTTTGCAGAATCCCTGCTTCGTCCATTTGACAGGCACATTATGCATCGATGATAAGCTGTCAAACATGAGCAGATCCTCTACGCCGGACGCATCGTGGCCGGC<br>ATCACCGGCGCCACAGGTGCGGTTGCTGGCGCCTATATCGCCGACATCACCGATGGGGAAGATCGGGCTCGCCACTTCGGGCTCATGAGCAAATATTTTATCTGGCGGCGCGCCA<br>TCGAATGGTGAAAAACCTTTCGCGGTATGGCATGATAGCGCCCGGAAGAGAGTCAATTCATGGAGGTGAATATGAAACCAGTAACGTTATACGATGTCGCAGAGTATGCCGGTGT<br>CTCTTATATGACCGTTTCCCGCGTGGTGAACCAGGCCAGCCACGTTTCTGCGAAAACGCGGGAAAAAGTGGAAGCGGCGATGGTGGAGCTGAATTACATTCCCAACCGCGTGGCA<br>CAACAACTGGCGGGCAAACAGTCGTTGCTGATTGGCGTTGCCACCTCCAGTCTGGCCCTGCACGCGCCGTCGCAAATTGTCGCGGCGATTAAATCTCGCGCCGATCAACTGGGTG<br>CCAGCGTGGTGGTGTCGATGGTAGAACGAAGCGGCGTCGAAGCCTGTAAAGCGGCGGTGCACAATCTTCTCGCGCAACGCGTCAGTGGGCTGATCATTAACTATCCGCTGGATGA<br>CCAGGATGCCATTGCTGTGGAAGCTGCCTGCACTAATGTTCCGGCGTTATTTCTTGATGTCTCTGACCAGACACCCATCAACAGTATTATTTACTCCCATGAGGACGGTACGCGA<br>CTGGGCGTGGAGCATCTGGTCGCATTGGGTCACCAGCAAATCGCGCTGTTAGCGGGCCCATTAAGTTCTGTCTCGGCGCGTCTGCGTCTGGCTGGCTGGCATAAATATCTCACTC<br>GCAATCAAATTCAGCCGATAGCGGAACGGGAAGGCGACTGGAGTGCCATGTCCGGTTTTCAACAAACCATGCAAATGCTGAATGAGGGCATCGTTCCCACTGCGATGCTGGTTGC<br>CAACGATCAGATGGCGCTGGGCGCAATGCGCGCCATTACCGAGTCCGGGCTGCGCGTTGGTGCGGATATCTCGGTAGTGGGATACGACGATACCGAAGATAGCTCATGTTATATC<br>CCGCCGTTAACCACCATCAAACAGGATTTTCGCCTGCTGGGGCAAACCAGCGTGGACCGCTTGCTGCAACTCTCTCAGGGCCAGGCGGTGAAGGGCAATCAGCTGTTGCCAGTCT<br>CACTGGTGAAAAGAAAAACCACCCTGGCGCCCAATACGCAAACCGCCTCTCCCCGCGCGTTGGCCGATTCATTAATGCAGCTGGCACGACAGGTTTCCCGACTGGAAAGCGGGCA<br>GTGAGTAATAATCAAGGTGGCAAAAAATGTCCAGATTAGATAAAAGTAAAGTGATTAACAGCGCATTAGAGCTGCTTAATGAGGTCGGAATCGAAGGTTTAACAACCCGTAAACT<br>CGCCCAGAAGCTAGGTGTAGAGCAGCCTACATTGTATTGGCATGTAAAAAATAAGCGGGCTTTGCTCGACGCCTTAGCCATTGAGATGTTAGATAGGCACCATACTCACTTTTGC<br>CCTTTAGAAGGGGAAAGCTGGCAAGATTTTTTACGTAATAACGCTAAAAGTTTTAGATGTGCTTTACTAAGTCATCGCGATGGAGCAAAAGTACATTTAGGTACACGGCCTACAG<br>AAAAACAGTATGAAACTCTCGAAAATCAATTAGCCTTTTTATGCCAACAAGGTTTTTCACTAGAGAATGCATTATATGCACTCAGCGCTGTGGGGCATTTTACTTTAGGTTGCGT<br>ATTGGAAGATCAAGAGCATCAAGTCGCTAAAGAAGAAAGGGAAACACCTACTACTGATAGTATGCCGCCATTATTACGACAAGCTATCGAATTATTTGATCACCAAGGTGCAGAG<br>CCAGCCTTCTTATTCGGCCTTGAATTGATCATATGCGGATTAGAAAAACAACTTAAATGTGAAAGTGGGTCCTAAGGATCCCAATGATGTGGGGTCCCGCTTTCAAGGAGGTAAA<br>TATGGCTGAAGCGCAAAATGATCCCCTGCTGCCGGGATACTCGTTTAATGCCCATCTGGTGGCGGGTTTAACGCCGATTGAGGCCAACGGTTATCTCGATTTTTTTATCGACCGA<br>CCGCTGGGAATGAAAGGTTATATTCTCAATCTCACCATTCGCGGTCAGGGGGTGGTGAAAAATCAGGGACGAGAATTTGTTTGCCGACCGGGTGATATTTTGCTGTTCCCGCCAG<br>GAGAGATTCATCACTACGGTCGTCATCCGGAGGCTCGCGAATGGTATCACCAGTGGGTTTACTTTCGTCCGCGCGCCTACTGGCATGAATGGCTTAACTGGCCGTCAATATTTGC<br>CAATACGGGGTTCTTTCGCCCGGATGAAGCGCACCAGCCGCATTTCAGCGACTTTTTGGGCAAATCATTAACGCCGGGCAAGGGGAAGGGCGCTATTCGGAGCTGCTGGCGATA<br>AATCTGCTTGAGCAATTGTTACTGCGGCGCATGCTAGCGATTAACGGATCGCTCCATCCACCGATGGATAATCGGGTACGCGAGGCTTGTCAGTACATCAGCGATCACCTGGCAG<br>ACAGCAATTTTGATATCGCCAGCGTCGCACAGCATGTTTGCTTGTCGCCGTCGCGTCTGTCACATCTTTTCCGCCAGCAGTTAGGGATTAGCGTCTTAAGCTGGCGCGAGGACCA<br>ACGTATCAGCCAGGCGAAGCTGCTTTTGAGCACCACCCGGATGCCTATCGCCACCGTCGGTCGCAATGTTGGTTTTGACGATCAACTCTATTTCTCGCGGGTATTTAAAAAATGC<br>ACCGGGGCCAGCCCGAGCGAGTTCCGTGCCGGTTTGGAAGAAAAAGTGAATGATGTAGCCGTCAAGTTGTCATGATAAGATCCTATTCCAGCGGGATTAAAGAGGAGCGATTAAG<br>CATGGTTACTATCAATACGGAATCTGCTTTAACGCCACGTTCTTTGCGGGATACGCGGCGTATGAATATGTTTGTTTCGGTAGCTGCTGCGGTCGCAGGATTGTTATTTGGTCTT<br>GATATCGGCGTAATCGCCGGAGCGTTGCCGTTCATTACCGATCACTTTGTGCTGACCAGTCGTTTGCAGGAATGGGTGGTTAGTAGCATGATGCTCGGTGCAGCAATTGGTGCGC<br>TGTTTAATGGTTGGCTGTCGTTCCGCCTGGGGCGTAAATACAGCCTGATGGCGGGGGCCATCCTGTTTGTACTCGGTTCTATAGGGTCCGCTTTTGCGACCAGCGTAGAGATGTT<br>AATCGCCGCTCGTGTGGTGCTGGGCATTGCTGTCGGGATCGCGTCTTACACCGCTCCTCTGTATCTTTCTGAAATGGCAAGTGAAAACGTTCGCGGTAAGATGATCAGTATGTAC<br>CAGTTGATGGTCACACTCGGCATCGTGCTGGCGTTTTTATCCGATACAGCGTTCAGTTATAGCGGTAACTGGCGCGCAATGTTGGGGGTTCTTGCTTTACCAGCAGTTCTGCTGA<br>TTATTCTGGTAGTATTTCTGCCAAATAGCCCGCGCTGGCTGGCGGAAAAGGGGCGTCATATTGAGGCGGAAGAAGTATTGCGTATGCTGCGCGATACGTCGGAAAAAGCGCGAGA<br>AGAACTCAACGAAATTCGTGAAAGCCTGAAGTTAAAACAGGGCGGTTGGGCACTGTTTAAGATCAACCGTAACGTCCGTCGTGCTGTGTTTCTCGGTATGTTGTTGCAGGCGATG<br>CAGCAGTTTACCGGTATGAACATCATCATGTACTACGCGCCGCGTATCTTCAAAATGGCGGGCTTTACGACCACAGAACAACAGATGATTGCGACTCTGGTCGTAGGGCTGACCT<br>TTATGTTCGCCACCTTTATTGCGGTGTTTACGGTAGATAAAGCAGGGCGTAAACCGGCTCTGAAAATTGGTTTCAGCGTGATGGCGTTAGGCACTCTGGTGCTGGGCTATTGCCT<br>GATGCAGTTTGATAACGGTACGGCTTCCAGTGGCTTGTCCTGGCTCTCTGTTGGCATGACGATGATGTGTATTGCCGGTTATGCGATGAGCGCCGCGCCAGTGGTGTGGATCCTG<br>TGCTCTGAAATTCAGCCGCTGAAATGCCGCGATTTCGGTATTACCTGTTCGACCACCACGAACTGGGTGTCGAATATGATTATCGGCGCGACCTTCCTGACACTGCTTGATAGCA<br>TTGGCGCTGCCGGTACGTTCTGGCTCTACACTGCGCTGAACATTGCGTTTGTGGGCATTACTTTCTGGCTCATTCCGGAAACCAAAAATGTCACGCTGGAACATATCGAACGCAA<br>ACTGATGGCAGGCGAGAAGTTGAGAAATATCGGCGTCTGAGATGTGGGGTCCGATCTAACTAAAAAGGCCGCTCTGCGGCCTTTTTTCTTTTCACTGGACAGTATTGAATGGACT<br>CTGATCAATGTTGACGGCTAGCTCAGTCCTAGGTACAGTGCTAGCGGAAGAGAGTCATTCATGGAGGTGAATATGCCGAAACTGGGTATGCAGAGCATTCGTCGTCGTCAGCTGA<br>TTGATGCAACCCTGGAAGCAATTAATGAAGTTGGTATGCATGATGCAACCATTGCACAGATTGCACGTCGTGCCGGTGTTAGCACCGGTATTATTAGCCATTATTTCCGCGATAA<br>AAACGGTCTACTGGAAGCAACCATGCGTGATATTACCAGCCAGCTGCGTGATGCAGTTCTGAATCGTCTGCATGCACTGCCGCAGGGTAGCGCAGAACAGCGTCTGCAGGCAATT<br>GTTGGTGGTAATTTTGATGAAACCCAGGTTAGCAGCGCAGCAATGAAAGCATGGCTGGCATTTTGGGCAATCAGCATGCATCAGCCGATGCTGTATCGTCTGCAGCAGGTTAGCA<br>GTCGTCGTCTGCTGAGCAATCTGGTTAGCGAATTTCGTCGTGAACTGCCTCGTGAACAGGCACAAGAGGCAGGTTATGGTCTGGCAGCACTGATTGATGGTCTGTGGCTGCGTGC<br>AGCACTGAGCGGTAAACCGCTGGATAAAACCCGTGCAAATAGCCTGACCCGTCATTTTATCACCCAGCATCTGCCGACCGATTGATTACGAATTAGATAAGTAGGTAAAACATGG<br>TTCGTCGTACCAAAGAAGAGGCACAAGAAACCCGTGCACAGATTATTGAAGCAGCAGAACGTGCATTCTATAAACGTGGTGTTGCACGTACCACCCTGGCAGATATTGCAGAACT<br>GGCAGGCGTTACCCGTGGTGCAATTTATTGGCATTTTAACAACAAAGCCGAACTGGTTCAGGCACTGCTGGATAGCCTGCATGAAACCCATGATCATCTGGCACGTGCAAGCGAA<br>AGCGAAGATGAAGTTGATCCGCTGGGTTGTATGCGTAAACTGCTGCTGCAGGTTTTTAATGAACTGGTTCTGGATGCACGTACCCGTCGTATTAATGAAATCCTGCATCACAAAT<br>GCGAGTTCACCGATGATATGTGTGAAATTCGTCAGCAGCATCAGAGCGCAGTTCTGGATTGTCATAAAGGTATTACCCTGACACTGGCAAATGTAGTTCGTCGTGGTCAGCTGCC<br>TGGTGAACTGGATGCAGAACGTGCCGCAGTTGCAATGTTTGCCTATGTTGATGGTCTGATTCGTCGTTGGCTGCTGCTGCCGGATAGCGTTGATCTGCTGGGTGATGTTGAAAAA<br>TGGGTTGATACCGGTCTGGATATGCTGCGTCTGAGTCCGGCACTGCGTAAATAATGACGCTTACAATAGACGAACAATAAAGGAGGAATTAACCGATGTGGTCGAACATGGATGA<br>CAAGAAAGTGAAAGAGGAGAATATTCTGCACAATTCCACCAACAAGAAGATCATCCGCCACGAAGATTTTGTAGCCGGCATTAGCAAAGGGATGGCGATTCTGGATTCGTTTGGT<br>ACAGATCGTCATCGCCTCAATATCACCATGGCCGCAGAGAAAACCGGTATGACACGTGCAGCAGCTCGTCGCCACCTGCTTACTCTGGAGTATCTGGGCTATCTGGAAAGTGACG<br>GCCACTACTTCTACTTAACTCCCAAAATCCTGAAATTCAGTGGTTCATATTTGGGTGGTGCTCAATTGCCGAAAATTTCCCAACCACTGTTGAACTTGCTTACGACCCAGACCAG<br>CCTGATTTACAGCGTGATGGTGTTGGATGGCTATGAAGCCATTACCATTGCGCGTTCTGCCGCTCATCAGCAAACCGACCGCGTTAACCCGTATGGTTTACATCTCGGGAATCGC<br>TTACCAGCGCATACAACGTCAGCGGGCAAAATCCTGTTAGCGTATTTGGATGACCATGCCCAGCAAGAGTGGCTCAATCAGTACCCTCTGCAACGGCTCACGAAATACACGTATA<br>CCAACCACATCGACTTTCTGCGCCTTTTGAGTGAAATCAAGGAACAGGGTTGGTGCTATAGTTCGGAAGAACACGAACTGGGAGTACACGCCCTTGCGGTTCCGATTTACGGACA<br>ACAGTCTCGCGTCGTAGCGGCACTGAACATTGTCAGCCCGACAATGCGGACCACGAAAGAATACCTGATTCAGCATATTCTGCCGTTACTGCAAGAAACTGCGCGTGAATTGCGC<br>AATATCCTGTAATGAACCCCCTATAAGAAAAAGACTTAACTATCCATGGAACTGCGTGACCTTGATTTAAACCTGCTGGTGGTGTTCAACCAGTTGCTGGTCGACAGACGCGTCT<br>CTGTCACTGCGGAGAACCTGGGCCTGACCCAGCCTGCCGTGAGCAATGCGCTGAAACGCCTGCGCACCTCGCTACAGGACCCACTCTTCGTGCGCACACATCAGGGAATGGAACC<br>CACACCCTATGCCGCGCATCTGGCCGAGCACGTCACTTCGGCCATGCACGCACTGCGCAACGCCCTACAGCACCATGAAAGCTTCGATCCGCTGACCAGCGAGCGTACCTTCACC<br>CTGGCCATGACCGACATTGGCGAGATCTACTTCATGCCGCGGCTGATGGATGCGCTGGCTCACCAGGCCCCCAATTGCGTGATCAGTACGGTGCGCGACAGTTCGATGAGCCTGA<br>TGCAGGCCTTGCAGAACGGAACCGTGGACTTGGCCGTGGGCCTGCTTCCCAATCTGCAAACTGGCTTCTTTCAGCGCCGGCTGCTCCAGAATCACTACGTGTGCCTATGTCGCAA |

GGACCATCCAGTCACCCGCGAACCCCTGACTCTGGAGCGCTTCTGTTCCTACGGCCACGTGCGTGTCATCGCCGCTGGCACCGGCCACGGCGAGGTGGACACGTACATGACACGG
GTCGGCATCCGGCGCGACATCCGTCTGGAAGTGCCGCACTTCGCCGCCGTTGGCCACATCCTCCAGCGCACCGATCTGCTCGCCACTGTGCCGATATGTTTAGCCGACTGCTGCG
TAGAGCCCTTCGGCCTAAGCGCCTTGCCGCACCCAGTCGTCTTGCCTGAAATAGCCATCAACATGTTCTGGCATGCGAAGTACCACAAGGACCTAGCCAATATTTGGTTGCGGCA
ACTGATGTTTGACCTGTTTACGGATTGATAAAGGTCCGAGACGCCCGTCAACGGAGAACGGCGAATGATTGAGAATACCTATAGCGAAAAGTTCGAGTCCGCGTTCGAACAGATC
AAAGCGGCGGCCAACGTGGATGCCGCCATCCGTATTCTCCAGGCGGAATATAACCTCGATTTCGTCACCTACCATCTCGCCCAGACAATCGCGAGCAAGATCGATTCGCCCTTCG
TGCGCACCACCTATCCGGATGCCTGGGTTTCCCGTTACCTCCTCAACTGCTATGTGAAGGTCGATCCGATCATCAAGCAGGGCTTCGAACGCCAGCTGCCCTTCGACTGGAGCGA
GGTCGAACCGACGCCGGAGGCCTATGCCATGCTGGTCGACGCCCAGAAACACGGCATCGATGACAATGGCTACTCCATCCCCGTCGCCGACAAGGCGCAGCGCCGCGCCCTGCTG
TCGCTGAATGCCCATATACCGGCCGACGAATGGACCGAGCTCGTGCGCCGCTGCCGCAATGAGTGGATCGAGATCGCCCATCTGATCCACCGCAAGGCCGTATATGAGCTGCATG
GCGAAAACGATCCGGTGCCGGCATTGTCGCCGCGCGAGATCGAGTGTCTGCACTGGACCGCCCTCGGCAAGGATTACAAGGATATTTCGGTCATCCTGGGCATATCAGAGCATAC
CACACGCGATTACCTGAAAACCGCCCGCTTCAGGCTCGGCTGCACCACGATCTCGGCCGCCGCGTCGCGGGCTGTTCAATTGCGCATCATCAATCCCTATAGGATCCGCATGACG
CGACGTAATTGGTAATAGGGATGAACTGTACAAATGAAACGCATGAGAAAGCCCCCGGAAGATCACCTTCCGGGGGCTTTTTTATTGCGCTCCTTGGCCCTCCATCCTTAGATAG
TTGCCGGGTCCATATGAATATCCTCCTTAGTTCCTATTCCGAAGTTCCTATTCTCTAGAAAGTATAGGAACTTCGGCGCGCCTACCTGTGACGGAAGATCACTTCGCAGAATAAA
TAAATCCTGGTGTCCCTGTTGATACCGGGAAGCCCTGGGCCAACTTTTGGCGAAAATGAGACGTTGATCGGCACGTAAGAGGTTCCAACTTTCACCATAATGAAATAAGATCACT
ACCGGGCGTATTTTTTGAGTTGTCGAGATTTTCAGGAGCTAAGGAAGCTAAAATGGAGAAAAAAATCACTGGATATACCACCGTTGATATATCCCAATGGCATCGTAAAGAACAT
TTTGAGGCATTTCAGTCAGTTGCTCAATGTACCTATAACCAGACCGTTCAGCTGGATATTACGGCCTTTTTAAAGACCGTAAAGAAAAATAAGCACAAGTTTTATCCGGCCTTTA
TTCACATTCTTGCCCGCCTGATGAATGCTCATCCGGAATTACGTATGGCAATGAAAGACGGTGAGCTGGTGATATGGGATAGTGTTCACCCTTGTTACACCGTTTTCCATGAGCA
AACTGAAACGTTTTCATCGCTCTGGAGTGAATACCACGACGATTTCCGGCAGTTTCTACACATATATTCGCAAGATGTGGCGTGTTACGGTGAAAACCTGGCCTATTTCCCTAAA
GGGTTTATTGAGAATATGTTTTTCGTCTCAGCCAATCCCTGGGTGAGTTTCACCAGTTTTGATTTAAACGTGGCCAATATGGACAACTTCTTCGCCCCCGTTTTCACCATGGGCA
AATATTATACGCAAGGCGACAAGGTGCTGATGCCGCTGGCGATTCAGGTTCATCATGCCGTTTGTGATGGCTTCCATGTCGGCAGAATGCTTAATGAATTACAACAGTACTGCGA
TGAGTGGCAGGGCGGGGCGTAAGGCGCGCCATTTAAATGAAGTTCCTATTCCGAAGTTCCTATTCTCTAGAAAGTATAGGAACTTCGAAGCAGCTCCAGCCTACACAATCGCTGA
CCAAATTCTCAGTGGTTGTCGCAGGCGGTGGAAGCACCTTTACGCCA

a) DNA sequence colors correspond to promoters (orange), RBSs (green), protein coding sequences (red), terminators (purple), spacers (black), Chloramphenicol resistance cassette (pink), and *glvC* homology regions (blue).
b) Sequence was verified by PCR and sequencing from the genome of Marionette.

**Supplementary Table 7: Plasmids parts list**

### Reporter only plasmids[a]

| Plasmid | Output promoter | Reporter | Origin | Resistance |
|---|---|---|---|---|
| pAJM.711 | $P_{PhlF}$ | *YFP* | p15A | Kan |
| pAJM.712 | $P_{CymRC}$ | *YFP* | p15A | Kan |
| pAJM.713 | $P_{LuxB}$ | *YFP* | p15A | Kan |
| pAJM.714 | $P_{VanCC}$ | *YFP* | p15A | Kan |
| pAJM.715 | $P_{Tac}$ | *YFP* | p15A | Kan |
| pAJM.717 | $P_{Tet*}$ | *YFP* | p15A | Kan |
| pAJM.716 | $P_{BAD}$ | *YFP* | p15A | Kan |
| pAJM.718 | $P_{BetI}$ | *YFP* | p15A | Kan |
| pAJM.719 | $P_{Ttg}$ | *YFP* | p15A | Kan |
| pAJM.1459 | $P_{3B5C}$ | *YFP* | p15A | Kan |
| pAJM.721 | $P_{SalTTC}$ | *YFP* | p15A | Kan |
| pAJM.944 | $P_{Cin}$ | *YFP* | p15A | Kan |

### Reporter and regulator plasmids[a]

| Plasmid | Output promoter | Reporter | Promoter | RBS | Regulator | Origin | Resistance |
|---|---|---|---|---|---|---|---|
| pAJM.847 | $P_{PhlF}$ | *YFP* | $P_{LacIQ}$ | phl2 | *phlF*$^{AM}$ | p15A | Kan |
| pAJM.657 | $P_{CymRC}$ | *YFP* | $P_{LacIQ}$ | cym1 | *cymR*$^{AM}$ | p15A | Kan |
| pAJM.474 | $P_{LuxB}$ | *YFP* | $P_{Lac}$ | lux1 | *luxR* | p15A | Kan |
| pAJM.773 | $P_{VanCC}$ | *YFP* | $P_{LacIQ}$ | van2 | *vanR*$^{AM}$ | p15A | Kan |
| pAJM.336 | $P_{Tac}$ | *YFP* | $P_{LacI}$ | lac1 | *lacI*$^{AM}$ | p15A | Kan |
| pAJM.011 | $P_{Tet*}$ | *YFP* | $P_{LacI}$ | tet1 | *tetR* | p15A | Kan |
| pAJM.677 | $P_{BAD}$ | *YFP* | $P_{LacIQ}$ | ara1/e1 | *araC*$^{AM}$ / *araE* | p15A | Kan |
| pAJM.683 | $P_{BetI}$ | *YFP* | $P_{LacIQ}$ | bet2 | *betI*$^{AM}$ | p15A | Kan |
| pAJM.661 | $P_{Ttg}$ | *YFP* | $P_{LacIQ}$ | ttg2 | *ttgR*$^{AM}$ | p15A | Kan |
| pAJM.690 | $P_{3B5B}$ | *YFP* | $P_{LacIQ}$ | pca2 | *pcaU*$^{AM}$ | p15A | Kan |
| pAJM.771 | $P_{SalTTC}$ | *YFP* | $P_{LacIQ}$ | nah2 | *nahR*$^{AM}$ | p15A | Kan |
| pAJM.1642 | $P_{Cin}$ | *YFP* | $P_{LacI}$ | cin1 | *cinR*$^{AM}$ | p15A | Kan |
| pAJM.884 | $P_{Acu}$ | *YFP* | $P_{LacIQ}$ | acu2 | *rhlR*$^{AM}$ | p15A | Kan |
| pAJM.969 | $P_{Mph}$ | *YFP* | $P_{LacIQ}$ | mph2/ery1 | *mphR*$^{AM}$ / *eryR* | p15A | Kan/Ery |

a) DNA sequence of all plasmids are provided in Supplementary Table 8

### Supplementary Table 8: Plasmid sequences

| Plasmid | Sequence[a] |
| --- | --- |
| pAJM.011<br><br>GenBank accession number: MH101732 | CCAATTATTGAAGGCCTCCCTAACGGGGGGCCTTTTTTTGTTTCTGGTCTCCCGCTTAACGATCGTTGGCTGTTTTCAGCAGGACGCACTGACCTCCCTATCAGTGATAGAGATT<br>GACATCCCTATCAGTGATAGAGATACTGAGCACAGCTGTCACCGGATGTGCTTTCCGGTCTGATGAGTCCGTGAGGACGAAACAGCCTCTACAAATAATTTTGTTTAATACTAGA<br>GAAAGAGGGGAAATACTAGATGGTGAGCAAGGGCGAGGAGCTGTTCACCGGGGTGGTGCCCATCCTGGTCGAGCTGGACGGCGACGTAAACGGCCACAAGTTCAGCGTGTCCGGC<br>GAGGGCGAGGGCGATGCCACCTACGGCAAGCTGACCCTGAAGTTCATCTGCACCACAGGCAAGCTGCCCGTGCCCTGGCCCACCCTCGTGACCACCTTCGGCTACGGCCTGCAAT<br>GCTTCGCCCGCTACCCCGACCACATGAAGCTGCACGACTTCTTCAAGTCCGCCATGCCCGAAGGCTACGTCCAGGAGCGCACCATCTTCTTCAAGGACGACGGCAACTACAAGAC<br>CCGCGCCGAGGTGAAGTTCGAGGGCGACACCCTGGTGAACCGCATCGAGCTGAAGGGCATCGACTTCAAGGAGGACGGCAACATCCTGGGGCACAAGCTGGAGTACAACTACAAC<br>AGCCACAACGTCTATATCATGGCCGACAAGCAGAAGAACGGCATCAAGGTGAACTTCAAGATCCGCCACAACATCGAGGACGGCAGCGTGCAGCTCGCCGACCACTACCAGCAGA<br>ACACCCCCAATCGGCGACGGCCCCGTGCTGCTGCCCGACAACCACTACCTTAGCTACCAGTCCGCCCTGAGCAAAGACCCCAACGAGAAGCGCGATCACATGGTCCTGCTGGAGTT<br>CGTGACCGCCGCCGGGATCACTCTCGGCATGGACGAGCTGTACAAGTAACTCGGTACCAAATTCCAGAAAAGAGGCCTCCCGAAAGGGGGGCCTTTTTTCGTTTTGGTCCAATGG<br>CGGCGCGCCATCGAATGGCGCAAAACCTTTCGCGGTATGGCATGATAGCGCCCGGAAGAGAGTCAATTCAGGGTGGTGAATATGTCCAGATTAGATAAAAGTAAAGTGATTAACA<br>GCGCATTAGAGCTGCTTAATGAGGTCGGAATCGAAGGTTTAACAACCCGTAAACTCGCCCAGAAGCTAGGTGTAGAGCAGCCTACATTGTATTGGCATGTAAAAAATAAGCGGGC<br>TTTGCTCGACGCCTTAGCCATTGAGATGTTAGATAGGCACCATACTCACTTTTGCCCTTTAGAAGGGGAAAGCTGGCAAGATTTTTTACGTAATAACGCTAAAAGTTTTAGATGT<br>GCTTTACTAAGTCATCGCGATGGAGCAAAAGTACATTTAGGTACACGGCCTACAGAAAAACAGTATGAAACTCTCGAAAATCAATTAGCCTTTTTATGCCAACAAGGTTTTTCAC<br>TAGAGAATGCATTATATGCACTCAGCGCTGTGGGGCATTTTACTTTAGGTTGCGTATTGGAAGATCAAGAGCATCAAGTCGCTAAAGAAGAAAGGGAAACACCTACTACTGATAG<br>TATGCCGCCATTATTACGACAAGCTATCGAATTATTTGATCACCAAGGTGCAGAGCCAGCCTTCTTATTCGGCCTTGAATTGATCATATGCGGATTAGAAAAACAACTTAAATGT<br>GAAAGTGGGTCCTGATAAGGATCCTAATTGGTAACGAATCAGAATTGACGGCTCGAGGGAGTAGCATAGGGTTTGCAGAATCCCTGCTTCGTCCATTTGACAGGCACATTATGCA<br>TCGATGATAAGCTGTCAAACATGAGCAGATCCTCTACGCCGGACGCATCGTGGCCGGCATCACCGGCGCCACAGGTGCGGTTGCTGGCGCCTATATCGCCGACATCACCGATGGG<br>GAAGATCGGGCTCGCCACTTCGGGCTCATGAGCAAATATTTTATCTGAGGTGCTTCCTCGCTCACTGACTCGCTGCACGAGGCAGACCTCAGCGCTAGCGGAGTGTATACTGGCT<br>TACTATGTTGGCACTGATGAGGGTGTCAGTGAAGTGCTTCATGTGGCAGGAGAAAAAAGGCTGCACCGGTGCGTCAGCAGAATATGTGATACAGGATATATTCCGCTTCCTCGCT<br>CACTGACTCGCTACGCTCGGTCGTTCGACTGCGGCGAGCGGAAATGGCTTACGAACGGGGCGGAGATTTCCTGGAAGATGCCAGGAAGATACTTAACAGGGAAGTGAGAGGGCCG<br>CGGCAAAGCCGTTTTTCCATAGGCTCCGCCCCCCTGACAAGCATCACGAAATCTGACGCTCAAATCAGTGGTGGCGAAACCCGACAGGACTATAAAGATACCAGGCGTTTCCCCC<br>TGGCGGCTCCCTCGTGCGCTCTCCTGTTCCTGCCTTTCGGTTTACCGGTGTCATTCCGCTGTTATGGCCGCGTTTGTCTCATTCCACGCCTGACACTCAGTTCCGGGTAGGCAGT<br>TCGCTCCAAGCTGGACTGTATGCACGAACCCCCCGTTCAGTCCGACCGCTGCGCCTTATCCGGTAACTATCGTCTTGAGTCCAACCCGGAAAGACATGCAAAAGCACCACTGGCA<br>GCAGCCACTGGTAATTGATTTAGAGGAGTTAGTCTTGAAGTCATGCGCCGGTTAAGGCTAAACTGAAAGGACAAGTTTTGGTGACTGCGCTCCTCCAAGCCAGTTACCTCGGTTC<br>AAAGAGTTGGTAGCTCAGAGAACCTTCGAAAAACCGCCCTGCAAGGCGGTTTTTTCGTTTTCAGAGCAAGAGATTACGCGCAGACCAAAACGATCTCAAGAAGATCATCTTATTA<br>AGGGGTCTGACGCTCAGTGGAACGAAAAATCAATCTAAAGTATATATGAGTAAACTTGGTCTGACAGTTACCATGAGCCATATTCAACGGGAAACGTCTTGCTCGAGGCCGCGAT<br>TAAATTCCAACATGGATGCTGATTTATATGGGTATAAATGGGCTCGCGATAATGTCGGGCAATCAGGTGCGACAATCTATCGATTGTATGGGAAGCCCGATGCGCCAGAGTTGTT<br>TCTGAAACATGGCAAAGGTAGCGTTGCCAATGATGTTACAGATGAGATGGTCAGACTAAACTGGCTGACGGAATTTATGCCTCTTCCGACCATCAAGCATTTTATCCGTACTCCT<br>GATGATGCATGGTTACTCACCACTGCGATCCCCGGGAAAACAGCATTCCAGGTATTAGAAGAATATCCTGATTCAGGTGAAAATATTGTTGATGCGCTGGCAGTGTTCCTGCGCC<br>GGTTGCATTCGATTCCTGTTTGTAATTGTCCTTTTAACAGCGATCGCGTATTTCGTCTCGCTCAGGCGCAATCACGAATGAATAACGGTTTGGTTGATGCGAGTGATTTTGATGA<br>CGAGCGTAATGGCTGGCCTGTTGAACAAGTCTGGAAAGAAATGCATAAGCTTTTGCCATTCTCACCGGATTCAGTCGTCACTCATGGTGATTTCTCACTTGATAACCTTATTTTT<br>GACGAGGGGAAATTAATAGGTTGTATTGATGTTGGACGAGTCGGAATCGCAGACCGATACCAGGATCTTGCCATCCTATGGAACTGCCTCGGTGAGTTTTCTCCTTCATTACAGA<br>AACGGCTTTTTCAAAAATATGGTATTGATAATCCTGATATGAATAAATTGCAGTTTCATTTGATGCTCGATGAGTTTTTCTAAAACACCCCTTGTATTACTGTTTATGTAAGCAG<br>ACAGTTTTATTGTTCATGATGATATATTTTTATCTTGTGCAATGTACATCAGAGATTTTGAGACACAA |
| pAJM.336<br><br>GenBank accession number: MH101731 | CCAATTATTGAAGGCCTCCCTAACGGGGGGCCTTTTTTTGTTTCTGGTCTCCCGCTTAACGATCGTTGGCTGTGTTGACAATTAATCATCGGCTCGTATAATGTGTGGAATTGTG<br>AGCGCTCACAATTAGCTGTCACCGGATGTGCTTTCCGGTCTGATGAGTCCGTGAGGACGAAACAGCCTCTACAAATAATTTTGTTTAATACTAGAGAAAGAGGGGAAATACTAGA<br>TGGTGAGCAAGGGCGAGGAGCTGTTCACCGGGGTGGTGCCCATCCTGGTCGAGCTGGACGGCGACGTAAACGGCCACAAGTTCAGCGTGTCCGGCGAGGGCGAGGGCGATGCCAC<br>CTACGGCAAGCTGACCCTGAAGTTCATCTGCACCACAGGCAAGCTGCCCGTGCCCTGGCCCACCCTCGTGACCACCTTCGGCTACGGCCTGCAATGCTTCGCCCGCTACCCCGAC<br>CACATGAAGCTGCACGACTTCTTCAAGTCCGCCATGCCCGAAGGCTACGTCCAGGAGCGCACCATCTTCTTCAAGGACGACGGCAACTACAAGACCCGCGCCGAGGTGAAGTTCG<br>AGGGCGACACCCTGGTGAACCGCATCGAGCTGAAGGGCATCGACTTCAAGGAGGACGGCAACATCCTGGGGCACAAGCTGGAGTACAACTACAACAGCCACAACGTCTATATCAT<br>GGCCGACAAGCAGAAGAACGGCATCAAGGTGAACTTCAAGATCCGCCACAACATCGAGGACGGCAGCGTGCAGCTCGCCGACCACTACCAGCAGAACACCCCCAATCGGCGACGGC<br>CCCGTGCTGCTGCCCGACAACCACTACCTTAGCTACCAGTCCGCCCTGAGCAAAGACCCCAACGAGAAGCGCGATCACATGGTCCTGCTGGAGTTCGTGACCGCCGCCGGGATCA<br>CTCTCGGCATGGACGAGCTGTACAAGTAACTCGGTACCAAATTCCAGAAAAGAGGCCTCCCGAAAGGGGGGCCTTTTTTCGTTTTGGTCCAATGGCGGCGCGCCATCGAATGGCG<br>CAAAACCTTTCGCGGTATGGCATGATAGCGCCCGGAAGAGAGTCAATTCAGGGTGGTGAATATGAAACCAGTAACGTTATACGATGTCGCAGAGTATGCCGGTGTCTCTTATATG<br>ACCGTTTCCCGCGTGGTGAACCAGGCCAGCCACGTTTCTGCGAAAACGCGGGAAAAAGTGGAAGCGGCGATGGTGGAGCTGAATTACATTCCCAACCGCGTGGCACAACAACTGG<br>CGGGCAAACAGTCGTTGCTGATTGGCGTTGCCACCTCCAGTCTGGCCCTGCACGCGCCGTCGCAAATTGTCGCGGCGATTAAATCTCGCGCCGATCAACTGGGTGCCAGCGTGGT<br>GGTGTCGATGGTAGAACGAAGCGGCGTCGAAGCCTGTAAAGCGGCGGTGCACAATCTTCTCGCGCAACGCGTCAGTGGGCTGATCATTAACTATCCGCTGGATGACCAGGATGCC<br>ATTGCTGTGGAAGCTGCCTGCACTAATGTTCCGGCGTTATTTCTTGATGTCTCTGACCAGACACCCATCAACAGTATTATTTACTCCCATGAGGACGGTACGCGACTGGGCGTGG<br>AGCATCTGGTCGCATTGGGTCACCAGCAAATCGCGCTGTTAGCGGGCCCATTAAGTTCTGTCTCGGCGCGTCTGCGTCTGGCTGGCTGGCATAAATATCTCACTCGCAATCAAAT<br>TCAGCCGATAGCGGAACGGGAAGGCGACTGGAGTGCCATGTCCGGTTTTCAACAAACCATGCAAATGCTGAATGAGGGCATCGTTCCCACTGCGATGCTGGTTGCCAACGATCAG<br>ATGGCGCTGGGCGCAATGCGCGCCATTACCGAGTCCGGGCTGCGCGTTGGTGCGGATATCTCGGTAGTGGGATACGACGATACCGAAGATAGCTCATGTTATATCCCGCCGTTAA<br>CCACCATCAAACAGGATTTTCGCCTGCTGGGGCAAACCAGCGTGGACCGCTTGCTGCAACTCTCTCAGGGCCAGGCGGTGAAGGGCAATCAGCTGTTGCCAGTCTCACTGGTGAA<br>AAGAAAAACCACCCTGGCGCCCAATACGCAAACCGCCTCTCCCCGCGCGTTGGCCGATTCATTAATGCAGCTGGCACGACAGGTTTCCCGACTGGAAAGCGGGCAGTGATAAGGA<br>TCCTAATTGGTAACGAATCAGACAATTGACGGCTCGAGGGAGTAGCATAGGGTTTGCAGAATCCCTGCTTCGTCCATTTGACAGGCACATTATGCATCGATGATAAGCTGTCAAA<br>CATGAGCAGATCCTCTACGCCGGACGCATCGTGGCCGGCATCACCGGCGCCACAGGTGCGGTTGCTGGCGCCTATATCGCCGACATCACCGATGGGGAAGATCGGGCTCGCCACT<br>TCGGGCTCATGAGCAAATATTTTATCTGAGGTGCTTCCTCGCTCACTGACTCGCTGCACGAGGCAGACCTCAGCGCTAGCGGAGTGTATACTGGCTTACTATGTTGGCACTGATG<br>AGGGTGTCAGTGAAGTGCTTCATGTGGCAGGAGAAAAAAGGCTGCACCGGTGCGTCAGCAGAATATGTGATACAGGATATATTCCGCTTCCTCGCTCACTGACTCGCTACGCTCG<br>GTCGTTCGACTGCGGCGAGCGGAAATGGCTTACGAACGGGGCGGAGATTTCCTGGAAGATGCCAGGAAGATACTTAACAGGGAAGTGAGAGGGCCGCGGCAAAGCCGTTTTTCCA<br>TAGGCTCCGCCCCCCTGACAAGCATCACGAAATCTGACGCTCAAATCAGTGGTGGCGAAACCCGACAGGACTATAAAGATACCAGGCGTTTCCCCCTGGCGGCTCCCTCGTGCGC<br>TCTCCTGTTCCTGCCTTTCGGTTTACCGGTGTCATTCCGCTGTTATGGCCGCGTTTGTCTCATTCCACGCCTGACACTCAGTTCCGGGTAGGCAGTTCGCTCCAAGCTGGACTGT<br>ATGCACGAACCCCCCGTTCAGTCCGACCGCTGCGCCTTATCCGGTAACTATCGTCTTGAGTCCAACCCGGAAAGACATGCAAAAGCACCACTGGCAGCAGCCACTGGTAATTGAT<br>TTAGAGGAGTTAGTCTTGAAGTCATGCGCCGGTTAAGGCTAAACTGAAAGGACAAGTTTTGGTGACTGCGCTCCTCCAAGCCAGTTACCTCGGTTCAAAGAGTTGGTAGCTCAGA<br>GAACCTTCGAAAAACCGCCCTGCAAGGCGGTTTTTTCGTTTTCAGAGCAAGAGATTACGCGCAGACCAAAACGATCTCAAGAAGATCATCTTATTAAGGGGTCTGACGCTCAGTG<br>GAACGAAAAATCAATCTAAAGTATATATGAGTAAACTTGGTCTGACAGTTACCATGAGCCATATTCAACGGGAAACGTCTTGCTCGAGGCCGCGATTAAATTCCAACATGGATGC<br>TGATTTATATGGGTATAAATGGGCTCGCGATAATGTCGGGCAATCAGGTGCGACAATCTATCGATTGTATGGGAAGCCCGATGCGCCAGAGTTGTTTCTGAAACATGGCAAAGGT<br>AGCGTTGCCAATGATGTTACAGATGAGATGGTCAGACTAAACTGGCTGACGGAATTTATGCCTCTTCCGACCATCAAGCATTTTATCCGTACTCCTGATGATGCATGGTTACTCA<br>CCACTGCGATCCCCGGGAAAACAGCATTCCAGGTATTAGAAGAATATCCTGATTCAGGTGAAAATATTGTTGATGCGCTGGCAGTGTTCCTGCGCCGGTTGCATTCGATTCCTGT<br>TTGTAATTGTCCTTTTAACAGCGATCGCGTATTTCGTCTCGCTCAGGCGCAATCACGAATGAATAACGGTTTGGTTGATGCGAGTGATTTTGATGACGAGCGTAATGGCTGGCCT<br>GTTGAACAAGTCTGGAAAGAAATGCATAAGCTTTTGCCATTCTCACCGGATTCAGTCGTCACTCATGGTGATTTCTCACTTGATAACCTTATTTTTGACGAGGGGAAATTAATAG<br>GTTGTATTGATGTTGGACGAGTCGGAATCGCAGACCGATACCAGGATCTTGCCATCCTATGGAACTGCCTCGGTGAGTTTTCTCCTTCATTACAGAAACGGCTTTTTCAAAAATA<br>TGGTATTGATAATCCTGATATGAATAAATTGCAGTTTCATTTGATGCTCGATGAGTTTTTCTAAAACACCCCTTGTATTACTGTTTATGTAAGCAGACAGTTTTATTGTTCATGA<br>TGATATATTTTTATCTTGTGCAATGTACATCAGAGATTTTGAGACACAA |
| pAJM.474<br><br>GenBank accession number: MH101729 | CCAATTATTGAAGGCCTCCCTAACGGGGGGCCTTTTTTTGTTTCTGGTCTCCCGCTTAACGATCGTTGGCTGACCTGTAGGATCGTACAGGTTTACGCAAGAAAATGGTTTGTTA<br>CAGTCGAATAAAAGCTGTCACCGGATGTGCTTTCCGGTCTGATGAGTCCGTGAGGACGAAACAGCCTCTACAAATAATTTTGTTTAATACTAGAGAAAGAGGGGAAATACTAGAT<br>GGTGAGCAAGGGCGAGGAGCTGTTCACCGGGGTGGTGCCCATCCTGGTCGAGCTGGACGGCGACGTAAACGGCCACAAGTTCAGCGTGTCCGGCGAGGGCGAGGGCGATGCCACC<br>TACGGCAAGCTGACCCTGAAGTTCATCTGCACCACAGGCAAGCTGCCCGTGCCCTGGCCCACCCTCGTGACCACCTTCGGCTACGGCCTGCAATGCTTCGCCCGCTACCCCGACC<br>ACATGAAGCTGCACGACTTCTTCAAGTCCGCCATGCCCGAAGGCTACGTCCAGGAGCGCACCATCTTCTTCAAGGACGACGGCAACTACAAGACCCGCGCCGAGGTGAAGTTCGA<br>GGGCGACACCCTGGTGAACCGCATCGAGCTGAAGGGCATCGACTTCAAGGAGGACGGCAACATCCTGGGGCACAAGCTGGAGTACAACTACAACAGCCACAACGTCTATATCATG<br>GCCGACAAGCAGAAGAACGGCATCAAGGTGAACTTCAAGATCCGCCACAACATCGAGGACGGCAGCGTGCAGCTCGCCGACCACTACCAGCAGAACACCCCCAATCGGCGACGGCC<br>CCGTGCTGCTGCCCGACAACCACTACCTTAGCTACCAGTCCGCCCTGAGCAAAGACCCCAACGAGAAGCGCGATCACATGGTCCTGCTGGAGTTCGTGACCGCCGCCGGGATCAC<br>TCTCGGCATGGACGAGCTGTACAAGTAACTCGGTACCAAATTCCAGAAAAGAGGCCTCCCGAAAGGGGGGCCTTTTTTCGTTTTGGTCCAATGGCGGCGCGCCATCGAATGGCGC<br>AAAACCTTTCGCGGTATGGCATGATAGCGCCCGGAAGAGAGTCAATTCAGGGTGGTGAATATGAAAAACATAAATGCCGACGACACATACAGAATAATTAATAAAATTAAAGCTT<br>GTAGAAGCAATAATGATATTAATCAATGCTTATCTGATATGACTAAAATGGTACATTGTGAATATTATTTACTCGCGATCATTTATCCTCATTCTATGGTTAAATCTGATATTTC<br>AATCCTAGATAATTACCCTAAAAAATGGAGGCAATATTATGATGACGCTAATTTAATAAAATATGATCCTATAGTAGATTATTCTAACTCCAATCATTCACCAATTAATTGGAAT<br>ATATTTGAAAACAATGCTGTAAATAAAAAATCTCCAAATGTAATTAAAGAAGCGAAAACATCAGGTCTTATCACTGGGTTTAGTTTCCCTATTCATACGGCTAACAATGGCTTCG<br>GAATGCTTAGTTTTGCACATTCAGAAAAAGACAACTATATAGATAGTTTATTTTTACATGCGTGTATGAACATACCATTAATTGTTCCTTCTCTAGTTGATAATTATCGAAAAAT<br>AAATATAGCAAATAATAAATCAAACAACGATTTAACCAAAAGAGAAAAAGAATGTTTAGCGTGGGCATGCGAAGGAAAAAGCTCTTGGGATATTTCAAAAATATTAGGTTGCAGT<br>GAGCGTACTGTCACTTTCCATTTAACCAATGCGCAAATGAAACTCAATACAACAAACCGCTGCCAAAGTATTTCTAAAGCAATTTTAACAGGAGCAATTGATTGCCCATACTTTA |

AAAATTGATAAGGATCCTAATTGGTAACGAATCAGACAATTGACGGCTCGAGGGAGTAGCATAGGGTTTGCAGAATCCCTGCTTCGTCCATTTGACAGGCACATTATGCATCGAT
GATAAGCTGTCAAACATGAGCAGATCCTCTACGCCGGACGCATCGTGGCCGGCATCACCGGCGCCACAGGTGCGGTTGCTGGCGCCTATATCGCCGACATCACCGATGGGGAAGA
TCGGGCTCGCCACTTCGGGCTCATGAGCAAATATTTTATCTGAGGTGCTTCCTCGCTCACTGACTCGCTGCACGAGGCAGACCTCAGCGCTAGCGGAGTGTATACTGGCTTACTA
TGTTGGCACTGATGAGGGTGTCAGTGAAGTGCTTCATGTGGCAGGAGAAAAAAGGCTGCACCGGTGCGTCAGCAGAATATGTGATACAGGATATATTCCGCTTCCTCGCTCACTG
ACTCGCTACGCTCGGTCGTTCGACTGCGGCGAGCGGAAATGGCTTACGAACGGGGCGGAGATTTCCTGGAAGATGCCAGGAAGATACTTAACAGGGAAGTGAGAGGGCCGCGGCA
AAGCCGTTTTTCCATAGGCTCCGCCCCCCTGACAAGCATCACGAAATCTGACGCTCAAATCAGTGGTGGCGAAACCCGACAGGACTATAAAGATACCAGGCGTTTCCCCCTGGCG
GCTCCCTCGTGCGCTCTCCTGTTCCTGCCTTTCGGTTTACCGGTGTCATTCCGCTGTTATGGCCGCGTTTGTCTCATTCCACGCCTGACACTCAGTTCCGGGTAGGCAGTTCGCT
CCAAGCTGGACTGTATGCACGAACCCCCCGTTCAGTCCGACCGCTGCGCCTTATCCGGTAACTATCGTCTTGAGTCCAACCCGGAAAGACATGCAAAAGCACCACTGGCAGCAGC
CACTGGTAATTGATTTAGAGGAGTTAGTCTTGAAGTCATGCGCCGGTTAAGGCTAAACTGAAAGGACAAGTTTTGGTGACTGCGCTCCTCCAAGCCAGTTACCTCGGTTCAAAGA
GTTGGTAGCTCAGAGAACCTTCGAAAAACCGCCCTGCAAGGCGGTTTTTTCGTTTTCAGAGCAAGAGATTACGCGCAGACCAAAACGATCTCAAGAAGATCATCTTATTAAGGGG
TCTGACGCTCAGTGGAACGAAAAATCAATCTAAAGTATATATGAGTAAACTTGGTCTGACAGTTACCATGAGCCATATTCAACGGGAAACGTCTTGCTCGAGGCCGCGATTAAAT
TCCAACATGGATGCTGATTTATATGGGTATAAATGGGCTCGCGATAATGTCGGGCAATCAGGTGCGACAATCTATCGATTGTATGGGAAGCCCGATGCGCCAGAGTTGTTTCTGA
AACATGGCAAAGGTAGCGTTGCCAATGATGTTACAGATGAGATGGTCAGACTAAACTGGCTGACGGAATTTATGCCTCTTCCGACCATCAAGCATTTTATCCGTACTCCTGATGA
TGCATGGTTACTCACCACTGCGATCCCCGGGAAAACAGCATTCCAGGTATTAGAAGAATATCCTGATTCAGGTGAAAATATTGTTGATGCGCTGGCAGTGTTCCTGCGCCGGTTG
CATTCGATTCCTGTTTGTAATTGTCCTTTTAACAGCGATCGCGTATTTCGTCTCGCTCAGGCGCAATCACGAATGAATAACGGTTTGGTTGATGCGAGTGATTTTGATGACGAGC
GTAATGGCTGGCCTGTTGAACAAGTCTGGAAAGAAATGCATAAGCTTTTGCCATTCTCACCGGATTCAGTCGTCACTCATGGTGATTTCTCACTTGATAACCTTATTTTTGACGA
GGGGAAATTAATAGGTTGTATTGATGTTGGACGAGTCGGAATCGCAGACCGATACCAGGATCTTGCCATCCTATGGAACTGCCTCGGTGAGTTTTCTCCTTCATTACAGAAACGG
CTTTTTCAAAAATATGGTATTGATAATCCTGATATGAATAAATTGCAGTTTCATTTGATGCTCGATGAGTTTTTCTAAAACACCCCTTGTATTACTGTTTATGTAAGCAGACAGT
TTTATTGTTCATGATGATATATTTTTATCTTGTGCAATGTACATCAGAGATTTTGAGACACAA

pAJM.657

GenBank accession number: MH101728

CCAATTATTGAAGGCCTCCCTAACGGGGGGCCTTTTTTTGTTTCTGGTCTCCCGCTTAACGATCGTTGGCTGAACAAACAGACAATCTGGTCTGTTTGTATTATGGAAAATTTTT
CTGTATAATAGATTCAACAAACAGACAATCTGGTCTGTTTGTATTATAGCTGTCACCGGATGTGCTTTCCGGTCTGATGAGTCCGTGAGGACGAAACAGCCTCTACAAATAATTT
TGTTTAATACTAGAGAAAGAGGGGAAATACTAGATGGTGAGCAAGGGCGAGGAGCTGTTCACCGGGGTGGTGCCCATCCTGGTCGAGCTGGACGGCGACGTAAACGGCCACAAGT
TCAGCGTGTCCGGCGAGGGCGAGGGCGATGCCACCTACGGCAAGCTGACCCTGAAGTTCATCTGCACCACAGGCAAGCTGCCCGTGCCCTGGCCCACCCTCGTGACCACCTTCGG
CTACGGCCTGCAATGCTTCGCCCGCTACCCCGACCACATGAAGCTGCACGACTTCTTCAAGTCCGCCATGCCCGAAGGCTACGTCCAGGAGCGCACCATCTTCTTCAAGGACGAC
GGCAACTACAAGACCCGCGCCGAGGTGAAGTTCGAGGGCGACACCCTGGTGAACCGCATCGAGCTGAAGGGCATCGACTTCAAGGAGGACGGCAACATCCTGGGGCACAAGCTGG
AGTACAACTACAACAGCCACAACGTCTATATCATGGCCGACAAGCAGAAGAACGGCATCAAGGTGAACTTCAAGATCCGCCACAACATCGAGGACGGCAGCGTGCAGCTCGCCGA
CCACTACCAGCAGAACACCCCAATCGGCGACGGCCCCGTGCTGCTGCCCGACAACCACTACCTTAGCTACCAGTCCGCCCTGAGCAAAGACCCCAACGAGAAGCGCGATCACATG
GTCCTGCTGGAGTTCGTGACCGCCGCCGGGATCACTCTCGGCATGGACGAGCTGTACAAGTAACTCGGTACCAAATTCCAGAAAAGAGGCCTCCCGAAAGGGGGGCCTTTTTTCG
TTTTGGTCCAATGGCGGCGCGCCATCGAATGGTGCAAAACCTTTCGCGGTATGGCATGATAGCGCCCGGAAGAGAGTCAATTCAGGGTGGTGAATATGAGCCCGAAACGTCGTAC
CCAGGCAGAACGTGCAATGGAAACCCAGGGTAAACTGATTGCAGCAGCACTGGGTGTTCTGCGTGAAAAAGGTTATGCAGGTTTTCGTATTGCAGATGTTCCGGGTGCAGCCGGT
GTTAGCCGTGGTGCACAGAGCCATCATTTTCCGACCAAACTGGAACTGCTGCTGGCAACCTTTGAATGGCTGTATGAGCAGATTACCGAACGTAGCCGTGCACGTCTGGCAAAAC
TGAAACCGGAAGATGATGTTATTCAGCAGATGCTGGATGATGCAGCAGAATTTTTTCTGGATGATGATTTTAGCATCGGCCTGGATCTGATTGTTGCAGCAGATCGTGATCCGGC
ACTGCGTGAAGGTATTCAGCGTACCGTTGAACGTAATCGTTTTGTTGTTGAAGATATGTGGCTGGGTGTGCTGGTGAGCCGTGGTCTGAGCCGTGATGATGCCGAAGATATTCTG
TGGCTGATTTTTAACAGCGTTCGTGGTCTGGTAGTTCGTAGCCTGTGGCAGAAAGATAAAGAACGTTTTGAACGTGTGCGTAATAGCACCCTGGAAATTGCACGTGAACGTTATG
CAAAATTCAAACGTTGATAAGGATCCTAATTGGTAACGAATCAGACAATTGACGGCTCGAGGGAGTAGCATAGGGTTTGCAGAATCCCTGCTTCGTCCATTTGACAGGCACATTA
TGCATCGATGATAAGCTGTCAAACATGAGCAGATCCTCTACGCCGGACGCATCGTGGCCGGCATCACCGGCGCCACAGGTGCGGTTGCTGGCGCCTATATCGCCGACATCACCGA
TGGGGAAGATCGGGCTCGCCACTTCGGGCTCATGAGCAAATATTTTATCTGAGGTGCTTCCTCGCTCACTGACTCGCTGCACGAGGCAGACCTCAGCGCTAGCGGAGTGTATACT
GGCTTACTATGTTGGCACTGATGAGGGTGTCAGTGAAGTGCTTCATGTGGCAGGAGAAAAAAGGCTGCACCGGTGCGTCAGCAGAATATGTGATACAGGATATATTCCGCTTCCT
CGCTCACTGACTCGCTACGCTCGGTCGTTCGACTGCGGCGAGCGGAAATGGCTTACGAACGGGGCGGAGATTTCCTGGAAGATGCCAGGAAGATACTTAACAGGGAAGTGAGAGG
GCCGCGGCAAAGCCGTTTTTCCATAGGCTCCGCCCCCCTGACAAGCATCACGAAATCTGACGCTCAAATCAGTGGTGGCGAAACCCGACAGGACTATAAAGATACCAGGCGTTTC
CCCCTGGCGGCTCCCTCGTGCGCTCTCCTGTTCCTGCCTTTCGGTTTACCGGTGTCATTCCGCTGTTATGGCCGCGTTTGTCTCATTCCACGCCTGACACTCAGTTCCGGGTAGG
CAGTTCGCTCCAAGCTGGACTGTATGCACGAACCCCCCGTTCAGTCCGACCGCTGCGCCTTATCCGGTAACTATCGTCTTGAGTCCAACCCGGAAAGACATGCAAAAGCACCACT
GGCAGCAGCCACTGGTAATTGATTTAGAGGAGTTAGTCTTGAAGTCATGCGCCGGTTAAGGCTAAACTGAAAGGACAAGTTTTGGTGACTGCGCTCCTCCAAGCCAGTTACCTCG
GTTCAAAGAGTTGGTAGCTCAGAGAACCTTCGAAAAACCGCCCTGCAAGGCGGTTTTTTCGTTTTCAGAGCAAGAGATTACGCGCAGACCAAAACGATCTCAAGAAGATCATCTT
ATTAAGGGGTCTGACGCTCAGTGGAACGAAAAATCAATCTAAAGTATATATGAGTAAACTTGGTCTGACAGTTACCATGAGCCATATTCAACGGGAAACGTCTTGCTCGAGGCCG
CGATTAAATTCCAACATGGATGCTGATTTATATGGGTATAAATGGGCTCGCGATAATGTCGGGCAATCAGGTGCGACAATCTATCGATTGTATGGGAAGCCCGATGCGCCAGAGT
TGTTTCTGAAACATGGCAAAGGTAGCGTTGCCAATGATGTTACAGATGAGATGGTCAGACTAAACTGGCTGACGGAATTTATGCCTCTTCCGACCATCAAGCATTTTATCCGTAC
TCCTGATGATGCATGGTTACTCACCACTGCGATCCCCGGGAAAACAGCATTCCAGGTATTAGAAGAATATCCTGATTCAGGTGAAAATATTGTTGATGCGCTGGCAGTGTTCCTG
CGCCGGTTGCATTCGATTCCTGTTTGTAATTGTCCTTTTAACAGCGATCGCGTATTTCGTCTCGCTCAGGCGCAATCACGAATGAATAACGGTTTGGTTGATGCGAGTGATTTTG
ATGACGAGCGTAATGGCTGGCCTGTTGAACAAGTCTGGAAAGAAATGCATAAGCTTTTGCCATTCTCACCGGATTCAGTCGTCACTCATGGTGATTTCTCACTTGATAACCTTAT
TTTTGACGAGGGGAAATTAATAGGTTGTATTGATGTTGGACGAGTCGGAATCGCAGACCGATACCAGGATCTTGCCATCCTATGGAACTGCCTCGGTGAGTTTTCTCCTTCATTA
CAGAAACGGCTTTTTCAAAAATATGGTATTGATAATCCTGATATGAATAAATTGCAGTTTCATTTGATGCTCGATGAGTTTTTCTAAAACACCCCTTGTATTACTGTTTATGTAA
GCAGACAGTTTTATTGTTCATGATGATATATTTTTATCTTGTGCAATGTACATCAGAGATTTTGAGACACAA

pAJM.661

GenBank accession number: MH101735

CCAATTATTGAAGGCCTCCCTAACGGGGGGCCTTTTTTTGTTTCTGGTCTCCCGCTTAACGATCGTTGGCTGCACCCAGCAGTATTTACAAACAACCATGAATGTAAGTATATTC
CTTAGCAAAGCTGTCACCGGATGTGCTTTCCGGTCTGATGAGTCCGTGAGGACGAAACAGCCTCTACAAATAATTTTGTTTAATACTAGAGAAAGAGGGGAAATACTAGATGGTG
AGCAAGGGCGAGGAGCTGTTCACCGGGGTGGTGCCCATCCTGGTCGAGCTGGACGGCGACGTAAACGGCCACAAGTTCAGCGTGTCCGGCGAGGGCGAGGGCGATGCCACCTACG
GCAAGCTGACCCTGAAGTTCATCTGCACCACAGGCAAGCTGCCCGTGCCCTGGCCCACCCTCGTGACCACCTTCGGCTACGGCCTGCAATGCTTCGCCCGCTACCCCGACCACAT
GAAGCTGCACGACTTCTTCAAGTCCGCCATGCCCGAAGGCTACGTCCAGGAGCGCACCATCTTCTTCAAGGACGACGGCAACTACAAGACCCGCGCCGAGGTGAAGTTCGAGGGC
GACACCCTGGTGAACCGCATCGAGCTGAAGGGCATCGACTTCAAGGAGGACGGCAACATCCTGGGGCACAAGCTGGAGTACAACTACAACAGCCACAACGTCTATATCATGGCCG
ACAAGCAGAAGAACGGCATCAAGGTGAACTTCAAGATCCGCCACAACATCGAGGACGGCAGCGTGCAGCTCGCCGACCACTACCAGCAGAACACCCCAATCGGCGACGGCCCCGT
GCTGCTGCCCGACAACCACTACCTTAGCTACCAGTCCGCCCTGAGCAAAGACCCCAACGAGAAGCGCGATCACATGGTCCTGCTGGAGTTCGTGACCGCCGCCGGGATCACTCTC
GGCATGGACGAGCTGTACAAGTAACTCGGTACCAAATTCCAGAAAAGAGGCCTCCCGAAAGGGGGGCCTTTTTTCGTTTTGGTCCAATGGCGGCGCGCCATCGAATGGTGCAAAA
CCTTTCGCGGTATGGCATGATAGCGCCCGGAAGAGAGTCAATTCAGGGAGGTGAATATGGTTCGTCGTACCAAAGAAGAGGCACAAGAAACCCGTGCACAGATTATTGAAGCAGC
AGAACGTGCATTCTATAAACGTGGTGTTGCACGTACCACCCTGGCAGATATTGCAGAACTGGCAGGCGTTACCCGTGGTGCAATTTATTGGCATTTTAACAACAAAGCCGAACTG
GTTCAGGCACTGCTGGATAGCCTGCATGAAACCCATGATCATCTGGCACGTGCAAGCGAAAGCGAAGATGAAGTTGATCCGCTGGGTTGTATGCGTAAACTGCTGCTGCAGGTTT
TTAATGAACTGGTTCTGGATGCACGTACCCGTCGTATTAATGAAATCCTGCATCACAAATGCGAGTTCACCGATGATATGTGTGAAATTCGTCAGCAGCATCAGAGCGCAGTTCT
GGATTGTCATAAAGGTATTACCCTGACACTGGCAAATGTAGTTCGTCGTGGTCAGCTGCCTGGTGAACTGGATGCAGAACGTGCCGCAGTTGCAATGTTTGCCTATGTTGATGGT
CTGATTCGTCGTTGGCTGCTGCTGCCGGATAGCGTTGATCTGCTGGGTGATGTTGAAAAATGGGTTGATACCGGTCTGGATATGCTGCGTCTGAGTCCGGCACTGCGTAAATAAT
GATAAGGATCCTAATTGGTAACGAATCAGACAATTGACGGCTCGAGGGAGTAGCATAGGGTTTGCAGAATCCCTGCTTCGTCCATTTGACAGGCACATTATGCATCGATGATAAG
CTGTCAAACATGAGCAGATCCTCTACGCCGGACGCATCGTGGCCGGCATCACCGGCGCCACAGGTGCGGTTGCTGGCGCCTATATCGCCGACATCACCGATGGGGAAGATCGGGC
TCGCCACTTCGGGCTCATGAGCAAATATTTTATCTGAGGTGCTTCCTCGCTCACTGACTCGCTGCACGAGGCAGACCTCAGCGCTAGCGGAGTGTATACTGGCTTACTATGTTGG
CACTGATGAGGGTGTCAGTGAAGTGCTTCATGTGGCAGGAGAAAAAAGGCTGCACCGGTGCGTCAGCAGAATATGTGATACAGGATATATTCCGCTTCCTCGCTCACTGACTCGC
TACGCTCGGTCGTTCGACTGCGGCGAGCGGAAATGGCTTACGAACGGGGCGGAGATTTCCTGGAAGATGCCAGGAAGATACTTAACAGGGAAGTGAGAGGGCCGCGGCAAAGCCG
TTTTTCCATAGGCTCCGCCCCCCTGACAAGCATCACGAAATCTGACGCTCAAATCAGTGGTGGCGAAACCCGACAGGACTATAAAGATACCAGGCGTTTCCCCCTGGCGGCTCCC
TCGTGCGCTCTCCTGTTCCTGCCTTTCGGTTTACCGGTGTCATTCCGCTGTTATGGCCGCGTTTGTCTCATTCCACGCCTGACACTCAGTTCCGGGTAGGCAGTTCGCTCCAAGC
TGGACTGTATGCACGAACCCCCCGTTCAGTCCGACCGCTGCGCCTTATCCGGTAACTATCGTCTTGAGTCCAACCCGGAAAGACATGCAAAAGCACCACTGGCAGCAGCCACTGG
TAATTGATTTAGAGGAGTTAGTCTTGAAGTCATGCGCCGGTTAAGGCTAAACTGAAAGGACAAGTTTTGGTGACTGCGCTCCTCCAAGCCAGTTACCTCGGTTCAAAGAGTTGGT
AGCTCAGAGAACCTTCGAAAAACCGCCCTGCAAGGCGGTTTTTTCGTTTTCAGAGCAAGAGATTACGCGCAGACCAAAACGATCTCAAGAAGATCATCTTATTAAGGGGTCTGAC
GCTCAGTGGAACGAAAAATCAATCTAAAGTATATATGAGTAAACTTGGTCTGACAGTTACCATGAGCCATATTCAACGGGAAACGTCTTGCTCGAGGCCGCGATTAAATTCCAAC
ATGGATGCTGATTTATATGGGTATAAATGGGCTCGCGATAATGTCGGGCAATCAGGTGCGACAATCTATCGATTGTATGGGAAGCCCGATGCGCCAGAGTTGTTTCTGAAACATG
GCAAAGGTAGCGTTGCCAATGATGTTACAGATGAGATGGTCAGACTAAACTGGCTGACGGAATTTATGCCTCTTCCGACCATCAAGCATTTTATCCGTACTCCTGATGATGCATG
GTTACTCACCACTGCGATCCCCGGGAAAACAGCATTCCAGGTATTAGAAGAATATCCTGATTCAGGTGAAAATATTGTTGATGCGCTGGCAGTGTTCCTGCGCCGGTTGCATTCG
ATTCCTGTTTGTAATTGTCCTTTTAACAGCGATCGCGTATTTCGTCTCGCTCAGGCGCAATCACGAATGAATAACGGTTTGGTTGATGCGAGTGATTTTGATGACGAGCGTAATG
GCTGGCCTGTTGAACAAGTCTGGAAAGAAATGCATAAGCTTTTGCCATTCTCACCGGATTCAGTCGTCACTCATGGTGATTTCTCACTTGATAACCTTATTTTTGACGAGGGGAA
ATTAATAGGTTGTATTGATGTTGGACGAGTCGGAATCGCAGACCGATACCAGGATCTTGCCATCCTATGGAACTGCCTCGGTGAGTTTTCTCCTTCATTACAGAAACGGCTTTTT
CAAAAATATGGTATTGATAATCCTGATATGAATAAATTGCAGTTTCATTTGATGCTCGATGAGTTTTTCTAAAACACCCCTTGTATTACTGTTTATGTAAGCAGACAGTTTTATT
GTTCATGATGATATATTTTTATCTTGTGCAATGTACATCAGAGATTTTGAGACACAA

pAJM.677

GenBank

CCAATTATTGAAGGCCTCCCTAACGGGGGGCCTTTTTTTGTTTCTGGTCTCCCGCTTAACGATCGTTGGCTGAGAAACCAATTGTCCATATTGCATCAGACATTGCCGTCACTGC
GTCTTTTACTGGCTCTTCTCGCTAACCAAACCGGTAACCCCGCTTATTAAAAGCATTCTGTAACAAAGCGGGACCAAAGCCATGACAAAAACGCGTAACAAAAGTGTCTATAATC
ACGGCAGAAAAGTCCACATTGATTATTTGCACGGCGTCACACTTTGCTATGCCATAGCATTTTTATCCATAAGATTAGCGGATCCTACCTGACGCTTTTTATCGCAACTCTCTAC
TGTTTCTCCATACCCGAGCTGTCACCGGATGTGCTTTCCGGTCTGATGAGTCCGTGAGGACGAAACAGCCTCTACAAATAATTTTGTTTAATACTAGAGAAAGAGGGGAAATACT

accession number: MH101733

AGATGGTGAGCAAGGGCGAGGAGCTGTTCACCGGGGTGGTGCCCATCCTGGTCGAGCTGGACGGCGACGTAAACGGCCACAAGTTCAGCGTGTCCGGCGAGGGCGAGGGCGATGC
CACCTACGGCAAGCTGACCCTGAAGTTCATCTGCACCACAGGCAAGCTGCCCGTGCCCTGGCCCACCCTCGTGACCACCTTCGGCTACGGCCTGCAATGCTTCGCCCGCTACCCC
GACCACATGAAGCTGCACGACTTCTTCAAGTCCGCCATGCCCGAAGGCTACGTCCAGGAGCGCACCATCTTCTTCAAGGACGACGGCAACTACAAGACCCGCGCCGAGGTGAAGT
TCGAGGGCGACACCCTGGTGAACCGCATCGAGCTGAAGGGCATCGACTTCAAGGAGGACGGCAACATCCTGGGGCACAAGCTGGAGTACAACTACAACAGCCACAACGTCTATAT
CATGGCCGACAAGCAGAAGAACGGCATCAAGGTGAACTTCAAGATCCGCCACAACATCGAGGACGGCAGCGTGCAGCTCGCCGACCACTACCAGCAGAACACCCCAATCGGCGAC
GGCCCCGTGCTGCTGCCCGACAACCACTACCTTAGCTACCAGTCCGCCCTGAGCAAAGACCCCAACGAGAAGCGCGATCACATGGTCCTGCTGGAGTTCGTGACCGCCGCCGGGA
TCACTCTCGGCATGGACGAGCTGTACAAGTAACTCGGTACCAAATTCCAGAAAAGAGGCCTCCCGAAAGGGGGGCCTTTTTTCGTTTTGGTCCAATGGCGGCGCGCCATCGAATG
GTGCAAAACCTTTCGCGGTATGGCATGATAGCGCCCGGAAGAGAGTCAATTCAGGGTGGTGAATATGGCTGAAGCGCAAAATGATCCCCTGCTGCCGGGATACTCGTTTAATGCC
CATCTGGTGGCGGGTTTAACGCCGATTGAGGCCAACGGTTATCTCGATTTTTTTATCGACCGACCGCTGGGAATGAAAGGTTATATTCTCAATCTCACCATTCGCGGTCAGGGGG
TGGTGAAAAATCAGGGACGAGAATTTGTTTGCCGACCGGGTGATATTTTGCTGTTCCCGCCAGGAGAGATTCATCACTACGGCCGTCATCCGGAGGCTCGCGAATGGTATCACCA
GTGGGTTTACTTTCGTCCGCGCGCCTACTGGCATGAATGGCTTAACTGGCCGTCAATATTTGCCAATACGGGGTTCTTTCGCCCGGATGAAGCGCACCAGCCGCATTTCAGCGAC
TTTTTTGGGCAAATCATTAACGCCGGGCAAGGGGAAGGGCGCTATTCGGAGCTGCTGGCGATAAATCTGCTTGAGCAATTGTTACTGCGGCGCATGCTAGCGATTAACGGATCGC
TCCATCCACCGATGGATAATCGGGTACGCGAGGCTTGTCAGTACATCAGCGATCACCTGGCAGACAGCAATTTTGATATCGCCAGCGTCGCACAGCATGTTTGCTTGTCGCCGTC
GCGTCTGTCACATCTTTTCCGCCAGCAGTTAGGGATTAGCGTCTTAAGCTGGCGCGAGGACCAACGTATCAGCCAGGCGAAGCTGCTTTTGAGCACCACCCGGATGCCTATCGCC
ACCGTCGGTCGCAATGTTGGTTTTGACGATCAACTCTATTTCTCGCGGGTATTTAAAAAATGCACCGGGGCCAGCCCGAGCGAGTTCCGTGCCGGTTTGGAAGAAAAAGTGAATG
ATGTAGCCGTCAAGTTGTCATGATAAGATCCTATTCCAGCGGGATTAAAGAGGAGCGATTAAGCATGGTTACTATCAATACGGAATCTGCTTTAACGCCACGTTCTTTGCGGGAT
ACGCGGCGTATGAATATGTTTGTTTCGGTAGCTGCTGCGGTCGCAGGATTGTTATTTGGTCTTGATATCGGCGTAATCGCCGGAGCGTTGCCGTTCATTACCGATCACTTTGTGC
TGACCAGTCGTTTGCAGGAATGGGTGGTTAGTAGCATGATGCTCGGTGCAGCAATTGGTGCGCTGTTTAATGGTTGGCTGTCGTTCCGCCTGGGGCGTAAATACAGCCTGATGGC
GGGGGCCATCCTGTTTGTACTCGGTTCTATAGGGTCCGCTTTTGCGACCAGCGTAGAGATGTTAATCGCCGCTCGTGTGGTGCTGGGCATTGCTGTCGGGATCGCGTCTTACACC
GCTCCTCTGTATCTTTCTGAAATGGCAAGTGAAAACGTTCGCGGTAAGATGATCAGTATGTACCAGTTGATGGTCACACTCGGCATCGTGCTGGCGTTTTTATCCGATACAGCGT
TCAGTTATAGCGGTAACTGGCGCGCAATGTTGGGGGTTCTTGCTTTACCAGCAGTTCTGCTGATTATTCTGGTAGTATTTCTGCCAAATAGCCCGCGCTGGCTGGCGGAAAAGGG
GCGTCATATTGAGGCGGAAGAAGTATTGCGTATGCTGCGCGATACGTCGGAAAAAGCGCGAGAAGAACTCAACGAAATTCGTGAAAGCCTGAAGTTAAAACAGGGCGGTTGGGCA
CTGTTTAAGATCAACCGTAACGTCCGTCGTGCTGTGTTTCTCGGTATGTTGTTGCAGGCGATGCAGCAGTTTACCGGTATGAACATCATCATGTACTACGCGCCGCGTATCTTCA
AAATGGCGGGCTTTACGACCACAGAACAACAGATGATTGCGACTCTGGTCGTAGGGCTGACCTTTATGTTCGCCACCTTTATTGCGGTGTTTACGGTAGATAAAGCAGGGCGTAA
ACCGGCTCTGAAAATTGGTTTCAGCGTGATGGCGTTAGGCACTCTGGTGCTGGGCTATTGCCTGATGCAGTTTGATAACGGTACGGCTTCCAGTGGCTTGTCCTGGCTCTCTGTT
GGCATGACGATGATGTGTATTGCCGGTTATGCGATGAGCGCCGCGCCAGTGGTGTGGATCCTGTGCTCTGAAATTCAGCCGCTGAAATGCCGCGATTTCGGTATTACCTGTTCGA
CCACCACGAACTGGGTGTCGAATATGATTATCGGCGCGACCTTCCTGACACTGCTTGATAGCATTGGCGCTGCCGGTACGTTCTGGCTCTACACTGCGCTGAACATTGCGTTTGT
GGGCATTACTTTCTGGCTCATTCCGGAAACCAAAAATGTCACGCTGGAACATATCGAACGCAAACTGATGGCAGGCGAGAAGTTGAGAAATATCGGCGTCTGATAAGGATCCTAA
TTGGTAACGAATCAGACAATTGACGGCTCGAGGGAGTAGCATAGGGTTTGCAGAATCCCTGCTTCGTCCATTTGACAGGCACATTATGCATCGATGATAAGCTGTCAAACATGAG
CAGATCCTCTACGCCGGACGCATCGTGGCCGGCATCACCGGCGCCACAGGTGCGGTTGCTGGCGCCTATATCGCCGACATCACCGATGGGGAAGATCGGGCTCGCCACTTCGGGC
TCATGAGCAAATATTTTATCTGAGGTGCTTCCTCGCTCACTGACTCGCTGCACGAGGCAGACCTCAGCGCTAGCGGAGTGTATACTGGCTTACTATGTTGGCACTGATGAGGGTG
TCAGTGAAGTGCTTCATGTGGCAGGAGAAAAAAGGCTGCACCGGTGCGTCAGCAGAATATGTGATACAGGATATATTCCGCTTCCTCGCTCACTGACTCGCTACGCTCGGTCGTT
CGACTGCGGCGAGCGGAAATGGCTTACGAACGGGGCGGAGATTTCCTGGAAGATGCCAGGAAGATACTTAACAGGGAAGTGAGAGGGCCGCGGCAAAGCCGTTTTTCCATAGGCT
CCGCCCCCCTGACAAGCATCACGAAATCTGACGCTCAAATCAGTGGTGGCGAAACCCGACAGGACTATAAAGATACCAGGCGTTTCCCCCTGGCGGCTCCCTCGTGCGCTCTCCT
GTTCCTGCCTTTCGGTTTACCGGTGTCATTCCGCTGTTATGGCCGCGTTTGTCTCATTCCACGCCTGACACTCAGTTCCGGGTAGGCAGTTCGCTCCAAGCTGGACTGTATGCAC
GAACCCCCCGTTCAGTCCGACCGCTGCGCCTTATCCGGTAACTATCGTCTTGAGTCCAACCCGGAAAGACATGCAAAAGCACCACTGGCAGCAGCCACTGGTAATTGATTTAGAG
GAGTTAGTCTTGAAGTCATGCGCCGGTTAAGGCTAAACTGAAAGGACAAGTTTTGGTGACTGCGCTCCTCCAAGCCAGTTACCTCGGTTCAAAGAGTTGGTAGCTCAGAGAACCT
TCGAAAAACCGCCCTGCAAGGCGGTTTTTTCGTTTTCAGAGCAAGAGATTACGCGCAGACCAAAACGATCTCAAGAAGATCATCTTATTAAGGGGTCTGACGCTCAGTGGAACGA
AAAATCAATCTAAAGTATATATGAGTAAACTTGGTCTGACAGTTACCATGAGCCATATTCAACGGGAAACGTCTTGCTCGAGGCCGCGATTAAATTCCAACATGGATGCTGATTT
ATATGGGTATAAATGGGCTCGCGATAATGTCGGGCAATCAGGTGCGACAATCTATCGATTGTATGGGAAGCCCGATGCGCCAGAGTTGTTTCTGAAACATGGCAAAGGTAGCGTT
GCCAATGATGTTACAGATGAGATGGTCAGACTAAACTGGCTGACGGAATTTATGCCTCTTCCGACCATCAAGCATTTTATCCGTACTCCTGATGATGCATGGTTACTCACCACTG
CGATCCCCGGGAAAACAGCATTCCAGGTATTAGAAGAATATCCTGATTCAGGTGAAAATATTGTTGATGCGCTGGCAGTGTTCCTGCGCCGGTTGCATTCGATTCCTGTTTGTAA
TTGTCCTTTTAACAGCGATCGCGTATTTCGTCTCGCTCAGGCGCAATCACGAATGAATAACGGTTTGGTTGATGCGAGTGATTTTGATGACGAGCGTAATGGCTGGCCTGTTGAA
CAAGTCTGGAAAGAAATGCATAAGCTTTTGCCATTCTCACCGGATTCAGTCGTCACTCATGGTGATTTCTCACTTGATAACCTTATTTTTGACGAGGGGAAATTAATAGGTTGTA
TTGATGTTGGACGAGTCGGAATCGCAGACCGATACCAGGATCTTGCCATCCTATGGAACTGCCTCGGTGAGTTTTCTCCTTCATTACAGAAACGGCTTTTTCAAAAATATGGTAT
TGATAATCCTGATATGAATAAATTGCAGTTTCATTTGATGCTCGATGAGTTTTTCTAAAACACCCCTTGTATTACTGTTTATGTAAGCAGACAGTTTTATTGTTCATGATGATAT
ATTTTTATCTTGTGCAATGTACATCAGAGATTTTGAGACACAA

pAJM.683

GenBank accession number: MH101734

CCAATTATTGAAGGCCTCCCTAACGGGGGGCCTTTTTTTGTTTCTGGTCTCCCGCTTAACGATCGTTGGCTGAGCGCGGGTGAGAGGGATTCGTTACCAATAGACAATTGATTGG
ACGTTCAATATAATGCTAGCAGCTGTCACCGGATGTGCTTTCCGGTCTGATGAGTCCGTGAGGACGAAACAGCCTCTACAAATAATTTTGTTTAATACTAGAGAAAGAGGGGAAA
TACTAGATGGTGAGCAAGGGCGAGGAGCTGTTCACCGGGGTGGTGCCCATCCTGGTCGAGCTGGACGGCGACGTAAACGGCCACAAGTTCAGCGTGTCCGGCGAGGGCGAGGGCG
ATGCCACCTACGGCAAGCTGACCCTGAAGTTCATCTGCACCACAGGCAAGCTGCCCGTGCCCTGGCCCACCCTCGTGACCACCTTCGGCTACGGCCTGCAATGCTTCGCCCGCTA
CCCCGACCACATGAAGCTGCACGACTTCTTCAAGTCCGCCATGCCCGAAGGCTACGTCCAGGAGCGCACCATCTTCTTCAAGGACGACGGCAACTACAAGACCCGCGCCGAGGTG
AAGTTCGAGGGCGACACCCTGGTGAACCGCATCGAGCTGAAGGGCATCGACTTCAAGGAGGACGGCAACATCCTGGGGCACAAGCTGGAGTACAACTACAACAGCCACAACGTCT
ATATCATGGCCGACAAGCAGAAGAACGGCATCAAGGTGAACTTCAAGATCCGCCACAACATCGAGGACGGCAGCGTGCAGCTCGCCGACCACTACCAGCAGAACACCCCAATCGG
CGACGGCCCCGTGCTGCTGCCCGACAACCACTACCTTAGCTACCAGTCCGCCCTGAGCAAAGACCCCAACGAGAAGCGCGATCACATGGTCCTGCTGGAGTTCGTGACCGCCGCC
GGGATCACTCTCGGCATGGACGAGCTGTACAAGTAACTCGGTACCAAATTCCAGAAAAGAGGCCTCCCGAAAGGGGGGCCTTTTTTCGTTTTGGTCCAATGGCGGCGCGCCATCG
AATGGTGCAAAACCTTTCGCGGTATGGCATGATAGCGCCCGGAAGAGAGTCAATTCACGGGGGTGAATATGCCGAAACTGGGTATGCAGAGCATTCGTCGTCGTCAGCTGATTGA
TGCAACCCTGGAAGCAATTAATGAAGTTGGTATGCATGATGCAACCATTGCACAGATTGCACGTCGTGCCGGTGTTAGCACCGGTATTATTAGCCATTATTTCCGCGATAAAAAC
GGTCTACTGGAAGCAACCATGCGTGATATTACCAGCCAGCTGCGTGATGCAGTTCTGAATCGTCTGCATGCACTGCCGCAGGGTAGCGCAGAACAGCGTCTGCAGGCAATTGTTG
GTGGTAATTTTGATGAAACCCAGGTTAGCAGCGCAGCAATGAAAGCATGGCTGGCATTTTGGGCAATCAGCATGCATCAGCCGATGCTGTATCGTCTGCAGCAGGTTAGCAGTCG
TCGTCTGCTGAGCAATCTGGTTAGCGAATTTCGTCGTGAACTGCCTCGTGAACAGGCACAAGAGGCAGGTTATGGTCTGGCAGCACTGATTGATGGTCTGTGGCTGCGTGCAGCA
CTGAGCGGTAAACCGCTGGATAAAACCCGTGCAAATAGCCTGACCCGTCATTTTATCACCCAGCATCTGCCGACCGATTGATAAGGATCCTAATTGGTAACGAATCAGACAATTG
ACGGCTCGAGGGAGTAGCATAGGGTTTGCAGAATCCCTGCTTCGTCCATTTGACAGGCACATTATGCATCGATGATAAGCTGTCAAACATGAGCAGATCCTCTACGCCGGACGCA
TCGTGGCCGGCATCACCGGCGCCACAGGTGCGGTTGCTGGCGCCTATATCGCCGACATCACCGATGGGGAAGATCGGGCTCGCCACTTCGGGCTCATGAGCAAATATTTTATCTG
AGGTGCTTCCTCGCTCACTGACTCGCTGCACGAGGCAGACCTCAGCGCTAGCGGAGTGTATACTGGCTTACTATGTTGGCACTGATGAGGGTGTCAGTGAAGTGCTTCATGTGGC
AGGAGAAAAAAGGCTGCACCGGTGCGTCAGCAGAATATGTGATACAGGATATATTCCGCTTCCTCGCTCACTGACTCGCTACGCTCGGTCGTTCGACTGCGGCGAGCGGAAATGG
CTTACGAACGGGGCGGAGATTTCCTGGAAGATGCCAGGAAGATACTTAACAGGGAAGTGAGAGGGCCGCGGCAAAGCCGTTTTTCCATAGGCTCCGCCCCCCTGACAAGCATCAC
GAAATCTGACGCTCAAATCAGTGGTGGCGAAACCCGACAGGACTATAAAGATACCAGGCGTTTCCCCCTGGCGGCTCCCTCGTGCGCTCTCCTGTTCCTGCCTTTCGGTTTACCG
GTGTCATTCCGCTGTTATGGCCGCGTTTGTCTCATTCCACGCCTGACACTCAGTTCCGGGTAGGCAGTTCGCTCCAAGCTGGACTGTATGCACGAACCCCCCGTTCAGTCCGACC
GCTGCGCCTTATCCGGTAACTATCGTCTTGAGTCCAACCCGGAAAGACATGCAAAAGCACCACTGGCAGCAGCCACTGGTAATTGATTTAGAGGAGTTAGTCTTGAAGTCATGCG
CCGGTTAAGGCTAAACTGAAAGGACAAGTTTTGGTGACTGCGCTCCTCCAAGCCAGTTACCTCGGTTCAAAGAGTTGGTAGCTCAGAGAACCTTCGAAAAACCGCCCTGCAAGGC
GGTTTTTTCGTTTTCAGAGCAAGAGATTACGCGCAGACCAAAACGATCTCAAGAAGATCATCTTATTAAGGGGTCTGACGCTCAGTGGAACGAAAAATCAATCTAAAGTATATAT
GAGTAAACTTGGTCTGACAGTTACCATGAGCCATATTCAACGGGAAACGTCTTGCTCGAGGCCGCGATTAAATTCCAACATGGATGCTGATTTATATGGGTATAAATGGGCTCGC
GATAATGTCGGGCAATCAGGTGCGACAATCTATCGATTGTATGGGAAGCCCGATGCGCCAGAGTTGTTTCTGAAACATGGCAAAGGTAGCGTTGCCAATGATGTTACAGATGAGA
TGGTCAGACTAAACTGGCTGACGGAATTTATGCCTCTTCCGACCATCAAGCATTTTATCCGTACTCCTGATGATGCATGGTTACTCACCACTGCGATCCCCGGGAAAACAGCATT
CCAGGTATTAGAAGAATATCCTGATTCAGGTGAAAATATTGTTGATGCGCTGGCAGTGTTCCTGCGCCGGTTGCATTCGATTCCTGTTTGTAATTGTCCTTTTAACAGCGATCGC
GTATTTCGTCTCGCTCAGGCGCAATCACGAATGAATAACGGTTTGGTTGATGCGAGTGATTTTGATGACGAGCGTAATGGCTGGCCTGTTGAACAAGTCTGGAAAGAAATGCATA
AGCTTTTGCCATTCTCACCGGATTCAGTCGTCACTCATGGTGATTTCTCACTTGATAACCTTATTTTTGACGAGGGGAAATTAATAGGTTGTATTGATGTTGGACGAGTCGGAAT
CGCAGACCGATACCAGGATCTTGCCATCCTATGGAACTGCCTCGGTGAGTTTTCTCCTTCATTACAGAAACGGCTTTTTCAAAAATATGGTATTGATAATCCTGATATGAATAAA
TTGCAGTTTCATTTGATGCTCGATGAGTTTTTCTAAAACACCCCTTGTATTACTGTTTATGTAAGCAGACAGTTTTATTGTTCATGATGATATATTTTTATCTTGTGCAATGTAC
ATCAGAGATTTTGAGACACAA

pAJM.690

GenBank accession number: MH101736

CCAATTATTGAAGGCCTCCCTAACGGGGGGCCTTTTTTTGTTTCTGGTCTCCCGCTTAACGATCGTTGGCTGTTTTGTTCGATTATCGAACAAATTATTGAAATATCGAACAAAA
CCTCTAAACTACTGTGGCACTGAATCAAAAAATTATAAACCATGATCAGAAGCTGTCACCGGATGTGCTTTCCGGTCTGATGAGTCCGTGAGGACGAAACAGCCTCTACAAATAA
TTTTGTTTAATACTAGAGAAAGAGGGGAAATACTAGATGGTGAGCAAGGGCGAGGAGCTGTTCACCGGGGTGGTGCCCATCCTGGTCGAGCTGGACGGCGACGTAAACGGCCACA
AGTTCAGCGTGTCCGGCGAGGGCGAGGGCGATGCCACCTACGGCAAGCTGACCCTGAAGTTCATCTGCACCACAGGCAAGCTGCCCGTGCCCTGGCCCACCCTCGTGACCACCTT
CGGCTACGGCCTGCAATGCTTCGCCCGCTACCCCGACCACATGAAGCTGCACGACTTCTTCAAGTCCGCCATGCCCGAAGGCTACGTCCAGGAGCGCACCATCTTCTTCAAGGAC
GACGGCAACTACAAGACCCGCGCCGAGGTGAAGTTCGAGGGCGACACCCTGGTGAACCGCATCGAGCTGAAGGGCATCGACTTCAAGGAGGACGGCAACATCCTGGGGCACAAGC
TGGAGTACAACTACAACAGCCACAACGTCTATATCATGGCCGACAAGCAGAAGAACGGCATCAAGGTGAACTTCAAGATCCGCCACAACATCGAGGACGGCAGCGTGCAGCTCGC
CGACCACTACCAGCAGAACACCCCAATCGGCGACGGCCCCGTGCTGCTGCCCGACAACCACTACCTTAGCTACCAGTCCGCCCTGAGCAAAGACCCCAACGAGAAGCGCGATCAC
ATGGTCCTGCTGGAGTTCGTGACCGCCGCCGGGATCACTCTCGGCATGGACGAGCTGTACAAGTAACTCGGTACCAAATTCCAGAAAAGAGGCCTCCCGAAAGGGGGGCCTTTTT
TCGTTTTGGTCCAATGGCGGCGCGCCATCGAATGGTGCAAAACCTTTCGCGGTATGGCATGATAGCGCCCGGAAGAGAGTCAATTCAGGGAGGTGAATATGTGGTCGAACATGGA
TGACAAGAAAGTGAAAGAGGAGAATATTCTGCACAATTCCACCAACAAGAAGATCATCCGCCACGAAGATTTTGTAGCCGGCATTAGCAAAGGGATGGCGATTCTGGATTCGTTT

GGTACAGATCGTCATCGCCTCAATATCACCATGGCCGCAGAGAAAACCGGTATGACACGTGCAGCAGCTCGTCGCCACCTGCTTACTCTGGAGTATCTGGGCTATCTGGAAAGTG
ACGGCCACTACTTCTACTTAACTCCCAAAATCCTGAAATTCAGTGGTTCATATTTGGGTGGTGCTCAATTGCCGAAAATTTCCCAACCACTGTTGAACTTGCTTACGACCCAGAC
CAGCCTGATTTACAGCGTGATGGTGTTGGATGGCTATGAAGCCATTACCATTGCGCGTTCTGCCGCTCATCAGCAAACCGACCGCGTTAACCCGTATGGTTTACATCTCGGGAAT
CGCTTACCAGCGCATACAACGTCAGCGGGCAAAATCCTGTTAGCGTATTTGGATGACCATGCCCAGCAAGAGTGGCTCAATCAGTACCCTCTGCAACGGCTCACGAAATACACGT
ATACCAACCACATCGACTTTCTGCGCCTTTTGAGTGAAATCAAGGAACAGGGTTGGTGCTATAGTTCGGAAGAACACGAACTGGGAGTACACGCCCTTGCGGTTCCGATTTACGG
ACAGCAGTCTCGCGTCGTAGCGGCACTGAACATTGTCAGCCCGACAATGCGGACCACGAAAGAATACCTGATTCAGCATATTCTGCCGTTACTGCAAGAAACTGCGCGTGAATTG
CGCAATATCCTGTAATGATAAGGATCCTAATTGGTAACGAATCAGACAATTGACGGCTCGAGGGAGTAGCATAGGGTTTGCAGAATCCCTGCTTCGTCCATTTGACAGGCACATT
ATGCATCGATGATAAGCTGTCAAACATGAGCAGATCCTCTACGCCGGACGCATCGTGGCCGGCATCACCGGCGCCACAGGTGCGGTTGCTGGCGCCTATATCGCCGACATCACCG
ATGGGGAAGATCGGGCTCGCCACTTCGGGCTCATGAGCAAATATTTTATCTGAGGTGCTTCCTCGCTCACTGACTCGCTGCACGAGGCAGACCTCAGCGCTAGCGGAGTGTATAC
TGGCTTACTATGTTGGCACTGATGAGGGTGTCAGTGAAGTGCTTCATGTGGCAGGAGAAAAAAGGCTGCACCGGTGCGTCAGCAGAATATGTGATACAGGATATATTCCGCTTCC
TCGCTCACTGACTCGCTACGCTCGGTCGTTCGACTGCGGCGAGCGGAAATGGCTTACGAACGGGGCGGAGATTTCCTGGAAGATGCCAGGAAGATACTTAACAGGGAAGTGAGAG
GGCCGCGGCAAAGCCGTTTTTCCATAGGCTCCGCCCCCCTGACAAGCATCACGAAATCTGACGCTCAAATCAGTGGTGGCGAAACCCGACAGGACTATAAAGATACCAGGCGTTT
CCCCCTGGCGGCTCCCTCGTGCGCTCTCCTGTTCCTGCCTTTCGGTTTACCGGTGTCATTCCGCTGTTATGGCCGCGTTTGTCTCATTCCACGCCTGACACTCAGTTCCGGGTAG
GCAGTTCGCTCCAAGCTGGACTGTATGCACGAACCCCCCGTTCAGTCCGACCGCTGCGCCTTATCCGGTAACTATCGTCTTGAGTCCAACCCGGAAAGACATGCAAAAGCACCAC
TGGCAGCAGCCACTGGTAATTGATTTAGAGGAGTTAGTCTTGAAGTCATGCGCCGGTTAAGGCTAAACTGAAAGGACAAGTTTTGGTGACTGCGCTCCTCCAAGCCAGTTACCTC
GGTTCAAAGAGTTGGTAGCTCAGAGAACCTTCGAAAAACCGCCCTGCAAGGCGGTTTTTTCGTTTTCAGAGCAAGAGATTACGCGCAGACCAAAACGATCTCAAGAAGATCATCT
TATTAAGGGGTCTGACGCTCAGTGGAACGAAAAATCAATCTAAAGTATATATGAGTAAACTTGGTCTGACAGTTACCATGAGCCATATTCAACGGGAAACGTCTTGCTCGAGGCC
GCGATTAAATTCCAACATGGATGCTGATTTATATGGGTATAAATGGGCTCGCGATAATGTCGGGCAATCAGGTGCGACAATCTATCGATTGTATGGGAAGCCCGATGCGCCAGAG
TTGTTTCTGAAACATGGCAAAGGTAGCGTTGCCAATGATGTTACAGATGAGATGGTCAGACTAAACTGGCTGACGGAATTTATGCCTCTTCCGACCATCAAGCATTTTATCCGTA
CTCCTGATGATGCATGGTTACTCACCACTGCGATCCCCGGGAAAACAGCATTCCAGGTATTAGAAGAATATCCTGATTCAGGTGAAAATATTGTTGATGCGCTGGCAGTGTTCCT
GCGCCGGTTGCATTCGATTCCTGTTTGTAATTGTCCTTTTAACAGCGATCGCGTATTTCGTCTCGCTCAGGCGCAATCACGAATGAATAACGGTTTGGTTGATGCGAGTGATTTT
GATGACGAGCGTAATGGCTGGCCTGTTGAACAAGTCTGGAAAGAAATGCATAAGCTTTTGCCATTCTCACCGGATTCAGTCGTCACTCATGGTGATTTCTCACTTGATAACCTTA
TTTTTGACGAGGGGAAATTAATAGGTTGTATTGATGTTGGACGAGTCGGAATCGCAGACCGATACCAGGATCTTGCCATCCTATGGAACTGCCTCGGTGAGTTTTCTCCTTCATT
ACAGAAACGGCTTTTTCAAAAATATGGTATTGATAATCCTGATATGAATAAATTGCAGTTTCATTTGATGCTCGATGAGTTTTTCTAAAACACCCCTTGTATTACTGTTTATGTA
AGCAGACAGTTTTATTGTTCATGATGATATATTTTTATCTTGTGCAATGTACATCAGAGATTTTGAGACACAA

pAJM.711

GenBank accession number: MH101715

CCAATTATTGAAGGCCTCCCTAACGGGGGGCCTTTTTTTGTTTCTGGTCTCCCGCTTAACGATCGTTGGCTGCGACGTACGGTGGAATCTGATTCGTTACCAATTGACATGATAC
GAAACGTACCGTATCGTTAAGGTAGCTGTCACCGGATGTGCTTTCCGGTCTGATGAGTCCGTGAGGACGAAACAGCCTCTACAAATAATTTTGTTTAATACTAGAGAAAGAGGGG
AAATACTAGATGGTGAGCAAGGGCGAGGAGCTGTTCACCGGGGTGGTGCCCATCCTGGTCGAGCTGGACGGCGACGTAAACGGCCACAAGTTCAGCGTGTCCGGCGAGGGCGAGG
GCGATGCCACCTACGGCAAGCTGACCCTGAAGTTCATCTGCACCACAGGCAAGCTGCCCGTGCCCTGGCCCACCCTCGTGACCACCTTCGGCTACGGCCTGCAATGCTTCGCCCG
CTACCCCGACCACATGAAGCTGCACGACTTCTTCAAGTCCGCCATGCCCGAAGGCTACGTCCAGGAGCGCACCATCTTCTTCAAGGACGACGGCAACTACAAGACCCGCGCCGAG
GTGAAGTTCGAGGGCGACACCCTGGTGAACCGCATCGAGCTGAAGGGCATCGACTTCAAGGAGGACGGCAACATCCTGGGGCACAAGCTGGAGTACAACTACAACAGCCACAACG
TCTATATCATGGCCGACAAGCAGAAGAACGGCATCAAGGTGAACTTCAAGATCCGCCACAACATCGAGGACGGCAGCGTGCAGCTCGCCGACCACTACCAGCAGAACACCCCCAAT
CGGCGACGGCCCCGTGCTGCTGCCCGACAACCACTACCTTAGCTACCAGTCCGCCCTGAGCAAAGACCCCAACGAGAAGCGCGATCACATGGTCCTGCTGGAGTTCGTGACCGCC
GCCGGGATCACTCTCGGCATGGACGAGCTGTACAAGTAACTCGGTACCAAATTCCAGAAAAGAGGCCTCCCGAAAGGGGGGCCTTTTTTCGTTTTGGTCCAATGGCGGCGCGCCA
TCGAATGAGGTGCTTCCTCGCTCACTGACTCGCTGCACGAGGCAGACCTCAGCGCTAGCGGAGTGTATACTGGCTTACTATGTTGGCACTGATGAGGGTGTCAGTGAAGTGCTTC
ATGTGGCAGGAGAAAAAAGGCTGCACCGGTGCGTCAGCAGAATATGTGATACAGGATATATTCCGCTTCCTCGCTCACTGACTCGCTACGCTCGGTCGTTCGACTGCGGCGAGCG
GAAATGGCTTACGAACGGGGCGGAGATTTCCTGGAAGATGCCAGGAAGATACTTAACAGGGAAGTGAGAGGGCCGCGGCAAAGCCGTTTTTCCATAGGCTCCGCCCCCCTGACAA
GCATCACGAAATCTGACGCTCAAATCAGTGGTGGCGAAACCCGACAGGACTATAAAGATACCAGGCGTTTCCCCCTGGCGGCTCCCTCGTGCGCTCTCCTGTTCCTGCCTTTCGG
TTTACCGGTGTCATTCCGCTGTTATGGCCGCGTTTGTCTCATTCCACGCCTGACACTCAGTTCCGGGTAGGCAGTTCGCTCCAAGCTGGACTGTATGCACGAACCCCCCGTTCAG
TCCGACCGCTGCGCCTTATCCGGTAACTATCGTCTTGAGTCCAACCCGGAAAGACATGCAAAAGCACCACTGGCAGCAGCCACTGGTAATTGATTTAGAGGAGTTAGTCTTGAAG
TCATGCGCCGGTTAAGGCTAAACTGAAAGGACAAGTTTTGGTGACTGCGCTCCTCCAAGCCAGTTACCTCGGTTCAAAGAGTTGGTAGCTCAGAGAACCTTCGAAAAACCGCCCT
GCAAGGCGGTTTTTTCGTTTTCAGAGCAAGAGATTACGCGCAGACCAAAACGATCTCAAGAAGATCATCTTATTAAGGGGTCTGACGCTCAGTGGAACGAAAAATCAATCTAAAG
TATATATGAGTAAACTTGGTCTGACAGTTACCATGAGCCATATTCAACGGGAAACGTCTTGCTCGAGGCCGCGATTAAATTCCAACATGGATGCTGATTTATATGGGTATAAATG
GGCTCGCGATAATGTCGGGCAATCAGGTGCGACAATCTATCGATTGTATGGGAAGCCCGATGCGCCAGAGTTGTTTCTGAAACATGGCAAAGGTAGCGTTGCCAATGATGTTACA
GATGAGATGGTCAGACTAAACTGGCTGACGGAATTTATGCCTCTTCCGACCATCAAGCATTTTATCCGTACTCCTGATGATGCATGGTTACTCACCACTGCGATCCCCGGGAAAA
CAGCATTCCAGGTATTAGAAGAATATCCTGATTCAGGTGAAAATATTGTTGATGCGCTGGCAGTGTTCCTGCGCCGGTTGCATTCGATTCCTGTTTGTAATTGTCCTTTTAACAG
CGATCGCGTATTTCGTCTCGCTCAGGCGCAATCACGAATGAATAACGGTTTGGTTGATGCGAGTGATTTTGATGACGAGCGTAATGGCTGGCCTGTTGAACAAGTCTGGAAAGAA
ATGCATAAGCTTTTGCCATTCTCACCGGATTCAGTCGTCACTCATGGTGATTTCTCACTTGATAACCTTATTTTTGACGAGGGGAAATTAATAGGTTGTATTGATGTTGGACGAG
TCGGAATCGCAGACCGATACCAGGATCTTGCCATCCTATGGAACTGCCTCGGTGAGTTTTCTCCTTCATTACAGAAACGGCTTTTTCAAAAATATGGTATTGATAATCCTGATAT
GAATAAATTGCAGTTTCATTTGATGCTCGATGAGTTTTTCTAAAACACCCCTTGTATTACTGTTTATGTAAGCAGACAGTTTTATTGTTCATGATGATATATTTTTATCTTGTGC
AATGTACATCAGAGATTTTGAGACACAA

pAJM.712

GenBank accession number: MH101716

CCAATTATTGAAGGCCTCCCTAACGGGGGGCCTTTTTTTGTTTCTGGTCTCCCGCTTAACGATCGTTGGCTGAACAAACAGACAATCTGGTCTGTTTGTATTATGGAAAATTTTT
CTGTATAATAGATTCAACAAACAGACAATCTGGTCTGTTTGTATTATAGCTGTCACCGGATGTGCTTTCCGGTCTGATGAGTCCGTGAGGACGAAACAGCCTCTACAAATAATTT
TGTTTAATACTAGAGAAAGAGGGGAAATACTAGATGGTGAGCAAGGGCGAGGAGCTGTTCACCGGGGTGGTGCCCATCCTGGTCGAGCTGGACGGCGACGTAAACGGCCACAAGT
TCAGCGTGTCCGGCGAGGGCGAGGGCGATGCCACCTACGGCAAGCTGACCCTGAAGTTCATCTGCACCACAGGCAAGCTGCCCGTGCCCTGGCCCACCCTCGTGACCACCTTCGG
CTACGGCCTGCAATGCTTCGCCCGCTACCCCGACCACATGAAGCTGCACGACTTCTTCAAGTCCGCCATGCCCGAAGGCTACGTCCAGGAGCGCACCATCTTCTTCAAGGACGAC
GGCAACTACAAGACCCGCGCCGAGGTGAAGTTCGAGGGCGACACCCTGGTGAACCGCATCGAGCTGAAGGGCATCGACTTCAAGGAGGACGGCAACATCCTGGGGCACAAGCTGG
AGTACAACTACAACAGCCACAACGTCTATATCATGGCCGACAAGCAGAAGAACGGCATCAAGGTGAACTTCAAGATCCGCCACAACATCGAGGACGGCAGCGTGCAGCTCGCCGA
CCACTACCAGCAGAACACCCCCAATCGGCGACGGCCCCGTGCTGCTGCCCGACAACCACTACCTTAGCTACCAGTCCGCCCTGAGCAAAGACCCCAACGAGAAGCGCGATCACATG
GTCCTGCTGGAGTTCGTGACCGCCGCCGGGATCACTCTCGGCATGGACGAGCTGTACAAGTAACTCGGTACCAAATTCCAGAAAAGAGGCCTCCCGAAAGGGGGGCCTTTTTTCG
TTTTGGTCCAATGGCGGCGCGCCATCGAATGAGGTGCTTCCTCGCTCACTGACTCGCTGCACGAGGCAGACCTCAGCGCTAGCGGAGTGTATACTGGCTTACTATGTTGGCACTG
ATGAGGGTGTCAGTGAAGTGCTTCATGTGGCAGGAGAAAAAAGGCTGCACCGGTGCGTCAGCAGAATATGTGATACAGGATATATTCCGCTTCCTCGCTCACTGACTCGCTACGC
TCGGTCGTTCGACTGCGGCGAGCGGAAATGGCTTACGAACGGGGCGGAGATTTCCTGGAAGATGCCAGGAAGATACTTAACAGGGAAGTGAGAGGGCCGCGGCAAAGCCGTTTTT
CCATAGGCTCCGCCCCCCTGACAAGCATCACGAAATCTGACGCTCAAATCAGTGGTGGCGAAACCCGACAGGACTATAAAGATACCAGGCGTTTCCCCCTGGCGGCTCCCTCGTG
CGCTCTCCTGTTCCTGCCTTTCGGTTTACCGGTGTCATTCCGCTGTTATGGCCGCGTTTGTCTCATTCCACGCCTGACACTCAGTTCCGGGTAGGCAGTTCGCTCCAAGCTGGAC
TGTATGCACGAACCCCCCGTTCAGTCCGACCGCTGCGCCTTATCCGGTAACTATCGTCTTGAGTCCAACCCGGAAAGACATGCAAAAGCACCACTGGCAGCAGCCACTGGTAATT
GATTTAGAGGAGTTAGTCTTGAAGTCATGCGCCGGTTAAGGCTAAACTGAAAGGACAAGTTTTGGTGACTGCGCTCCTCCAAGCCAGTTACCTCGGTTCAAAGAGTTGGTAGCTC
AGAGAACCTTCGAAAAACCGCCCCGCAAGGCGGTTTTTTCGTTTTCAGAGCAAGAGATTACGCGCAGACCAAAACGATCTCAAGAAGATCATCTTATTAAGGGGTCTGACGCTCA
GTGGAACGAAAAATCAATCTAAAGTATATATGAGTAAACTTGGTCTGACAGTTACCATGAGCCATATTCAACGGGAAACGTCTTGCTCGAGGCCGCGATTAAATTCCAACATGGA
TGCTGATTTATATGGGTATAAATGGGCTCGCGATAATGTCGGGCAATCAGGTGCGACAATCTATCGATTGTATGGGAAGCCCGATGCGCCAGAGTTGTTTCTGAAACATGGCAAA
GGTAGCGTTGCCAATGATGTTACAGATGAGATGGTCAGACTAAACTGGCTGACGGAATTTATGCCTCTTCCGACCATCAAGCATTTTATCCGTACTCCTGATGATGCATGGTTAC
TCACCACTGCGATCCCCGGGAAAACAGCATTCCAGGTATTAGAAGAATATCCTGATTCAGGTGAAAATATTGTTGATGCGCTGGCAGTGTTCCTGCGCCGGTTGCATTCGATTCC
TGTTTGTAATTGTCCTTTTAACAGCGATCGCGTATTTCGTCTCGCTCAGGCGCAATCACGAATGAATAACGGTTTGGTTGATGCGAGTGATTTTGATGACGAGCGTAATGGCTGG
CCTGTTGAACAAGTCTGGAAAGAAATGCATAAGCTTTTGCCATTCTCACCGGATTCAGTCGTCACTCATGGTGATTTCTCACTTGATAACCTTATTTTTGACGAGGGGAAATTAA
TAGGTTGTATTGATGTTGGACGAGTCGGAATCGCAGACCGATACCAGGATCTTGCCATCCTATGGAACTGCCTCGGTGAGTTTTCTCCTTCATTACAGAAACGGCTTTTTCAAAA
ATATGGTATTGATAATCCTGATATGAATAAATTGCAGTTTCATTTGATGCTCGATGAGTTTTTCTAAAACACCCCTTGTATTACTGTTTATGTAAGCAGACAGTTTTATTGTTCA
TGATGATATATTTTTATCTTGTGCAATGTACATCAGAGATTTTGAGACACAA

pAJM.713

GenBank accession number: MH101717

CCAATTATTGAAGGCCTCCCTAACGGGGGGCCTTTTTTTGTTTCTGGTCTCCCGCTTAACGATCGTTGGCTGACCTGTAGGATCGTACAGGTTTACGCAAGAAAATGGTTTGTTA
CAGTCGAATAAAAGCTGTCACCGGATGTGCTTTCCGGTCTGATGAGTCCGTGAGGACGAAACAGCCTCTACAAATAATTTTGTTTAATACTAGAGAAAGAGGGGAAATACTAGAT
GGTGAGCAAGGGCGAGGAGCTGTTCACCGGGGTGGTGCCCATCCTGGTCGAGCTGGACGGCGACGTAAACGGCCACAAGTTCAGCGTGTCCGGCGAGGGCGAGGGCGATGCCACC
TACGGCAAGCTGACCCTGAAGTTCATCTGCACCACAGGCAAGCTGCCCGTGCCCTGGCCCACCCTCGTGACCACCTTCGGCTACGGCCTGCAATGCTTCGCCCGCTACCCCGACC
ACATGAAGCTGCACGACTTCTTCAAGTCCGCCATGCCCGAAGGCTACGTCCAGGAGCGCACCATCTTCTTCAAGGACGACGGCAACTACAAGACCCGCGCCGAGGTGAAGTTCGA
GGGCGACACCCTGGTGAACCGCATCGAGCTGAAGGGCATCGACTTCAAGGAGGACGGCAACATCCTGGGGCACAAGCTGGAGTACAACTACAACAGCCACAACGTCTATATCATG
GCCGACAAGCAGAAGAACGGCATCAAGGTGAACTTCAAGATCCGCCACAACATCGAGGACGGCAGCGTGCAGCTCGCCGACCACTACCAGCAGAACACCCCCAATCGGCGACGGCC
CCGTGCTGCTGCCCGACAACCACTACCTTAGCTACCAGTCCGCCCTGAGCAAAGACCCCAACGAGAAGCGCGATCACATGGTCCTGCTGGAGTTCGTGACCGCCGCCGGGATCAC
TCTCGGCATGGACGAGCTGTACAAGTAACTCGGTACCAAATTCCAGAAAAGAGGCCTCCCGAAAGGGGGGCCTTTTTTCGTTTTGGTCCAATGGCGGCGCGCCATCGAATGAGGT
GCTTCCTCGCTCACTGACTCGCTGCACGAGGCAGACCTCAGCGCTAGCGGAGTGTATACTGGCTTACTATGTTGGCACTGATGAGGGTGTCAGTGAAGTGCTTCATGTGGCAGGA
GAAAAAAGGCTGCACCGGTGCGTCAGCAGAATATGTGATACAGGATATATTCCGCTTCCTCGCTCACTGACTCGCTACGCTCGGTCGTTCGACTGCGGCGAGCGGAAATGGCTTA
CGAACGGGGCGGAGATTTCCTGGAAGATGCCAGGAAGATACTTAACAGGGAAGTGAGAGGGCCGCGGCAAAGCCGTTTTTCCATAGGCTCCGCCCCCCTGACAAGCATCACGAAA
TCTGACGCTCAAATCAGTGGTGGCGAAACCCGACAGGACTATAAAGATACCAGGCGTTTCCCCCTGGCGGCTCCCTCGTGCGCTCTCCTGTTCCTGCCTTTCGGTTTACCGGTGT
CATTCCGCTGTTATGGCCGCGTTTGTCTCATTCCACGCCTGACACTCAGTTCCGGGTAGGCAGTTCGCTCCAAGCTGGACTGTATGCACGAACCCCCCGTTCAGTCCGACCGCTG

CGCCTTATCCGGTAACTATCGTCTTGAGTCCAACCCGGAAAGACATGCAAAAGCACCACTGGCAGCAGCCACTGGTAATTGATTTAGAGGAGTTAGTCTTGAAGTCATGCGCCGG
TTAAGGCTAAACTGAAAGGACAAGTTTTGGTGACTGCGCTCCTCCAAGCCAGTTACCTCGGTTCAAAGAGTTGGTAGCTCAGAGAACCTTCGAAAAACCGCCCTGCAAGGCGGTT
TTTTCGTTTTCAGAGCAAGAGATTACGCGCAGACCAAAACGATCTCAAGAAGATCATCTTATTAAGGGGTCTGACGCTCAGTGGAACGAAAAATCAATCTAAAGTATATATGAGT
AAACTTGGTCTGACAGTTACCATGAGCCATATTCAACGGGAAACGTCTTGCTCGAGGCCGCGATTAAATTCCAACATGGATGCTGATTTATATGGGTATAAATGGGCTCGCGATA
ATGTCGGGCAATCAGGTGCGACAATCTATCGATTGTATGGGAAGCCCGATGCGCCAGAGTTGTTTCTGAAACATGGCAAAGGTAGCGTTGCCAATGATGTTACAGATGAGATGGT
CAGACTAAACTGGCTGACGGAATTTATGCCTCTTCCGACCATCAAGCATTTTATCCGTACTCCTGATGATGCATGGTTACTCACCACTGCGATCCCCGGGAAAACAGCATTCCAG
GTATTAGAAGAATATCCTGATTCAGGTGAAAATATTGTTGATGCGCTGGCAGTGTTCCTGCGCCGGTTGCATTCGATTCCTGTTTGTAATTGTCCTTTTAACAGCGATCGCGTAT
TTCGTCTCGCTCAGGCGCAATCACGAATGAATAACGGTTTGGTTGATGCGAGTGATTTTGATGACGAGCGTAATGGCTGGCCTGTTGAACAAGTCTGGAAAGAAATGCATAAGCT
TTTGCCATTCTCACCGGATTCAGTCGTCACTCATGGTGATTTCTCACTTGATAACCTTATTTTTGACGAGGGGAAATTAATAGGTTGTATTGATGTTGGACGAGTCGGAATCGCA
GACCGATACCAGGATCTTGCCATCCTATGGAACTGCCTCGGTGAGTTTTCTCCTTCATTACAGAAACGGCTTTTTCAAAAATATGGTATTGATAATCCTGATATGAATAAATTGC
AGTTTCATTTGATGCTCGATGAGTTTTTCTAAAACACCCCTTGTATTACTGTTTATGTAAGCAGACAGTTTTATTGTTCATGATGATATATTTTTATCTTGTGCAATGTACATCA
GAGATTTTGAGACACAA

pAJM.714

GenBank accession number: MH101718

CCAATTATTGAAGGCCTCCCTAACGGGGGGCCTTTTTTTGTTTCTGGTCTCCCGCTTAACGATCGTTGGCTGATTGGATCCAATTGACAGCTAGCTCAGTCCTAGGTACCATTGG
ATCCAATAGCTGTCACCGGATGTGCTTTCCGGTCTGATGAGTCCGTGAGGACGAAACAGCCTCTACAAATAATTTTGTTTAATACTAGAGAAAGAGGGGAAATACTAGATGGTGA
GCAAGGGCGAGGAGCTGTTCACCGGGGTGGTGCCCATCCTGGTCGAGCTGGACGGCGACGTAAACGGCCACAAGTTCAGCGTGTCCGGCGAGGGCGAGGGCGATGCCACCTACGG
CAAGCTGACCCTGAAGTTCATCTGCACCACAGGCAAGCTGCCCGTGCCCTGGCCCACCCTCGTGACCACCTTCGGCTACGGCCTGCAATGCTTCGCCCGCTACCCCGACCACATG
AAGCTGCACGACTTCTTCAAGTCCGCCATGCCCGAAGGCTACGTCCAGGAGCGCACCATCTTCTTCAAGGACGACGGCAACTACAAGACCCGCGCCGAGGTGAAGTTCGAGGGCG
ACACCCTGGTGAACCGCATCGAGCTGAAGGGCATCGACTTCAAGGAGGACGGCAACATCCTGGGGCACAAGCTGGAGTACAACTACAACAGCCACAACGTCTATATCATGGCCGA
CAAGCAGAAGAACGGCATCAAGGTGAACTTCAAGATCCGCCACAACATCGAGGACGGCAGCGTGCAGCTCGCCGACCACTACCAGCAGAACACCCCAATCGGCGACGGCCCCGTG
CTGCTGCCCGACAACCACTACCTTAGCTACCAGTCCGCCCTGAGCAAAGACCCCAACGAGAAGCGCGATCACATGGTCCTGCTGGAGTTCGTGACCGCCGCCGGGATCACTCTCG
GCATGGACGAGCTGTACAAGTAACTCGGTACCAAATTCCAGAAAAGAGGCCTCCCGAAAGGGGGGCCTTTTTTCGTTTTGGTCCAATGGCGGCGCGCCATCGAATGAGGTGCTTC
CTCGCTCACTGACTCGCTGCACGAGGCAGACCTCAGCGCTAGCGGAGTGTATACTGGCTTACTATGTTGGCACTGATGAGGGTGTCAGTGAAGTGCTTCATGTGGCAGGAGAAAA
AAGGCTGCACCGGTGCGTCAGCAGAATATGTGATACAGGATATATTCCGCTTCCTCGCTCACTGACTCGCTACGCTCGGTCGTTCGACTGCGGCGAGCGGAAATGGCTTACGAAC
GGGGCGGAGATTTCCTGGAAGATGCCAGGAAGATACTTAACAGGGAAGTGAGAGGGCCGCGGCAAAGCCGTTTTTCCATAGGCTCCGCCCCCCTGACAAGCATCACGAAATCTGA
CGCTCAAATCAGTGGTGGCGAAACCCGACAGGACTATAAAGATACCAGGCGTTTCCCCCTGGCGGCTCCCTCGTGCGCTCTCCTGTTCCTGCCTTTCGGTTTACCGGTGTCATTC
CGCTGTTATGGCCGCGTTTGTCTCATTCCACGCCTGACACTCAGTTCCGGGTAGGCAGTTCGCTCCAAGCTGGACTGTATGCACGAACCCCCCGTTCAGTCCGACCGCTGCGCCT
TATCCGGTAACTATCGTCTTGAGTCCAACCCGGAAAGACATGCAAAAGCACCACTGGCAGCAGCCACTGGTAATTGATTTAGAGGAGTTAGTCTTGAAGTCATGCGCCGGTTAAG
GCTAAACTGAAAGGACAAGTTTTGGTGACTGCGCTCCTCCAAGCCAGTTACCTCGGTTCAAAGAGTTGGTAGCTCAGAGAACCTTCGAAAAACCGCCCTGCAAGGCGGTTTTTTC
GTTTTCAGAGCAAGAGATTACGCGCAGACCAAAACGATCTCAAGAAGATCATCTTATTAAGGGGTCTGACGCTCAGTGGAACGAAAAATCAATCTAAAGTATATATGAGTAAACT
TGGTCTGACAGTTACCATGAGCCATATTCAACGGGAAACGTCTTGCTCGAGGCCGCGATTAAATTCCAACATGGATGCTGATTTATATGGGTATAAATGGGCTCGCGATAATGTC
GGGCAATCAGGTGCGACAATCTATCGATTGTATGGGAAGCCCGATGCGCCAGAGTTGTTTCTGAAACATGGCAAAGGTAGCGTTGCCAATGATGTTACAGATGAGATGGTCAGAC
TAAACTGGCTGACGGAATTTATGCCTCTTCCGACCATCAAGCATTTTATCCGTACTCCTGATGATGCATGGTTACTCACCACTGCGATCCCCGGGAAAACAGCATTCCAGGTATT
AGAAGAATATCCTGATTCAGGTGAAAATATTGTTGATGCGCTGGCAGTGTTCCTGCGCCGGTTGCATTCGATTCCTGTTTGTAATTGTCCTTTTAACAGCGATCGCGTATTTCGT
CTCGCTCAGGCGCAATCACGAATGAATAACGGTTTGGTTGATGCGAGTGATTTTGATGACGAGCGTAATGGCTGGCCTGTTGAACAAGTCTGGAAAGAAATGCATAAGCTTTTGC
CATTCTCACCGGATTCAGTCGTCACTCATGGTGATTTCTCACTTGATAACCTTATTTTTGACGAGGGGAAATTAATAGGTTGTATTGATGTTGGACGAGTCGGAATCGCAGACCG
ATACCAGGATCTTGCCATCCTATGGAACTGCCTCGGTGAGTTTTCTCCTTCATTACAGAAACGGCTTTTTCAAAAATATGGTATTGATAATCCTGATATGAATAAATTGCAGTTT
CATTTGATGCTCGATGAGTTTTTCTAAAACACCCCTTGTATTACTGTTTATGTAAGCAGACAGTTTTATTGTTCATGATGATATATTTTTATCTTGTGCAATGTACATCAGAGAT
TTTGAGACACAA

pAJM.715

GenBank accession number: MH101719

CCAATTATTGAAGGCCTCCCTAACGGGGGGCCTTTTTTTGTTTCTGGTCTCCCGCTTAACGATCGTTGGCTGTGTTGACAATTAATCATCGGCTCGTATAATGTGTGGAATTGTG
AGCGCTCACAATTAGCTGTCACCGGATGTGCTTTCCGGTCTGATGAGTCCGTGAGGACGAAACAGCCTCTACAAATAATTTTGTTTAATACTAGAGAAAGAGGGGAAATACTAGA
TGGTGAGCAAGGGCGAGGAGCTGTTCACCGGGGTGGTGCCCATCCTGGTCGAGCTGGACGGCGACGTAAACGGCCACAAGTTCAGCGTGTCCGGCGAGGGCGAGGGCGATGCCAC
CTACGGCAAGCTGACCCTGAAGTTCATCTGCACCACAGGCAAGCTGCCCGTGCCCTGGCCCACCCTCGTGACCACCTTCGGCTACGGCCTGCAATGCTTCGCCCGCTACCCCGAC
CACATGAAGCTGCACGACTTCTTCAAGTCCGCCATGCCCGAAGGCTACGTCCAGGAGCGCACCATCTTCTTCAAGGACGACGGCAACTACAAGACCCGCGCCGAGGTGAAGTTCG
AGGGCGACACCCTGGTGAACCGCATCGAGCTGAAGGGCATCGACTTCAAGGAGGACGGCAACATCCTGGGGCACAAGCTGGAGTACAACTACAACAGCCACAACGTCTATATCAT
GGCCGACAAGCAGAAGAACGGCATCAAGGTGAACTTCAAGATCCGCCACAACATCGAGGACGGCAGCGTGCAGCTCGCCGACCACTACCAGCAGAACACCCCAATCGGCGACGGC
CCCGTGCTGCTGCCCGACAACCACTACCTTAGCTACCAGTCCGCCCTGAGCAAAGACCCCAACGAGAAGCGCGATCACATGGTCCTGCTGGAGTTCGTGACCGCCGCCGGGATCA
CTCTCGGCATGGACGAGCTGTACAAGTAACTCGGTACCAAATTCCAGAAAAGAGGCCTCCCGAAAGGGGGGCCTTTTTTCGTTTTGGTCCAATGGCGGCGCGCCATCGAATGAGG
TGCTTCCTCGCTCACTGACTCGCTGCACGAGGCAGACCTCAGCGCTAGCGGAGTGTATACTGGCTTACTATGTTGGCACTGATGAGGGTGTCAGTGAAGTGCTTCATGTGGCAGG
AGAAAAAAGGCTGCACCGGTGCGTCAGCAGAATATGTGATACAGGATATATTCCGCTTCCTCGCTCACTGACTCGCTACGCTCGGTCGTTCGACTGCGGCGAGCGGAAATGGCTT
ACGAACGGGGCGGAGATTTCCTGGAAGATGCCAGGAAGATACTTAACAGGGAAGTGAGAGGGCCGCGGCAAAGCCGTTTTTCCATAGGCTCCGCCCCCCTGACAAGCATCACGAA
ATCTGACGCTCAAATCAGTGGTGGCGAAACCCGACAGGACTATAAAGATACCAGGCGTTTCCCCCTGGCGGCTCCCTCGTGCGCTCTCCTGTTCCTGCCTTTCGGTTTACCGGTG
TCATTCCGCTGTTATGGCCGCGTTTGTCTCATTCCACGCCTGACACTCAGTTCCGGGTAGGCAGTTCGCTCCAAGCTGGACTGTATGCACGAACCCCCCGTTCAGTCCGACCGCT
GCGCCTTATCCGGTAACTATCGTCTTGAGTCCAACCCGGAAAGACATGCAAAAGCACCACTGGCAGCAGCCACTGGTAATTGATTTAGAGGAGTTAGTCTTGAAGTCATGCGCCG
GTTAAGGCTAAACTGAAAGGACAAGTTTTGGTGACTGCGCTCCTCCAAGCCAGTTACCTCGGTTCAAAGAGTTGGTAGCTCAGAGAACCTTCGAAAAACCGCCCTGCAAGGCGGT
TTTTTCGTTTTCAGAGCAAGAGATTACGCGCAGACCAAAACGATCTCAAGAAGATCATCTTATTAAGGGGTCTGACGCTCAGTGGAACGAAAAATCAATCTAAAGTATATATGAG
TAAACTTGGTCTGACAGTTACCATGAGCCATATTCAACGGGAAACGTCTTGCTCGAGGCCGCGATTAAATTCCAACATGGATGCTGATTTATATGGGTATAAATGGGCTCGCGAT
AATGTCGGGCAATCAGGTGCGACAATCTATCGATTGTATGGGAAGCCCGATGCGCCAGAGTTGTTTCTGAAACATGGCAAAGGTAGCGTTGCCAATGATGTTACAGATGAGATGG
TCAGACTAAACTGGCTGACGGAATTTATGCCTCTTCCGACCATCAAGCATTTTATCCGTACTCCTGATGATGCATGGTTACTCACCACTGCGATCCCCGGGAAAACAGCATTCCA
GGTATTAGAAGAATATCCTGATTCAGGTGAAAATATTGTTGATGCGCTGGCAGTGTTCCTGCGCCGGTTGCATTCGATTCCTGTTTGTAATTGTCCTTTTAACAGCGATCGCGTA
TTTCGTCTCGCTCAGGCGCAATCACGAATGAATAACGGTTTGGTTGATGCGAGTGATTTTGATGACGAGCGTAATGGCTGGCCTGTTGAACAAGTCTGGAAAGAAATGCATAAGC
TTTTGCCATTCTCACCGGATTCAGTCGTCACTCATGGTGATTTCTCACTTGATAACCTTATTTTTGACGAGGGGAAATTAATAGGTTGTATTGATGTTGGACGAGTCGGAATCGC
AGACCGATACCAGGATCTTGCCATCCTATGGAACTGCCTCGGTGAGTTTTCTCCTTCATTACAGAAACGGCTTTTTCAAAAATATGGTATTGATAATCCTGATATGAATAAATTG
CAGTTTCATTTGATGCTCGATGAGTTTTTCTAAAACACCCCTTGTATTACTGTTTATGTAAGCAGACAGTTTTATTGTTCATGATGATATATTTTTATCTTGTGCAATGTACATC
AGAGATTTTGAGACACAA

pAJM.716

GenBank accession number: MH101721

CCAATTATTGAAGGCCTCCCTAACGGGGGGCCTTTTTTTGTTTCTGGTCTCCCGCTTAACGATCGTTGGCTGAGAAACCAATTGTCCATATTGCATCAGACATTGCCGTCACTGC
GTCTTTTACTGGCTCTTCTCGCTAACCAAACCGGTAACCCCGCTTATTAAAAGCATTCTGTAACAAAGCGGGACCAAAGCCATGACAAAAACGCGTAACAAAAGTGTCTATAATC
ACGGCAGAAAAGTCCACATTGATTATTTGCACGGCGTCACACTTTGCTATGCCATAGCATTTTTATCCATAAGATTAGCGGATCCTACCTGACGCTTTTTATCGCAACTCTCTAC
TGTTTCTCCATACCCGAGCTGTCACCGGATGTGCTTTCCGGTCTGATGAGTCCGTGAGGACGAAACAGCCTCTACAAATAATTTTGTTTAATACTAGAGAAAGAGGGGAAATACT
AGATGGTGAGCAAGGGCGAGGAGCTGTTCACCGGGGTGGTGCCCATCCTGGTCGAGCTGGACGGCGACGTAAACGGCCACAAGTTCAGCGTGTCCGGCGAGGGCGAGGGCGATGC
CACCTACGGCAAGCTGACCCTGAAGTTCATCTGCACCACAGGCAAGCTGCCCGTGCCCTGGCCCACCCTCGTGACCACCTTCGGCTACGGCCTGCAATGCTTCGCCCGCTACCCC
GACCACATGAAGCTGCACGACTTCTTCAAGTCCGCCATGCCCGAAGGCTACGTCCAGGAGCGCACCATCTTCTTCAAGGACGACGGCAACTACAAGACCCGCGCCGAGGTGAAGT
TCGAGGGCGACACCCTGGTGAACCGCATCGAGCTGAAGGGCATCGACTTCAAGGAGGACGGCAACATCCTGGGGCACAAGCTGGAGTACAACTACAACAGCCACAACGTCTATAT
CATGGCCGACAAGCAGAAGAACGGCATCAAGGTGAACTTCAAGATCCGCCACAACATCGAGGACGGCAGCGTGCAGCTCGCCGACCACTACCAGCAGAACACCCCAATCGGCGAC
GGCCCCGTGCTGCTGCCCGACAACCACTACCTTAGCTACCAGTCCGCCCTGAGCAAAGACCCCAACGAGAAGCGCGATCACATGGTCCTGCTGGAGTTCGTGACCGCCGCCGGGA
TCACTCTCGGCATGGACGAGCTGTACAAGTAACTCGGTACCAAATTCCAGAAAAGAGGCCTCCCGAAAGGGGGGCCTTTTTTCGTTTTGGTCCAATGGCGGCGCGCCATCGAATG
AGGTGCTTCCTCGCTCACTGACTCGCTGCACGAGGCAGACCTCAGCGCTAGCGGAGTGTATACTGGCTTACTATGTTGGCACTGATGAGGGTGTCAGTGAAGTGCTTCATGTGGC
AGGAGAAAAAAGGCTGCACCGGTGCGTCAGCAGAATATGTGATACAGGATATATTCCGCTTCCTCGCTCACTGACTCGCTACGCTCGGTCGTTCGACTGCGGCGAGCGGAAATGG
CTTACGAACGGGGCGGAGATTTCCTGGAAGATGCCAGGAAGATACTTAACAGGGAAGTGAGAGGGCCGCGGCAAAGCCGTTTTTCCATAGGCTCCGCCCCCCTGACAAGCATCAC
GAAATCTGACGCTCAAATCAGTGGTGGCGAAACCCGACAGGACTATAAAGATACCAGGCGTTTCCCCCTGGCGGCTCCCTCGTGCGCTCTCCTGTTCCTGCCTTTCGGTTTACCG
GTGTCATTCCGCTGTTATGGCCGCGTTTGTCTCATTCCACGCCTGACACTCAGTTCCGGGTAGGCAGTTCGCTCCAAGCTGGACTGTATGCACGAACCCCCCGTTCAGTCCGACC
GCTGCGCCTTATCCGGTAACTATCGTCTTGAGTCCAACCCGGAAAGACATGCAAAAGCACCACTGGCAGCAGCCACTGGTAATTGATTTAGAGGAGTTAGTCTTGAAGTCATGCG
CCGGTTAAGGCTAAACTGAAAGGACAAGTTTTGGTGACTGCGCTCCTCCAAGCCAGTTACCTCGGTTCAAAGAGTTGGTAGCTCAGAGAACCTTCGAAAAACCGCCCTGCAAGGC
GGTTTTTTCGTTTTCAGAGCAAGAGATTACGCGCAGACCAAAACGATCTCAAGAAGATCATCTTATTAAGGGGTCTGACGCTCAGTGGAACGAAAAATCAATCTAAAGTATATAT
GAGTAAACTTGGTCTGACAGTTACCATGAGCCATATTCAACGGGAAACGTCTTGCTCGAGGCCGCGATTAAATTCCAACATGGATGCTGATTTATATGGGTATAAATGGGCTCGC
GATAATGTCGGGCAATCAGGTGCGACAATCTATCGATTGTATGGGAAGCCCGATGCGCCAGAGTTGTTTCTGAAACATGGCAAAGGTAGCGTTGCCAATGATGTTACAGATGAGA
TGGTCAGACTAAACTGGCTGACGGAATTTATGCCTCTTCCGACCATCAAGCATTTTATCCGTACTCCTGATGATGCATGGTTACTCACCACTGCGATCCCCGGGAAAACAGCATT
CCAGGTATTAGAAGAATATCCTGATTCAGGTGAAAATATTGTTGATGCGCTGGCAGTGTTCCTGCGCCGGTTGCATTCGATTCCTGTTTGTAATTGTCCTTTTAACAGCGATCGC
GTATTTCGTCTCGCTCAGGCGCAATCACGAATGAATAACGGTTTGGTTGATGCGAGTGATTTTGATGACGAGCGTAATGGCTGGCCTGTTGAACAAGTCTGGAAAGAAATGCATA
AGCTTTTGCCATTCTCACCGGATTCAGTCGTCACTCATGGTGATTTCTCACTTGATAACCTTATTTTTGACGAGGGGAAATTAATAGGTTGTATTGATGTTGGACGAGTCGGAAT
CGCAGACCGATACCAGGATCTTGCCATCCTATGGAACTGCCTCGGTGAGTTTTCTCCTTCATTACAGAAACGGCTTTTTCAAAAATATGGTATTGATAATCCTGATATGAATAAA

TTGCAGTTTCATTTGATGCTCGATGAGTTTTTCTAAAACACCCCTTGTATTACTGTTTATGTAAGCAGACAGTTTTATTGTTCATGATGATATATTTTTATCTTGTGCAATGTAC
ATCAGAGATTTTGAGACACAA

pAJM.717

GenBank accession number: MH101720

CCAATTATTGAAGGCCTCCCTAACGGGGGGCCTTTTTTTGTTTCTGGTCTCCCGCTTAACGATCGTTGGCTGTTTTCAGCAGGACGCACTGACCTCCCTATCAGTGATAGAGATT
GACATCCCTATCAGTGATAGAGATACTGAGCACAGCTGTCACCGGATGTGCTTTCCGGTCTGATGAGTCCGTGAGGACGAAACAGCCTCTACAAATAATTTTGTTTAATACTAGA
GAAAGAGGGGAAATACTAGATGGTGAGCAAGGGCGAGGAGCTGTTCACCGGGGTGGTGCCCATCCTGGTCGAGCTGGACGGCGACGTAAACGGCCACAAGTTCAGCGTGTCCGGC
GAGGGCGAGGGCGATGCCACCTACGGCAAGCTGACCCTGAAGTTCATCTGCACCACAGGCAAGCTGCCCGTGCCCTGGCCCACCCTCGTGACCACCTTCGGCTACGGCCTGCAAT
GCTTCGCCCGCTACCCCGACCACATGAAGCTGCACGACTTCTTCAAGTCCGCCATGCCCGAAGGCTACGTCCAGGAGCGCACCATCTTCTTCAAGGACGACGGCAACTACAAGAC
CCGCGCCGAGGTGAAGTTCGAGGGCGACACCCTGGTGAACCGCATCGAGCTGAAGGGCATCGACTTCAAGGAGGACGGCAACATCCTGGGGCACAAGCTGGAGTACAACTACAAC
AGCCACAACGTCTATATCATGGCCGACAAGCAGAAGAACGGCATCAAGGTGAACTTCAAGATCCGCCACAACATCGAGGACGGCAGCGTGCAGCTCGCCGACCACTACCAGCAGA
ACACCCCAATCGGCGACGGCCCCGTGCTGCTGCCCGACAACCACTACCTTAGCTACCAGTCCGCCCTGAGCAAAGACCCCAACGAGAAGCGCGATCACATGGTCCTGCTGGAGTT
CGTGACCGCCGCCGGGATCACTCTCGGCATGGACGAGCTGTACAAGTAACTCGGTACCAAATTCCAGAAAAGAGGCCTCCCGAAAGGGGGGCCTTTTTTCGTTTTGGTCCAATGG
CGGCGCGCCATCGAATGAGGTGCTTCCTCGCTCACTGACTCGCTGCACGAGGCAGACCTCAGCGCTAGCGGAGTGTATACTGGCTTACTATGTTGGCACTGATGAGGGTGTCAGT
GAAGTGCTTCATGTGGCAGGAGAAAAAAGGCTGCACCGGTGCGTCAGCAGAATATGTGATACAGGATATATTCCGCTTCCTCGCTCACTGACTCGCTACGCTCGGTCGTTCGACT
GCGGCGAGCGGAAATGGCTTACGAACGGGGCGGAGATTTCCTGGAAGATGCCAGGAAGATACTTAACAGGGAAGTGAGAGGGCCGCGGCAAAGCCGTTTTTCCATAGGCTCCGCC
CCCCTGACAAGCATCACGAAATCTGACGCTCAAATCAGTGGTGGCGAAACCCGACAGGACTATAAAGATACCAGGCGTTTCCCCCTGGCGGCTCCCTCGTGCGCTCTCCTGTTCC
TGCCTTTCGGTTTACCGGTGTCATTCCGCTGTTATGGCCGCGTTTGTCTCATTCCACGCCTGACACTCAGTTCCGGGTAGGCAGTTCGCTCCAAGCTGGACTGTATGCACGAACC
CCCCGTTCAGTCCGACCGCTGCGCCTTATCCGGTAACTATCGTCTTGAGTCCAACCCGGAAAGACATGCAAAAGCACCACTGGCAGCAGCCACTGGTAATTGATTTAGAGGAGTT
AGTCTTGAAGTCATGCGCCGGTTAAGGCTAAACTGAAAGGACAAGTTTTGGTGACTGCGCTCCTCCAAGCCAGTTACCTCGGTTCAAAGAGTTGGTAGCTCAGAGAACCTTCGAA
AAACCGCCCTGCAAGGCGGTTTTTTCGTTTTCAGAGCAAGAGATTACGCGCAGACCAAAACGATCTCAAGAAGATCATCTTATTAAGGGGTCTGACGCTCAGTGGAACGAAAAAT
CAATCTAAAGTATATATGAGTAAACTTGGTCTGACAGTTACCATGAGCCATATTCAACGGGAAACGTCTTGCTCGAGGCCGCGATTAAATTCCAACATGGATGCTGATTTATATG
GGTATAAATGGGCTCGCGATAATGTCGGGCAATCAGGTGCGACAATCTATCGATTGTATGGGAAGCCCGATGCGCCAGAGTTGTTTCTGAAACATGGCAAAGGTAGCGTTGCCAA
TGATGTTACAGATGAGATGGTCAGACTAAACTGGCTGACGGAATTTATGCCTCTTCCGACCATCAAGCATTTTATCCGTACTCCTGATGATGCATGGTTACTCACCACTGCGATC
CCCGGGAAAACAGCATTCCAGGTATTAGAAGAATATCCTGATTCAGGTGAAAATATTGTTGATGCGCTGGCAGTGTTCCTGCGCCGGTTGCATTCGATTCCTGTTTGTAATTGTC
CTTTTAACAGCGATCGCGTATTTCGTCTCGCTCAGGCGCAATCACGAATGAATAACGGTTTGGTTGATGCGAGTGATTTTGATGACGAGCGTAATGGCTGGCCTGTTGAACAAGT
CTGGAAAGAAATGCATAAGCTTTTGCCATTCTCACCGGATTCAGTCGTCACTCATGGTGATTTCTCACTTGATAACCTTATTTTTGACGAGGGGAAATTAATAGGTTGTATTGAT
GTTGGACGAGTCGGAATCGCAGACCGATACCAGGATCTTGCCATCCTATGGAACTGCCTCGGTGAGTTTTCTCCTTCATTACAGAAACGGCTTTTTCAAAAATATGGTATTGATA
ATCCTGATATGAATAAATTGCAGTTTCATTTGATGCTCGATGAGTTTTTCTAAAACACCCCTTGTATTACTGTTTATGTAAGCAGACAGTTTTATTGTTCATGATGATATATTTT
TATCTTGTGCAATGTACATCAGAGATTTTGAGACACAA

pAJM.718

GenBank accession number: MH101722

CCAATTATTGAAGGCCTCCCTAACGGGGGGCCTTTTTTTGTTTCTGGTCTCCCGCTTAACGATCGTTGGCTGAGCGCGGGTGAGAGGGATTCGTTACCAATAGACAATTGATTGG
ACGTTCAATATAATGCTAGCAGCTGTCACCGGATGTGCTTTCCGGTCTGATGAGTCCGTGAGGACGAAACAGCCTCTACAAATAATTTTGTTTAATACTAGAGAAAGAGGGGAAA
TACTAGATGGTGAGCAAGGGCGAGGAGCTGTTCACCGGGGTGGTGCCCATCCTGGTCGAGCTGGACGGCGACGTAAACGGCCACAAGTTCAGCGTGTCCGGCGAGGGCGAGGGCG
ATGCCACCTACGGCAAGCTGACCCTGAAGTTCATCTGCACCACAGGCAAGCTGCCCGTGCCCTGGCCCACCCTCGTGACCACCTTCGGCTACGGCCTGCAATGCTTCGCCCGCTA
CCCCGACCACATGAAGCTGCACGACTTCTTCAAGTCCGCCATGCCCGAAGGCTACGTCCAGGAGCGCACCATCTTCTTCAAGGACGACGGCAACTACAAGACCCGCGCCGAGGTG
AAGTTCGAGGGCGACACCCTGGTGAACCGCATCGAGCTGAAGGGCATCGACTTCAAGGAGGACGGCAACATCCTGGGGCACAAGCTGGAGTACAACTACAACAGCCACAACGTCT
ATATCATGGCCGACAAGCAGAAGAACGGCATCAAGGTGAACTTCAAGATCCGCCACAACATCGAGGACGGCAGCGTGCAGCTCGCCGACCACTACCAGCAGAACACCCCAATCGG
CGACGGCCCCGTGCTGCTGCCCGACAACCACTACCTTAGCTACCAGTCCGCCCTGAGCAAAGACCCCAACGAGAAGCGCGATCACATGGTCCTGCTGGAGTTCGTGACCGCCGCC
GGGATCACTCTCGGCATGGACGAGCTGTACAAGTAACTCGGTACCAAATTCCAGAAAAGAGGCCTCCCGAAAGGGGGGCCTTTTTTCGTTTTGGTCCAATGGCGGCGCGCCATCG
AATGAGGTGCTTCCTCGCTCACTGACTCGCTGCACGAGGCAGACCTCAGCGCTAGCGGAGTGTATACTGGCTTACTATGTTGGCACTGATGAGGGTGTCAGTGAAGTGCTTCATG
TGGCAGGAGAAAAAAGGCTGCACCGGTGCGTCAGCAGAATATGTGATACAGGATATATTCCGCTTCCTCGCTCACTGACTCGCTACGCTCGGTCGTTCGACTGCGGCGAGCGGAA
ATGGCTTACGAACGGGGCGGAGATTTCCTGGAAGATGCCAGGAAGATACTTAACAGGGAAGTGAGAGGGCCGCGGCAAAGCCGTTTTTCCATAGGCTCCGCCCCCCTGACAAGCA
TCACGAAATCTGACGCTCAAATCAGTGGTGGCGAAACCCGACAGGACTATAAAGATACCAGGCGTTTCCCCCTGGCGGCTCCCTCGTGCGCTCTCCTGTTCCTGCCTTTCGGTTT
ACCGGTGTCATTCCGCTGTTATGGCCGCGTTTGTCTCATTCCACGCCTGACACTCAGTTCCGGGTAGGCAGTTCGCTCCAAGCTGGACTGTATGCACGAACCCCCCGTTCAGTCC
GACCGCTGCGCCTTATCCGGTAACTATCGTCTTGAGTCCAACCCGGAAAGACATGCAAAAGCACCACTGGCAGCAGCCACTGGTAATTGATTTAGAGGAGTTAGTCTTGAAGTCA
TGCGCCGGTTAAGGCTAAACTGAAAGGACAAGTTTTGGTGACTGCGCTCCTCCAAGCCAGTTACCTCGGTTCAAAGAGTTGGTAGCTCAGAGAACCTTCGAAAAACCGCCCTGCA
AGGCGGTTTTTTCGTTTTCAGAGCAAGAGATTACGCGCAGACCAAAACGATCTCAAGAAGATCATCTTATTAAGGGGTCTGACGCTCAGTGGAACGAAAAATCAATCTAAAGTAT
ATATGAGTAAACTTGGTCTGACAGTTACCATGAGCCATATTCAACGGGAAACGTCTTGCTCGAGGCCGCGATTAAATTCCAACATGGATGCTGATTTATATGGGTATAAATGGGC
TCGCGATAATGTCGGGCAATCAGGTGCGACAATCTATCGATTGTATGGGAAGCCCGATGCGCCAGAGTTGTTTCTGAAACATGGCAAAGGTAGCGTTGCCAATGATGTTACAGAT
GAGATGGTCAGACTAAACTGGCTGACGGAATTTATGCCTCTTCCGACCATCAAGCATTTTATCCGTACTCCTGATGATGCATGGTTACTCACCACTGCGATCCCCGGGAAAACAG
CATTCCAGGTATTAGAAGAATATCCTGATTCAGGTGAAAATATTGTTGATGCGCTGGCAGTGTTCCTGCGCCGGTTGCATTCGATTCCTGTTTGTAATTGTCCTTTTAACAGCGA
TCGCGTATTTCGTCTCGCTCAGGCGCAATCACGAATGAATAACGGTTTGGTTGATGCGAGTGATTTTGATGACGAGCGTAATGGCTGGCCTGTTGAACAAGTCTGGAAAGAAATG
CATAAGCTTTTGCCATTCTCACCGGATTCAGTCGTCACTCATGGTGATTTCTCACTTGATAACCTTATTTTTGACGAGGGGAAATTAATAGGTTGTATTGATGTTGGACGAGTCG
GAATCGCAGACCGATACCAGGATCTTGCCATCCTATGGAACTGCCTCGGTGAGTTTTCTCCTTCATTACAGAAACGGCTTTTTCAAAAATATGGTATTGATAATCCTGATATGAA
TAAATTGCAGTTTCATTTGATGCTCGATGAGTTTTTCTAAAACACCCCTTGTATTACTGTTTATGTAAGCAGACAGTTTTATTGTTCATGATGATATATTTTTATCTTGTGCAAT
GTACATCAGAGATTTTGAGACACAA

pAJM.719

GenBank accession number: MH101723

CCAATTATTGAAGGCCTCCCTAACGGGGGGCCTTTTTTTGTTTCTGGTCTCCCGCTTAACGATCGTTGGCTGCACCCAGCAGTATTTACAAACAACCATGAATGTAAGTATATTC
CTTAGCAAAGCTGTCACCGGATGTGCTTTCCGGTCTGATGAGTCCGTGAGGACGAAACAGCCTCTACAAATAATTTTGTTTAATACTAGAGAAAGAGGGGAAATACTAGATGGTG
AGCAAGGGCGAGGAGCTGTTCACCGGGGTGGTGCCCATCCTGGTCGAGCTGGACGGCGACGTAAACGGCCACAAGTTCAGCGTGTCCGGCGAGGGCGAGGGCGATGCCACCTACG
GCAAGCTGACCCTGAAGTTCATCTGCACCACAGGCAAGCTGCCCGTGCCCTGGCCCACCCTCGTGACCACCTTCGGCTACGGCCTGCAATGCTTCGCCCGCTACCCCGACCACAT
GAAGCTGCACGACTTCTTCAAGTCCGCCATGCCCGAAGGCTACGTCCAGGAGCGCACCATCTTCTTCAAGGACGACGGCAACTACAAGACCCGCGCCGAGGTGAAGTTCGAGGGC
GACACCCTGGTGAACCGCATCGAGCTGAAGGGCATCGACTTCAAGGAGGACGGCAACATCCTGGGGCACAAGCTGGAGTACAACTACAACAGCCACAACGTCTATATCATGGCCG
ACAAGCAGAAGAACGGCATCAAGGTGAACTTCAAGATCCGCCACAACATCGAGGACGGCAGCGTGCAGCTCGCCGACCACTACCAGCAGAACACCCCAATCGGCGACGGCCCCGT
GCTGCTGCCCGACAACCACTACCTTAGCTACCAGTCCGCCCTGAGCAAAGACCCCAACGAGAAGCGCGATCACATGGTCCTGCTGGAGTTCGTGACCGCCGCCGGGATCACTCTC
GGCATGGACGAGCTGTACAAGTAACTCGGTACCAAATTCCAGAAAAGAGGCCTCCCGAAAGGGGGGCCTTTTTTCGTTTTGGTCCAATGGCGGCGCGCCATCGAATGAGGTGCTT
CCTCGCTCACTGACTCGCTGCACGAGGCAGACCTCAGCGCTAGCGGAGTGTATACTGGCTTACTATGTTGGCACTGATGAGGGTGTCAGTGAAGTGCTTCATGTGGCAGGAGAAA
AAAGGCTGCACCGGTGCGTCAGCAGAATATGTGATACAGGATATATTCCGCTTCCTCGCTCACTGACTCGCTACGCTCGGTCGTTCGACTGCGGCGAGCGGAAATGGCTTACGAA
CGGGGCGGAGATTTCCTGGAAGATGCCAGGAAGATACTTAACAGGGAAGTGAGAGGGCCGCGGCAAAGCCGTTTTTCCATAGGCTCCGCCCCCCTGACAAGCATCACGAAATCTG
ACGCTCAAATCAGTGGTGGCGAAACCCGACAGGACTATAAAGATACCAGGCGTTTCCCCCTGGCGGCTCCCTCGTGCGCTCTCCTGTTCCTGCCTTTCGGTTTACCGGTGTCATT
CCGCTGTTATGGCCGCGTTTGTCTCATTCCACGCCTGACACTCAGTTCCGGGTAGGCAGTTCGCTCCAAGCTGGACTGTATGCACGAACCCCCCGTTCAGTCCGACCGCTGCGCC
TTATCCGGTAACTATCGTCTTGAGTCCAACCCGGAAAGACATGCAAAAGCACCACTGGCAGCAGCCACTGGTAATTGATTTAGAGGAGTTAGTCTTGAAGTCATGCGCCGGTTAA
GGCTAAACTGAAAGGACAAGTTTTGGTGACTGCGCTCCTCCAAGCCAGTTACCTCGGTTCAAAGAGTTGGTAGCTCAGAGAACCTTCGAAAAACCGCCCTGCAAGGCGGTTTTTT
CGTTTTCAGAGCAAGAGATTACGCGCAGACCAAAACGATCTCAAGAAGATCATCTTATTAAGGGGTCTGACGCTCAGTGGAACGAAAAATCAATCTAAAGTATATATGAGTAAAC
TTGGTCTGACAGTTACCATGAGCCATATTCAACGGGAAACGTCTTGCTCGAGGCCGCGATTAAATTCCAACATGGATGCTGATTTATATGGGTATAAATGGGCTCGCGATAATGT
CGGGCAATCAGGTGCGACAATCTATCGATTGTATGGGAAGCCCGATGCGCCAGAGTTGTTTCTGAAACATGGCAAAGGTAGCGTTGCCAATGATGTTACAGATGAGATGGTCAGA
CTAAACTGGCTGACGGAATTTATGCCTCTTCCGACCATCAAGCATTTTATCCGTACTCCTGATGATGCATGGTTACTCACCACTGCGATCCCCGGGAAAACAGCATTCCAGGTAT
TAGAAGAATATCCTGATTCAGGTGAAAATATTGTTGATGCGCTGGCAGTGTTCCTGCGCCGGTTGCATTCGATTCCTGTTTGTAATTGTCCTTTTAACAGCGATCGCGTATTTCG
TCTCGCTCAGGCGCAATCACGAATGAATAACGGTTTGGTTGATGCGAGTGATTTTGATGACGAGCGTAATGGCTGGCCTGTTGAACAAGTCTGGAAAGAAATGCATAAGCTTTTG
CCATTCTCACCGGATTCAGTCGTCACTCATGGTGATTTCTCACTTGATAACCTTATTTTTGACGAGGGGAAATTAATAGGTTGTATTGATGTTGGACGAGTCGGAATCGCAGACC
GATACCAGGATCTTGCCATCCTATGGAACTGCCTCGGTGAGTTTTCTCCTTCATTACAGAAACGGCTTTTTCAAAAATATGGTATTGATAATCCTGATATGAATAAATTGCAGTT
TCATTTGATGCTCGATGAGTTTTTCTAAAACACCCCTTGTATTACTGTTTATGTAAGCAGACAGTTTTATTGTTCATGATGATATATTTTTATCTTGTGCAATGTACATCAGAGA
TTTTGAGACACAA

pAJM.721

GenBank accession number: MH101725

CCAATTATTGAAGGCCTCCCTAACGGGGGGCCTTTTTTTGTTTCTGGTCTCCCGCTTAACGATCGTTGGCTGGGGGCCTCGCTTGGGTTATTGCTGGTGCCCGGCCGGGCGCAAT
ATTCATGTTGATGATTTATTATATATCGAGTGGTGTATTTATTTATATTGTTTGCTCCGTTACCGTTATTAACAGCTGTCACCGGATGTGCTTTCCGGTCTGATGAGTCCGTGAG
GACGAAACAGCCTCTACAAATAATTTTGTTTAATACTAGAGAAAGAGGGGAAATACTAGATGGTGAGCAAGGGCGAGGAGCTGTTCACCGGGGTGGTGCCCATCCTGGTCGAGCT
GGACGGCGACGTAAACGGCCACAAGTTCAGCGTGTCCGGCGAGGGCGAGGGCGATGCCACCTACGGCAAGCTGACCCTGAAGTTCATCTGCACCACAGGCAAGCTGCCCGTGCCC
TGGCCCACCCTCGTGACCACCTTCGGCTACGGCCTGCAATGCTTCGCCCGCTACCCCGACCACATGAAGCTGCACGACTTCTTCAAGTCCGCCATGCCCGAAGGCTACGTCCAGG
AGCGCACCATCTTCTTCAAGGACGACGGCAACTACAAGACCCGCGCCGAGGTGAAGTTCGAGGGCGACACCCTGGTGAACCGCATCGAGCTGAAGGGCATCGACTTCAAGGAGGA
CGGCAACATCCTGGGGCACAAGCTGGAGTACAACTACAACAGCCACAACGTCTATATCATGGCCGACAAGCAGAAGAACGGCATCAAGGTGAACTTCAAGATCCGCCACAACATC
GAGGACGGCAGCGTGCAGCTCGCCGACCACTACCAGCAGAACACCCCAATCGGCGACGGCCCCGTGCTGCTGCCCGACAACCACTACCTTAGCTACCAGTCCGCCCTGAGCAAAG
ACCCCAACGAGAAGCGCGATCACATGGTCCTGCTGGAGTTCGTGACCGCCGCCGGGATCACTCTCGGCATGGACGAGCTGTACAAGTAACTCGGTACCAAATTCCAGAAAAGAGG
CCTCCCGAAAGGGGGGCCTTTTTTCGTTTTGGTCCAATGGCGGCGCGCCATCGAATGAGGTGCTTCCTCGCTCACTGACTCGCTGCACGAGGCAGACCTCAGCGCTAGCGGAGTG
TATACTGGCTTACTATGTTGGCACTGATGAGGGTGTCAGTGAAGTGCTTCATGTGGCAGGAGAAAAAAGGCTGCACCGGTGCGTCAGCAGAATATGTGATACAGGATATATTCCG

CTTCCTCGCTCACTGACTCGCTACGCTCGGTCGTTCGACTGCGGCGAGCGGAAATGGCTTACGAACGGGGCGGAGATTTCCTGGAAGATGCCAGGAAGATACTTAACAGGGAAGT
GAGAGGGCCGCGGCAAAGCCGTTTTTCCATAGGCTCCGCCCCCCTGACAAGCATCACGAAATCTGACGCTCAAATCAGTGGTGGCGAAACCCGACAGGACTATAAAGATACCAGG
CGTTTCCCCCTGGCGGCTCCCTCGTGCGCTCTCCTGTTCCTGCCTTTCGGTTTACCGGTGTCATTCCGCTGTTATGGCCGCGTTTGTCTCATTCCACGCCTGACACTCAGTTCCG
GGTAGGCAGTTCGCTCCAAGCTGGACTGTATGCACGAACCCCCCGTTCAGTCCGACCGCTGCGCCTTATCCGGTAACTATCGTCTTGAGTCCAACCCGGAAAGACATGCAAAAGC
ACCACTGGCAGCAGCCACTGGTAATTGATTTAGAGGAGTTAGTCTTGAAGTCATGCGCCGGTTAAGGCTAAACTGAAAGGACAAGTTTTGGTGACTGCGCTCCTCCAAGCCAGTT
ACCTCGGTTCAAAGAGTTGGTAGCTCAGAGAACCTTCGAAAAACCGCCCTGCAAGGCGGTTTTTTCGTTTTCAGAGCAAGAGATTACGCGCAGACCAAAACGATCTCAAGAAGAT
CATCTTATTAAGGGGTCTGACGCTCAGTGGAACGAAAAATCAATCTAAAGTATATATGAGTAAACTTGGTCTGACAGTTACCATGAGCCATATTCAACGGGAAACGTCTTGCTCG
AGGCCGCGATTAAATTCCAACATGGATGCTGATTTATATGGGTATAAATGGGCTCGCGATAATGTCGGGCAATCAGGTGCGACAATCTATCGATTGTATGGGAAGCCCGATGCGC
CAGAGTTGTTTCTGAAACATGGCAAAGGTAGCGTTGCCAATGATGTTACAGATGAGATGGTCAGACTAAACTGGCTGACGGAATTTATGCCTCTTCCGACCATCAAGCATTTTAT
CCGTACTCCTGATGATGCATGGTTACTCACCACTGCGATCCCCGGGAAAACAGCATTCCAGGTATTAGAAGAATATCCTGATTCAGGTGAAAATATTGTTGATGCGCTGGCAGTG
TTCCTGCGCCGGTTGCATTCGATTCCTGTTTGTAATTGTCCTTTTAACAGCGATCGCGTATTTCGTCTCGCTCAGGCGCAATCACGAATGAATAACGGTTTGGTTGATGCGAGTG
ATTTTGATGACGAGCGTAATGGCTGGCCTGTTGAACAAGTCTGGAAAGAAATGCATAAGCTTTTGCCATTCTCACCGGATTCAGTCGTCACTCATGGTGATTTCTCACTTGATAA
CCTTATTTTTGACGAGGGGAAATTAATAGGTTGTATTGATGTTGGACGAGTCGGAATCGCAGACCGATACCAGGATCTTGCCATCCTATGGAACTGCCTCGGTGAGTTTTCTCCT
TCATTACAGAAACGGCTTTTTCAAAAATATGGTATTGATAATCCTGATATGAATAAATTGCAGTTTCATTTGATGCTCGATGAGTTTTTCTAAAACACCCCTTGTATTACTGTTT
ATGTAAGCAGACAGTTTTATTGTTCATGATGATATATTTTTATCTTGTGCAATGTACATCAGAGATTTTGAGACACAA

pAJM.771

GenBank accession number: MH101737

CCAATTATTGAAGGCCTCCCTAACGGGGGGCCTTTTTTTGTTTCTGGTCTCCCGCTTAACGATCGTTGGCTGGGGGCCTCGCTTGGGTTATTGCTGGTGCCCGGCCGGGCGCAAT
ATTCATGTTGATGATTTATTATATATCGAGTGGTGTATTTATTTATATTGTTTGCTCCGTTACCGTTATTAACAGCTGTCACCGGATGTGCTTTCCGGTCTGATGAGTCCGTGAG
GACGAAACAGCCTCTACAAATAATTTTGTTTAATACTAGAGAAAGAGGGGAAATACTAGATGGTGAGCAAGGGCGAGGAGCTGTTCACCGGGGTGGTGCCCATCCTGGTCGAGCT
GGACGGCGACGTAAACGGCCACAAGTTCAGCGTGTCCGGCGAGGGCGAGGGCGATGCCACCTACGGCAAGCTGACCCTGAAGTTCATCTGCACCACAGGCAAGCTGCCCGTGCCC
TGGCCCACCCTCGTGACCACCTTCGGCTACGGCCTGCAATGCTTCGCCCGCTACCCCGACCACATGAAGCTGCACGACTTCTTCAAGTCCGCCATGCCCGAAGGCTACGTCCAGG
AGCGCACCATCTTCTTCAAGGACGACGGCAACTACAAGACCCGCGCCGAGGTGAAGTTCGAGGGCGACACCCTGGTGAACCGCATCGAGCTGAAGGGCATCGACTTCAAGGAGGA
CGGCAACATCCTGGGGCACAAGCTGGAGTACAACTACAACAGCCACAACGTCTATATCATGGCCGACAAGCAGAAGAACGGCATCAAGGTGAACTTCAAGATCCGCCACAACATC
GAGGACGGCAGCGTGCAGCTCGCCGACCACTACCAGCAGAACACCCCCAATCGGCGACGGCCCCGTGCTGCTGCCCGACAACCACTACCTTAGCTACCAGTCCGCCCTGAGCAAAG
ACCCCAACGAGAAGCGCGATCACATGGTCCTGCTGGAGTTCGTGACCGCCGCCGGGATCACTCTCGGCATGGACGAGCTGTACAAGTAACTCGGTACCAAATTCCAGAAAAGAGG
CCTCCCGAAAGGGGGGCCTTTTCGTTTTGGTCCAATGGCGGCGCGCCATCGAATGGTGCAAAACCTTTCGCGGTATGGCATGATAGCGCCCGGAAGAGAGTCAATTCATGGGGG
TTGAATATGGAACTGCGTGACCTGGATTTAAACCTGCTGGTGGTGTTCAACCAGTTGCTGGTCGACAGACGCGTCTCTGTCACTGCGGAGAACCTGGGCCTGACCCAGCCTGCCG
TGAGCAATGCGCTGAAACGCCTGCGCACCTCGCTACAGGACCCACTCTTCGTGCGCACACATCAGGGAATGGAACCCACACCCTATGCCGCGCATCTGGCCGAGCACGTCACTTC
GGCCATGCACGCACTGCGCAACGCCCTACAGCACCATGAAAGCTTCGATCCGCTGACCAGCGAGCGTACCTTCACCCTGGCCATGACCGACATTGGCGAGATCTACTTCATGCCG
CGGCTGATGGATGCGCTGGCTCACCAGGCCCCCAATTGCGTGATCAGTACGGTGCGCGACAGTTCGATGAGCCTGATGCAGGCCTTGCAGAACGGAACCGTGGACTTGGCCGTGG
GCCTGCTTCCCAATCTGCAAACTGGCTTCTTTCAGCGCCGGCTGCTCCAGAATCACTACGTGTGCCTATGTCGCAAGGACCATCCAGTCACCCGCGAACCCCTGACTCTGGAGCG
CTTCTGTTCCTACGGCCACGTGCGTGTCATCGCCGCTGGCACCGGCCACGGCGAGGTGGACACGTACATGACACGGGTCGGCATCCGGCGCGACATCCGTCTGGAAGTGCCGCAC
TTCGCCGCCGTTGGCCACATCCTCCAGCGCACCGATCTGCTCGCCACTGTGCCGATATGTTTAGCCGACTGCTGCGTAGAGCCCTTCGGCCTAAGCGCCTTGCCGCACCCAGTCG
TCTTGCCTGAAATAGCCATCAACATGTTCTGGCATGCGAAGTACCACAAGGACCTAGCCAATATTTGGTTGCGGCAACTGATGTTTGACCTGTTTACGGATTGATAAGGATCCTA
ATTGGTAACGAATCAGACAATTGACGGCTCGAGGGAGTAGCATAGGGTTTGCAGAATCCCTGCTTCGTCCATTTGACAGGCACATTATGCATCGATGATAAGCTGTCAAACATGA
GCAGATCCTCTACGCCGGACGCATCGTGGCCGGCATCACCGGCGCCACAGGTGCGGTTGCTGGCGCCTATATCGCCGACATCACCGATGGGGAAGATCGGGCTCGCCACTTCGGG
CTCATGAGCAAATATTTTATCTGAGGTGCTTCCTCGCTCACTGACTCGCTGCACGAGGCAGACCTCAGCGCTAGCGGAGTGTATACTGGCTTACTATGTTGGCACTGATGAGGGT
GTCAGTGAAGTGCTTCATGTGGCAGGAGAAAAAAGGCTGCACCGGTGCGTCAGCAGAATATGTGATACAGGATATATTCCGCTTCCTCGCTCACTGACTCGCTACGCTCGGTCGT
TCGACTGCGGCGAGCGGAAATGGCTTACGAACGGGGCGGAGATTTCCTGGAAGATGCCAGGAAGATACTTAACAGGGAAGTGAGAGGGCCGCGGCAAAGCCGTTTTTCCATAGGC
TCCGCCCCCCTGACAAGCATCACGAAATCTGACGCTCAAATCAGTGGTGGCGAAACCCGACAGGACTATAAAGATACCAGGCGTTTCCCCCTGGCGGCTCCCTCGTGCGCTCTCC
TGTTCCTGCCTTTCGGTTTACCGGTGTCATTCCGCTGTTATGGCCGCGTTTGTCTCATTCCACGCCTGACACTCAGTTCCGGGTAGGCAGTTCGCTCCAAGCTGGACTGTATGCA
CGAACCCCCCGTTCAGTCCGACCGCTGCGCCTTATCCGGTAACTATCGTCTTGAGTCCAACCCGGAAAGACATGCAAAAGCACCACTGGCAGCAGCCACTGGTAATTGATTTAGA
GGAGTTAGTCTTGAAGTCATGCGCCGGTTAAGGCTAAACTGAAAGGACAAGTTTTGGTGACTGCGCTCCTCCAAGCCAGTTACCTCGGTTCAAAGAGTTGGTAGCTCAGAGAACC
TTCGAAAAACCGCCCTGCAAGGCGGTTTTTTCGTTTTCAGAGCAAGAGATTACGCGCAGACCAAAACGATCTCAAGAAGATCATCTTATTAAGGGGTCTGACGCTCAGTGGAACG
AAAAATCAATCTAAAGTATATATGAGTAAACTTGGTCTGACAGTTACCATGAGCCATATTCAACGGGAAACGTCTTGCTCGAGGCCGCGATTAAATTCCAACATGGATGCTGATT
TATATGGGTATAAATGGGCTCGCGATAATGTCGGGCAATCAGGTGCGACAATCTATCGATTGTATGGGAAGCCCGATGCGCCAGAGTTGTTTCTGAAACATGGCAAAGGTAGCGT
TGCCAATGATGTTACAGATGAGATGGTCAGACTAAACTGGCTGACGGAATTTATGCCTCTTCCGACCATCAAGCATTTTATCCGTACTCCTGATGATGCATGGTTACTCACCACT
GCGATCCCCGGGAAAACAGCATTCCAGGTATTAGAAGAATATCCTGATTCAGGTGAAAATATTGTTGATGCGCTGGCAGTGTTCCTGCGCCGGTTGCATTCGATTCCTGTTTGTA
ATTGTCCTTTTAACAGCGATCGCGTATTTCGTCTCGCTCAGGCGCAATCACGAATGAATAACGGTTTGGTTGATGCGAGTGATTTTGATGACGAGCGTAATGGCTGGCCTGTTGA
ACAAGTCTGGAAAGAAATGCATAAGCTTTTGCCATTCTCACCGGATTCAGTCGTCACTCATGGTGATTTCTCACTTGATAACCTTATTTTTGACGAGGGGAAATTAATAGGTTGT
ATTGATGTTGGACGAGTCGGAATCGCAGACCGATACCAGGATCTTGCCATCCTATGGAACTGCCTCGGTGAGTTTTCTCCTTCATTACAGAAACGGCTTTTTCAAAAATATGGTA
TTGATAATCCTGATATGAATAAATTGCAGTTTCATTTGATGCTCGATGAGTTTTTCTAAAACACCCCTTGTATTACTGTTTATGTAAGCAGACAGTTTTATTGTTCATGATGATA
TATTTTTATCTTGTGCAATGTACATCAGAGATTTTGAGACACAA

pAJM.773

GenBank accession number: MH101730

CCAATTATTGAAGGCCTCCCTAACGGGGGGCCTTTTTTTGTTTCTGGTCTCCCGCTTAACGATCGTTGGCTGATTGGATCCAATTGACAGCTAGCTCAGTCCTAGGTACCATTGG
ATCCAATAGCTGTCACCGGATGTGCTTTCCGGTCTGATGAGTCCGTGAGGACGAAACAGCCTCTACAAATAATTTTGTTTAATACTAGAGAAAGAGGGGAAATACTAGATGGTGA
GCAAGGGCGAGGAGCTGTTCACCGGGGTGGTGCCCATCCTGGTCGAGCTGGACGGCGACGTAAACGGCCACAAGTTCAGCGTGTCCGGCGAGGGCGAGGGCGATGCCACCTACGG
CAAGCTGACCCTGAAGTTCATCTGCACCACAGGCAAGCTGCCCGTGCCCTGGCCCACCCTCGTGACCACCTTCGGCTACGGCCTGCAATGCTTCGCCCGCTACCCCGACCACATG
AAGCTGCACGACTTCTTCAAGTCCGCCATGCCCGAAGGCTACGTCCAGGAGCGCACCATCTTCTTCAAGGACGACGGCAACTACAAGACCCGCGCCGAGGTGAAGTTCGAGGGCG
ACACCCTGGTGAACCGCATCGAGCTGAAGGGCATCGACTTCAAGGAGGACGGCAACATCCTGGGGCACAAGCTGGAGTACAACTACAACAGCCACAACGTCTATATCATGGCCGA
CAAGCAGAAGAACGGCATCAAGGTGAACTTCAAGATCCGCCACAACATCGAGGACGGCAGCGTGCAGCTCGCCGACCACTACCAGCAGAACACCCCCAATCGGCGACGGCCCCGTG
CTGCTGCCCGACAACCACTACCTTAGCTACCAGTCCGCCCTGAGCAAAGACCCCAACGAGAAGCGCGATCACATGGTCCTACTGGAGTTCGTGACCGCCGCCGGGATCACTCTCG
GCATGGACGAGCTGTACAAGTAACTCGGTACCAAATTCCAGAAAAGAGGCCTCCCGAAAGGGGGGCCTTTTTTCGTTTTGGTCCAATGGCGGCGCGCCATCGAATGGTGCAAAAC
CTTTCGCGGTATGGCATGATAGCGCCCGGAAGAGAGTCAATTCAGGGGGGTGAATAATGGACATGCCTCGTATTAAACCGGGTCAGCGTGTTATGATGGCACTGCGTAAAATGAT
TGCAAGCGGTGAAATCAAAAGTGGTGAACGTATTGCAGAAATTCCGACCGCAGCAGCACTGGGTGTTAGCCGTATGCCGGTTCGTATCGCACTGCGTTCACTGGAACAAGAAGGT
CTGGTTGTTCGTCTGGGTGCACGTGGTTATGCAGCCCGTGGTGTTAGCAGCGATCAGATTCGTGATGCAATTGAAGTTCGTGGTGTTCTGGAAGGTTTTGCAGCACGTCGTCTGG
CAGAACGTGGTATGACCGCAGAAACCCATGCACGTTTTGTTGTACTGATTGCAGAAGGTGAAGCACTGTTTGCAGCCGGTCGCCTGAATGGTGAAGATCTGGATCGTTATGCCGC
ATATAATCAGGCATTTCATGATACCCTGGTTAGCGCAGCAGGTAATGGTGCAGTTGAAAGCGCACTGGCACGTAATGGTTTTGAACCGTTTGCAGCAGCCGGTGCACTGGCCCTG
GATCTGATGGACCTGTCTGCCGAATATGAACATCTGCTGGCAGCACATCGTCAGCATCAGGCAGTTCTGGATGCAGTTAGCTGTGGTGATGCCGAAGGTGCAGAACGTATTATGC
GTGATCATGCACTGGCAGCAATTCGTAATGCAAAAGTTTTTGAAGCAGCAGCAAGCGCAGGCGCACCGCTGGGTGCAGCATGGTCAATTCGTGCAGATTGATAAGGATCCTAATT
GGTAACGAATCAGACAATTGACGGCTCGAGGGAGTAGCATAGGGTTTGCAGAATCCCTGCTTCGTCCATTTGACAGGCACATTATGCATCGATGATAAGCTGTCAAACATGAGCA
GATCCTCTACGCCAGACGCATCGTGGCCGGCATCACCGGCGCCACAGGTGCGGTTGCTGGCGCCTATATCGCCGACATCACCGATGGGGAAGATCGGGCTCGCCACTTCGGGCTC
ATGAGCAAATATTTTATCTGAGGTGCTTCCTCGCTCACTGACTCGCTGCACGAGGCAGACCTCAGCGCTAGCGGAGTGTATACTGGCTTACTATGTTGGCACTGATGAGGGTGTC
AGTGAAGTGCTTCATGTGGCAGGAGAAAAAAGGCTGCACCGGTGCGTCAGCAGAATATGTGATACAGGATATATTCCGCTTCCTCGCTCACTGACTCGCTACGCTCGGTCGTTCG
ACTGCGGCGAGCGGAAATGGCTTACGAACGGGGCGGAGATTTCCTGGAAGATGCCAGGAAGATACTTAACAGGGAAGTGAGAGGGCCGCGGCAAAGCCGTTTTTCCATAGGCTCC
GCCCCCCTGACAAGCATCACGAAATCTGACGCTCAAATCAGTGGTGGCGAAACCCGACAGGACTATAAAGATACCAGGCGTTTCCCCCTGGCGGCTCCCTCGTGCGCTCTCCTGT
TCCTGCCTTTCGGTTTACCGGTGTCATTCCGCTGTTATGGCCGCGTTTGTCTCATTCCACGCCTGACACTCAGTTCCGGGTAGGCAGTTCGCTCCAAGCTGGACTGTATGCACGA
ACCCCCCGTTCAGTCCGACCGCTGCGCCTTATCCGGTAACTATCGTCTTGAGTCCAACCCGGAAAGACATGCAAAAGCACCACTGGCAGCAGCCACTGGTAATTGATTTAGAGGA
GTTAGTCTTGAAGTCATGCGCCGGTTAAGGCTAAACTGAAAGGACAAGTTTTGGTGACTGCGCTCCTCCAAGCCAGTTACCTCGGTTCAAAGAGTTGGTAGCTCAGAGAACCTTC
GAAAAACCGCCCTGCAAGGCGGTTTTTTCGTTTTCAGAGCAAGAGATTACGCGCAGACCAAAACGATCTCAAGAAGATCATCTTATTAAGGGGTCTGACGCTCAGTGGAACGAAA
AATCAATCTAAAGTATATATGAGTAAACTTGGTCTGACAGTTACCATGAGCCATATTCAACGGGAAACGTCTTGCTCGAGGCCGCGATTAAATTCCAACATGGATGCTGATTTAT
ATGGGTATAAATGGGCTCGCGATAATGTCGGGCAATCAGGTGCGACAATCTATCGATTGTATGGGAAGCCCGATGCGCCAGAGTTGTTTCTGAAACATGGCAAAGGTAGCGTTGC
CAATGATGTTACAGATGAGATGGTCAGACTAAACTGGCTGACGGAATTTATGCCTCTTCCGACCATCAAGCATTTTATCCGTACTCCTGATGATGCATGGTTACTCACCACTGCG
ATCCCCGGGAAAACAGCATTCCAGGTATTAGAAGAATATCCTGATTCAGGTGAAAATATTGTTGATGCGCTGGCAGTGTTCCTGCGCCGGTTGCATTCGATTCCTGTTTGTAATT
GTCCTTTTAACAGCGATCGCGTATTTCGTCTCGCTCAGGCGCAATCACGAATGAATAACGGTTTGGTTGATGCGAGTGATTTTGATGACGAGCGTAATGGCTGGCCTGTTGAACA
AGTCTGGAAAGAAATGCATAAGCTTTTGCCATTCTCACCGGATTCAGTCGTCACTCATGGTGATTTCTCACTTGATAACCTTATTTTTGACGAGGGGAAATTAATAGGTTGTATT
GATGTTGGACGAGTCGGAATCGCAGACCGATACCAGGATCTTGCCATCCTATGGAACTGCCTCGGTGAGTTTTCTCCTTCATTACAGAAACGGCTTTTTCAAAAATATGGTATTG
ATAATCCTGATATGAATAAATTGCAGTTTCATTTGATGCTCGATGAGTTTTTCTAAAACACCCCTTGTATTACTGTTTATGTAAGCAGACAGTTTTATTGTTCATGATGATATAT
TTTTATCTTGTGCAATGTACATCAGAGATTTTGAGACACAA

pAJM.847

GenBank

CCAATTATTGAAGGCCTCCCTAACGGGGGGCCTTTTTTTGTTTCTGGTCTCCCGCTTAACGATCGTTGGCTGCGACGTACGGTGGAATCTGATTCGTTACCAATTGACATGATAC
GAAACGTACCGTATCGTTAAGGTAGCTGTCACCGGATGTGCTTTCCGGTCTGATGAGTCCGTGAGGACGAAACAGCCTCTACAAATAATTTTGTTTAATACTAGAGAAAGAGGGG
AAATACTAGATGGTGAGCAAGGGCGAGGAGCTGTTCACCGGGGTGGTGCCCATCCTGGTCGAGCTGGACGGCGACGTAAACGGCCACAAGTTCAGCGTGTCCGGCGAGGGCGAGG
GCGATGCCACCTACGGCAAGCTGACCCTGAAGTTCATCTGCACCACAGGCAAGCTGCCCGTGCCCTGGCCCACCCTCGTGACCACCTTCGGCTACGGCCTGCAATGCTTCGCCCG

accession number: MH101727

CTACCCCGACCACATGAAGCTGCACGACTTCTTCAAGTCCGCCATGCCCGAAGGCTACGTCCAGGAGCGCACCATCTTCTTCAAGGACGACGGCAACTACAAGACCCGCGCCGAG
GTGAAGTTCGAGGGCGACACCCTGGTGAACCGCATCGAGCTGAAGGGCATCGACTTCAAGGAGGACGGCAACATCCTGGGGCACAAGCTGGAGTACAACTACAACAGCCACAACG
TCTATATCATGGCCGACAAGCAGAAGAACGGCATCAAGGTGAACTTCAAGATCCGCCACAACATCGAGGACGGCAGCGTGCAGCTCGCCGACCACTACCAGCAGAACACCCCCAAT
CGGCGACGGCCCCGTGCTGCTGCCCGACAACCACTACCTTAGCTACCAGTCCGCCCTGAGCAAAGACCCCAACGAGAAGCGCGATCACATGGTCCTGCTGGAGTTCGTGACCGCC
GCCGGGATCACTCTCGGCATGGACGAGCTGTACAAGTAACTCGGTACCAAATTCCAGAAAAGAGGCCTCCCGAAAGGGGGGCCTTTTTTCGTTTTGGTCCAATGGCGGCGCGCCA
TCGAATGGTGCAAAACCTTTCGCGGTATGGCATGATAGCGCCCGGAAGAGAGTCAATTCATGGGGGTGAATATGGCACGTACCCCGAGCCGTAGCAGCATTGGTAGCCTGCGTAG
TCCGCATACCCATAAAGCAATTCTGACCAGCACCATTGAAATCCTGAAAGAATGTGGTTATAGCGGTCTGAGCATTGAAAGCGTGGCACGTCGCGCCGGTGCAGGCAAACCGACC
ATTTATCGTTGGTGGACCAACAAAGCAGCACTGATTGCCGAAGTGTATGAAAATGAAATCGAACAGGTACGTAAATTTCCGGATTTGGGTAGCTTTAAAGCCGATCTGGATTTTC
TGCTGCATAATCTGTGGAAAGTTTGGCGTGAAACCATTTGTGGTGAAGCATTTCGTTGTGTTATTGCAGAAGCACAGTTGGACCCTGTAACCCTGACCCAACTGAAAGATCAGTT
TATGGAACGTCGTCGTGAGATACCGAAAAAACTGGTTGAAGATGCCATTAGCAATGGTGAACTGCCGAAAGATATCAATCGTGAACTGCTGCTGGATATGATTTTTGGTTTTTGT
TGGTATCGCCTGCTGACCGAACAGTTGACCGTTGAACAGGATATTGAAGAATTTACCTTCCTGCTGATTAATGGTGTTTGTCCGGGTACACAGTGTTGATAAGGATCCTAATTGG
TAACGAATCAGACAATTGACGGCTCGAGGGAGTAGCATAGGGTTTGCAGAATCCCTGCTTCGTCCATTTGACAGGCACATTATGCATCGATGATAAGCTGTCAAACATGAGCAGA
TCCTCTACGCCGGACGCATCGTGGCCGGCATCACCGGCGCCACAGGTGCGGTTGCTGGCGCCTATATCGCCGACATCACCGATGGGGAAGATCGGGCTCGCCACTTCGGGCTCAT
GAGCAAATATTTTATCTGAGGTGCTTCCTCGCTCACTGACTCGCTGCACGAGGCAGACCTCAGCGCTAGCGGAGTGTATACTGGCTTACTATGTTGGCACTGATGAGGGTGTCAG
TGAAGTGCTTCATGTGGCAGGAGAAAAAAGGCTGCACCGGTGCGTCAGCAGAATATGTGATACAGGATATATTCCGCTTCCTCGCTCACTGACTCGCTACGCTCGGTCGTTCGAC
TGCGGCGAGCGGAAATGGCTTACGAACGGGGCGGAGATTTCCTGGAAGATGCCAGGAAGATACTTAACAGGGAAGTGAGAGGGCCGCGGCAAAGCCGTTTTTCCATAGGCTCCGC
CCCCCTGACAAGCATCACGAAATCTGACGCTCAAATCAGTGGTGGCGAAACCCGACAGGACTATAAAGATACCAGGCGTTTCCCCCTGGCGGCTCCCTCGTGCGCTCTCCTGTTC
CTGCCTTTCGGTTTACCGGTGTCATTCCGCTGTTATGGCCGCGTTTGTCTCATTCCACGCCTGACACTCAGTTCCGGGTAGGCAGTTCGCTCCAAGCTGGACTGTATGCACGAAC
CCCCCGTTCAGTCCGACCGCTGCGCCTTATCCGGTAACTATCGTCTTGAGTCCAACCCGGAAAGACATGCAAAAGCACCACTGGCAGCAGCCACTGGTAATTGATTTAGAGGAGT
TAGTCTTGAAGTCATGCGCCGGTTAAGGCTAAACTGAAAGGACAAGTTTTGGTGACTGCGCTCCTCCAAGCCAGTTACCTCGGTTCAAAGAGTTGGTAGCTCAGAGAACCTTCGA
AAAACCGCCCTGCAAGGCGGTTTTTTCGTTTTCAGAGCAAGAGATTACGCGCAGACCAAAACGATCTCAAGAAGATCATCTTATTAAGGGGTCTGACGCTCAGTGGAACGAAAAA
TCAATCTAAAGTATATATGAGTAAACTTGGTCTGACAGTTACCATGAGCCATATTCAACGGGAAACGTCTTGCTCGAGGCCGCGATTAAATTCCAACATGGATGCTGATTTATAT
GGGTATAAATGGGCTCGCGATAATGTCGGGCAATCAGGTGCGACAATCTATCGATTGTATGGGAAGCCCGATGCGCCAGAGTTGTTTCTGAAACATGGCAAAGGTAGCGTTGCCA
ATGATGTTACAGATGAGATGGTCAGACTAAACTGGCTGACGGAATTTATGCCTCTTCCGACCATCAAGCATTTTATCCGTACTCCTGATGATGCATGGTTACTCACCACTGCGAT
CCCCGGGAAAACAGCATTCCAGGTATTAGAAGAATATCCTGATTCAGGTGAAAATATTGTTGATGCGCTGGCAGTGTTCCTGCGCCGGTTGCATTCGATTCCTGTTTGTAATTGT
CCTTTTAACAGCGATCGCGTATTTCGTCTCGCTCAGGCGCAATCACGAATGAATAACGGTTTGGTTGATGCGAGTGATTTTGATGACGAGCGTAATGGCTGGCCTGTTGAACAAG
TCTGGAAAGAAATGCATAAGCTTTTGCCATTCTCACCGGATTCAGTCGTCACTCATGGTGATTTCTCACTTGATAACCTTATTTTTGACGAGGGGAAATTAATAGGTTGTATTGA
TGTTGGACGAGTCGGAATCGCAGACCGATACCAGGATCTTGCCATCCTATGGAACTGCCTCGGTGAGTTTTCTCCTTCATTACAGAAACGGCTTTTTCAAAAATATGGTATTGAT
AATCCTGATATGAATAAATTGCAGTTTCATTTGATGCTCGATGAGTTTTTCTAAAACACCCCTTGTATTACTGTTTATGTAAGCAGACAGTTTTATTGTTCATGATGATATATTT
TTATCTTGTGCAATGTACATCAGAGATTTTGAGACACAA

pAJM.884

GenBank accession number: MH101739

CCAATTATTGAAGGCCTCCCTAACGGGGGGCCTTTTTTTGTTTCTGGTCTCCCGCTTAACGATCGTTGGCTGCGCTAGCAAGTAAGGCCGACGCTTCACAACCGCACTTGATTTA
ATAGACCATACCGTCTATTATTTCTGGCCATAGCTGTCACCGGATGTGCTTTCCGGTCTGATGAGTCCGTGAGGACGAAACAGCCTCTACAAATAATTTTGTTTAATACTAGAGA
AAGAGGGGAAATACTAGATGGTGAGCAAGGGCGAGGAGCTGTTCACCGGGGTGGTGCCCATCCTGGTCGAGCTGGACGGCGACGTAAACGGCCACAAGTTCAGCGTGTCCGGCGA
GGGCGAGGGCGATGCCACCTACGGCAAGCTGACCCTGAAGTTCATCTGCACCACAGGCAAGCTGCCCGTGCCCTGGCCCACCCTCGTGACCACCTTCGGCTACGGCCTGCAATGC
TTCGCCCGCTACCCCGACCACATGAAGCTGCACGACTTCTTCAAGTCCGCCATGCCCGAAGGCTACGTCCAGGAGCGCACCATCTTCTTCAAGGACGACGGCAACTACAAGACCC
GCGCCGAGGTGAAGTTCGAGGGCGACACCCTGGTGAACCGCATCGAGCTGAAGGGCATCGACTTCAAGGAGGACGGCAACATCCTGGGGCACAAGCTGGAGTACAACTACAACAG
CCACAACGTCTATATCATGGCCGACAAGCAGAAGAACGGCATCAAGGTGAACTTCAAGATCCGCCACAACATCGAGGACGGCAGCGTGCAGCTCGCCGACCACTACCAGCAGAAC
ACCCCAATCGGCGACGGCCCCGTGCTGCTGCCCGACAACCACTACCTTAGCTACCAGTCCGCCCTGAGCAAAGACCCCAACGAGAAGCGCGATCACATGGTCCTGCTGGAGTTCG
TGACCGCCGCCGGGATCACTCTCGGCATGGACGAGCTGTACAAGTAACTCGGTACCAAATTCCAGAAAAGAGGCCTCCCGAAAGGGGGGCCTTTTTTCGTTTTGGTCCAATGGCG
GCGCGCCATCGAATGGTGCAAAACCTTTCGCGGTATGGCATGATAGCGCCCGGAAGAGAGTCAATTCAGGGAGGTGAATATGCCGCTGACCGACACCCCGCCGTCTGTTCCGCAG
AAACCGCGTCGTGGTCGTCCGCGTGGTGCTCCGGACGCTTCTCTTGCTCACCAGTCTCTGATCCGTGCTGGTCTGGAACACCTGACCGAAAAAGGTTACTCTTCGGTTGGTGTTG
ACGAAATCCTGAAAGCTGCTCGTGTTCCGAAAGGTTCTTTCTACCACTACTTCCGTAACAAAGCTGACTTCGGTCTGGCTCTGATCAAAGCTTACGACACCTACTTCGCTCGTCT
CCTCGACCAGGCGTTCCTGGACGGTTCGCTGGCTCCGCTGGCTCGTCTGCGTCTGTTCACCCGTATGGCTGAAGAAGGTATGGCTCGTCACGGTTTCCGTCGTGGTTGCCTGGTT
GGTAACCTGGGTCAGGAAATGGGCGCTCTGCCGGACGACTTCCGTGCTGCTCTGATCGGTGTTCTGGAAACCTGGCAACATCGTACCGCTCAGCTGTTCCGTGAAGCTCAGGCTT
GCGGTGAACTGTCTGCTGACCATGACCCGGACGCTCTGGCTGAAGCTTTCTGGATCGGATGGGAAGGTGCTATCCTGCGTGCTAAACTGGAACTGCGTCCGGACCCGATGCACTC
TTTCACCCGTACCTTCGGTCGTCACTTCGTTACCCGTACCCAGGAATAATGATAAGGATCCTAATTGGTAACGAATCAGACAATTGACGGCTCGAGGGAGTAGCATAGGGTTTGC
AGAATCCCTGCTTCGTCCATTTGACAGGCACATTATGCATCGATGATAAGCTGTCAAACATGAGCAGATCCTCTACGCCGGACGCATCGTGGCCGGCATCACCGGCGCCACAGGT
GCGGTTGCTGGCGCCTATATCGCCGACATCACCGATGGGGAAGATCGGGCTCGCCACTTCGGGCTCATGAGCAAATATTTTATCTGAGGTGCTTCCTCGCTCACTGACTCGCTGC
ACGAGGCAGACCTCAGCGCTAGCGGAGTGTATACTGGCTTACTATGTTGGCACTGATGAGGGTGTCAGTGAAGTGCTTCATGTGGCAGGAGAAAAAAGGCTGCACCGGTGCGTCA
GCAGAATATGTGATACAGGATATATTCCGCTTCCTCGCTCACTGACTCGCTACGCTCGGTCGTTCGACTGCGGCGAGCGGAAATGGCTTACGAACGGGGCGGAGATTTCCTGGAA
GATGCCAGGAAGATACTTAACAGGGAAGTGAGAGGGCCGCGGCAAAGCCGTTTTTCCATAGGCTCCGCCCCCCTGACAAGCATCACGAAATCTGACGCTCAAATCAGTGGTGGCG
AAACCCGACAGGACTATAAAGATACCAGGCGTTTCCCCCTGGCGGCTCCCTCGTGCGCTCTCCTGTTCCTGCCTTTCGGTTTACCGGTGTCATTCCGCTGTTATGGCCGCGTTTG
TCTCATTCCACGCCTGACACTCAGTTCCGGGTAGGCAGTTCGCTCCAAGCTGGACTGTATGCACGAACCCCCCGTTCAGTCCGACCGCTGCGCCTTATCCGGTAACTATCGTCTT
GAGTCCAACCCGGAAAGACATGCAAAAGCACCACTGGCAGCAGCCACTGGTAATTGATTTAGAGGAGTTAGTCTTGAAGTCATGCGCCGGTTAAGGCTAAACTGAAAGGACAAGT
TTTGGTGACTGCGCTCCTCCAAGCCAGTTACCTCGGTTCAAAGAGTTGGTAGCTCAGAGAACCTTCGAAAAACCGCCCTGCAAGGCGGTTTTTTCGTTTTCAGAGCAAGAGATTA
CGCGCAGACCAAAACGATCTCAAGAAGATCATCTTATTAAGGGGTCTGACGCTCAGTGGAACGAAAAATCAATCTAAAGTATATATGAGTAAACTTGGTCTGACAGTTACCATGA
GCCATATTCAACGGGAAACGTCTTGCTCGAGGCCGCGATTAAATTCCAACATGGATGCTGATTTATATGGGTATAAATGGGCTCGCGATAATGTCGGGCAATCAGGTGCGACAAT
CTATCGATTGTATGGGAAGCCCGATGCGCCAGAGTTGTTTCTGAAACATGGCAAAGGTAGCGTTGCCAATGATGTTACAGATGAGATGGTCAGACTAAACTGGCTGACGGAATTT
ATGCCTCTTCCGACCATCAAGCATTTTATCCGTACTCCTGATGATGCATGGTTACTCACCACTGCGATCCCCGGGAAAACAGCATTCCAGGTATTAGAAGAATATCCTGATTCAG
GTGAAAATATTGTTGATGCGCTGGCAGTGTTCCTGCGCCGGTTGCATTCGATTCCTGTTTGTAATTGTCCTTTTAACAGCGATCGCGTATTTCGTCTCGCTCAGGCGCAATCACG
AATGAATAACGGTTTGGTTGATGCGAGTGATTTTGATGACGAGCGTAATGGCTGGCCTGTTGAACAAGTCTGGAAAGAAATGCATAAGCTTTTGCCATTCTCACCGGATTCAGTC
GTCACTCATGGTGATTTCTCACTTGATAACCTTATTTTTGACGAGGGGAAATTAATAGGTTGTATTGATGTTGGACGAGTCGGAATCGCAGACCGATACCAGGATCTTGCCATCC
TATGGAACTGCCTCGGTGAGTTTTCTCCTTCATTACAGAAACGGCTTTTTCAAAAATATGGTATTGATAATCCTGATATGAATAAATTGCAGTTTCATTTGATGCTCGATGAGTT
TTTCTAAAACACCCCTTGTATTACTGTTTATGTAAGCAGACAGTTTTATTGTTCATGATGATATATTTTTATCTTGTGCAATGTACATCAGAGATTTTGAGACACAA

pAJM.944

GenBank accession number: MH101726

CCAATTATTGAAGGCCTCCCTAACGGGGGGCCTTTTTTTGTTTCTGGTCTCCCGCTTAACGATCGTTGGCTGCCCTTTGTGCGTCCAAACGGACGCACGGCGCTCTAAAGCGGGT
CGCGATCTTTCAGATTCGCTCCTCGCGCTTTCAGTCTTTGTTTTGGCGCATGTCGTTATCGCAAAACCGCTGCACACTTTTGCGCGACATGCTCTGATCCCCCTCATCTGGGGGG
GCCTATCTGAGGGAATTTCCGATCCGGCTCGCCTGAACCATTCTGCTTTCCACGAACTTGAAAACGCTAGCTGTCACCGGATGTGCTTTCCGGTCTGATGAGTCCGTGAGGACGA
AACAGCCTCTACAAATAATTTTGTTTAATACTAGAGAAAGAGGGGAAATACTAGATGGTGAGCAAGGGCGAGGAGCTGTTCACCGGGGTGGTGCCCATCCTGGTCGAGCTGGACG
GCGACGTAAACGGCCACAAGTTCAGCGTGTCCGGCGAGGGCGAGGGCGATGCCACCTACGGCAAGCTGACCCTGAAGTTCATCTGCACCACAGGCAAGCTGCCCGTGCCCTGGCC
CACCCTCGTGACCACCTTCGGCTACGGCCTGCAATGCTTCGCCCGCTACCCCGACCACATGAAGCTGCACGACTTCTTCAAGTCCGCCATGCCCGAAGGCTACGTCCAGGAGCGC
ACCATCTTCTTCAAGGACGACGGCAACTACAAGACCCGCGCCGAGGTGAAGTTCGAGGGCGACACCCTGGTGAACCGCATCGAGCTGAAGGGCATCGACTTCAAGGAGGACGGCA
ACATCCTGGGGCACAAGCTGGAGTACAACTACAACAGCCACAACGTCTATATCATGGCCGACAAGCAGAAGAACGGCATCAAGGTGAACTTCAAGATCCGCCACAACATCGAGGA
CGGCAGCGTGCAGCTCGCCGACCACTACCAGCAGAACACCCCAATCGGCGACGGCCCCGTGCTGCTGCCCGACAACCACTACCTTAGCTACCAGTCCGCCCTGAGCAAAGACCCC
AACGAGAAGCGCGATCACATGGTCCTGCTGGAGTTCGTGACCGCCGCCGGGATCACTCTCGGCATGGACGAGCTGTACAAGTAACTCGGTACCAAATTCCAGAAAAGAGGCCTCC
CGAAAGGGGGGCCTTTTTTCGTTTTGGTCCAATGGCGGCGCGCCATCGAATGAGGTGCTTCCTCGCTCACTGACTCGCTGCACGAGGCAGACCTCAGCGCTAGCGGAGTGTATAC
TGGCTTACTATGTTGGCACTGATGAGGGTGTCAGTGAAGTGCTTCATGTGGCAGGAGAAAAAAGGCTGCACCGGTGCGTCAGCAGAATATGTGATACAGGATATATTCCGCTTCC
TCGCTCACTGACTCGCTACGCTCGGTCGTTCGACTGCGGCGAGCGGAAATGGCTTACGAACGGGGCGGAGATTTCCTGGAAGATGCCAGGAAGATACTTAACAGGGAAGTGAGAG
GGCCGCGGCAAAGCCGTTTTTCCATAGGCTCCGCCCCCCTGACAAGCATCACGAAATCTGACGCTCAAATCAGTGGTGGCGAAACCCGACAGGACTATAAAGATACCAGGCGTTT
CCCCCTGGCGGCTCCCTCGTGCGCTCTCCTGTTCCTGCCTTTCGGTTTACCGGTGTCATTCCGCTGTTATGGCCGCGTTTGTCTCATTCCACGCCTGACACTCAGTTCCGGGTAG
GCAGTTCGCTCCAAGCTGGACTGTATGCACGAACCCCCCGTTCAGTCCGACCGCTGCGCCTTATCCGGTAACTATCGTCTTGAGTCCAACCCGGAAAGACATGCAAAAGCACCAC
TGGCAGCAGCCACTGGTAATTGATTTAGAGGAGTTAGTCTTGAAGTCATGCGCCGGTTAAGGCTAAACTGAAAGGACAAGTTTTGGTGACTGCGCTCCTCCAAGCCAGTTACCTC
GGTTCAAAGAGTTGGTAGCTCAGAGAACCTTCGAAAAACCGCCCTGCAAGGCGGTTTTTTCGTTTTCAGAGCAAGAGATTACGCGCAGACCAAAACGATCTCAAGAAGATCATCT
TATTAAGGGGTCTGACGCTCAGTGGAACGAAAAATCAATCTAAAGTATATATGAGTAAACTTGGTCTGACAGTTACCATGAGCCATATTCAACGGGAAACGTCTTGCTCGAGGCC
GCGATTAAATTCCAACATGGATGCTGATTTATATGGGTATAAATGGGCTCGCGATAATGTCGGGCAATCAGGTGCGACAATCTATCGATTGTATGGGAAGCCCGATGCGCCAGAG
TTGTTTCTGAAACATGGCAAAGGTAGCGTTGCCAATGATGTTACAGATGAGATGGTCAGACTAAACTGGCTGACGGAATTTATGCCTCTTCCGACCATCAAGCATTTTATCCGTA
CTCCTGATGATGCATGGTTACTCACCACTGCGATCCCCGGGAAAACAGCATTCCAGGTATTAGAAGAATATCCTGATTCAGGTGAAAATATTGTTGATGCGCTGGCAGTGTTCCT
GCGCCGGTTGCATTCGATTCCTGTTTGTAATTGTCCTTTTAACAGCGATCGCGTATTTCGTCTCGCTCAGGCGCAATCACGAATGAATAACGGTTTGGTTGATGCGAGTGATTTT
GATGACGAGCGTAATGGCTGGCCTGTTGAACAAGTCTGGAAAGAAATGCATAAGCTTTTGCCATTCTCACCGGATTCAGTCGTCACTCATGGTGATTTCTCACTTGATAACCTTA
TTTTTGACGAGGGGAAATTAATAGGTTGTATTGATGTTGGACGAGTCGGAATCGCAGACCGATACCAGGATCTTGCCATCCTATGGAACTGCCTCGGTGAGTTTTCTCCTTCATT
ACAGAAACGGCTTTTTCAAAAATATGGTATTGATAATCCTGATATGAATAAATTGCAGTTTCATTTGATGCTCGATGAGTTTTTCTAAAACACCCCTTGTATTACTGTTTATGTA
AGCAGACAGTTTTATTGTTCATGATGATATATTTTTATCTTGTGCAATGTACATCAGAGATTTTGAGACACAA

pAJM.969

GenBank accession number: MH101740

CCAATTATTGAAGGCCTCCCTAACGGGGGGCCTTTTTTTGTTTCTGGTCTCCCGCTTAACGATCGTTGGCTGGGATTGAATATAACCGACGTGACTGTTACATTTAGGTGGCTAA
ACCCGTCAAAGCTGTCACCGGATGTGCTTTCCGGTCTGATGAGTCCGTGAGGACGAAACAGCCTCTACAAATAATTTTGTTTAATACTAGAGAAAGAGGGGAAATACTAGATGGT
GAGCAAGGGCGAGGAGCTGTTCACCGGGGTGGTGCCCATCCTGGTCGAGCTGGACGGCGACGTAAACGGCCACAAGTTCAGCGTGTCCGGCGAGGGCGAGGGCGATGCCACCTAC
GGCAAGCTGACCCTGAAGTTCATCTGCACCACAGGCAAGCTGCCCGTGCCCTGGCCCACCCTCGTGACCACCTTCGGCTACGGCCTGCAATGCTTCGCCCGCTACCCCGACCACA
TGAAGCTGCACGACTTCTTCAAGTCCGCCATGCCCGAAGGCTACGTCCAGGAGCGCACCATCTTCTTCAAGGACGACGGCAACTACAAGACCCGCGCCGAGGTGAAGTTCGAGGG
CGACACCCTGGTGAACCGCATCGAGCTGAAGGGCATCGACTTCAAGGAGGACGGCAACATCCTGGGGCACAAGCTGGAGTACAACTACAACAGCCACAACGTCTATATCATGGCC
GACAAGCAGAAGAACGGCATCAAGGTGAACTTCAAGATCCGCCACAACATCGAGGACGGCAGCGTGCAGCTCGCCGACCACTACCAGCAGAACACCCCAATCGGCGACGGCCCCG
TGCTGCTGCCCGACAACCACTACCTTAGCTACCAGTCCGCCCTGAGCAAAGACCCCAACGAGAAGCGCGATCACATGGTCCTGCTGGAGTTCGTGACCGCCGCCGGGATCACTCT
CGGCATGGACGAGCTGTACAAGTAACTCGGTACCAAATTCCAGAAAAGAGGCCTCCCGAAAGGGGGGCCTTTTTTCGTTTTGGTCCAATGGCGGCGCGCCATCGAATGGTGCAAA
ACCTTTCGCGGTATGGCATGATAGCGCCCGGAAGAGAGTCAATTTAGGGAGGTGAATATGCCTCGTCCGAAACTGAAAAGTGATGATGAAGTTCTGGAAGCAGCAACCGTTGTTC
TGAAACGTTGTGGTCCGATTGAATTTACCCTGAGCGGTGTTGCAAAAGAAGTTGGTCTGAGTCGCGCAGCACTGATTCAGCGTTTTACCAATCGTGATACCCTGCTGGTTCGTAT
GATGGAACGTGGTGTTGAACAGGTTCGTCATTATCTGAATGCAATTCCGATTGGTGCAGGTCCGCAGGGTCTGTGGGAATTTCTGCAGGTTCTGGTTCGTAGCATGGATACCCGT
AATGATTTCAGCGTGAACTATCTGATCAGCTGGTATGAACTGCAGGTTCCGGAACTGCGTACCCTGGCAATTCAGCGTAATCGTGCAGTTGTTGAAGGTATTCGTAAACGTCTGC
CTCCGGGTGCACCGGCAGCAGCAGAACTGCTGCTGCATAGCGTTATTGCCGGTGCAACCATGCAGTGGGCAGTTGATCCGGATGGTGAACTGGCAGATCATGTTCTGGCACAGAT
TGCAGCAATTCTGTGTCTGATGTTTCCGGAACATGATGATTTTCAGCTGCTGCAGGCACATGCATAATAATATTGAAAAAGGAAGAGTATGAACGAGAAAAATATAAAACACAGT
CAAAACTTTATTACTTCAAAACATAATATAGATAAAATAATGACAAATATAAGATTAAATGAACATGATAATATCTTTGAAATCGGCTCAGGAAAAGGCCATTTTACCCTTGAAT
TAGTAAAGAGGTGTAATTTCGTAACTGCCATTGAAATAGACCATAAATTATGCAAAACTACAGAAAATAAACTTGTTGATCACGATAATTTCCAAGTTTTAAACAAGGATATATT
GCAGTTTAAATTTCCTAAAAACCAATCCTATAAAATATATGGTAATATACCTTATAACATAAGTACGGATATAATACGCAAAATTGTTTTTGATAGTATAGCTAATGAGATTTAT
TTAATCGTGGAATACGGGTTTGCTAAAAGATTATTAAATACAAAACGCTCATTGGCATTACTTTTAATGGCAGAAGTTGATATTTCTATATTAAGTATGGTTCCAAGAGAATATT
TTCATCCTAAACCTAAAGTGAATAGCTCACTTATCAGATTAAGTAGAAAAAAATCAAGAATATCACACAAAGATAAACAAAAGTATAATTATTTCGTTATGAAATGGGTTAACAA
AGAATACAAGAAAATATTTACAAAAAATCAATTTAACAATTCCTTAAAACATGCAGGAATTGACGATTTAAACAATATTAGCTTTGAACAATTCTTATCTCTTTTCAATAGCTAT
AAATTATTTAATAAGTGATAAGGATCCTAATTGGTAACGAATCAGACAATTGACGGCTCGAGGGAGTAGCATAGGGTTTGCAGAATCCCTGCTTCGTCCATTTGACAGGCACATT
ATGCATCGATGATAAGCTGTCAAACATGAGCAGATCCTCTACGCCGGACGCATCGTGGCCGGCATCACCGGCGCCACAGGTGCGGTTGCTGGCGCCTATATCGCCGACATCACCG
ATGGGGAAGATCGGGCTCGCCACTTCGGGCTCATGAGCAAATATTTTATCTGAGGTGCTTCCTCGCTCACTGACTCGCTGCACGAGGCAGACCTCAGCGCTAGCGGAGTGTATAC
TGGCTTACTATGTTGGCACTGATGAGGGTGTCAGTGAAGTGCTTCATGTGGCAGGAGAAAAAAGGCTGCACCGGTGCGTCAGCAGAATATGTGATACAGGATATATTCCGCTTCC
TCGCTCACTGACTCGCTACGCTCGGTCGTTCGACTGCGGCGAGCGGAAATGGCTTACGAACGGGGCGGAGATTTCCTGGAAGATGCCAGGAAGATACTTAACAGGGAAGTGAGAG
GGCCGCGGCAAAGCCGTTTTTCCATAGGCTCCGCCCCCCTGACAAGCATCACGAAATCTGACGCTCAAATCAGTGGTGGCGAAACCCGACAGGACTATAAAGATACCAGGCGTTT
CCCCCTGGCGGCTCCCTCGTGCGCTCTCCTGTTCCTGCCTTTCGGTTTACCGGTGTCATTCCGCTGTTATGGCCGCGTTTGTCTCATTCCACGCCTGACACTCAGTTCCGGGTAG
GCAGTTCGCTCCAAGCTGGACTGTATGCACGAACCCCCCGTTCAGTCCGACCGCTGCGCCTTATCCGGTAACTATCGTCTTGAGTCCAACCCGGAAAGACATGCAAAAGCACCAC
TGGCAGCAGCCACTGGTAATTGATTTAGAGGAGTTAGTCTTGAAGTCATGCGCCGGTTAAGGCTAAACTGAAAGGACAAGTTTTGGTGACTGCGCTCCTCCAAGCCAGTTACCTC
GGTTCAAAGAGTTGGTAGCTCAGAGAACCTTCGAAAAACCGCCCTGCAAGGCGGTTTTTTCGTTTTCAGAGCAAGAGATTACGCGCAGACCAAAACGATCTCAAGAAGATCATCT
TATTAAGGGGTCTGACGCTCAGTGGAACGAAAAATCAATCTAAAGTATATATGAGTAAACTTGGTCTGACAGTTACCATGAGCCATATTCAACGGGAAACGTCTTGCTCGAGGCC
GCGATTAAATTCCAACATGGATGCTGATTTATATGGGTATAAATGGGCTCGCGATAATGTCGGGCAATCAGGTGCGACAATCTATCGATTGTATGGGAAGCCCGATGCGCCAGAG
TTGTTTCTGAAACATGGCAAAGGTAGCGTTGCCAATGATGTTACAGATGAGATGGTCAGACTAAACTGGCTGACGGAATTTATGCCTCTTCCGACCATCAAGCATTTTATCCGTA
CTCCTGATGATGCATGGTTACTCACCACTGCGATCCCCGGGAAAACAGCATTCCAGGTATTAGAAGAATATCCTGATTCAGGTGAAAATATTGTTGATGCGCTGGCAGTGTTCCT
GCGCCGGTTGCATTCGATTCCTGTTTGTAATTGTCCTTTTAACAGCGATCGCGTATTTCGTCTCGCTCAGGCGCAATCACGAATGAATAACGGTTTGGTTGATGCGAGTGATTTT
GATGACGAGCGTAATGGCTGGCCTGTTGAACAAGTCTGGAAAGAAATGCATAAGCTTTTGCCATTCTCACCGGATTCAGTCGTCACTCATGGTGATTTCTCACTTGATAACCTTA
TTTTTGACGAGGGGAAATTAATAGGTTGTATTGATGTTGGACGAGTCGGAATCGCAGACCGATACCAGGATCTTGCCATCCTATGGAACTGCCTCGGTGAGTTTTCTCCTTCATT
ACAGAAACGGCTTTTTCAAAAATATGGTATTGATAATCCTGATATGAATAAATTGCAGTTTCATTTGATGCTCGATGAGTTTTTCTAAAACACCCCTTGTATTACTGTTTATGTA
AGCAGACAGTTTTATTGTTCATGATGATATATTTTTATCTTGTGCAATGTACATCAGAGATTTTGAGACACAA

pAJM.1459

GenBank accession number: MH101724

CCAATTATTGAAGGCCTCCCTAACGGGGGGCCTTTTTTTGTTTCTGGTCTCCCGCTTAACGATCGTTGGCTGTTTTGTTCGATTATCGAACAAATTATTGAAATATCGAACAAAA
CCTCTAAACTACTGTGGCACTGAATCAAAAAATTATAAACCCTGATCAGAAGCTGTCACCGGATGTGCTTTCCGGTCTGATGAGTCCGTGAGGACGAAACAGCCTCTACAAATAA
TTTTGTTTAATACTAGAGAAAGAGGGGAAATACTAGATGGTGAGCAAGGGCGAGGAGCTGTTCACCGGGGTGGTGCCCATCCTGGTCGAGCTGGACGGCGACGTAAACGGCCACA
AGTTCAGCGTGTCCGGCGAGGGCGAGGGCGATGCCACCTACGGCAAGCTGACCCTGAAGTTCATCTGCACCACAGGCAAGCTGCCCGTGCCCTGGCCCACCCTCGTGACCACCTT
CGGCTACGGCCTGCAATGCTTCGCCCGCTACCCCGACCACATGAAGCTGCACGACTTCTTCAAGTCCGCCATGCCCGAAGGCTACGTCCAGGAGCGCACCATCTTCTTCAAGGAC
GACGGCAACTACAAGACCCGCGCCGAGGTGAAGTTCGAGGGCGACACCCTGGTGAACCGCATCGAGCTGAAGGGCATCGACTTCAAGGAGGACGGCAACATCCTGGGGCACAAGC
TGGAGTACAACTACAACAGCCACAACGTCTATATCATGGCCGACAAGCAGAAGAACGGCATCAAGGTGAACTTCAAGATCCGCCACAACATCGAGGACGGCAGCGTGCAGCTCGC
CGACCACTACCAGCAGAACACCCCAATCGGCGACGGCCCCGTGCTGCTGCCCGACAACCACTACCTTAGCTACCAGTCCGCCCTGAGCAAAGACCCCAACGAGAAGCGCGATCAC
ATGGTCCTGCTGGAGTTCGTGACCGCCGCCGGGATCACTCTCGGCATGGACGAGCTGTACAAGTAACTCGGTACCAAATTCCAGAAAAGAGGCCTCCCGAAAGGGGGGCCTTTTT
TCGTTTTGGTCCAATGGCGGCGCGCCATCGAATGAGGTGCTTCCTCGCTCACTGACTCGCTGCACGAGGCAGACCTCAGCGCTAGCGGAGTGTATACTGGCTTACTATGTTGGCA
CTGATGAGGGTGTCAGTGAAGTGCTTCATGTGGCAGGAGAAAAAAGGCTGCACCGGTGCGTCAGCAGAATATGTGATACAGGATATATTCCGCTTCCTCGCTCACTGACTCGCTA
CGCTCGGTCGTTCGACTGCGGCGAGCGGAAATGGCTTACGAACGGGGCGGAGATTTCATGGAAGATGCCAGGAAGATACTTAACAGGGAAGTGAGAGGGCCGCGGCAAAGCCGTT
TTTCCATAGGCTCCGCCCCCCTGACAAGCATCACGAAATCTGACGCTCAAATCAGTGGTGGCGAAACCCGACAGGACTATAAAGATACCAGGCGTTTCCCCCTGGCGGCTCCCTC
GTGCGCTCTCCTGTTCCTGCCTTTCGGTTTACCGGTGTCATTCCGCTGTTATGGCCGCGTTTGTCTCATTCCACGCCTGACACTCAGTTCCGGGTAGGCAGTTCGCTCCAAGCTG
GACTGTATGCACGAACCCCCCGTTCAGTCCGACCGCTGCGCCTTATCCGGTAACTATCGTCTTGAGTCCAACCCGGAAAGACATGCAAAAGCACCACTGGCAGCAGCCACTGGTA
ATTGATTTAGAGGAGTTAGTCTTGAAGTCATGCGCCGGTTAAGGCTAAACTGAAAGGACAAGTTTTGGTGACTGCGCTCCTCCAAGCCAGTTACCTCGGTTCAAAGAGTTGGTAG
CTCAGAGAACCTTCGAAAAACCGCCCTGCAAGGCGGTTTTTTCGTTTTCAGAGCAAGAGATTACGCGCAGACCAAAACGATCTCAAGAAGATCATCTTATTAAGGGGTCTGACGC
TCAGTGGAACGAAAAATCAATCTAAAGTATATATGAGTAAACTTGGTCTGACAGTTACCATGAGCCATATTCAACGGGAAACGTCTTGCTCGAGGCCGCGATTAAATTCCAACAT
GGATGCTGATTTATATGGGTATAAATGGGCTCGCGATAATGTCGGGCAATCAGGTGCGACAATCTATCGATTGTATGGGAAGCCCGATGCGCCAGAGTTGTTTCTGAAACATGGC
AAAGGTAGCGTTGCCAATGATGTTACAGATGAGATGGTCAGACTAAACTGGCTGACGGAATTTATGCCTCTTCCGACCATCAAGCATTTTATCCGTACTCCTGATGATGCATGGT
TACTCACCACTGCGATCCCCGGGAAAACAGCATTCCAGGTATTAGAAGAATATCCTGATTCAGGTGAAAATATTGTTGATGCGCTGGCAGTGTTCCTGCGCCGGTTGCATTCGAT
TCCTGTTTGTAATTGTCCTTTTAACAGCGATCGCGTATTTCGTCTCGCTCAGGCGCAATCACGAATGAATAACGGTTTGGTTGATGCGAGTGATTTTGATGACGAGCGTAATGGC
TGGCCTGTTGAACAAGTCTGGAAAGAAATGCATAAGCTTTTGCCATTCTCACCGGATTCAGTCGTCACTCATGGTGATTTCTCACTTGATAACCTTATTTTTGACGAGGGGAAAT
TAATAGGTTGTATTGATGTTGGACGAGTCGGAATCGCAGACCGATACCAGGATCTTGCCATCCTATGGAACTGCCTCGGTGAGTTTTCTCCTTCATTACAGAAACGGCTTTTTCA
AAAATATGGTATTGATAATCCTGATATGAATAAATTGCAGTTTCATTTGATGCTCGATGAGTTTTTCTAAAACACCCCTTGTATTACTGTTTATGTAAGCAGACAGTTTTATTGT
TCATGATGATATATTTTTATCTTGTGCAATGTACATCAGAGATTTTGAGACACAA

pAJM.1642

GenBank accession number: MH101738

CCAATTATTGAAGGCCTCCCTAACGGGGGGCCTTTTTTTGTTTCTGGTCTCCCGCTTAACGATCGTTGGCTGCCCTTTGTGCGTCCAAACGGACGCACGGCGCTCTAAAGCGGGT
CGCGATCTTTCAGATTCGCTCCTCGCGCTTTCAGTCTTTGTTTTGGCGCATGTCGTTATCGCAAAACCGCTGCACACTTTTGCGCGACATGCTCTGATCCCCCTCATCTGGGGGG
GCCTATCTGAGGGAATTTCCGATCCGGCTCGCCTGAACCATTCTGCTTTCCACGAACTTGAAAACGCTAGCTGTCACCGGATGTGCTTTCCGGTCTGATGAGTCCGTGAGGACGA
AACAGCCTCTACAAATAATTTTGTTTAATACTAGAGAAAGAGGGGAAATACTAGATGGTGAGCAAGGGCGAGGAGCTGTTCACCGGGGTGGTGCCCATCCTGGTCGAGCTGGACG
GCGACGTAAACGGCCACAAGTTCAGCGTGTCCGGCGAGGGCGAGGGCGATGCCACCTACGGCAAGCTGACCCTGAAGTTCATCTGCACCACAGGCAAGCTGCCCGTGCCCTGGCC
CACCCTCGTGACCACCTTCGGCTACGGCCTGCAATGCTTCGCCCGCTACCCCGACCACATGAAGCTGCACGACTTCTTCAAGTCCGCCATGCCCGAAGGCTACGTCCAGGAGCGC
ACCATCTTCTTCAAGGACGACGGCAACTACAAGACCCGCGCCGAGGTGAAGTTCGAGGGCGACACCCTGGTGAACCGCATCGAGCTGAAGGGCATCGACTTCAAGGAGGACGGCA
ACATCCTGGGGCACAAGCTGGAGTACAACTACAACAGCCACAACGTCTATATCATGGCCGACAAGCAGAAGAACGGCATCAAGGTGAACTTCAAGATCCGCCACAACATCGAGGA
CGGCAGCGTGCAGCTCGCCGACCACTACCAGCAGAACACCCCAATCGGCGACGGCCCCGTGCTGCTGCCCGACAACCACTACCTTAGCTACCAGTCCGCCCTGAGCAAAGACCCC
AACGAGAAGCGCGATCACATGGTCCTGCTGGAGTTCGTGACCGCCGCCGGGATCACTCTCGGCATGGACGAGCTGTACAAGTAACTCGGTACCAAATTCCAGAAAAGAGGCCTCC
CGAAAGGGGGGCCTTTTTTCGTTTTGGTCCAATGGCGGCGCGCCATCGAATGGCGCAAAACCTTTCGCGGTATGGCATGATAGCGCCCGGAAGAGAGTCAATTCAGGGTGGTGAA
TATGATTGAGAATACCTATAGCGAAAAGTTCGAGTCCGCGTTCGAACAGATCAAAGCGGCGGCCAACGTGGATGCCGCCATCCGTATTCTCCAGGCGGAATATAACCTCGATTTC
GTCACCTACCATCTCGCCCAGACAATCGCGAGCAAGATCGATTCGCCCTTCGTGCGCACCACCTATCCGGATGCCTGGGTTTCCCGTTACCTCCTCAACTGCTATGTGAAGGTCG
ATCCGATCATCAAGCAGGGCTTCGAACGCCAGCTGCCCTTCGACTGGAGCGAGGTCGAACCGACGCCGGAGGCCTATGCCATGCTGGTCGACGCCCAGAAACACGGCATCGATGA
CAATGGCTACTCCATCCCCGTCGCCGACAAGGCGCAGCGCCGCGCCCTGCTGTCGCTGAATGCCCATATACCGGCCGACGAATGGACCGAGCTCGTGCGCCGCTGCCGCAATGAG
TGGATCGAGATCGCCCATCTGATCCACCGCAAGGCCGTATATGAGCTGCATGGCGAAAACGATCCGGTGCCGGCATTGTCGCCGCGCGAGATCGAGTGTCTGCACTGGACCGCCC
TCGGCAAGGATTACAAGGATATTTCGGTCATCCTGGGCATATCAGAGCATACCACACGCGATTACCTGAAAACCGCCCGCTTCAGGCTCGGCTGCACCACGATCTCGGCCGCCGC
GTCGCGGGCTGTTCAATTGCGCATCATCAATCCCTATAGGATCCGCATGACGCGACGTAATTGGTAATAGCCTAATTGGTAACGAATCAGACAATTGACGGCTCGAGGGAGTAGC
ATAGGGTTTGCAGAATCCCTGCTTCGTCCATTTGACAGGCACATTATGCATCGATGATAAGCTGTCAAACATGAGCAGATCCTCTACGCCGGACGCATCGTGGCCGGCATCACCG
GCGCCACAGGTGCGGTTGCTGGCGCCTATATCGCCGACATCACCGATGGGGAAGATCGGGCTCGCCACTTCGGGCTCATGAGCAAATATTTTATCTGAGGTGCTTCCTCGCTCAC
TGACTCGCTGCACGAGGCAGACCTCAGCGCTAGCGGAGTGTATACTGGCTTACTATGTTGGCACTGATGAGGGTGTCAGTGAAGTGCTTCATGTGGCAGGAGAAAAAAGGCTGCA
CCGGTGCGTCAGCAGAATATGTGATACAGGATATATTCCGCTTCCTCGCTCACTGACTCGCTACGCTCGGTCGTTCGACTGCGGCGAGCGGAAATGGCTTACGAACGGGGCGGAG
ATTTCCTGGAAGATGCCAGGAAGATACTTAACAGGGAAGTGAGAGGGCCGCGGCAAAGCCGTTTTTCCATAGGCTCCGCCCCCCTGACAAGCATCACGAAATCTGACGCTCAAAT
CAGTGGTGGCGAAACCCGACAGGACTATAAAGATACCAGGCGTTTCCCCCTGGCGGCTCCCTCGTGCGCTCTCCTGTTCCTGCCTTTCGGTTTACCGGTGTCATTCCGCTGTTAT
GGCCGCGTTTGTCTCATTCCACGCCTGACACTCAGTTCCGGGTAGGCAGTTCGCTCCAAGCTGGACTGTATGCACGAACCCCCCGTTCAGTCCGACCGCTGCGCCTTATCCGGTA
ACTATCGTCTTGAGTCCAACCCGGAAAGACATGCAAAAGCACCACTGGCAGCAGCCACTGGTAATTGATTTAGAGGAGTTAGTCTTGAAGTCATGCGCCGGTTAAGGCTAAACTG

AAAGGACAAGTTTTGGTGACTGCGCTCCTCCAAGCCAGTTACCTCGGTTCAAAGAGTTGGTAGCTCAGAGAACCTTCGAAAAACCGCCCTGCAAGGCGGTTTTTTCGTTTTCAGA
GCAAGAGATTACGCGCAGACCAAAACGATCTCAAGAAGATCATCTTATTAAGGGGTCTGACGCTCAGTGGAACGAAAAATCAATCTAAAGTATATATGAGTAAACTTGGTCTGAC
AGTTACCATGAGCCATATTCAACGGGAAACGTCTTGCTCGAGGCCGCGATTAAATTCCAACATGGATGCTGATTTATATGGGTATAAATGGGCTCGCGATAATGTCGGGCAATCA
GGTGCGACAATCTATCGATTGTATGGGAAGCCCGATGCGCCAGAGTTGTTTCTGAAACATGGCAAAGGTAGCGTTGCCAATGATGTTACAGATGAGATGGTCAGACTAAACTGGC
TGACGGAATTTATGCCTCTTCCGACCATCAAGCATTTTATCCGTACTCCTGATGATGCATGGTTACTCACCACTGCGATCCCCGGGAAAACAGCATTCCAGGTATTAGAAGAATA
TCCTGATTCAGGTGAAAATATTGTTGATGCGCTGGCAGTGTTCCTGCGCCGGTTGCATTCGATTCCTGTTTGTAATTGTCCTTTTAACAGCGATCGCGTATTTCGTCTCGCTCAG
GCGCAATCACGAATGAATAACGGTTTGGTTGATGCGAGTGATTTTGATGACGAGCGTAATGGCTGGCCTGTTGAACAAGTCTGGAAAGAAATGCATAAGCTTTTGCCATTCTCAC
CGGATTCAGTCGTCACTCATGGTGATTTCTCACTTGATAACCTTATTTTTGACGAGGGGAAATTAATAGGTTGTATTGATGTTGGACGAGTCGGAATCGCAGACCGATACCAGGA
TCTTGCCATCCTATGGAACTGCCTCGGTGAGTTTTCTCCTTCATTACAGAAACGGCTTTTTCAAAAATATGGTATTGATAATCCTGATATGAATAAATTGCAGTTTCATTTGATG
CTCGATGAGTTTTTCTAAAACACCCCTTGTATTACTGTTTATGTAAGCAGACAGTTTTATTGTTCATGATGATATATTTTTATCTTGTGCAATGTACATCAGAGATTTTGAGACA
CAA

RPU standard

CCAATTATTGAAGGCCTCCCTAACGGGGGGCCTTTTTTTGTTTCTGGTCTCCCGCTTGATAAGTCCCTAACTTTTACAGCTAGCTCAGTCCTAGGTATTATGCTAGCCTGAAGCT
GTCACCGGATGTGCTTTCCGGTCTGATGAGTCCGTGAGGACGAAACAGCCTCTACAAATAATTTTGTTTAATACTAGAGAAAGAGGGGAAATACTAGATGGTGAGCAAGGGCGAG
GAGCTGTTCACCGGGGTGGTGCCCATCCTGGTCGAGCTGGACGGCGACGTAAACGGCCACAAGTTCAGCGTGTCCGGCGAGGGCGAGGGCGATGCCACCTACGGCAAGCTGACCC
TGAAGTTCATCTGCACCACAGGCAAGCTGCCCGTGCCCTGGCCCACCCTCGTGACCACCTTCGGCTACGGCCTGCAATGCTTCGCCCGCTACCCCGACCACATGAAGCTGCACGA
CTTCTTCAAGTCCGCCATGCCCGAAGGCTACGTCCAGGAGCGCACCATCTTCTTCAAGGACGACGGCAACTACAAGACCCGCGCCGAGGTGAAGTTCGAGGGCGACACCCTGGTG
AACCGCATCGAGCTGAAGGGCATCGACTTCAAGGAGGACGGCAACATCCTGGGGCACAAGCTGGAGTACAACTACAACAGCCACAACGTCTATATCATGGCCGACAAGCAGAAGA
ACGGCATCAAGGTGAACTTCAAGATCCGCCACAACATCGAGGACGGCAGCGTGCAGCTCGCCGACCACTACCAGCAGAACACCCCAATCGGCGACGGCCCCGTGCTGCTGCCCGA
CAACCACTACCTTAGCTACCAGTCCGCCCTGAGCAAAGACCCCAACGAGAAGCGCGATCACATGGTCCTGCTGGAGTTCGTGACCGCCGCCGGGATCACTCTCGGCATGGACGAG
CTGTACAAGTAACTCGGTACCAAATTCCAGAAAAGAGGCCTCCCGAAAGGGGGGCCTTTTTTCGTTTTGGTCCGATCCTCTACGCCGGACGCATCGTGGCCGGCATCACCGGCGC
CACAGGTGCGGTTGCTGGCGCCTATATCGCCGACATCACCGATGGGGAAGATCGGGCTCGCCACTTCGGGCTCATGAGCAAATATTTTATCTGAGGTGCTTCCTCGCTCACTGAC
TCGCTGCACGAGGCAGACCTCAGCGCTAGCGGAGTGTATACTGGCTTACTATGTTGGCACTGATGAGGGTGTCAGTGAAGTGCTTCATGTGGCAGGAGAAAAAAGGCTGCACCGG
TGCGTCAGCAGAATATGTGATACAGGATATATTCCGCTTCCTCGCTCACTGACTCGCTACGCTCGGTCGTTCGACTGCGGCGAGCGGAAATGGCTTACGAACGGGGCGGAGATTT
CATGGAAGATGCCAGGAAGATACTTAACAGGGAAGTGAGAGGGCCGCGGCAAAGCCGTTTTTCCATAGGCTCCGCCCCCCTGACAAGCATCACGAAATCTGACGCTCAAATCAGT
GGTGGCGAAACCCGACAGGACTATAAAGATACCAGGCGTTTCCCCCTGGCGGCTCCCTCGTGCGCTCTCCTGTTCCTGCCTTTCGGTTTACCGGTGTCATTCCGCTGTTATGGCC
GCGTTTGTCTCATTCCACGCCTGACACTCAGTTCCGGGTAGGCAGTTCGCTCCAAGCTGGACTGTATGCACGAACCCCCCGTTCAGTCCGACCGCTGCGCCTTATCCGGTAACTA
TCGTCTTGAGTCCAACCCGGAAAGACATGCAAAAGCACCACTGGCAGCAGCCACTGGTAATTGATTTAGAGGAGTTAGTCTTGAAGTCATGCGCCGGTTAAGGCTAAACTGAAAG
GACAAGTTTTGGTGACTGCGCTCCTCCAAGCCAGTTACCTCGGTTCAAAGAGTTGGTAGCTCAGAGAACCTTCGAAAAACCGCCCTGCAAGGCGGTTTTTTCGTTTTCAGAGCAA
GAGATTACGCGCAGACCAAAACGATCTCAAGAAGATCATCTTATTAAGGGGTCTGACGCTCAGTGGAACGAAAAATCAATCTAAAGTATATATGAGTAAACTTGGTCTGACAGTT
ACCATGAGCCATATTCAACGGGAAACGTCTTGCTCGAGGCCGCGATTAAATTCCAACATGGATGCTGATTTATATGGGTATAAATGGGCTCGCGATAATGTCGGGCAATCAGGTG
CGACAATCTATCGATTGTATGGGAAGCCCGATGCGCCAGAGTTGTTTCTGAAACATGGCAAAGGTAGCGTTGCCAATGATGTTACAGATGAGATGGTCAGACTAAACTGGCTGAC
GGAATTTATGCCTCTTCCGACCATCAAGCATTTTATCCGTACTCCTGATGATGCATGGTTACTCACCACTGCGATCCCCGGGAAAACAGCATTCCAGGTATTAGAAGAATATCCT
GATTCAGGTGAAAATATTGTTGATGCGCTGGCAGTGTTCCTGCGCCGGTTGCATTCGATTCCTGTTTGTAATTGTCCTTTTAACAGCGATCGCGTATTTCGTCTCGCTCAGGCGC
AATCACGAATGAATAACGGTTTGGTTGATGCGAGTGATTTTGATGACGAGCGTAATGGCTGGCCTGTTGAACAAGTCTGGAAAGAAATGCATAAGCTTTTGCCATTCTCACCGGA
TTCAGTCGTCACTCATGGTGATTTCTCACTTGATAACCTTATTTTTGACGAGGGGAAATTAATAGGTTGTATTGATGTTGGACGAGTCGGAATCGCAGACCGATACCAGGATCTT
GCCATCCTATGGAACTGCCTCGGTGAGTTTTCTCCTTCATTACAGAAACGGCTTTTTCAAAAATATGGTATTGATAATCCTGATATGAATAAATTGCAGTTTCATTTGATGCTCG
ATGAGTTTTTCTAAAACACCCCTTGTATTACTGTTTATGTAAGCAGACAGTTTTATTGTTCATGATGATATATTTTTATCTTGTGCAATGTACATCAGAGATTTTGAGACACAA

a) DNA sequence colors correspond to promoters (orange), ribozyme insulators (blue), RBSs (green), protein coding sequences (red), terminators (purple), and plasmid backbone and spacers (black).
